# Structure Activity of β-Amidomethyl Vinyl Sulfones as Covalent Inhibitors of *Chikungunya* nsP2 Cysteine Protease with Anti-alphavirus Activity

**DOI:** 10.1101/2024.06.12.598722

**Authors:** Anirban Ghoshal, Kesatebrhan Haile Asressu, Mohammad Anwar Hossain, Peter J. Brown, Eric M. Merten, John D. Sears, Sumera Perveen, Kenneth H. Pearce, Konstantin I. Popov, Nathaniel J. Moorman, Mark T. Heise, Timothy M. Willson

## Abstract

Despite their widespread impact on human health there are no approved drugs for combating alphavirus infections. The heterocyclic β-aminomethyl vinyl sulfone RA-0002034 (**1a**) is a potent irreversible covalent inhibitor of the alphavirus nsP2 cysteine protease with broad spectrum antiviral activity. Analogs of **1a** that varied each of three regions of the molecule were synthesized to establish structure-activity relationships for inhibition of *Chikungunya* (CHIKV) nsP2 protease and viral replication. The covalent warhead was highly sensitive to modifications of the sulfone or vinyl substituents. However, numerous alterations to the core 5-membered heterocycle and its aryl substituent were well tolerated and several analogs were identified that enhanced CHIKV nsP2 binding. For example, the 4-cyanopyrazole analog **8d** exhibited a *k_inact_*/*K_i_* ratio >10,000 M^-1^s^-1^. 3-Arylisoxazole was identified an isosteric replacement for the 5-membered heterocycle, which circumvented the intramolecular cyclization that complicated the synthesis of pyrazole-based inhibitors like **1a**. The accumulated structure-activity data was used to build a ligand-based model of the enzyme active site, which can be used to guide the design of covalent nsP2 protease inhibitors as potential therapeutics against alphaviruses.

## Introduction

Alphaviruses, a group of widespread and enveloped RNA viruses, pose a significant threat to public health due to their transmission by broadly distributed mosquito vectors, such as *Aedes aegypti* and *Aedes albopictus*.^1^ These viruses have been placed into two distinct categories based on their geographical origin: Old World alphaviruses and New World alphaviruses, each associated with unique clinical manifestations. Old World alphaviruses, which include chikungunya virus (CHIKV), Ross River virus (RRV), and O’nyong-nyong virus (ONNV), are primarily characterized by their symptomatology of rash, fever, and prolonged arthralgia.^2^ These symptoms can persist for months post-infection, leading to significant morbidity and impacting the quality of life of affected individuals. New World alphaviruses, such as Venezuelan (VEEV), Western (WEEV), and Eastern (EEEV) Equine Encephalitis viruses, along with Mayaro virus (MAYV), present a different clinical picture often associated with encephalitis-like neurological symptoms, including fever, headache, and nausea. The prognosis for these infections can be dire, with EEEV cases resulting in a mortality rate of 30–50%.^3^ The clinical severity of these viruses is underscored by the lack of FDA-approved drugs for any alphavirus-caused disease. The urgency for the development of alphavirus therapeutics is further reinforced by the potential for weaponization, as evidenced by historical attempts with VEEV during the Cold War.^4^ Moreover, the possibility of the emergence of naturally occurring variants with high pandemic potential that could infect large portions of the human population cannot be overlooked.^5^

The largest non-structural protein in the alphavirus genome, nsP2, is essential for viral replication.^6^ nsP2 contains an N-terminal RNA-specific helicase domain, a C-terminal cysteine protease domain, and putative methyltransferase-like domain. The nsP2 protease uses a catalytic dyad of cysteine and histidine residues to effect cleavage of the viral polyprotein P1234 generating the individual nsP1–4 that comprise the viral RNA replicase whose maturation is crucial for the viral replication cycle. Accordingly, nsP2 protease is an attractive target for development of anti-alphavirus therapeutics. β-Aminomethyl vinyl sulfone RA-0002034 (**1a**, Figure 1A) was recently identified as a potent irreversible covalent inhibitor of CHIKV nsP2 protease with IC_50_ = 60 nM.^7^ Vinyl sulfone **1a** also demonstrated potent antiviral activity, inhibiting CHIKV and VEEV replication with EC_50_ = 0.01 and 0.3 µM, respectively, and decreasing viral titer across a wide range of New and Old World alphaviruses.^7^

The tunability and utility of vinyl sulfones as covalent warheads for irreversible cysteine targeting has been studied extensively.^8^ Notably, **1a** was remarkably selective across a wide panel of diverse cysteine proteases.^7^ Furthermore, the clinical development of vinyl sulfones has been demonstrated by K777 (Figure 1A),^9^ a covalent cysteine protease inhibitor that was effective in animal models of Chagas disease, schistosomiasis, hookworm infection, and cryptosporidiosis and VVD-133214, a covalent WRN helicase inhibitor that was recently advanced to clinical trials for cancers with microsatellite instability.^10^ One potential liability of **1a** is its propensity for intramolecular cyclization to an inactive dihydropyrazolo[1,5-*a*]pyrazin-4(5*H*)-one,^11^ which required the development of synthetic and analytical methods to ensure the exclusive synthesis of the acyclic vinyl sulfone **1a**.^11^ Herein, we report a detailed exploration of the structure activity of anti-alphaviral nsP2 protease inhibitors through systematic modification of three regions of the heterocyclic β-amidomethyl vinyl sulfone (Figure 1B).

**Figure 1.**
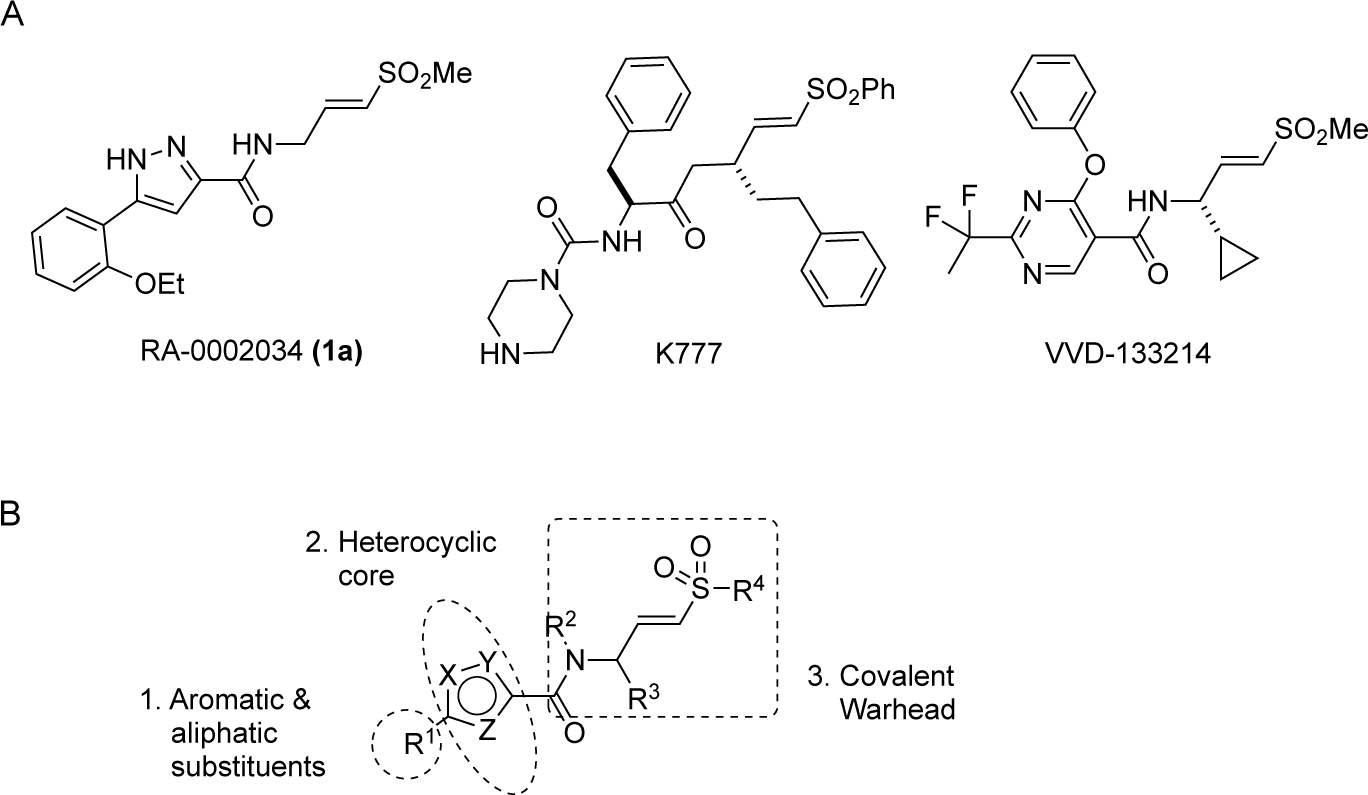
A. Vinyl sulfone covalent cysteine protease (**1a** and K777) and helicase (VVD-133214) inhibitors. B. Three regions of structure activity exploration.

## Results and Discussion

### Chemistry

Synthesis of the analogs **1**–**31** to explore each of the three regions of the covalent nsP2 protease inhibitor **1a** (Figure 1) was accomplished by amide coupling reaction between an appropriate heterocyclic carboxylic acid and an aminomethyl substituted warhead (Schemes 1–4). In most cases, the heterocyclic carboxylic acid was synthesized in multiple steps from available building blocks. Likewise, many of the aminomethyl substituted warheads were not commercially available and required development of a suitable synthesis.

5-aryl-1*H*-pyrazole-3-carboxylic acids **IVa**–**q** were readily prepared from the corresponding acetophenones **Ia**–**q** employing a three-step reaction sequence (Scheme 1A). Claisen condensation of substituted acetophenones **Ia**–**q** with diethyl oxalate in lithium diisopropylamide (LDA) solution afforded ethyl-2,4-dioxo-4-arylbutanoates **IIa**–**q**, which on treatment with hydrazine hydrate under acidic condition yielded the corresponding ethyl 5-aryl-1*H*-pyrazole-3-carboxylates **IIIa**–**q**. Pyrazole esters **IIIa**–**q** on subsequent hydrolysis afforded 5-aryl-1*H*-pyrazole-3-carboxylic acids **IVa**–**q**. Amide coupling of **IV** with (*E*)-3-(methylsulfonyl)prop-2-en-1-amine (**V**) afforded the corresponding pyrazole-substituted β-amidomethyl-vinyl sulfones **1a**⎼**q**. Similarly, 1-cyclohexylethan-1-one (**XII**) and 1,1-diphenylpropan-2-one (**XIV**) afforded analogs **4c** and **4g** following the same reaction sequence. **4d** and **4e** were synthesized using the appropriate THP-protected pyrazole carboxylic acids **XIXa**–**b** and amine **V**. **XIXa**–**b** were synthesized from 1-(2-ethoxyphenyl)propan-2-one (**XVIIa**) and 3-(2-ethoxyphenyl)butan-2-one (**XVIIb**) following a sequence of pyrazole formation/THP protection/ester hydrolysis reactions. 5-(2-ethoxyphenyl)-1-(tetrahydro-2*H*-pyran-2-yl)-1*H*-pyrazole-3-carboxylic acid (**VII**) and 5-(2-ethoxyphenyl)-1-((2-(trimethylsilyl)ethoxy)methyl)-1*H*-pyrazole-3-carboxylic acid (**IX**) could be accessed following a two-step reaction sequence from ethyl 5-(2-ethoxyphenyl)-1*H*-pyrazole-3-carboxylate (**IIIa**). Ethyl 4-(2-ethoxyphenyl)-2,4-dioxobutanoate (**IIa**) when treated with hydroxylamine hydrochloride afforded ethyl 3-(2-ethoxyphenyl)isoxazole-5-carboxylate (**X**), subsequent hydrolysis yielded acid **XI**, which was coupled with **V** to yield isoxazole vinyl sulfone **10**. Analogs **4h**, **4i**, **5**, **6**, **20** were synthesized by amide coupling between amine **V** and corresponding acid counterpart **XX**–**XXIV** (Scheme 1B). Analogs **2**, **3**, **24f**, **28**, **29**, **30**, and **31** were synthesized using 5-(2-ethoxyphenyl)-1*H*-pyrazole-3-carboxylic acid **IVa** and appropriate amine counterparts **XXVa**–**g** (Scheme 1C).

**Scheme 1.**
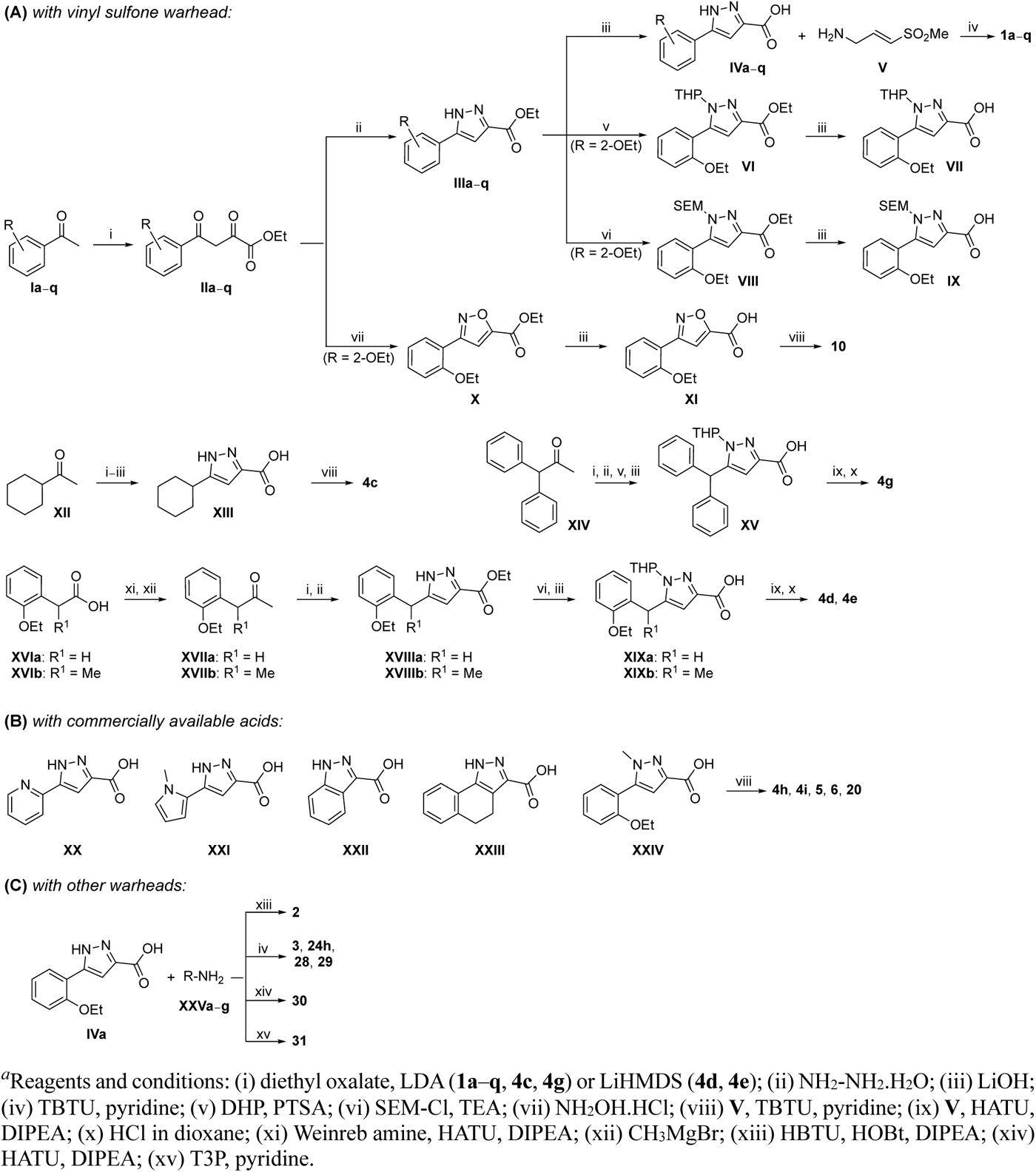
Synthesis of **1a**–**q**, **2**, **3**, **4c**–**e**, **4g**–**i**, **5**, **6**, **24f**, **28**–**31***^a^*

3-amino substituted pyrazole analogs **7a**–**f** were synthesized using the appropriate pyrazole carboxylic acids and amine **V** (Scheme 2). Ethyl 5-amino-1*H*-pyrazole-3-carboxylate (**XXVI**) on treatment with either benzene sulfonyl chloride or phenyl isocyanate, followed by hydrolysis afforded intermediates **XXVIIa**–**b**, which on amide coupling yielded **7a** and **7d** respectively. Amide coupling of **XXVI** with benzoic acid (or 2-ethoxybenzoic acid), followed by THP protection yielded benzamide-containing 1-(tetrahydro-2*H*-pyran-2-yl)-1*H*-pyrazole-3-carboxylates **XXIXa**–**b**. Compounds **7b** and **7c** were synthesized using a three-step sequence from **XXIXa** and **XXIXb** respectively. *N*-Methylation of **XXIXa**– **b**, followed by an ester hydrolysis/amide coupling/THP deprotection sequence afforded **7e** and **7f** respectively. Compound **8a** was prepared from pyrazole intermediate **XXXVI**, which was synthesized from methyl 5-bromo-4-methyl-1*H*-pyrazole-3-carboxylate (**XXXVa**) following a sequence of Suzuki coupling/THP protection/ester hydrolysis. SEM-protected intermediate **XXXVII** was synthesized from ethyl 5-bromo-1*H*-pyrazole-3-carboxylate (**XXXVb**) by a five-step protocol. Amide coupling and THP deprotection yielded **8a** and **8b** respectively. Analogs **12**, **13**, and **22** were synthesized from methyl 5-bromo-1*H*-pyrrole-3-carboxylate (**XXXVc**), ethyl 2-bromooxazole-5-carboxylate (**XXXVd**), and ethyl 2-bromo-5-methyl-1*H*-imidazole-4-carboxylate (**XXXVf**) respectively following a three-step reaction sequence involving Suzuki coupling/ester hydrolysis/amide coupling. 2-bromothiazole-5-carboxylic acid (**XXXVe**) yielded **14** by a two-step Suzuki coupling/amide coupling protocol.

**Scheme 2.**
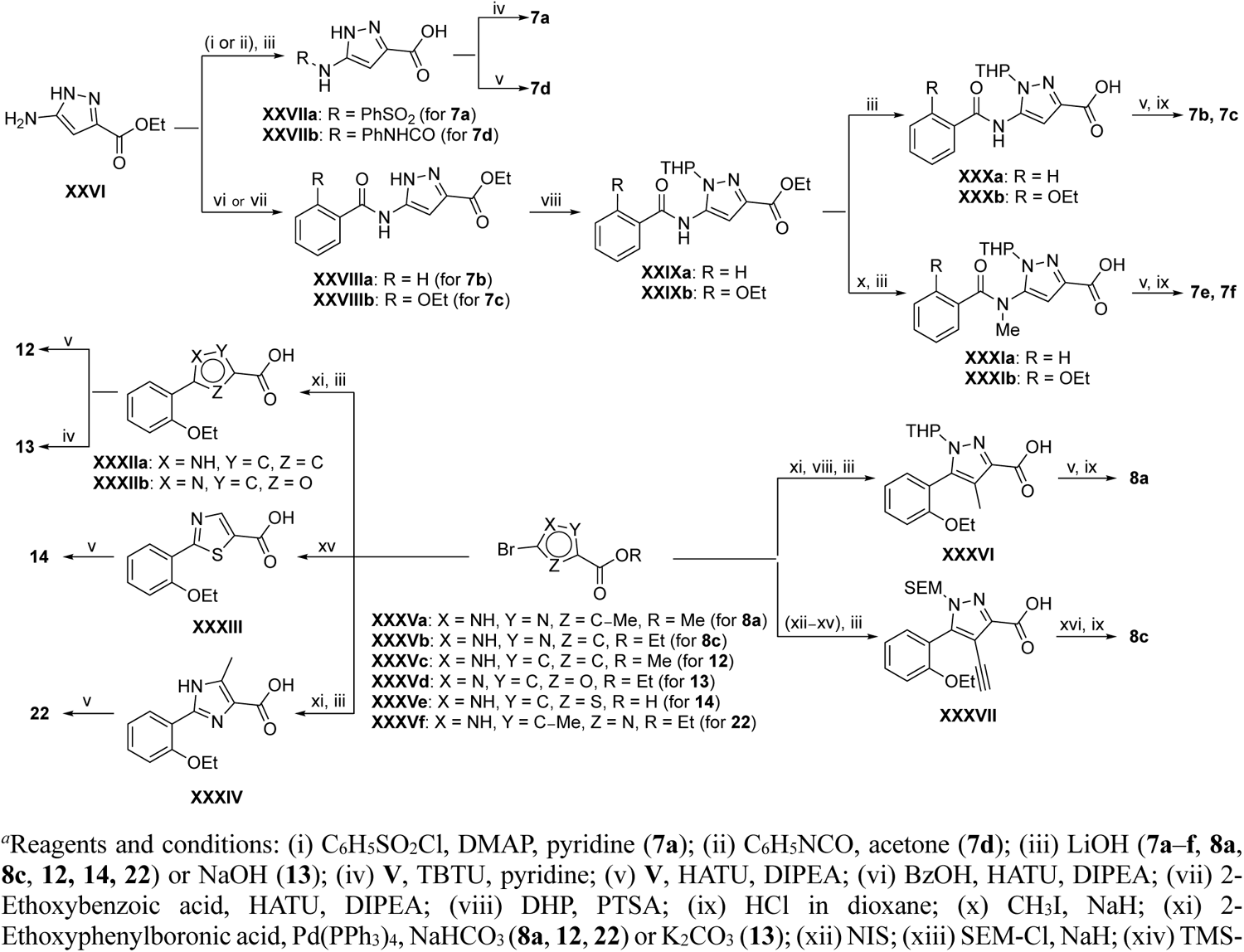
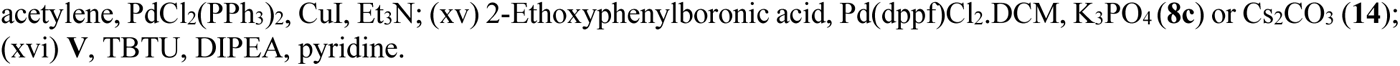
Synthesis of **7a**–**f**, **8a**, **8c**, **12**–**14**, **22***^a^*

Analogs **8b**, **8d**, **15**, and **21** were synthesized from 1-(2-ethoxyphenyl)ethan-1-one (**Ia**) (Scheme 3). Ethyl 5-(2-ethoxyphenyl)-1-(tetrahydro-2*H*-pyran-2-yl)-1*H*-pyrazole-3-carboxylate (**VI**) was synthesized from 1-(2-ethoxyphenyl)ethan-1-one (**Ia**) (Scheme 1A) and used to prepare 4-chloro and 4-cyano substituted pyrazole intermediates **XXXVIII** and **XXXIX** respectively. Amide coupling/THP deprotection yielded analogs **8b** and **8d**. **Ia** on treatment with ethyl 2-isocyanoacetate, followed by ester hydrolysis afforded 5-(2-ethoxyphenyl)oxazole-2-carboxylic acid (**XL**); while amide coupling with **V** yielded compound **15**. 5-(2-ethoxyphenyl)-2-methyl-1*H*-pyrrole-3-carboxylic acid (**XLI**) was synthesized from **Ia** by a four-step reaction sequence. Amide coupling with **V** yielded analog **21**. 3-(2-ethoxyphenyl)isothiazole-5-carboxylic acid (**XLIII**) and 5-(2-ethoxyphenyl)-1,3,4-oxadiazole-2-carboxylic acid (**XLIV**) were synthesized from 2-ethoxybenzamide (**XLIIa**) and 2-ethoxybenzohydrazide (**XLIIb**) respectively. **XLIIa** on treatment with chlorocarbonylsulfenyl chloride/ethyl propiolate/LiOH afforded intermediate **XLIII**, while **XLIIb** on treatment with ethyl 2-chloro-2-oxoacetate/P_2_O_5_/LiOH yielded **XLIV**. Amide coupling with **V** yielded analogs **11** and **18**. Analogs **16** and **17** were synthesized from (2-ethoxyphenyl)boronic acid (**XLV**). Suzuki coupling with 5-bromofuran-2-carboxylate, followed by ester hydrolysis afforded 5-(2-ethoxyphenyl)furan-2-carboxylic acid (**XLVI**); subsequent amide coupling afforded analog **16**. Suzuki coupling with 5-bromothiophene-2-carboxylic acid afforded 5-(2-ethoxyphenyl)thiophene-2-carboxylic acid (**XLVII**), followed by amide coupling to generate analog **17**. 5-(2-ethoxyphenyl)-1,2,4-oxadiazole-3-carboxylic acid (**XLIX**) was synthesized by a three-step reaction sequence starting from ethyl cyanoformate **XLVIII**. Amide coupling of **XLIX** with **V** yielded analog **19**. Diethyl 2-oxosuccinate (**L**) on treatment with hydroxylamine hydrochloride, and subsequent bromination afforded ethyl 5-bromoisoxazole-3-carboxylate (**LI**). Suzuki coupling/ester hydrolysis sequence afforded 5-(2-ethoxyphenyl)isoxazole-3-carboxylic acid (**LII**); amide coupling with **V** yielded analog **9**.

**Scheme 3.**
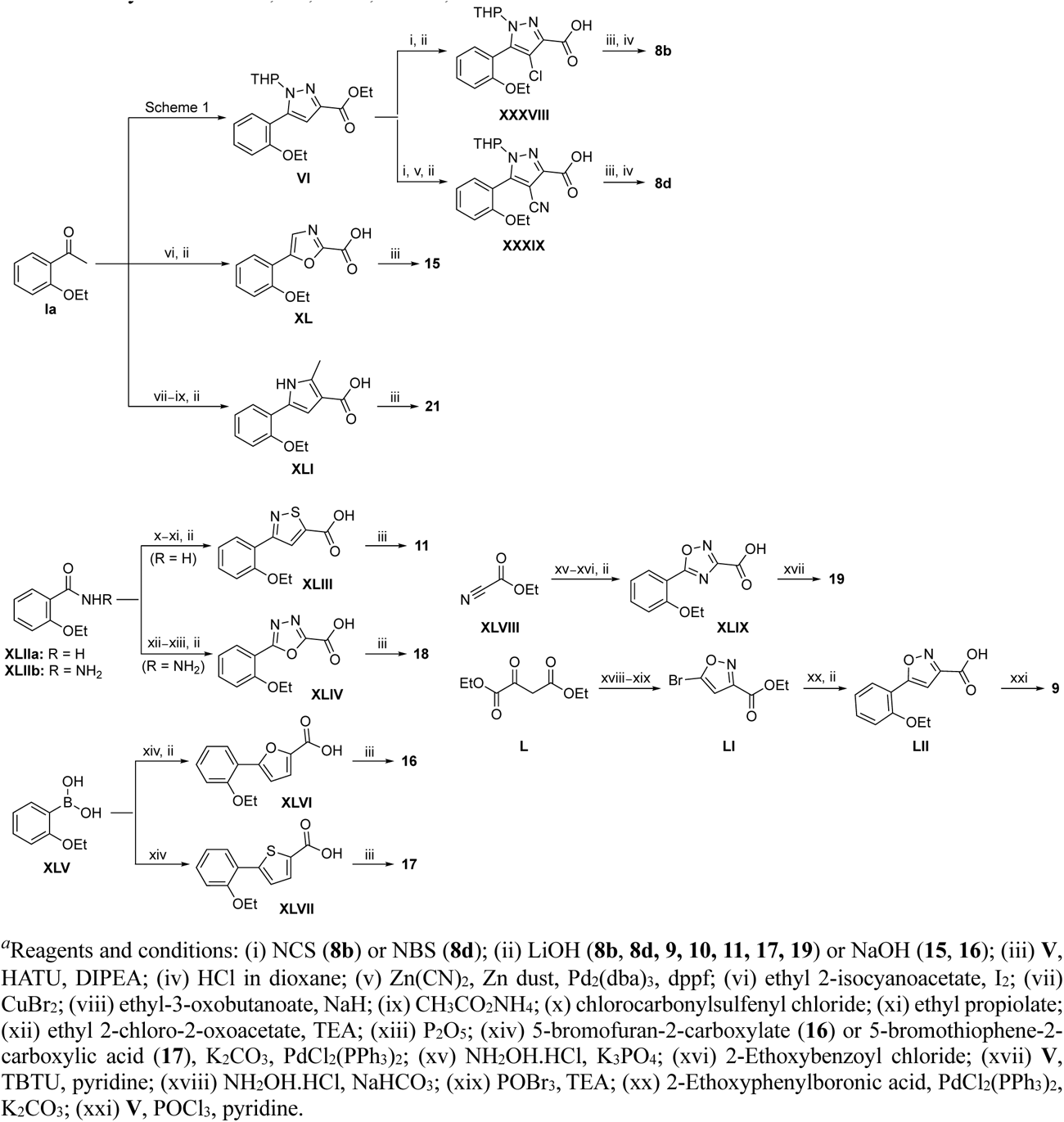
Synthesis of **8b**, **8d**, **9**–**11**, **15**–**19**, **21***^a^*

Modified aminomethyl vinyl sulfones were synthesized by multistep sequences (Scheme 4). Propargylamine **LIIIa** was subjected to a sequence of Boc-protection/radical reaction with tributyltin hydride/iodination to afford *tert*-butyl (*E*)-(3-iodoallyl)carbamate (**LIV**). Intermediate **LIV** was subjected to different reaction conditions to yield vinyl sulfone/sulfoxides **LV**–**LVII**. Treatment with sodium cyclopropanesulfinate and subsequent Boc-deprotection yielded (*E*)-3-(cyclopropylsulfonyl)prop-2-en-1-amine (**LV**); which was followed by a sequence of amide coupling/THP deprotection to yield analog **25a**. Intermediate **LIV** on reaction with pyridine-2-thiol, followed by m-CPBA oxidation, and Boc-deprotection afforded (*E*)-3-(pyridin-2-ylsulfonyl)prop-2-en-1-amine (**LVI**); amide coupling/THP deprotection yielded **25g**. Treatment of **LIV** with sodium thiomethoxide and CuI, followed by m-CPBA oxidation afforded (*E*)-3-(methylsulfinyl)prop-2-en-1-amine (**LVII**). Amide coupling with **VII**, followed by THP deprotection yielded **26**. Sulfoximine-containing analog **27** was synthesized by amide coupling with **XI** followed by treatment with ammonium carbamate and iodobenzene diacetate. 1,1-Dimethylpropargylamine (**LIIIb**) was subjected to a three-step reaction sequence to afford *tert*-butyl (*E*)-(4-iodo-2-methylbut-3-en-2-yl)carbamate (**LVIII**). Subsequent treatment with sodium methanesulfinate/Boc-deprotection afforded (*E*)-2-methyl-4-(methylsulfonyl)but-3-en-2-amine (**LIX**); which on amide coupling with **VII** and THP deprotection afforded analog **24b**. Treatment of **LIIIa** with sodium thiomethoxide and Pd(OAc)_2_ afforded *tert*-butyl (2-(methylthio)allyl)carbamate (**LX**); subsequent m-CPBA oxidation/Boc deprotection yielded 2-(methylsulfonyl)prop-2-en-1-amine (**LXI**). Amide coupling and THP deprotection resulted in analog **24e**.

**Scheme 4.**
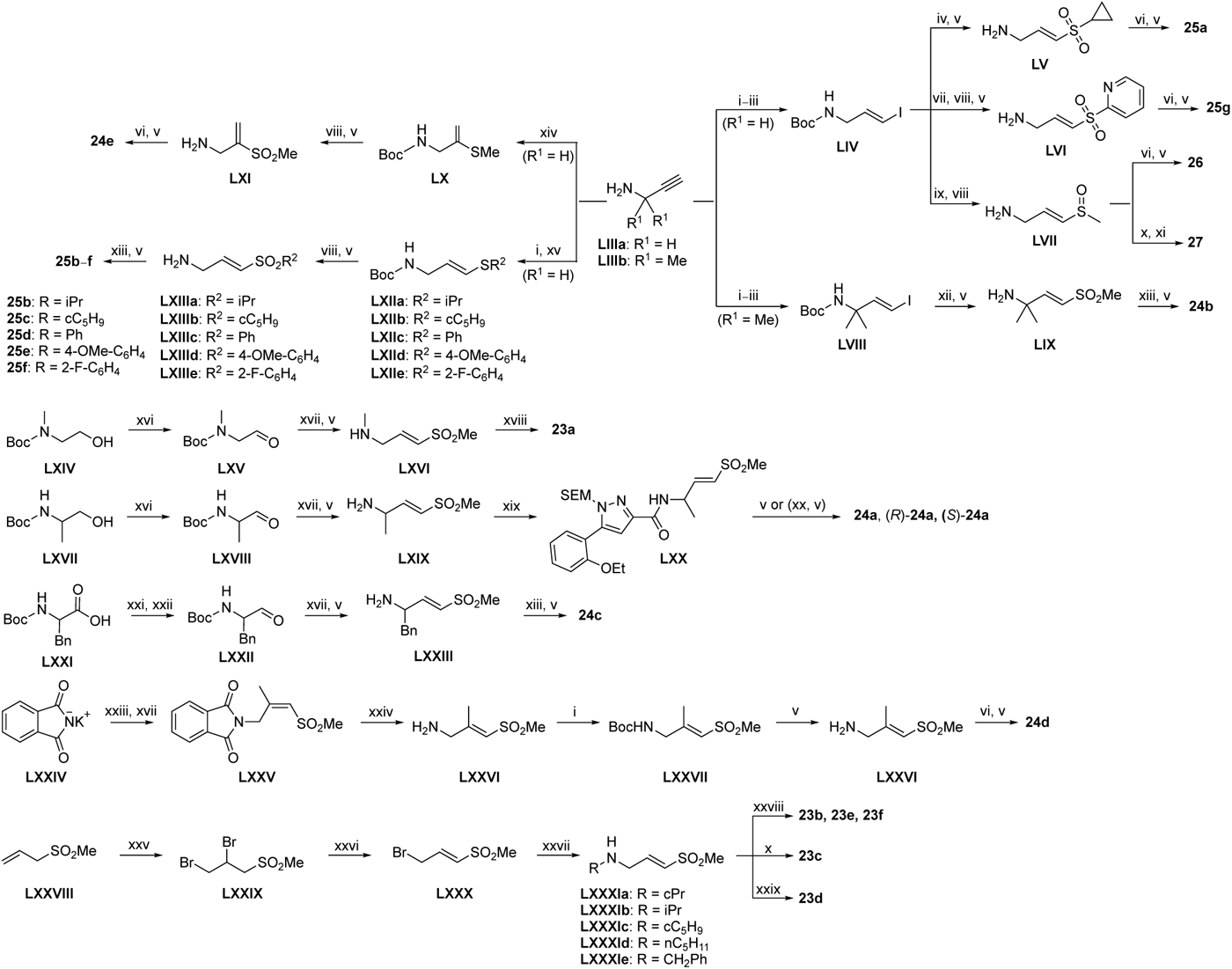
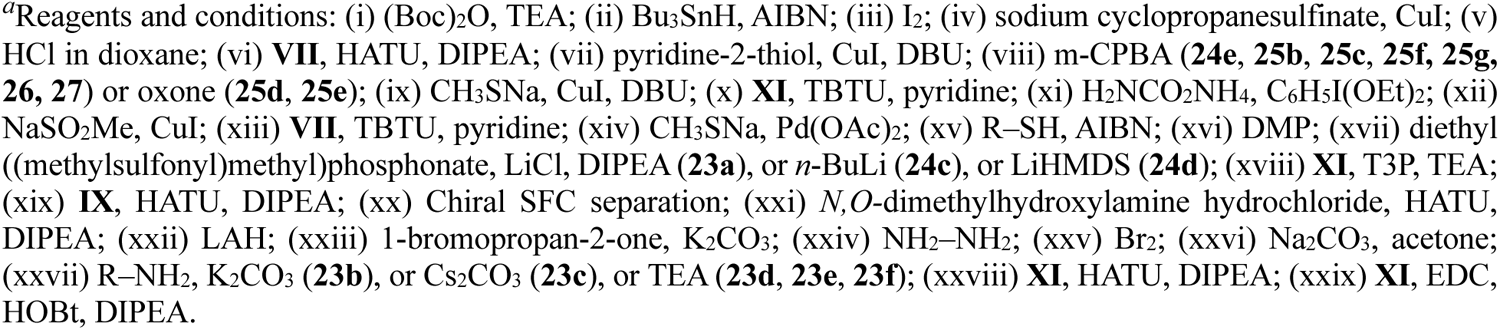
Synthesis of **23a**–**f**, **24a**–**e**, **25a**–**g**, **26**, **27***^a^*

Boc-protected propargylamine on treatment with the corresponding alkyl or aryl thiols resulted in intermediates **LXIIa**–**e**; subsequent m-CPBA or oxone-mediated oxidation and Boc deprotection afforded sulfone amines **LXIIIa**–**e**. Amide coupling and THP deprotection yielded analogs **25b**–**f**. The synthesis of **23a** and **24a** was achieved from the corresponding *tert*-butyl (1-alkyl)carbamates **LXIV** and **LXVII** respectively. A sequence of DMP oxidation/sulfone formation/Boc deprotection provided vinyl sulfones **LXVI** and **LXIX** respectively. Amide coupling of **LXVI** with **XI** yielded **23a**. Amide coupling of intermediate **LXIX** with **IX** afforded **LX**; subsequent SEM deprotection yielded analog **24a**. SFC chiral separation of **LXX**, followed by SEM deprotection afforded the individual enantiomers (*R*)-**24a** and (*S*)-**24a**. Boc-phenylalanine **LXXI** was converted to *tert*-butyl (1-oxo-3-phenylpropan-2-yl)carbamate (**LXXII**) over two steps. Sulfone formation and Boc deprotection provided intermediate **LXXIII**. Amide coupling and THP deprotection yielded analog **24c**.

(*E*)-2-methyl-3-(methylsulfonyl)prop-2-en-1-amine (**LXXVI**) was synthesized using a five-step reaction sequence from potassium phthalimide **LXXIV**. **LXXIV** on treatment with 1-bromopropan-2-one, followed by sulfone formation afforded **LXXV**; subsequent reaction with hydrazine provided intermediate **LXXVI**. Intermediate **LXXVI** was highly polar and UV-inactive which led to difficulty in purification and isolation. However, the Boc-protected intermediate **LXXVII** was easier to purify; subsequent Boc deprotection afforded vinyl sulfone amine **LXXVI** which on amide coupling and THP deprotection yielded the **24d** as an inseparable 1:3 mixture of allyl:vinyl regioisomers. 5-phenyl 2-carboxamido isoxazoles **23b**– **f** were synthesized from their corresponding *N*-substituted vinyl sulfone amines **LXXIa**–**e**, which were accessed from 3-(methylsulfonyl)prop-1-ene (**LXXVIII**) following a three-step reaction protocol. Amide coupling under different conditions furnished **23b**–**f**.

### Biological Evaluation

As expected for an irreversible covalent inhibitor, **1a** demonstrated a time-dependent inhibition of the nsP2 enzyme.^7^ To support structure-activity studies an optimized CHIKV nsP2 protease assay was developed using a 30 min incubation with the test inhibitor. Under these conditions **1a** had an IC_50_ = 60 nM.^7^ Antiviral activity of nsP2 inhibitors was measured using a CHIKV-nLuc viral replication in human fibroblast MRC5 cells at 6 h post inoculation with the virus.^7^ In the CHIKV-nLuc assay **1a** demonstrated an EC_50_ = 40 nM (Table 1).

**Table 1.**
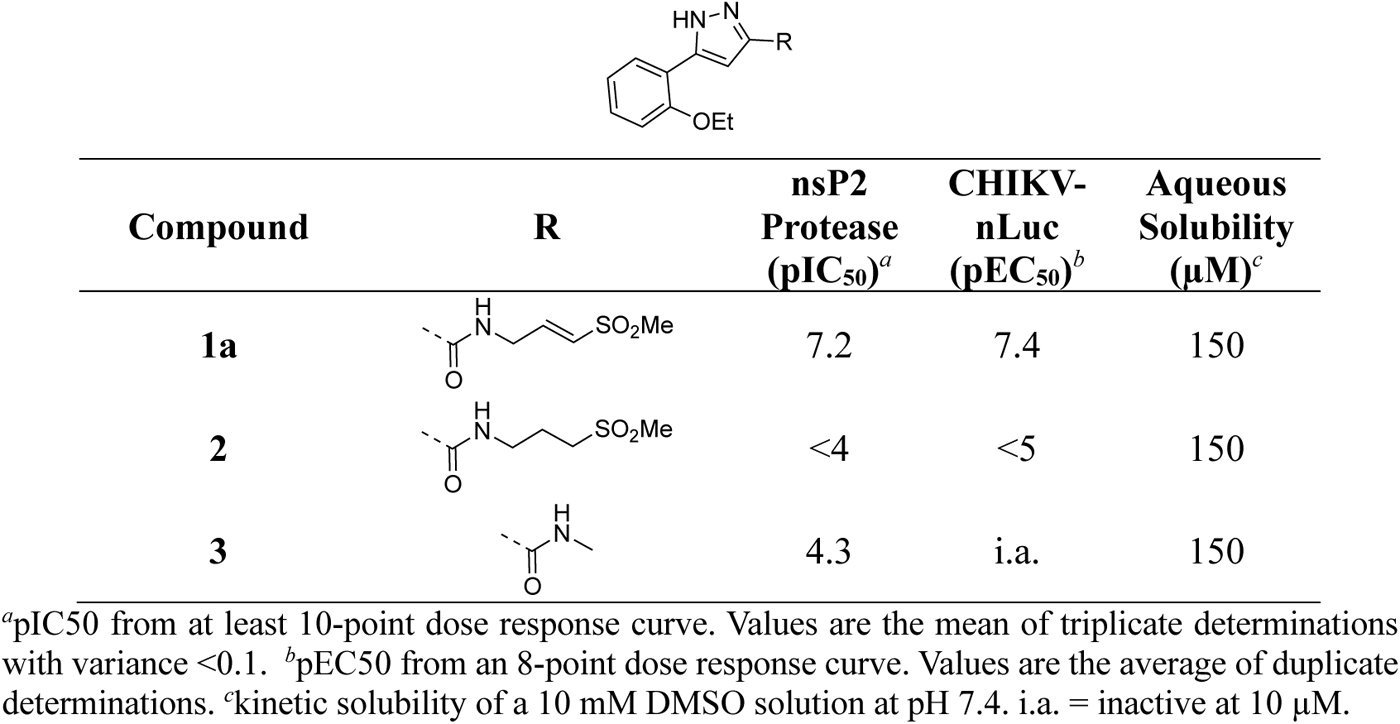
nsP2 Protease Inhibition, Antiviral Activity, and Solubility of 1a, 2, and 3.

| Compound | R | nsP2<br>Protease<br>(pIC<sub>50</sub>)<sup>a</sup> | CHIKV-nLuc<br>(pEC<sub>50</sub>)<sup>b</sup> | Aqueous<br>Solubility<br>(μM)<sup>c</sup> |
| --- | --- | --- | --- | --- |
| **1a** |  | 7.2 | 7.4 | 150 |
| **2** |  | <4 | <5 | 150 |
| **3** |  | 4.3 | i.a. | 150 |

<sup>a</sup>pIC<sub>50</sub> from at least 10-point dose response curve. Values are the mean of triplicate determinations with variance <0.1. <sup>b</sup>pEC<sub>50</sub> from an 8-point dose response curve. Values are the average of duplicate determinations. <sup>c</sup>kinetic solubility of a 10 mM DMSO solution at pH 7.4. i.a. = inactive at 10 μM.

Two analogs were synthesized to demonstrate the critical role of the vinyl sulfone for covalent inhibition of the viral cysteine protease (Table 1). The saturated sulfone **2** and the acetamide **3** were >1000-fold less active in the nsP2 protease and CHIKV antiviral assays. These results demonstrated the critical role of the vinyl sulfone warhead in the nsP2 protease inhibition and antiviral activity of **1a**. The three 5-aryl pyrazoles (**1a, 2–3**) showed good aqueous solubility, with no effect from modification or deletion of the vinyl sulfone warhead. Furthermore, **1a** showed no evidence of compound aggregation, which can confound cell-based antiviral activity,^12^ at concentrations below 100 μM by dynamic light scattering (Figure S1). The increase in laser intensity was linear until the limit of aqueous solubility.

To explore the contribution of the 5-aryl group to the activity of **1a**, a series of analogs were synthesized in which the ring substituents were modified (Table 2). The analog **1b** with an unsubstituted phenyl substituent had nearly equivalent activity to **1a**, demonstrating that the 2-ethoxy group contributed relatively little to the potency of protease inhibition. Within a series of 2-substituted phenyl analogs **1c–f**, analogs with chloro (**1c**) and methyl (**1e**) substituents maintained good activity while the phenol (**1d**) and trifluoromethyl (**1f**) analogs were less active. The 3-alkoxy phenyl analogs **1g–h** showed potent protease inhibition and antiviral activity, while the 3-chlorophenyl analog **1i** was potent in the protease assay but much weaker in the antiviral assay. For the 4-substituted phenyl analogs, electron donating groups (**1j–l**) resulted in good activity while the trifluoromethyl (**1m**) analog was less active. A series of disubstituted phenyl analogs **1n– q** mirrored the activity of the monosubstituted analogs, with 2,5-dimethoxy phenyl analog (**1o**) being marginally more potent in both the protease and replicon assays. Overall, the substituted phenyl analogs showed that small neutral or electron donating substituents were well tolerated but contributed little additional potency to the cysteine protease inhibition effected by the ß-amidomethyl vinyl sulfones. Analogs with the 2-aryl substituted pyrazoles had good aqueous solubility, while those with 3-and 4-aryl substituents often showed reduced solubility.

**Table 2.** nsP2 Protease Inhibition, Antiviral Activity, and Solubility of 1b–q.

| Compound | R | nsP2<br>Protease<br>(pIC<sub>50</sub>)<sup>a</sup> | CHIKV-nLuc<br>(pEC<sub>50</sub>)<sup>b</sup> | Aqueous<br>Solubility<br>(μM)<sup>c</sup> |
| --- | --- | --- | --- | --- |
| **1b** | – | 6.9 | 7.3 | 150 |
| **1c** | 2-Cl | 7.1 | 7.3 | 150 |
| **1d** | 2-OH | 6.4 | 6.1 | 180 |
| **1e** | 2-Me | 7.2 | 7.1 | 150 |
| **1f** | 2-CF<sub>3</sub> | 6.4 | 6.3 | 150 |
| **1g** | 3-OMe | 7.2 | 7.1 | 140 |
| **1h** | 3-OEt | 7.5 | 7.0 | 60 |
| **1i** | 3-Cl | 7.1 | 5.8 | 60 |
| **1j** | 4-OEt | 6.6 | 6.6 | 70 |
| **1k** | 4-SMe | 6.8 | 6.7 | 10 |
| **1l** | 4-NMe<sub>2</sub> | 6.8 | 7.4 | 20 |
| **1m** | 4-CF<sub>3</sub> | 5.9 | 5.5 | <10 |
| **1n** | 2,4-Me | 7.2 | 7.2 | 50 |
| **1o** | 2,5-OMe | 7.6 | 7.4 | 10 |
| **1p** | 2,6-Cl | 6.5 | 5.9 | 140 |
| **1q** | 3,5-Cl | 7.3 | 5.2 | <10 |

<sup>a</sup>pIC<sub>50</sub> from at least 10-point dose response curve. Values are the mean of triplicate determinations with variance <0.1. <sup>b</sup>pEC<sub>50</sub> from an 8-point dose response curve. Values are the average of duplicate determinations. <sup>c</sup>kinetic solubility of a 10 mM DMSO solution at pH 7.4.

Since small variations in the 5-aryl group substituents resulted in only minor changes in nsP2 protease inhibition a new series of analogs was synthesized with larger modifications (Table 3). The 5-methyl (**4a**) and 5-trifluromethyl (**4b**) pyrazoles were inactive. However, the saturated 5-cyclohexyl pyrazole (**4c**) was as active as the best of the 5-aryl groups, indicating that any lipophilic substitution at this position was sufficient for potent nsP2 protease and antiviral activity. Analogs **4d–f** with 5-benzyl groups were also active, further demonstrating that the pyrazole did not require direct conjugation to an aromatic substituent. The retention of activity in the 1,1-diphenylmethane analog **4g** showed that the nsP2 protease could accommodate a large substituent at this position, although the lack of further increase in activity suggested that there was no additional binding interaction resulting from addition of the extra phenyl ring. The potent protease inhibition of the pyridyl (**4h**) and *N*-methyl pyrrole (**4i**) analogs showed that this region of the enzyme could accommodate some limited polarity. However, the decreased activity of the pyridyl analog **4h** in the antiviral assay suggested that it may have lower cell penetration. Notably, these analogs with variations in the pyrazole 5-substituent retained good aqueous solubility despite the large variations in atom count and molecular weight.

**Table 3.**
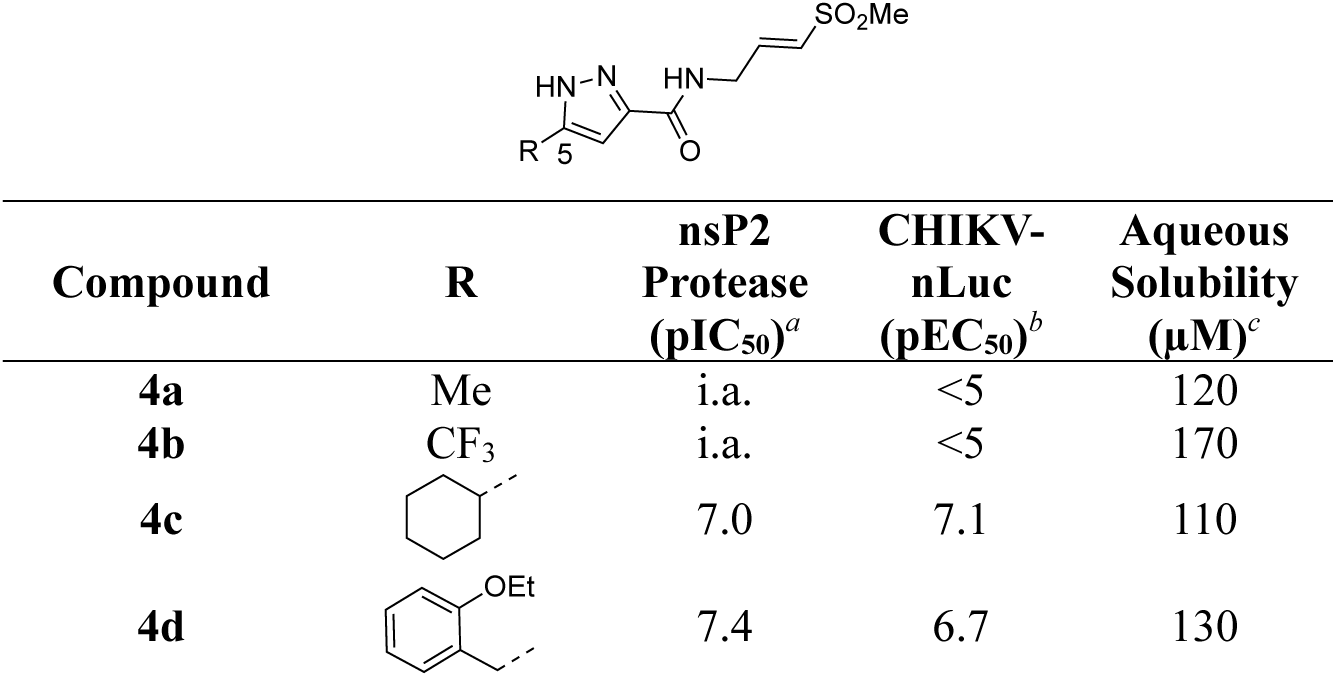
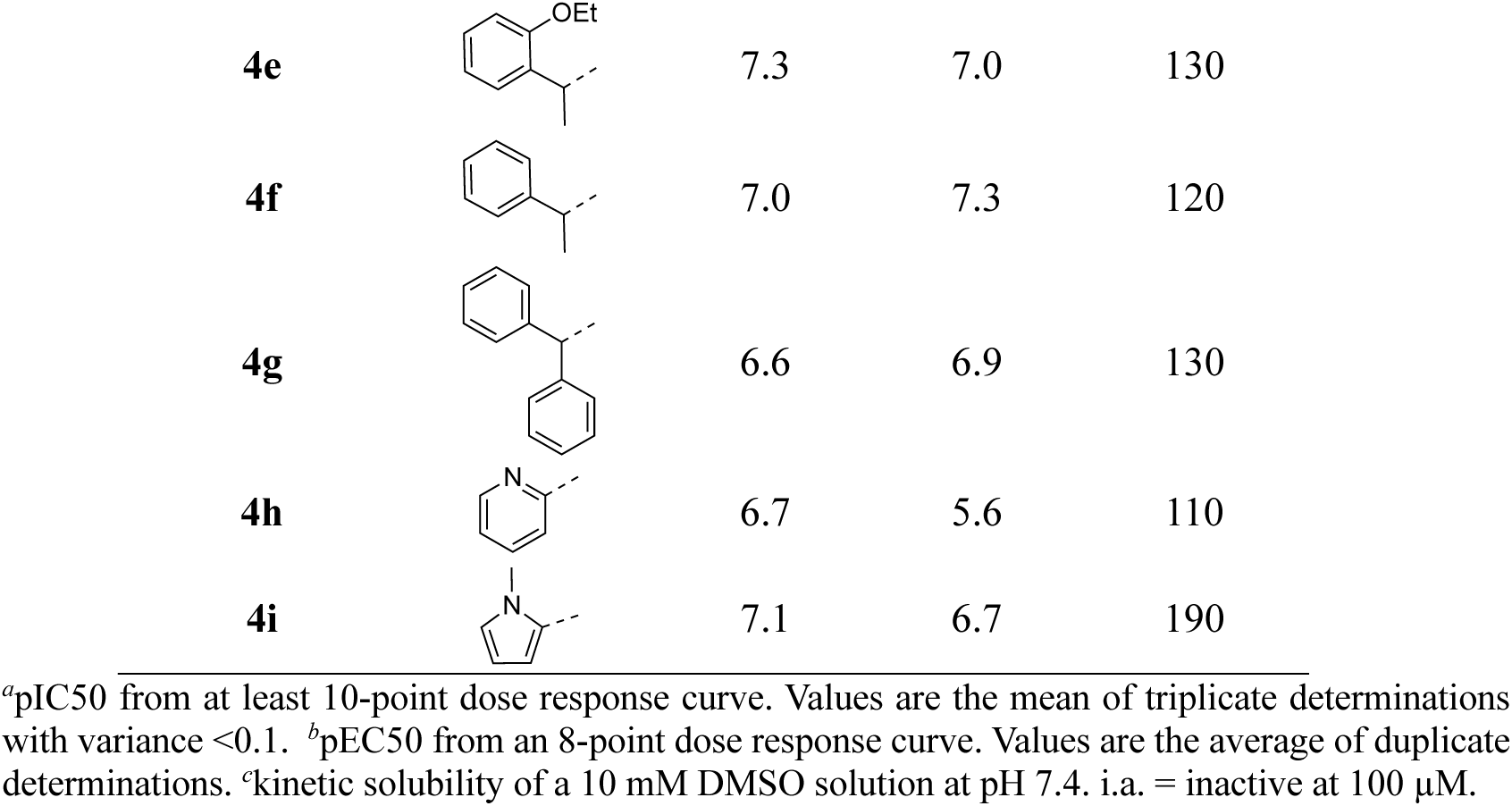
nsP2 Protease Inhibition, Antiviral Activity, and Solubility of 4a–i.

| Compound | R | nsP2<br>Protease<br>(pIC<sub>50</sub>)<sup>a</sup> | CHIKV-nLuc<br>(pEC<sub>50</sub>)<sup>b</sup> | Aqueous<br>Solubility<br>(μM)<sup>c</sup> |
| --- | --- | --- | --- | --- |
| **4a** | Me | i.a. | <5 | 120 |
| **4b** | CF<sub>3</sub> | i.a. | <5 | 170 |
| **4c** |  | 7.0 | 7.1 | 110 |
| **4d** |  | 7.4 | 6.7 | 130 |
| **4e** |  | 7.3 | 7.0 | 130 |
| **4f** |  | 7.0 | 7.3 | 120 |
| **4g** |  | 6.6 | 6.9 | 130 |
| **4h** |  | 6.7 | 5.6 | 110 |
| **4i** |  | 7.1 | 6.7 | 190 |

<sup>a</sup>pIC<sub>50</sub> from at least 10-point dose response curve. Values are the mean of triplicate determinations with variance <0.1. <sup>b</sup>pEC<sub>50</sub> from an 8-point dose response curve. Values are the average of duplicate determinations. <sup>c</sup>kinetic solubility of a 10 mM DMSO solution at pH 7.4. i.a. = inactive at 100 μM.

To further explore the contribution of the pyrazole 5-phenyl group to nsP2 protease inhibition, two analogs were synthesized with ring fusions to the heterocycle (Table 4). Indazole **5**, which can be viewed as a 4,5-fusion of the phenyl ring to the pyrazole, retained sub-micromolar potency as a protease inhibitor and in the antiviral assay. The 4,5-dihydro-1*H*-benzo[*g*]indazole **6** had reduced nsP2 protease activity and reduced solubility compared to the indazole **5** but still demonstrated sub-micromolar activity in the antiviral assay.

**Table 4.**
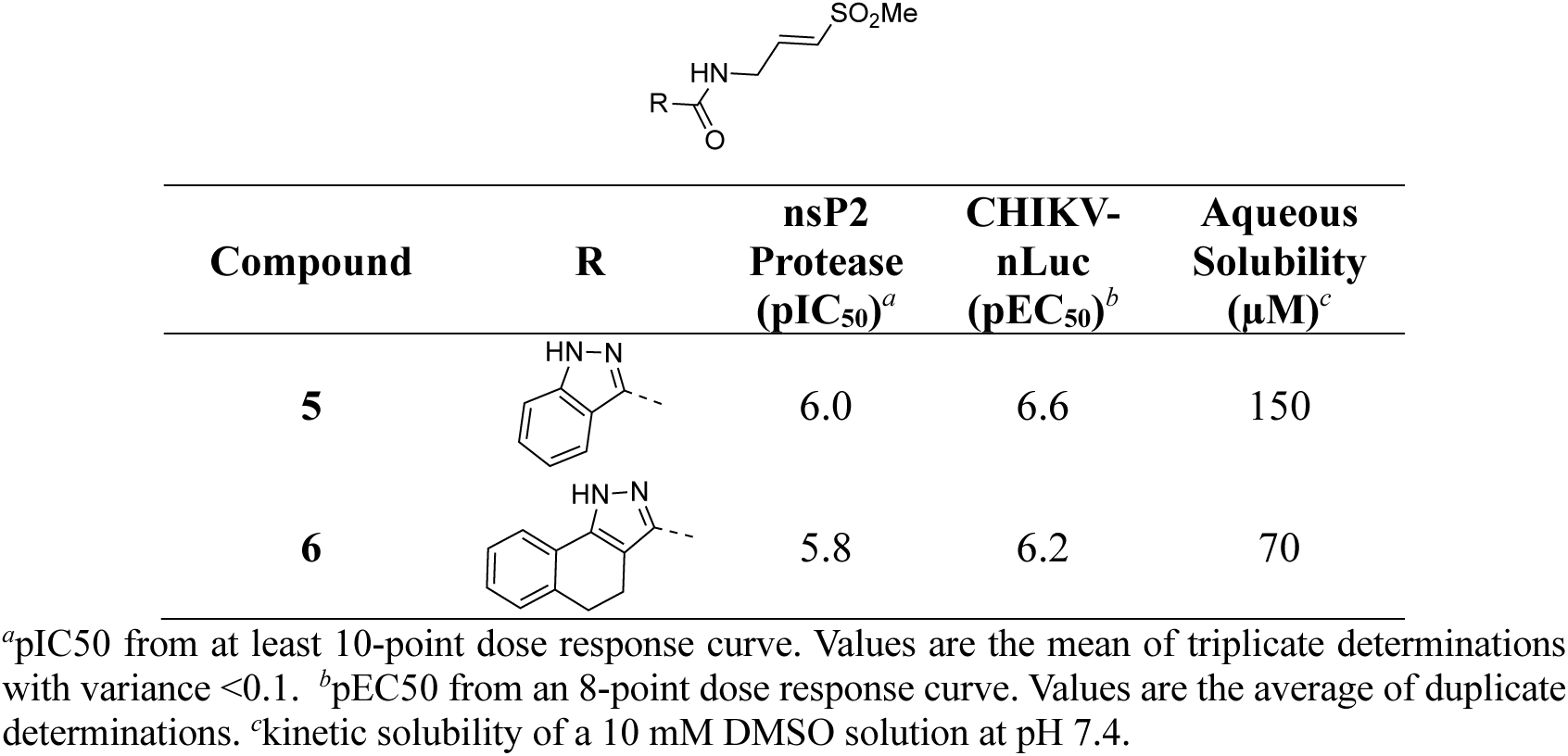
nsP2 Protease Inhibition, Antiviral Activity, and Solubility of 5 and 6.

| Compound | R | nsP2<br>Protease<br>(pIC<sub>50</sub>)<sup>a</sup> | CHIKV-nLuc<br>(pEC<sub>50</sub>)<sup>b</sup> | Aqueous<br>Solubility<br>(μM)<sup>c</sup> |
| --- | --- | --- | --- | --- |
| **5** |  | 6.0 | 6.6 | 150 |
| **6** |  | 5.8 | 6.2 | 70 |

<sup>a</sup>pIC<sub>50</sub> from at least 10-point dose response curve. Values are the mean of triplicate determinations with variance <0.1. <sup>b</sup>pEC<sub>50</sub> from an 8-point dose response curve. Values are the average of duplicate determinations. <sup>c</sup>kinetic solubility of a 10 mM DMSO solution at pH 7.4.

A series of 3-amino substituted pyrazole analogs were synthesized (Table 5). The sulfonamide **7a** had reduced activity in the nsP2 protease assay and was inactive in the antiviral assay. The benzamide **7b** showed good activity as a protease inhibitor, but it had lower aqueous solubility. Its modest potency in the antiviral assay may be due to poor physiochemical properties, since it had a calculated Balanced Permeability Index (BPI)^13^ score <0.5 that is often associated with poor cell penetration. Analog **7c** which added a 2-ethoxy group to the benzamide and the urea analog **7d** also showed good protease inhibition but lower activity in the antiviral assay. These two analogs had BPI scores <0.5 predicting low cell permeability. Two tertiary amides (**7e–f**) demonstrated a ∼10-fold decrease in potency for protease inhibition compared to the respective secondary amides (**7b–c**) and were inactive in the antiviral assay; despite their improved solubility the MPI scores remained below 0.5.

**Table 5.**
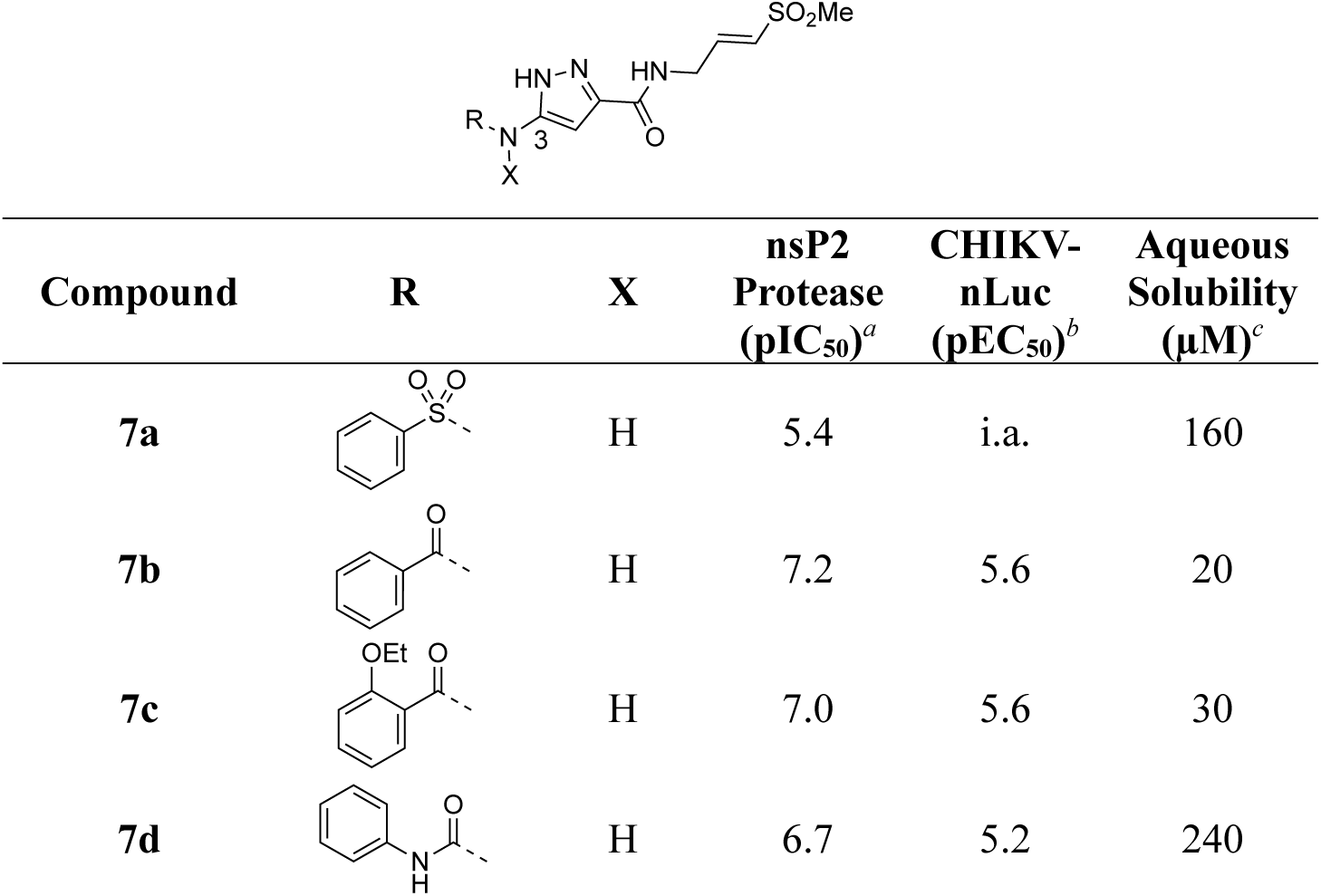
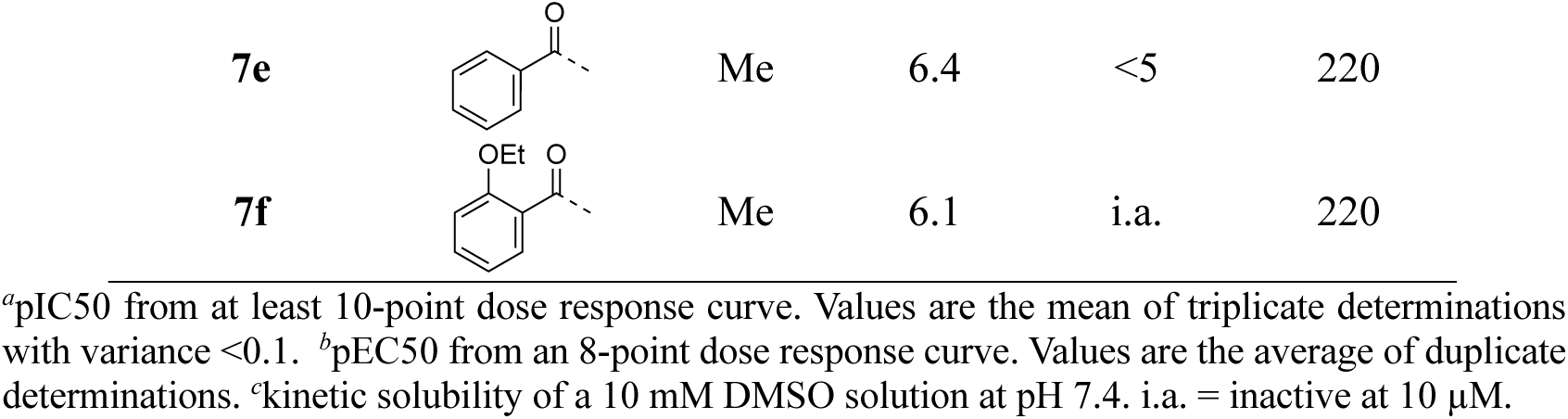
nsP2 Protease Inhibition, Antiviral Activity, and Solubility of 7a–f.

| Compound | R | X | nsP2<br>Protease<br>(pIC<sub>50</sub>)<sup>a</sup> | CHIKV-nLuc<br>(pEC<sub>50</sub>)<sup>b</sup> | Aqueous<br>Solubility<br>(μM)<sup>c</sup> |
| --- | --- | --- | --- | --- | --- |
| **7a** |  | H | 5.4 | i.a. | 160 |
| **7b** |  | H | 7.2 | 5.6 | 20 |
| **7c** |  | H | 7.0 | 5.6 | 30 |
| **7d** |  | H | 6.7 | 5.2 | 240 |
| **7e** |  | Me | 6.4 | <5 | 220 |
| **7f** |  | Me | 6.1 | i.a. | 220 |

<sup>a</sup>pIC<sub>50</sub> from at least 10-point dose response curve. Values are the mean of triplicate determinations with variance <0.1. <sup>b</sup>pEC<sub>50</sub> from an 8-point dose response curve. Values are the average of duplicate determinations. <sup>c</sup>kinetic solubility of a 10 mM DMSO solution at pH 7.4. i.a. = inactive at 10 μM.

Addition of small substituents to the 4-position of the pyrazole were explored but were generally not well tolerated. The 4-methyl (**8a**), 4-chloro (**8b**), and 4-ethynyl (**8c**) pyrazoles were poor inhibitors of the nsP2 protease and less active in the CHIKV antiviral assay (Table 6). Surprisingly, however, the 4-cyano pyrazole **8d** was 100-fold more potent than the other 4-substituted pyrazoles as a protease inhibitor and demonstrated sub-micromolar activity in the antiviral assay. The robust activity of the 4-cyano analog **8d** suggested that a small pocket was present in the protease, but it was very sensitive to the chemical properties of the pyrazole 4-substituent.

**Table 6.** nsP2 Protease Inhibition, Antiviral Activity, and Solubility of 8a–d.

| Compound | R | nsP2<br>Protease<br>(pIC<sub>50</sub>)<sup>a</sup> | CHIKV-nLuc<br>(pEC<sub>50</sub>)<sup>b</sup> | Aqueous<br>Solubility<br>(μM)<sup>c</sup> |
| --- | --- | --- | --- | --- |
| **8a** | Me | 5.0 | 5.8 | 200 |
| **8b** | Cl | 5.1 | <5 | 50 |
| **8c** | C≡C | 5.5 | 5.4 | 140 |
| **8d** | CN | 7.6 | 6.5 | 170 |

<sup>a</sup>pIC<sub>50</sub> from at least 10-point dose response curve. Values are the mean of triplicate determinations with variance <0.1. <sup>b</sup>pEC<sub>50</sub> from an 8-point dose response curve. Values are the average of duplicate determinations. <sup>c</sup>kinetic solubility of a 10 mM DMSO solution at pH 7.4.

Pyrazole **1a** was previously shown to undergo an intramolecular cyclization onto the vinyl sulfone warhead to form a dihydropyrazolo[1,5-*a*]pyrazin-4(5*H*)-one that was inactive as an nsP2 protease inhibitor.^11^ Although **1a** and related pyrazoles were stable as their TFA or HCl salts in 10 mM DMSO, cyclization was shown to occur over several days at pH 7.4 in phosphate buffer.^11^ To identify isosteric replacements of the pyrazole that were no longer prone to intramolecular cyclization a series of alternative 5-membered heterocycles were synthesized (Table 7). The 5-phenyl isoxazole **9** and the isomeric 3-phenyl isoxazole **10** were both effective replacements for the pyrazole, with the latter showing slightly improved protease inhibition and antiviral activity. The corresponding 3-phenyl isothiazole **11**, 5-phenyl pyrrole **12**, and 2-phenyl oxazole **13** were less potent as nsP2 protease inhibitors. However, the 2-phenyl thiazole **14** and 5-phenyl oxazole **15** were as potent as the original 5-phenyl pyrazole **1a**. Surprisingly, the 5-phenyl thiophene **16** and 5-phenyl furan **17**, which lacked nitrogen atoms in the heterocycle, were still active in the antiviral assay. Although the 5-phenyl 1,2,5-oxadiazole **18** was less active as an nsP2 protease inhibitor, the isomeric 5-phenyl 1,2,4-oxadiazole **19** had sub-micromolar potency in the protease assay and an EC_50_ = 100 nM in the antiviral assay. The *N*-methyl pyrazole **20**, 2-methyl pyrrole **21**, and 5-methyl imidazole **22**, which contained methyl substituents on the heterocyclic ring, were less active as nsP2 protease inhibitors suggesting that there was very limited room in the enzyme to accommodate even a small alkyl substituent. In sum, minor changes to the presence and arrangement of the heteroatoms in the central heterocycle led to subtle differences in their activity as nsP2 protease inhibitors. No single heteroatom appeared to be essential for enzyme inhibition, which suggested the absence of a specific H-bond between the heterocyclic core and the nsP2 enzyme. Importantly, none of the alternative heterocycles (**9–22**) formed cyclic byproducts arising from intramolecular cyclization onto the vinyl sulfone. Thus, several bioisosteric replacements of the pyrazole heterocycle were identified with the 3-phenyl isoxazole **10** being the most potent of these analogs for nsP2 protease and CHIKV inhibition.

**Table 7.** nsP2 Protease Inhibition, Antiviral Activity, and Solubility of 9–22.

X = C, N, O Y/Z = C, N, O, S
| Compound | R | nsP2<br>Protease<br>(pIC<sub>50</sub>)<sup>a</sup> | CHIKV-nLuc<br>(pEC<sub>50</sub>)<sup>b</sup> | Aqueous<br>Solubility<br>(μM)<sup>c</sup> |
| --- | --- | --- | --- | --- |
| 9 |  | 6.5 | 7.1 | 220 |
| 10 |  | 7.4 | 7.3 | 20 |
| 11 |  | 6.0 | 6.6 | 130 |
| 12 |  | 6.0 | 6.5 | 220 |
| 13 |  | 6.3 | 6.7 | 170 |
| 14 |  | 6.9 | 7.2 | 160 |
| 15 |  | 7.2 | 7.2 | 160 |
| 16 |  | 6.2 | 6.9 | 210 |
| 17 |  | 6.3 | 6.8 | 10 |
| 18 |  | 5.5 | 6.4 | 160 |
| 19 |  | 6.3 | 7.0 | 190 |
| 20 |  | 5.9 | 6.2 | 190 |
| 21 |  | 5.5 | 6.0 | 220 |
| 22 |  | 5.4 | 5.4 | 160 |

<sup>a</sup>pIC<sub>50</sub> from at least 10-point dose response curve. Values are the mean of triplicate determinations with variance <0.1. <sup>b</sup>pEC<sub>50</sub> from an 8-point dose response curve. Values are the average of duplicate determinations. <sup>c</sup>kinetic solubility of a 10 mM DMSO solution at pH 7.4.

The 3-phenyl isoxazole heterocycle was used as an isosteric pyrazole replacement for the synthesis of several modified warheads that were highly prone to cyclization to the corresponding dihydropyrazolo[1,5-*a*]pyrazin-4(5*H*)-one. For example, adding an alkyl substituent to the amide linker between the heterocycle and vinyl sulfone resulted in isolation of only cyclic dihydropyrazolo[1,5-*a*]pyrazin-4(*5H*)-ones when attempted in the pyrazole series. However, switching to the bioisosteric 3-aryl 5-carboxamido isoxazoles, which were unable to undergo intramolecular cyclization, enabled synthesis of **23a–f** (Table 8) and their activity on nsP2 protease inhibition could be determined. The *N*-methyl tertiary amide **23a** was less active as a protease inhibitor but was slightly improved in the cyclopropyl amide **23b**. Surprisingly, switching the tertiary amide substituent to isopropyl (**23c**) or cyclopentyl (**23d**) led to a large decrease in nsP2 protease inhibition. The linear *n*-pentyl substituted tertiary amide **23e** had improved protease inhibition but was not potent in the antiviral assay. Similarly, the *N*-benzyl tertiary amide **23f** had only moderate activity compared to the unsubstituted secondary amide **10**. Overall, there was no advantage gained by incorporation of tertiary amides into the covalent warhead for nsP2 protease inhibition, especially with larger substituents that had only modest antiviral activity and lower aqueous solubility. However, the sub-micromolar nsP2 protease inhibition of the tertiary amides with cyclopropyl (**23b**) and *n*-pentyl (**23e**) substituents showed that there was room in the enzyme to accommodate these alkyl groups.

**Table 8.** nsP2 Protease Inhibition, Antiviral Activity, and Solubility of 23a–f.

| Compound | R | nsP2<br>Protease<br>(pIC<sub>50</sub>)<sup>a</sup> | CHIKV-nLuc<br>(pEC<sub>50</sub>)<sup>b</sup> | Aqueous<br>Solubility<br>(μM)<sup>c</sup> |
| --- | --- | --- | --- | --- |
| **23a** | Me | 5.9 | 5.8 | 200 |
| **23b** | cPr | 6.4 | 6.5 | 210 |
| **23c** | iPr | 4.5 | <5 | 240 |
| **23d** | cC<sub>5</sub>H<sub>9</sub> | i.a. | 5.0 | 50 |
| **23e** | nC<sub>5</sub>H<sub>11</sub> | 6.2 | 5.7 | 40 |
| **23f** | CH<sub>2</sub>Ph | 6.0 | 5.8 | 10 |

<sup>a</sup>pIC<sub>50</sub> from at least 10-point dose response curve. Values are the mean of triplicate determinations with variance <0.1. <sup>b</sup>pEC<sub>50</sub> from an 8-point dose response curve. Values are the average of duplicate determinations. <sup>c</sup>kinetic solubility of a 10 μM DMSO solution at pH 7.4. i.a. = inactive at 100 μM.

Changes to the vinyl sulfone warhead were explored to determine how it would affect nsP2 protease inhibition and antiviral activity (Table 9). In the pyrazole series addition of a methyl substituent to the γ-carbon of the β-amidomethyl vinyl sulfone gave the racemic allylic sulfone **24a** which had good nsP2 protease inhibition. Synthesis and testing of the individual enantiomers showed that activity resided in the (*R*)-**24a**, which was ∼100-fold more active than (*S*)-**24a** in the protease and antiviral assays. The dimethyl substituted analog **24b** was inactive, while the racemic benzyl substituted analog **24c** showed modest activity for protease inhibition but good antiviral activity despite its lower aqueous solubility. Analogs **24a– e** showed that a pocket must be present in the nsP2 protease enzyme that can accommodate substituents with an *R*-configuration on the γ-carbon of the β-amidomethyl vinyl sulfone. Synthesis of vinyl sulfones with modification of the α- and β-carbon was also explored. The β-methyl substituted analog **24d**, which was found to exist as a 1:3 mixture of allyl and vinyl sulfone regioisomers, was essentially inactive. The alternative α-substituted vinyl sulfone **24e** had poor activity compared to the β-substituted vinyl sulfone **1a**. A cyclic vinyl sulfone **24f** was inactive as an nsP2 protease inhibitor. All of pyrazoles with the modified warheads had good aqueous solubility except the benzyl substituted analog **24c**.

**Table 9.**
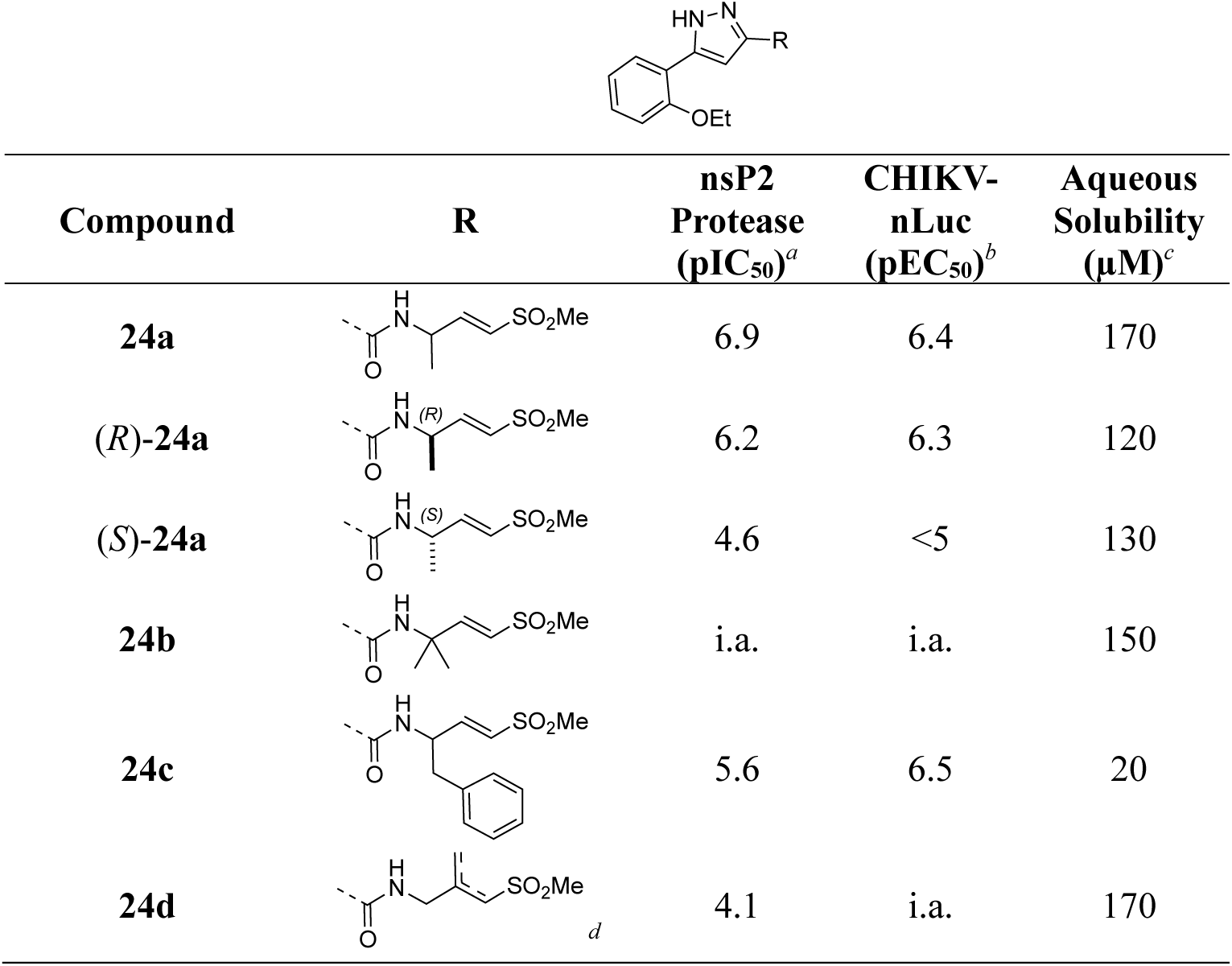
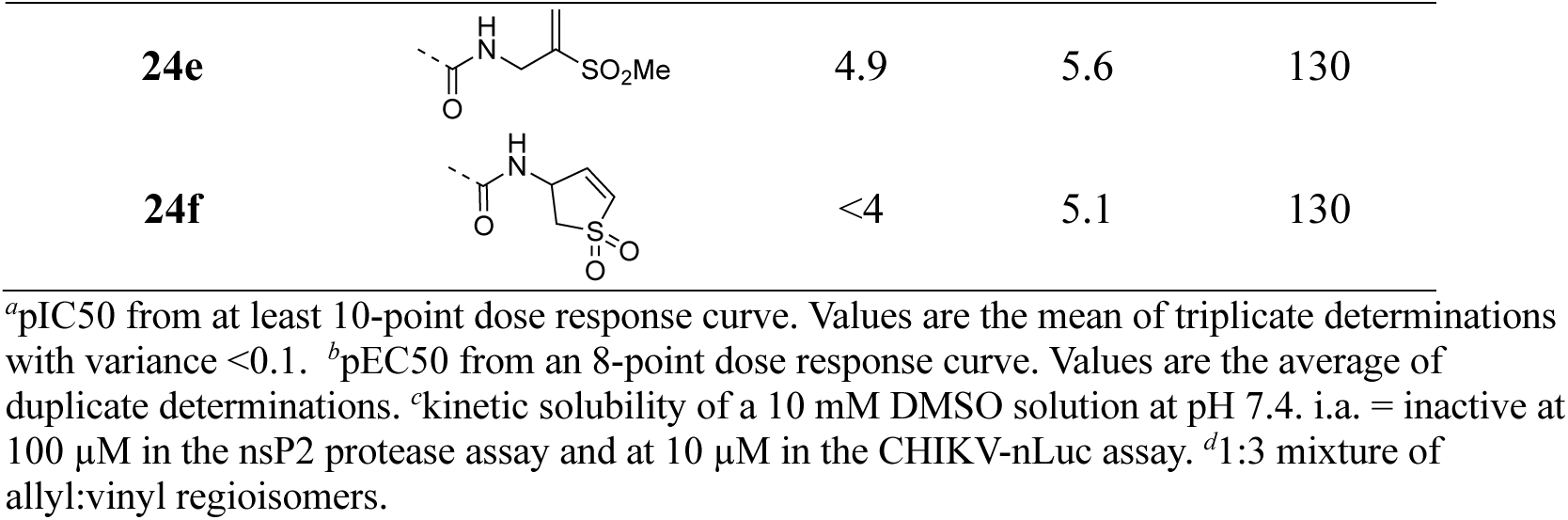
nsP2 Protease Inhibition, Antiviral Activity, and Solubility of 24a–h.

| Compound | R | nsP2<br>Protease<br>(pIC<sub>50</sub>)<sup>a</sup> | CHIKV-nLuc<br>(pEC<sub>50</sub>)<sup>b</sup> | Aqueous<br>Solubility<br>(μM)<sup>c</sup> |
| --- | --- | --- | --- | --- |
| **24a** |  | 6.9 | 6.4 | 170 |
| **(R)-24a** |  | 6.2 | 6.3 | 120 |
| **(S)-24a** |  | 4.6 | <5 | 130 |
| **24b** |  | i.a. | i.a. | 150 |
| **24c** |  | 5.6 | 6.5 | 20 |
| **24d** |  | 4.1 | i.a. | 170 |
| **24e** |  | 4.9 | 5.6 | 130 |
| **24f** |  | <4 | 5.1 | 130 |

<sup>a</sup>pIC<sub>50</sub> from at least 10-point dose response curve. Values are the mean of triplicate determinations with variance <0.1. <sup>b</sup>pEC<sub>50</sub> from an 8-point dose response curve. Values are the average of duplicate determinations. <sup>c</sup>kinetic solubility of a 10 mM DMSO solution at pH 7.4. i.a. = inactive at 100 μM in the nsP2 protease assay and at 10 μM in the CHIKV-nLuc assay. <sup>d</sup>1:3 mixture of allyl:vinyl regioisomers.

Small changes to the vinyl sulfone showed that cysteine protease inhibition was very sensitive to modification of the warhead (Table 10). The cyclopropyl sulfone **25a** had 10-fold lower activity compared to the methyl sulfone **1a** and the isopropyl sulfone **25b** was a further 5-fold less active as an nsP2 protease inhibitor. The cyclopentyl sulfone **25c** showed an additional decrease in enzyme inhibition. These data highlight the tight steric tolerance for variation of the alkyl substituent on the vinyl sulfone. The warhead was more tolerant to the addition of aryl substituents onto the sulfone. The phenyl sulfone **25d** was a good inhibitor of the protease but had decreased activity in the antiviral assay. Switching to the 4-methoxyphenyl sulfone **25e** restored antiviral activity. However, the 2-fluorophenyl (**25f**) and 2-pyridyl (**25g**) sulfones had weaker activity in the antiviral assay despite maintaining sub-micromolar nsP2 protease inhibition. Notably, several of the modified vinyl sulfones (**25c–g**) had lower aqueous solubility than the methyl sulfone **1a** suggesting that in some cases (e.g. **25d** and **25f**) their physiochemical properties may negatively impact antiviral activity in cells.

**Table 10.**
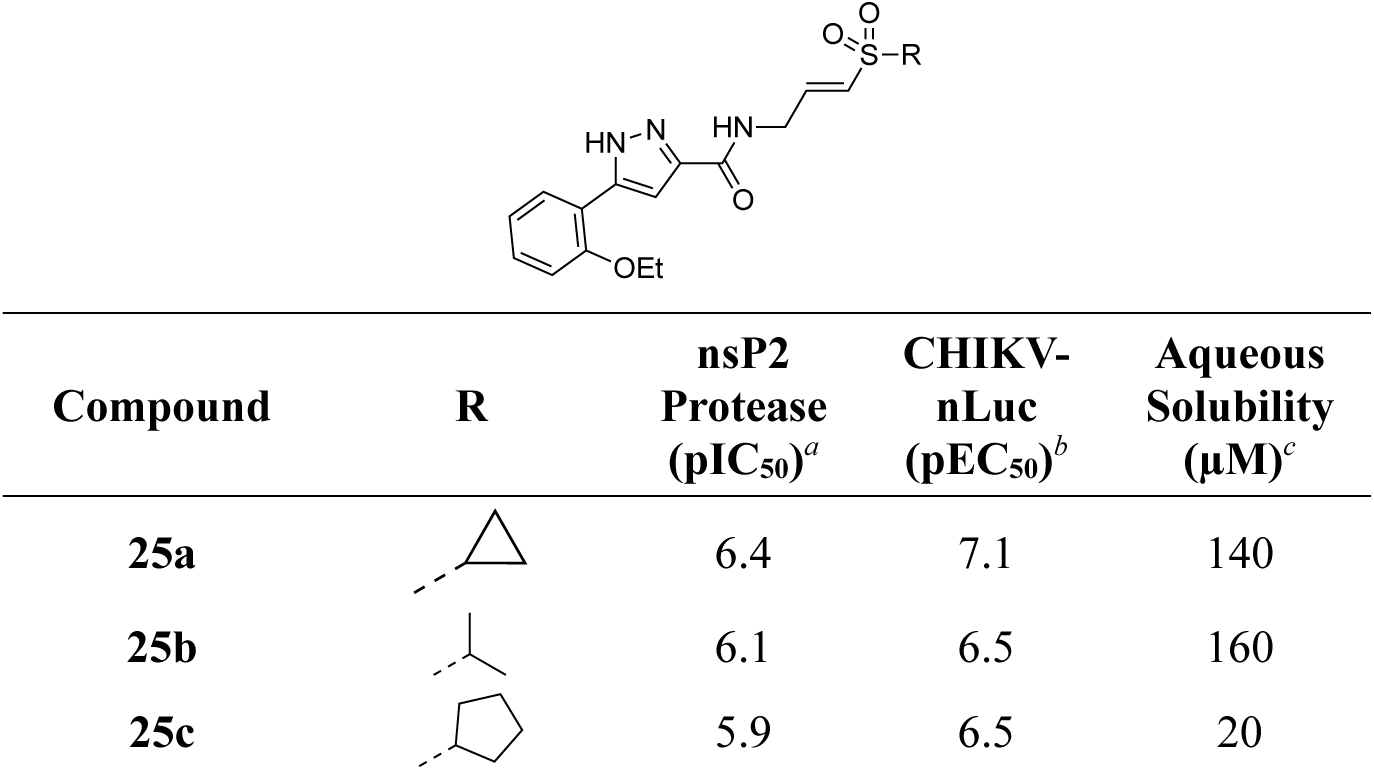
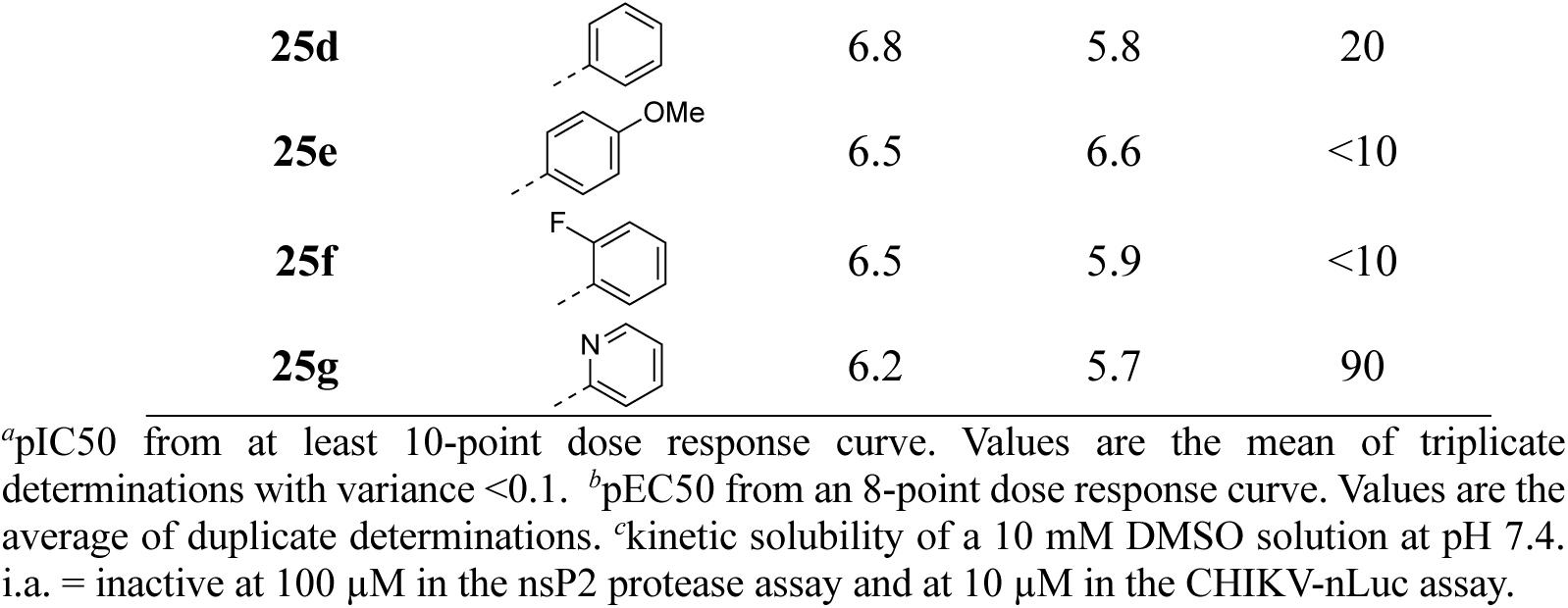
nsP2 Protease Inhibition, Antiviral Activity, and Solubility of 25a–g.

|  |  |  |  |  |
| --- | --- | --- | --- | --- |
| Compound | R | nsP2<br>Protease<br>(pIC<sub>50</sub>)<sup>a</sup> | CHIKV-nLuc<br>(pEC<sub>50</sub>)<sup>b</sup> | Aqueous<br>Solubility<br>(μM)<sup>c</sup> |
| **25a** |  | 6.4 | 7.1 | 140 |
| **25b** |  | 6.1 | 6.5 | 160 |
| **25c** |  | 5.9 | 6.5 | 20 |
| **25d** |  | 6.8 | 5.8 | 20 |
| **25e** |  | 6.5 | 6.6 | <10 |
| **25f** |  | 6.5 | 5.9 | <10 |
| **25g** |  | 6.2 | 5.7 | 90 |

<sup>a</sup>pIC<sub>50</sub> from at least 10-point dose response curve. Values are the mean of triplicate determinations with variance <0.1. <sup>b</sup>pEC<sub>50</sub> from an 8-point dose response curve. Values are the average of duplicate determinations. <sup>c</sup>kinetic solubility of a 10 mM DMSO solution at pH 7.4. i.a. = inactive at 100 μM in the nsP2 protease assay and at 10 μM in the CHIKV-nLuc assay.

Changing the vinyl sulfone warhead of **1a** to vinyl sulfoxide **26** resulted in loss of nsP2 protease inhibition (Table 11). However, in the isoxazole series the racemic vinyl sulfoximine **27** demonstrated only a 10–30-fold decrease in activity compared to the vinyl sulfone **10**. The corresponding pyrazole could not be synthesized due to intramolecular cyclization.

**Table 11.**
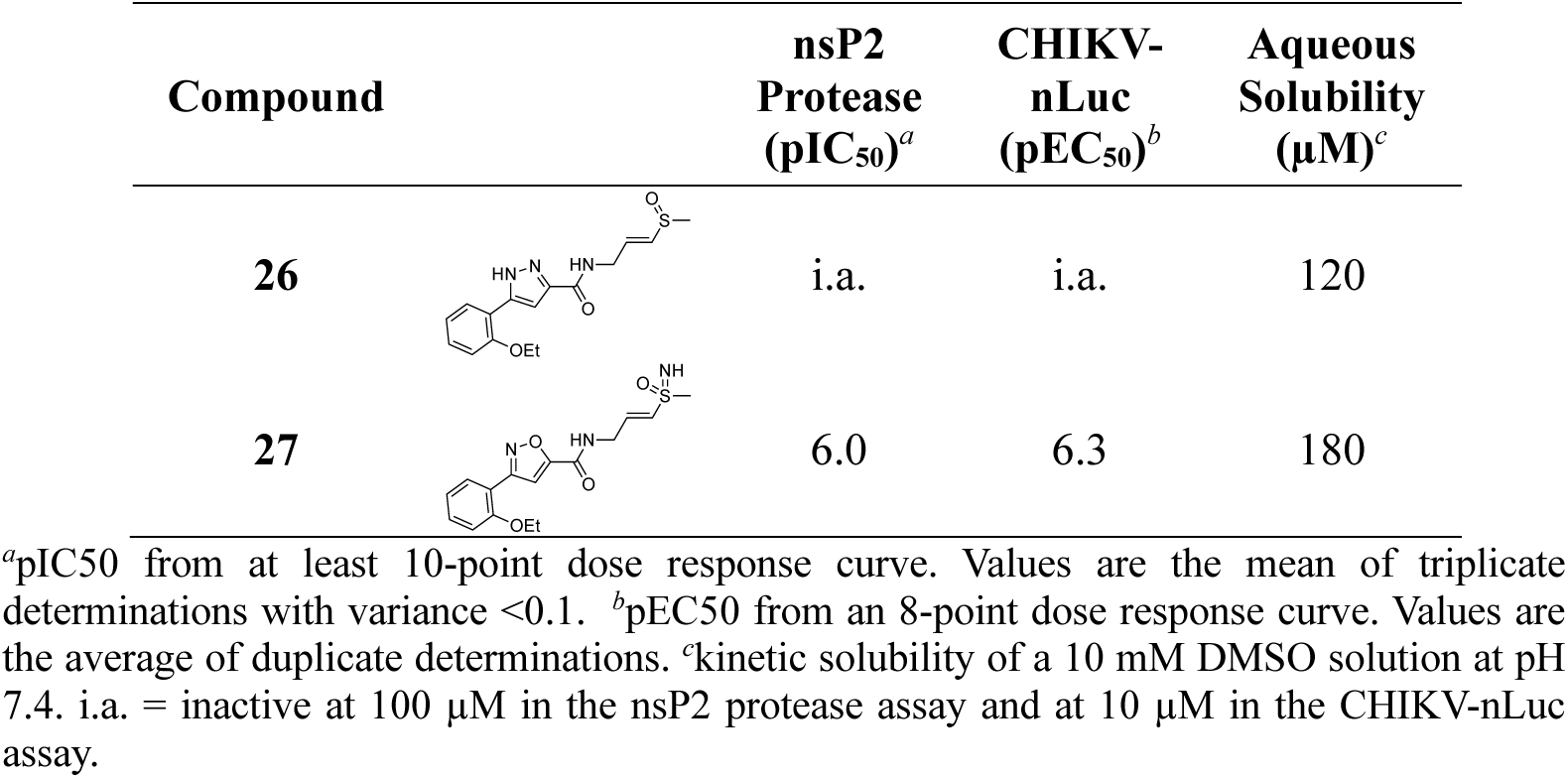
nsP2 Protease Inhibition, Antiviral Activity, and Solubility of 26–27.

| Compound |  | nsP2<br>Protease<br>(pIC<sub>50</sub>)<sup>a</sup> | CHIKV-nLuc<br>(pEC<sub>50</sub>)<sup>b</sup> | Aqueous<br>Solubility<br>(μM)<sup>c</sup> |
| --- | --- | --- | --- | --- |
| **26** |  | i.a. | i.a. | 120 |
| **27** |  | 6.0 | 6.3 | 180 |

<sup>a</sup>pIC<sub>50</sub> from at least 10-point dose response curve. Values are the mean of triplicate determinations with variance <0.1. <sup>b</sup>pEC<sub>50</sub> from an 8-point dose response curve. Values are the average of duplicate determinations. <sup>c</sup>kinetic solubility of a 10 mM DMSO solution at pH 7.4. i.a. = inactive at 100 μM in the nsP2 protease assay and at 10 μM in the CHIKV-nLuc assay.

An alternative series of non-sulfone covalent warheads was also explored (Table 12). The vinyl ester **28** had reduced activity and the vinyl amide **29** was inactive. The nitrile **30** was also inactive. However, the pyrrolidinyl nitrile **31** showed modest inhibition of the protease and sub-micromolar antiviral activity. These changes confirmed that the β-amidomethyl vinyl sulfone had superior enzyme inhibition and was a privileged covalent warhead for the nsP2 cysteine proteases.

**Table 12.**
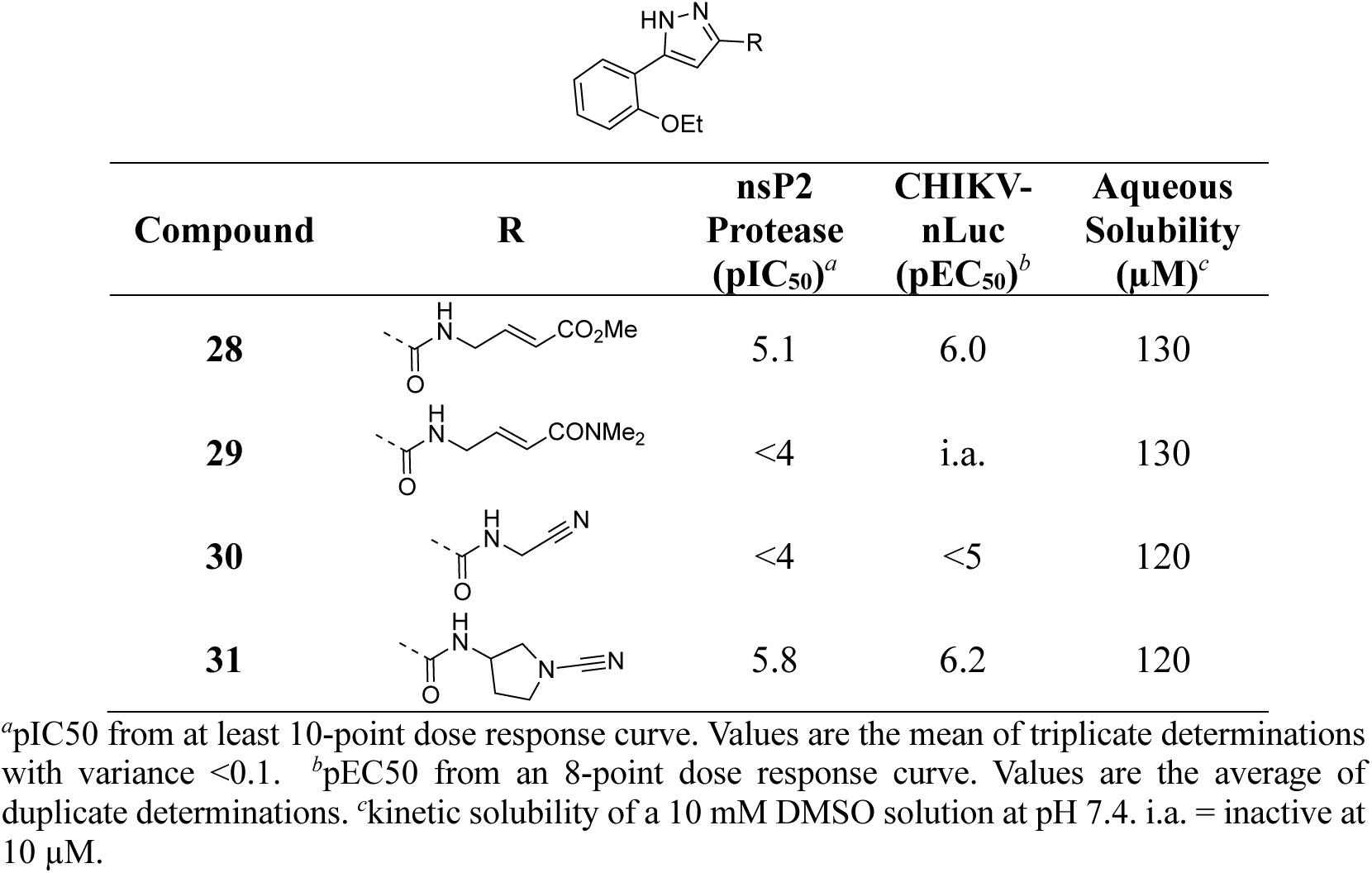
nsP2 Protease Inhibition, Antiviral Activity, and Solubility of 28–31.

| Compound | R | nsP2<br>Protease<br>(pIC<sub>50</sub>)<sup>a</sup> | CHIKV-nLuc<br>(pEC<sub>50</sub>)<sup>b</sup> | Aqueous<br>Solubility<br>(μM)<sup>c</sup> |
| --- | --- | --- | --- | --- |
| **28** |  | 5.1 | 6.0 | 130 |
| **29** |  | <4 | i.a. | 130 |
| **30** |  | <4 | <5 | 120 |
| **31** |  | 5.8 | 6.2 | 120 |

<sup>a</sup>pIC<sub>50</sub> from at least 10-point dose response curve. Values are the mean of triplicate determinations with variance <0.1. <sup>b</sup>pEC<sub>50</sub> from an 8-point dose response curve. Values are the average of duplicate determinations. <sup>c</sup>kinetic solubility of a 10 mM DMSO solution at pH 7.4. i.a. = inactive at 10 μM.

### Structure-activity of Covalent nsP2 Protease Inhibition

To further elucidate the structure-activity relationships within the series and to understand the contribution of the covalent warhead to the inhibition of nsP2 protease, several of the β-amidomethyl vinyl sulfones were selected for measurement of their binding kinetics and efficiency of enzyme inactivation (Table S1). *k_obs_* values over serial-dilutions of inhibitor were determined and fit with the equation (*k_obs_* = (*k_inact_* x [I]) / (*K_i_* + [I]) to determine the potency of the initial binding event (*K_i_*) and maximum potential rate of covalent bond formation (*k_inact_*).

The resulting *k_inact_/K_i_* ratios represent the efficiency of the covalent bond formation between the nsP2 cysteine protease and the vinyl sulfones (Table 13). Only one of the inhibitors (**20**) had a *k_inact_/K_i_* ratio below 1000. The other 17 inhibitors showed rapid inactivation of the enzyme, with 4-cyanopyrazole (**8d**) producing a *k_inact_/K_i_* >10,000 which marks it as a highly efficient non-peptide covalent cysteine protease inhibitor.^14^

**Table 13.**
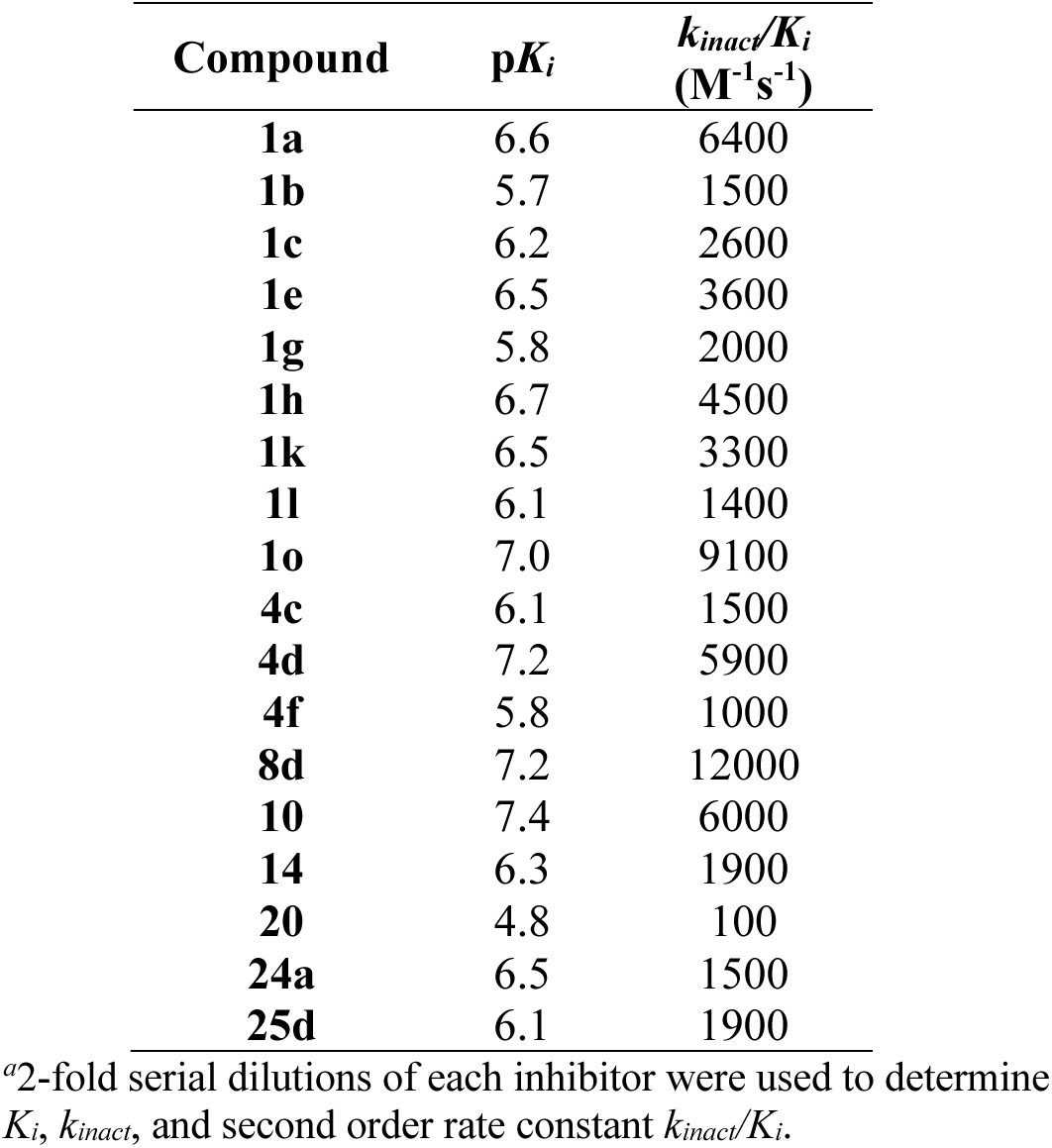
Binding and Kinetics of nsP2 Enzyme Inactivation*^a^*.

Analysis of the p*K_i_* data (Table 13) provided evidence that the structure of the heterocycle and its substituents influenced the recognition of the inhibitor by the enzyme. Comparison of p*K_i_* with nsP2 protease pIC_50_ showed that compounds with the best binding constant had the highest potency for enzyme inhibition (Figure 2A). However, several compounds fell above or below the correlation line (*R*^2^ = 0.70) suggesting that binding alone did not account for all variation in inhibition potency. A slightly stronger correlation was observed with the *k_inact_/K_i_* ratio (Figure 2B, *R*^2^ = 0.81) indicating that the efficiency of enzyme inactivation by the covalent warhead was a dominant factor in driving the potency of enzyme inhibition.

**Figure 2.**
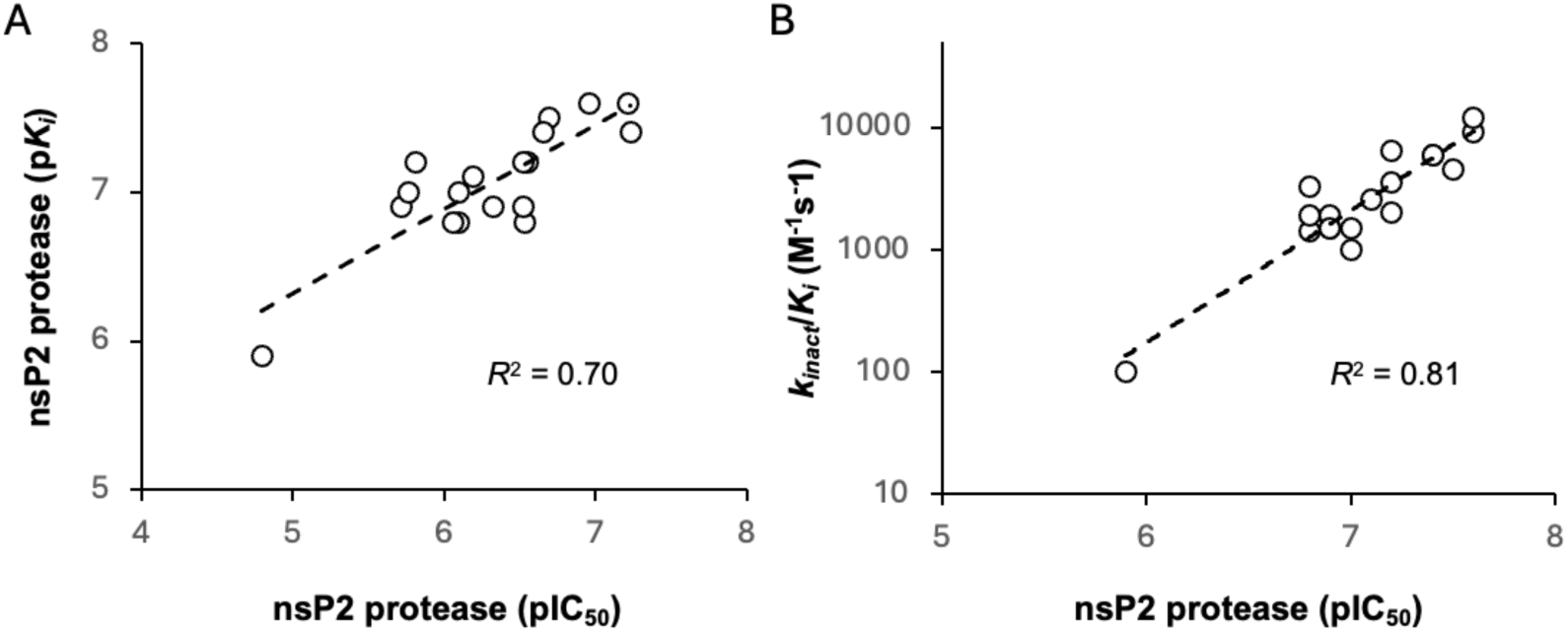
Relationship between nsP2 protease inhibition and kinetic parameters for binding and inactivation. A. p*K_i_* vs pIC_50_. B. *k_inact_/K_i_* vs pIC_50_. *R*^2^ was calculated from the trendline using Microsoft Excel v16.

Comparison of the potency of enzyme inhibition with antiviral activity across the full set of inhibitors for which IC_50_ and EC_50_ values could be determinized (Figure 3) gave only a modest correlation (*R*^2^ = 0.25). However, it is still evident that enzyme inhibition was a primary driver of antiviral activity for these covalent inhibitors, with the most potent nsP2 protease inhibitors showing the best antiviral activity. The efficiency of cell penetration may contribute to the discrepancy between biochemical enzyme inhibition and expression of antiviral activity in the replicon assay. Notably, several of the compounds that fell below the correlation line contained additional H-bond donors in the pyrazole 5-substituent (e.g. **7b–d**) that may contribute to poorer cell penetration and lower potency in the antiviral assay than would be predicted solely from their enzyme inhibition.

**Figure 3.**
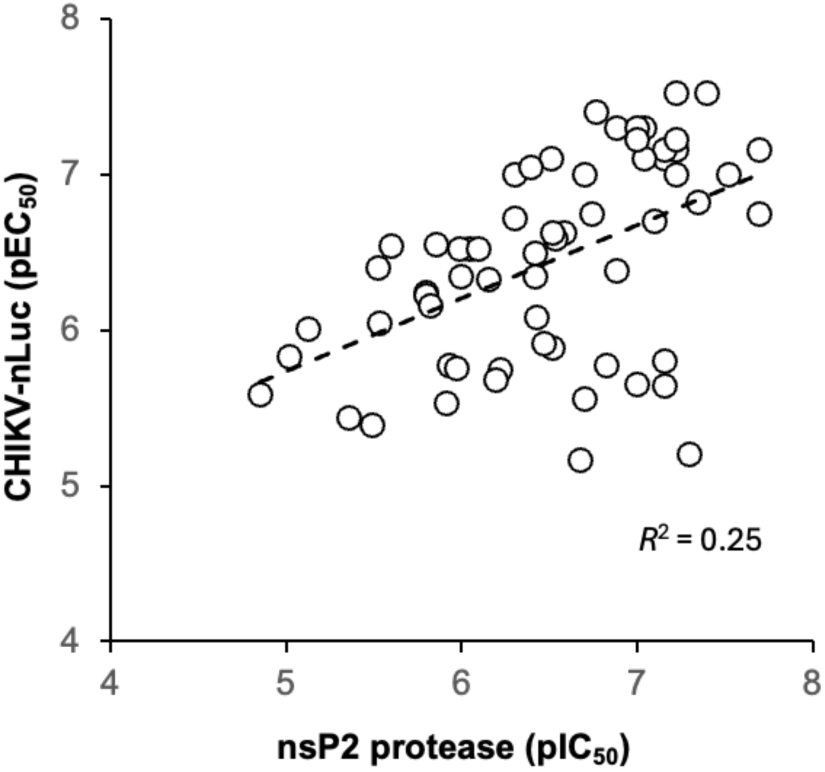
Relationship between nsP2 protease inhibition and antiviral activity. Those compounds where an nsP2 protease IC_50_ or CHIKV-nLuc EC_50_ could not be determined were excluded from the correlation. *R*^2^ was calculated from the trendline using Microsoft Excel v16.

The nsP2 cysteine protease cleaves the CHIKV viral polyprotein at three sites to generate the non-structural proteins nsP1–4. Alignment of the substrate cleavage sites^15^ showed that the enzyme recognized a conserved P_1_–P_4_ sequence (Figure 4A). The P_1_, P_2_, and P_3_ amino acids are dominated by alanine and glycine, which would predict that the complimentary substrate binding sites (S_1_–S_3_) could only accommodate small amino acids. The P_4_ amino acid is restricted to arginine, which implies that the S_4_ pocket may contain an acidic amino acid to allow formation of a salt bridge. In contrast the (S_-1_–S_-4_) binding sites should be less restrictive given the wide variation in amino acids in the P_-1_–P_-4_ substrate sequences. By combining these insights into substrate recognition with the ligand-base structure activity contained in Tables 1–13, a simple model was built of the potential enzyme active site (Figure 4B). Annotation of the binding site model with the CHIKV nsP2 substrate binding pockets (S_4_–S_-2_) showed a potential alignment with the inhibitor structure-activity. In the model, the 5-membered heterocycle was proposed to sit in a tight S_3_ pocket, which prefers only an alanine sidechain and may account for the inability to accommodate an additional methyl substituent on the heterocycle. In contrast, the wide range of active analogs with lipophilic substituents at the heterocycle 5-position might fit into a large S_4_ pocket. However, it is possible that additional binding affinity might be found if a basic substituent could be identified that was complimentary to the arginine substrate pocket in S_4_. The β-amidomethyl vinyl sulfone was proposed to sit in the S_2_ and S_1_ pockets, which allowed some small alkyl substitutions in the inhibitor. Notably, sulfone substituents larger than methyl could be potentially accommodated in the S_-1_ and S_-2_ pockets that are less restrictive in their substrate preferences. In the absence of experimental structural information of the inhibitor-enzyme complex, this ligand-based model (Figure 4B) may be useful in the design of new covalent nsP2 protease inhibitors.

**Figure 4.**
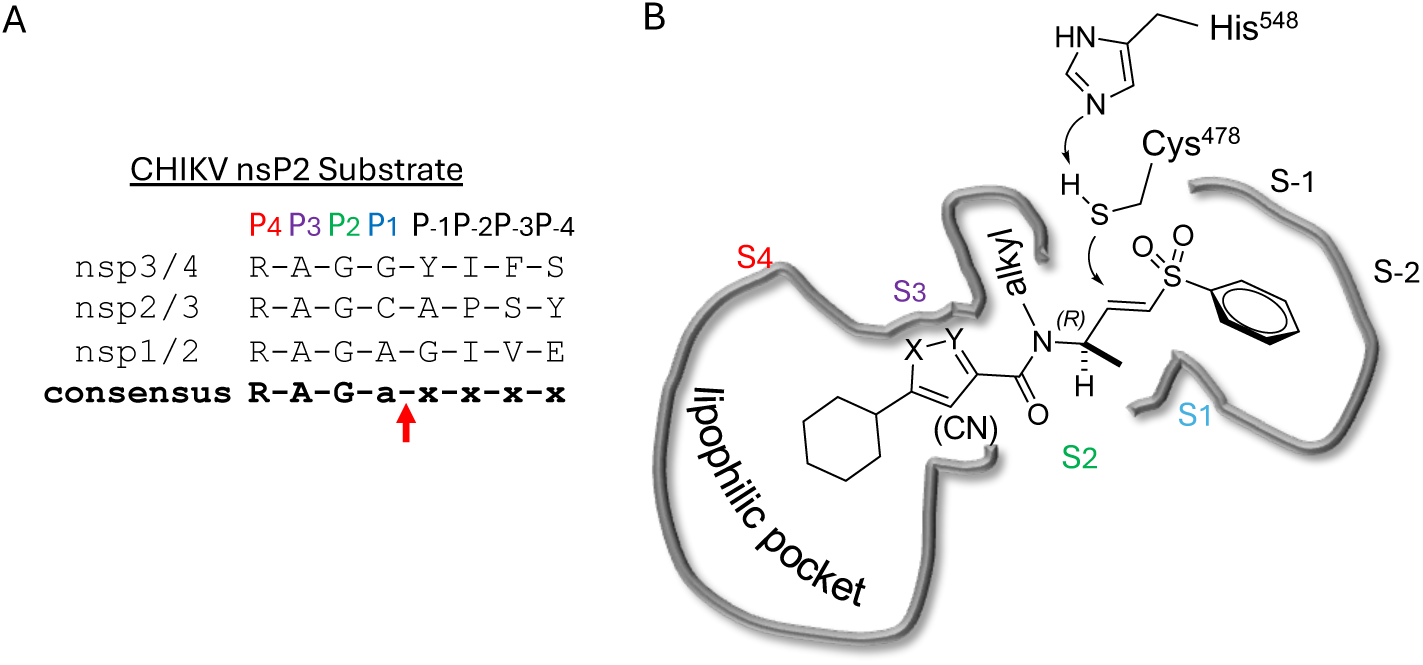
Model of the CHIKV nsP2 protease active site. A. CHIKV nsP2 substrate preference. The protease cleavage site of the viral polyproteins is indicated with a red arrow. B. Ligand-based model of the enzyme active site encompassing SAR data from Tables 1–12. The proposed substrate pockets are marked as S_4_–S_-2_.

## Conclusion

In summary, our exploration of three regions of the heterocyclic β-aminomethyl vinyl sulfone **1a** defined the structure-activity relationship for inhibition of CHIKV nsP2 protease and demonstrated the critical contribution of the covalent warhead for enzyme inhibition and antiviral activity. In general, the heterocyclic β-aminomethyl vinyl sulfones had good physiochemical properties, often with aqueous solubilities >100 μM. Notably, analogs with anti-CHIKV activity also showed anti-VEEV activity (Table S2) confirming the broad-spectrum potential of these covalent nsP2 inhibitors.^7^ Although the warhead was very sensitive to modification of either the sulfone or vinyl substituents, many changes to the core 5-membered heterocycle or its aryl substituent were identified that improved binding to nsP2 such as the 4-cyanopyrazole analog **8d** that demonstrated a *k_inact_/K_i_* >10,000 (Table 13). Modification of the 5-membered heterocycle identified the 3-arylisoxazole (e.g. **10**) as an isosteric replacement that was no longer prone to the intramolecular cyclization that complicated the synthesis of the pyrazole-based inhibitors such as **1a**.^11^ The combined data were used to generate a simple model of the enzyme active site (Figure 4B) that will aid the design of covalent nsP2 protease inhibitors as potential anti-alphavirus therapeutics.

## Experimental Section

### General Information

All reactions were performed in oven-dried glassware under an atmosphere of dry N_2_ unless otherwise stated. All reagents and solvents used were purchased from commercial sources and were used without further purification. No unexpected safety hazards were encountered during chemical synthesis. Analytical thin layer chromatography (TLC) was performed on pre-coated silica gel plates, 200 μm with an F254 indicator. TLC plates were visualized by fluorescence quenching under UV light or by staining with iodine and KMnO_4_. Column chromatography was performed using pre-loaded silica gel cartridges on a Biotage automated purification system. ^1^H and ^13^C NMR spectra were collected in CDCl_3_, MeOD, or DMSO-*d*_6_ at 400/500 and 101/126/214 MHz on Bruker spectrometers. Characteristic chemical shifts (*δ*) are reported in parts per million (ppm) downfield from tetramethylsilane (for ^1^H NMR) with designation of major peaks as s (singlet), d (doublet), t (triplet), q (quartet), m (multiplet), dd (doublet of doublets), dt (doublet of triplets), td (triplet of doublets), or ddd (doublet of doublets of doublets). HRMS samples were analyzed at the UNC Department of Chemistry Mass Spectrometry Core Laboratory with a Q Exactive HF-X mass spectrometer. LCMS were run on an Agilent 1290 Infinity II LC System using an Agilent Infinity Lab PoroShell 120 EC-C18 column (30 °C, 2.7 μm, 2.1 × 50 mm), eluent 5−95% CH_3_CN in water; modifier formic acid 0.2% (v/v) concentration, flow rate of 1 mL/min. Preparative HPLC was performed using an Agilent 1260 Infinity II LC System equipped with a Phenomenex C18 column (PhenylHexyl, 30 °C, 5 μm, 75 x 30 mm), eluent 5−95% CH_3_CN in water; modifier trifluoroacetic acid 0.05% (v/v) concentration, flow rate of 30 mL/min. Analytical HPLC data were recorded on Waters Alliance HPLC with PDA detector or Agilent 1260 Infinity II series with PDA detector (EC-C18, 100 mm x 4.6 mm, 3.5µm), eluent 10−90% CH_3_CN in water, flow rate of 1 mL/min. Chiral SFC separation was performed with Shimadzu LC-20AP preparative pump with UV-detector at 250 nm, mobile phase: (A) 0.1% NH_3_ in hexane, (B) 0.1% NH_3_ in IPA:ACN using a Chiralpak IC column (250 x 30 mm, particle size 5 µm), flow rate: 25 mL/min. Optical rotations were measured with an Autopol IV Automatic Polarimeter from Rudolph Research using a 10 cm microcell. All compounds were >95% pure by HPLC analysis. Additional QC of pyrazole analogs that had the potential for intramolecular cyclization was performed by HPLC prior to bioassay (Figures S154–S206).

### Chemistry

Methylamine (**XXVb**), 3-amino-2,3-dihydrothiophene 1,1-dioxide (**XXVc**), 2-aminoacetonitrile (**XXVf**), 3-aminopyrrolidine-1-carbonitrile (**XXVg**), analogs **4a**, **4b**, and **4f** were obtained from commercial sources. (*E*)-3-(methylsulfonyl)prop-2-en-1-amine (**V**),^16^ 3-(methylsulfonyl)propan-1-amine (**XXVa**),^17^ methyl (*E*)-4-aminobut-2-enoate (**XXVd**),^16^ (*E*)-4-amino-*N*,*N*-dimethylbut-2-enamide (**XXVe**),^18^ **IVa**–**q**^19^ were synthesized using the reported procedures.

### General Methods. General Procedure-I

To a stirred solution of acid (1.0 eq.) and 2-(1*H*-benzotriazole-1-yl)-1,1,3,3-tetramethylaminium tetrafluoroborate (TBTU) (1.5 eq.) in pyridine was added corresponding amine substituent (1.2 eq.) and the reaction was stirred at 25 °C for 2 h. On completion of the reaction (based on TLC and LCMS analysis), the reaction was poured into water, extracted with EtOAc, combined organic layers dried over anhydrous Na_2_SO_4_, filtered, and concentrated. The crude was purified by combiflash or preparative HPLC to afford the desired product.

### General Procedure-II

To a stirred a solution of acid (1.0 eq.) in DMF were added 1-[bis(dimethylamino)methylene]-1*H*-1,2,3-triazolo[4,5-*b*]pyridinium 3-oxid hexafluorophosphate (HATU) (1.5 eq.), *N*-ethyldiisopropylamine (DIPEA) (3.0 eq.) and corresponding amine substituent (1.1 eq.). The reaction mixture was stirred at 25 °C for 16 h. On completion of the reaction (based on TLC and LCMS), the reaction mixture was poured into water, extracted with EtOAc, washed with brine, and the combined organic layers dried over anhydrous Na_2_SO_4_, filtered, and concentrated. The crude product was purified by combiflash or preparative HPLC to afford the desired product.

### 22General Procedure-III

To a stirred solution of corresponding ester intermediate (1.0 eq.) in MeOH/THF/H_2_O was added lithium hydroxide monohydrate (3.0 eq.) and the reaction was stirred at 25 °C for 6 h. After completion (based on TLC and LCMS), the reaction mixture was concentrated under reduced pressure and the residue was acidified to pH 3 with 1N HCl. Organic layers were extracted with EtOAc, dried over anhydrous Na_2_SO_4_, filtered, and concentrated. The residue was washed with hexane to afford the desired product as a solid.

### General Procedure-IV

To a stirred solution of compound (1.0 eq.) in THF at 0 °C were added 3,4-dihydro-2*H*-pyran (1.5 eq.) and p-toluenesulfonic acid monohydrate (PTSA) (1.5 eq.). The reaction was stirred at 25 °C for 12 h. After completion, the reaction was diluted with water, extracted with EtOAc, washed with brine, dried with anhydrous Na_2_SO_4_, filtered, and concentrated. The resulting crude was purified by combiflash to furnish the desired THP-protected intermediate.

### General Procedure-V

To a stirred a solution of THP-or Boc-protected intermediate (1.0 eq.) in DCM (10 volumes) was added 4M HCl in dioxane (5 volumes) or trifluoroacetic acid (10 volumes) and the mixture stirred at 25 °C for 2–12 h. After completion, the reaction mixture was concentrated, and the resulting crude product was purified using preparative HPLC (if required) to afford the desired compound.

### Synthesis of 1a–q

Amide coupling of 5-aryl-1*H*-pyrazole-3-carboxylic acids **IVa**–**q** with (*E*)-3-(methylsulfonyl)prop-2-en-1-amine (**V**) following General Procedure-I afforded **1a**–**q**.

**(*E*)-5-(2-Ethoxyphenyl)-*N*-(3-(methylsulfonyl)allyl)-1*H*-pyrazole-3-carboxamide (1a).** White solid (44% yield, m.p 70 °C). ^1^H NMR (500 MHz, DMSO-*d*_6_): *δ* 8.62 (t, *J* = 5.9 Hz, 1H), 7.74 (dd, *J* = 7.6, 1.8 Hz, 1H), 7.34 (ddd, *J* = 8.5, 7.4, 1.7 Hz, 1H), 7.16 – 7.10 (m, 2H), 7.03 (td, *J* = 7.5, 1.1 Hz, 1H), 6.82 (dt, *J* = 15.2, 4.4 Hz, 1H), 6.69 (dt, *J* = 15.2, 1.8 Hz, 1H), 4.16 (q, *J* = 6.9 Hz, 2H), 4.11 (ddd, *J* = 6.1, 4.4, 1.8 Hz, 2H), 3.01 (s, 3H), 1.41 (t, *J* = 6.9 Hz, 3H). ^13^C NMR (DMSO-*d*_6_, 126 MHz): *δ* 161.5, 155.0, 143.5, 130.0, 129.7, 127.8, 120.6, 112.8, 105.6, 63.7, 42.2, 38.7, 14.6. HRMS (ESI) *m/z*: [M+H]^+^ calculated for C_16_H_20_N_3_O_4_S: 350.1175; found 350.1172. HPLC purity (254 nm) >99%.

**(*E*)-*N*-(3-(Methylsulfonyl)allyl)-3-phenyl-1*H*-pyrazole-5-carboxamide (1b).** White sticky solid (26% yield). ^1^H NMR (400 MHz, DMSO-*d*_6_): *δ* 13.70 (s, 1H), 8.75 (d, *J* = 116.9 Hz, 1H), 7.80 (d, *J* = 7.7 Hz, 2H), 7.48 (t, *J* = 7.6 Hz, 2H), 7.41 – 7.29 (m, 1H), 7.10 (s, 1H), 6.85 – 6.63 (m, 2H), 4.13 (d, *J* = 17.1 Hz, 2H), 3.01 (s, 3H). ^13^C NMR (176 MHz, DMSO-*d*_6_): *δ* 161.8, 147.4, 143.5, 130.0, 129.1, 128.5, 125.3, 102.8, 42.1, 38.7. HRMS (ESI) *m/z*: [M+H]^+^ calculated for C_14_H_16_N_3_O_3_S: 306.0912; found 306.0701. HPLC purity (254 nm) >99%.

**(*E*)-5-(2-Chlorophenyl)-*N*-(3-(methylsulfonyl)allyl)-1*H*-pyrazole-3-carboxamide (1c).** White solid (40% yield, m.p. 140–142 °C). ^1^H NMR (400 MHz, DMSO-*d*_6_): *δ* 8.79 (s, 1H), 7.74 (d, *J* = 8.3 Hz, 1H), 7.58 (td, *J* = 7.4, 1.9 Hz, 1H), 7.48 – 7.44 (m, 1H), 7.43 – 7.39 (m, 1H), 7.17 (d, *J* = 27.2 Hz, 1H), 6.82 (dt, *J* = 15.2, 4.1 Hz, 1H), 6.73 (d, *J* = 15.4 Hz, 1H), 4.13 (ddd, *J* = 6.0, 4.2, 1.6 Hz, 2H), 3.01 (s, 3H). ^13^C NMR (176 MHz, DMSO-*d*_6_): *δ* 143.2, 131.1, 130.4(9), 130.4(6), 130.4, 130.3, 130.2, 127.6, 127.5, 108.8, 106.2, 42.1, 38.7. HRMS (ESI) *m/z*: [M+H]^+^ calculated for C_14_H_15_ClN_3_O_3_S: 340.0523; found 340.0520. HPLC purity (254 nm) >99%.

**(*E*)-5-(2-Hydroxyphenyl)-*N*-(3-(methylsulfonyl)allyl)-1*H*-pyrazole-3-carboxamide (1d)**. White solid (47% yield, m.p. 188–189 °C). ^1^H NMR (500 MHz, DMSO-*d*_6_): *δ* 10.28 (s, 1H), 8.65 (s, 1H), 7.67 (dd, *J* = 7.7, 1.7 Hz, 1H), 7.19 (ddd, *J* = 8.1, 7.3, 1.7 Hz, 1H), 6.97 (dd, *J* = 8.3, 1.2 Hz, 1H), 6.90 (td, *J* = 7.6, 1.2 Hz, 1H), 6.82 (dt, *J* = 15.2, 4.3 Hz, 1H), 6.71 (d, *J* = 15.5 Hz, 1H), 4.14 – 4.09 (m, 2H), 3.01 (s, 3H). ^13^C NMR (126 MHz, DMSO-*d*_6_): *δ* 161.1, 154.4, 143.4, 130.1, 129.3, 127.3, 119.4, 116.4, 104.5, 42.2, 38.8. HRMS (ESI) *m/z*: [M+H]^+^ calculated for C_14_H_16_N_3_O_4_S: 322.0862, found: 322.0859; HPLC (Method A): *t*_ret_ 3.13 min, purity (254 nm) >99%.

**(*E*)-*N*-(3-(Methylsulfonyl)allyl)-3-(o-tolyl)-1*H*-pyrazole-5-carboxamide (1e).** White solid (36% yield). ^1^H NMR (500 MHz, DMSO-*d*_6_): *δ* 13.41 (s, 1H), 8.69 (s, 1H), 7.50 – 7.45 (m, 1H), 7.33 – 7.29 (m, 3H), 6.92 (s, 1H), 6.82 (dt, *J* = 15.2, 4.3 Hz, 1H), 6.72 (dt, *J* = 15.1, 1.8 Hz, 1H), 4.12 (ddd, *J* = 6.1, 4.3, 1.8 Hz, 2H), 3.02 (s, 3H), 2.39 (s, 3H). ^13^C NMR (214 MHz, DMSO-*d*_6_): *δ* 143.4, 135.5, 135.4, 130.9, 130.8, 130.1, 128.9, 128.8, 128.7, 128.4, 126.1, 126.0, 107.9, 105.4, 42.1, 38.7, 20.6. HRMS (ESI) *m/z*: [M+H]^+^ calculated for C_15_H_18_N_3_O_3_S: 320.1069; found 320.1065. HPLC purity (254 nm) >99%.

**(*E*)-*N*-(3-(Methylsulfonyl)allyl)-5-(2-(trifluoromethyl)phenyl)-1*H*-pyrazole-3-carboxamide (1f).** White solid (32% yield, m.p. 174 °C). ^1^H NMR (400 MHz, CD_3_CN): *δ* 11.54 (s, 1H), 6.89 (d, *J* = 4.0 Hz, 1H), 7.75 – 7.64 (m, 3H), 6.93 – 6.87 (m, 2H), 6.63 (d, *J* = 16 Hz, 1H), 4.22 (q, *J*_1_= 4 Hz, *J*_2_= 4 Hz, 2H), 2.91 (s, 3H). ^13^C NMR (101 MHz, CD_3_CN): *δ* 163.3, 159.1, 158.3, 145.2, 144.2, 133.4, 131.3, 129.3, 126.4, 123.7, 120.1, 117.3, 111.6, 107.8, 42.9, 40.2. HRMS (ESI) *m/z*: [M+H]^+^ calculated for C_15_H_15_F_3_N_3_O_3_S: 374.0786, found 374.0786. HPLC purity (254 nm) 97%.

**(*E*)-5-(3-Methoxyphenyl)-*N*-(3-(methylsulfonyl)allyl)-1*H*-pyrazole-3-carboxamide (1g).** White solid (29% yield, m.p. 198–200 °C). ^1^H NMR (500 MHz, DMSO-*d*_6_): *δ* 13.69 (s, 1H), 8.64 (s, 1H), 7.37 (s, 3H), 7.18 (s, 1H), 6.98 – 6.78 (m, 2H), 6.71 (d, *J* = 16.6 Hz, 1H), 4.12 (s, 2H), 3.81 (s, 3H), 3.01 (s, 3H). ^13^C NMR (126 MHz, DMSO-*d*_6_): *δ* 161.7, 159.7, 147.3, 143.4, 130.1, 130.1, 117.5, 114.1, 110.5, 102.9, 55.2, 42.1, 38.7. HRMS (ESI) *m/z*: [M+H]^+^ calculated for C_15_H_18_N_3_O_4_S: 336.1018; found 336.1020. HPLC purity (254 nm) >99%.

**(*E*)-5-(3-Ethoxyphenyl)-*N*-(3-(methylsulfonyl)allyl)-1*H*-pyrazole-3-carboxamide (1h).** White solid (40% yield). ^1^H NMR (400 MHz, DMSO-*d*_6_): *δ* 13.66 (s, 1H), 8.61 (s, 1H), 7.38 – 7.33 (m, 3H), 7.14 (s, 1H), 6.92 (s, 1H), 6.81 (dt, *J* = 15.0, 4.3 Hz, 1H), 6.71 (s, 1H), 4.09 (q, *J* = 6.5 Hz, 4H), 3.01 (s, 3H), 1.35 (t, *J* = 7.0 Hz, 3H). ^13^C NMR (214 MHz, DMSO-*d*_6_): *δ* 161.8, 158.9, 157.7, 157.6, 147.3, 143.4, 130.2, 130.0, 117.5, 114.6, 111.1, 103.0, 63.2, 42.1, 38.7, 14.6. HRMS (ESI) *m/z*: [M+H]^+^ calculated for C_16_H_20_N_3_O_4_S: 350.1175, found 350.1178. HPLC purity (254 nm) >99%.

**(*E*)-5-(3-Chlorophenyl)-*N*-(3-(methylsulfonyl)allyl)-1*H*-pyrazole-3-carboxamide (1i).** White solid (20% yield, m.p. 200 °C). ^1^H NMR (500 MHz, DMSO-*d*_6_): *δ* 13.79 (s, 1H), 8.76 (d, *J* = 93.1 Hz, 1H), 7.98 – 7.71 (m, 2H), 7.54 – 7.37 (m, 2H), 7.25 (s, 1H), 6.85 – 6.62 (m, 2H), 4.12 (s, 2H), 3.01 (s, 3H). ^13^C NMR (126 MHz, DMSO-*d*_6_): *δ* 161.6, 158.6, 149.2, 147.5, 143.5, 142.8, 142.1, 138.3, 135.2, 133.9, 130.9, 130.7, 130.3, 130.1, 128.2, 127.5, 124.9, 124.6, 124.0, 123.6, 103.7, 102.7, 42.1, 38.7. HRMS (ESI) *m/z*: [M+H]^+^ calculated for C_14_H_15_ClN_3_O_3_S: 340.0523, found 340.0513. HPLC purity (254 nm) >99%.

**(*E*)-5-(4-Ethoxyphenyl)-*N*-(3-(methylsulfonyl)allyl)-1*H*-pyrazole-3-carboxamide (1j).** White solid (37% yield). ^1^H NMR (500 MHz, MeOD): *δ* 7.62 (d, *J* = 8.3 Hz, 2H), 7.01 – 6.97 (m, 2H), 6.97 – 6.92 (m, 2H), 6.70 (d, *J* = 14.9 Hz, 1H), 4.24 (dd, *J* = 4.4, 1.9 Hz, 2H), 4.08 (q, *J* = 6.9 Hz, 2H), 2.98 (s, 3H), 1.41 (t, *J* = 7.0 Hz, 4H). ^13^C NMR (100 MHz, DMSO-*d*_6_): *δ* 161.9, 158.7, 147.3, 143.6, 130.0, 126.8, 121.3, 114.9, 101.8, 63.2, 42.2, 38.7, 14.6. HRMS (ESI) *m/z*: [M+H]^+^ calculated for C16H_20_N_3_O_4_S: 350.1175, found 350.1171. HPLC purity (254 nm) >99%.

**(*E*)-*N*-(3-(Methylsulfonyl)allyl)-5-(4-(methylthio)phenyl)-1*H*-pyrazole-3-carboxamide (1k)**. White solid (29% yield). ^1^H NMR (500 MHz, MeOD): *δ* 7.64 (d, *J* = 8.0 Hz, 2H), 7.35 (d, *J* = 8.1 Hz, 2H), 7.02 (s, 1H), 6.95 (dt, *J* = 15.2, 4.5 Hz, 1H), 6.71 (d, *J* = 15.2 Hz, 1H), 4.24 (dd, *J* = 4.5, 1.9 Hz, 1H), 2.98 (s, 2H), 2.52 (s, 2H). ^13^C NMR (126 MHz, MeOD): *δ* 161.8, 147.4, 143.5, 143.2, 138.8, 130.0, 126.2, 126.1, 125.8, 125.5, 125.2, 102.5, 42.1, 38.7, 14.5. HRMS (ESI) *m/z*: [M+H]^+^ calculated for C_15_H_18_N_3_O_3_S_2_: 352.0790, found 352.0789. HPLC purity (254 nm) >99%.

**(*E*)-5-(4-(Dimethylamino)phenyl)-*N*-(3-(methylsulfonyl)allyl)-1*H*-pyrazole-3-carboxamide (1l).** White solid (27% yield). ^1^H NMR (500 MHz, MeOD): *δ* 7.85 (d, *J* = 8.1 Hz, 2H), 7.42 (d, *J* = 9.1 Hz, 2H), 7.09 (s, 1H), 6.95 (dt, *J* = 15.2, 4.5 Hz, 1H), 6.71 (dt, *J* = 15.2, 1.9 Hz, 1H), 4.24 (dd, *J* = 4.5, 1.9 Hz, 2H), 3.23 (s, 6H), 2.98 (s, 3H). ^13^C NMR (126 MHz, DMSO-*d*_6_): *δ* 167.5, 159.3, 152.9, 139.5, 135.7, 122.3, 110.5, 64.4, 51.6, 49.7, 48.2. HRMS (ESI) *m/z*: [M+H]^+^ calculated for C_16_H_21_N_4_O_3_S: 349.1334, found 349.1335. HPLC purity (254 nm) >99%.

**(*E*)-*N*-(3-(Methylsulfonyl)allyl)-5-(4-(trifluoromethyl)phenyl)-1*H*-pyrazole-3-carboxamide (1m).** White solid (36% yield, m.p. 241–242 °C). ^1^H NMR (400 MHz, DMSO-*d*_6_): *δ* 13.93 (s, 1H), 8.81 (d, *J* = 93.1 Hz, 1H), 8.02 (d, *J* = 7.9 Hz, 2H), 7.82 (d, *J* = 8.0 Hz, 2H), 7.34 (s, 1H), 6.91 – 6.63 (m, 2H), 4.14 (s, 2H), 3.02 (s, 3H). ^13^C NMR (214 MHz, DMSO-*d*_6_): *δ* 161.6, 158.7, 149.3, 147.7, 143.4, 142.8, 142.1, 138.5, 137.1, 132.6, 130.2, 128.3, 125.9, 124.9, 123.6, 122.3, 118.1, 104.2, 103.0, 42.2, 38.8. HRMS (ESI) *m/z*: [M+H]^+^ calculated for C_15_H_15_F_3_N_3_O_3_S: 374.0786, found 374.0782. HPLC purity (254 nm) >99%.

(***E*)-5-(2,4-Dimethylphenyl)-*N*-(3-(methylsulfonyl)allyl)-1*H*-pyrazole-3-carboxamide (1n).** White solid (40% yield, m.p. 177–178 °C). ^1^H NMR (500 MHz, DMSO-*d*_6_): *δ* 13.39 (s, 1H), 8.61 (s, 1H), 7.36 (s, 1H), 7.19 – 7.04 (m, 2H), 6.90 – 6.62 (m, 3H), 4.11 (t, *J* = 4.7 Hz, 2H), 3.01 (s, 3H), 2.33 (d, *J* = 19.3 Hz, 6H). ^13^C NMR (126 MHz, DMSO-*d*_6_): *δ* 161.9, 146.7, 143.6, 142.8, 138.1, 135.3, 131.5, 130.0, 128.7, 126.7, 126.0, 105.3, 42.1, 38.7, 20.7, 20.4. HRMS (ESI) *m/z*: [M+H]^+^ calculated for C_16_H_20_N_3_O_3_S: 334.1225, found 334.1214. HPLC purity (254 nm) >99%.

**(*E*)-5-(2,5-Dimethoxyphenyl)-*N*-(3-(methylsulfonyl)allyl)-1*H*-pyrazole-3-carboxamide (1o).** White solid (35% yield, m.p. 75 °C). ^1^H NMR (500 MHz, DMSO-*d*_6_): *δ* 8.62 (s, 1H), 7.36 (d, *J* = 3.1 Hz, 1H), 7.18 (s, 1H), 7.09 (d, *J* = 9.0 Hz, 1H), 6.94 (dd, *J* = 9.0, 3.0 Hz, 1H), 6.82 (dt, *J* = 15.2, 4.4 Hz, 1H), 6.70 (dt, *J* = 15.4, 1.8 Hz, 1H), 4.12 (ddd, *J* = 6.1, 4.4, 1.8 Hz, 2H), 3.86 (s, 3H), 3.78 (s, 3H), 3.02 (s, 3H). ^13^C NMR (126 MHz, DMSO-*d*_6_): *δ* 153.2, 150.0, 143.4, 130.0, 114.6, 113.2, 112.8, 105.7, 55.9, 55.5, 42.1, 38.7. HRMS (ESI) *m/z*: [M+H]^+^ calculated for C_16_H_20_N_3_O_5_S: 366.1124, found 366.1127; HPLC purity (254 nm) >99%.

**(*E*)-5-(2,6-Dichlorophenyl)-*N*-(3-(methylsulfonyl)allyl)-1*H*-pyrazole-3-carboxamide (1p).** White solid (25% yield, m.p. 213–214 °C). ^1^H NMR (500 MHz, DMSO-*d*_6_): *δ* 13.74 (d, *J* = 162.0 Hz, 1H), 8.68 (s, 1H), 7.63 (d, *J* = 8.0 Hz, 2H), 7.52 (s, 1H), 6.88 – 6.66 (m, 3H), 4.12 (s, 2H), 3.02 (s, 3H). ^13^C NMR (126 MHz, DMSO-*d*_6_): *δ* 161.6, 158.6, 146.6, 143.4, 142.8, 137.2, 135.1, 132.0, 130.9, 130.1, 128.5, 128.2, 107.0, 106.4, 42.1, 38.7. HRMS (ESI) *m/z*: [M+H]^+^ calculated for C_14_H_14_Cl_2_N_3_O_3_S: 374.0133, found 374.0124; HPLC (Method A): *t*_ret_ 1.22 min, purity (254 nm) >99%.

**(*E*)-5-(3,5-Dichlorophenyl)-*N*-(3-(methylsulfonyl)allyl)-1*H*-pyrazole-3-carboxamide (1q).** White solid (33% yield, m.p. >300 °C). ^1^H NMR (500 MHz, DMSO-*d*_6_): *δ* 13.90 (dd, *J* = 54.4, 2.0 Hz, 1H), 8.91 – 8.63 (m, 1H), 7.94 (t, *J* = 1.7 Hz, 1H), 7.77 (d, *J* = 1.9 Hz, 1H), 7.60 (dt, *J* = 21.1, 2.0 Hz, 1H), 7.37 (dd, *J* = 30.6, 2.1 Hz, 1H), 6.85 – 6.61 (m, 2H), 4.18 – 4.07 (m, 2H), 3.01 (d, *J* = 4.2 Hz, 3H). ^13^C NMR (126 MHz, DMSO-*d*_6_): *δ* 161.5, 158.6, 158.5, 148.0, 147.9, 147.7, 147.6, 143.4, 142.7, 140.9, 140.7, 138.6, 138.4, 136.6, 136.5, 134.9, 134.7, 132.0, 131.9, 130.3, 130.1, 127.7, 127.1, 123.8, 123.5, 104.5, 103.2, 42.1, 38.7(4), 38.7(1). HRMS (ESI) *m/z*: [M+H]^+^ calculated for C_14_H_14_Cl_2_N_3_O_3_S: 374.0133, found 374.0128; HPLC purity (254 nm) >99%.

### Synthesis of Intermediates VI-XI

To a stirred solution of ethyl 5-(2-ethoxyphenyl)-1*H*-pyrazole-3-carboxylate (**IIIa**) (5.0 g, 19.2 mmol, 1.0 eq.) in THF (50 mL) at 0 °C was added 3,4-dihydro-2*H*-pyran (0.4 g, 28.8 mmol, 1.5 eq.) and PTSA (4.9 g, 28.8 mmol, 1.5 eq.). The reaction was stirred at 25 °C for 12 h. On completion of the reaction (based on TLC and LCMS), the mixture was diluted with water (50 mL) and the aqueous layer was extracted with EtOAc (2 x 50 mL), washed with brine, dried with anhydrous Na_2_SO_4_ and concentrated. The resulting crude was purified by combiflash (eluted with 30 % EtOAc in hexane) to afford intermediate **VI** as a yellow liquid (1.0 g, 15% yield). MS (ESI) *m/z*: 345.3 [M+H]^+^.

To a stirred solution of ethyl 5-(2-ethoxyphenyl)-1-(tetrahydro-2*H*-pyran-2-yl)-1*H*-pyrazole-3-carboxylate (**VI**) (0.5 g, 1.45 mmol, 1.0 eq.) in THF (5 mL) and water (3 mL) was added lithium hydroxide monohydrate (0.178 g, 4.36 mmol, 3.0 eq.) at 0 °C. The reaction mixture was stirred at 25 °C for 6 h. On completion of the reaction (based on TLC and LCMS), the reaction mixture was concentrated under reduced pressure, residue was acidified with 1N HCl until pH∼3. The organic layer was extracted with EtOAc (50 mL x 3), combined organic layers dried over anhydrous Na_2_SO_4_ and concentrated. The residue was washed with hexane to afford 5-(2-ethoxyphenyl)-1-(tetrahydro-2*H*-pyran-2-yl)-1*H*-pyrazole-3-carboxylic acid (**VII**) as a white solid (0.4 g, 87% yield). MS (ESI) *m/z*: 315.2 [M+H]^+^. ^1^H NMR (400 MHz, DMSO-*d*_6_): *δ* 12.83 (br s, 1H), 7.49 (t, *J* = 8.0 Hz, 1H), 7.32 (d, *J* = 8.0 Hz, 1H), 7.18 (d, *J* = 8.0 Hz, 1H), 7.08 (t, *J* = 8.0 Hz, 1H), 6.68 (s, 1H) 5.12 (d, *J* = 12 Hz, 1H), 4.12 (dd, *J* = 8 Hz, 2H), 3.88 (d, *J* = 12 Hz, 1H), 2.33 (d, *J* = 12 Hz, 1H), 1.96 (br s, 1H), 1.85 (t, *J* = 4 Hz, 1H), 1.55 – 1.47 (m, 4H), 1.11 (t, *J* = 8 Hz, 3H).

To a stirred solution of ethyl 5-(2-ethoxyphenyl)-1*H*-pyrazole-3-carboxylate (**IIIa**) (1.0 g, 3.84 mmol, 1.0 eq.) in DCM (10 mL) at 0 °C was added triethylamine (1.61 mL, 11.52 mmol, 1.5 eq.). Reaction mixture stirred for 10 min and 2-(trimethylsilyl)ethoxymethyl chloride (1.92 g, 11.52 mmol, 3.0 eq.) was added dropwise. Reaction mixture was stirred at 25 °C for 3 h. On completion of the reaction (based on TLC and LCMS), the reaction mixture was diluted with water (150 mL), organic layer was extracted with EtOAc (3 x 300 mL) and concentrated. The resulting crude was purified by combiflash (eluted with 5% EtOAc in hexane) to afford ethyl 5-(2-ethoxyphenyl)-1-((2-(trimethylsilyl)ethoxy)methyl)-1*H*-pyrazole-3-carboxylate (**VIII**) (0.6 g, 40% yield) as colorless oil. MS (ESI) *m/z*: 391.3 [M+H]^+^.

To a stirred solution of ethyl 5-(2-ethoxyphenyl)-1-((2-(trimethylsilyl)ethoxy)methyl)-1*H*-pyrazole-3-carboxylate (**VIII**) (0.6 g, 1.53 mmol, 1.0 eq.) in MeOH (6 mL) was added lithium hydroxide (0.11 g, 4.60 mmol, 3.0 eq.) in water (1 mL). Reaction mixture was stirred at 50 °C for 3 h. On completion of the reaction (based on TLC and LCMS), the mixture was quenched with 1N HCl (40 mL), organic layer extracted with EtOAc (3 x 100 mL) and concentrated to afford 5-(2-ethoxyphenyl)-1-((2-(trimethylsilyl)ethoxy)methyl)-1*H*-pyrazole-3-carboxylic acid (**IX**) (0.5 g, 90% yield) as a white solid. MS (ESI) *m/z*: 363.2 [M+H]^+^.

To a stirred solution of ethyl 4-(2-ethoxyphenyl)-2,4-dioxobutanoate (**IIa**) (1.0 g, 3.7 mmol, 1.0 eq.) in EtOH (10 mL) at 0 °C was added NH_2_OH.HCl (0.395 g, 5.6 mmol, 1.5 eq.) and the reaction was stirred at 60 °C for 6 h. On completion of the reaction (based on TLC and LCMS), the mixture was diluted with water (30 mL), organic layer extracted with EtOAc (3 x 50 mL), washed with brine, dried over anhydrous Na_2_SO_4_, filtered, and concentrated. The resulting crude was purified by combiflash (eluted with 30% EtOAc in hexane) to afford ethyl 3-(2-ethoxyphenyl)isoxazole-5-carboxylate (**X**) as a white solid. (0.5 g, 51% yield). MS (ESI) *m/z*: 262.3 [M+H]^+^.

To a stirred solution of ethyl 3-(2-ethoxyphenyl)isoxazole-5-carboxylate (**X**) (0.5 g, 1.9 mmol, 1.0 eq.) in THF (5 mL) and water (3 mL) was added lithium hydroxide (0.025 g, 5.7 mmol, 3.0 eq.). The reaction was stirred at 25 °C for 6 h. On completion of the reaction (based on TLC and LCMS), the reaction mixture was concentrated, and the residue was acidified with 1N HCl until pH∼3. The organic layer was extracted with EtOAc, combined organic layers were dried over anhydrous Na_2_SO_4_ and concentrated. The resulting crude was washed with hexane to afford 3-(2-ethoxyphenyl)isoxazole-5-carboxylic acid (**XI**) as a white solid (0.04 g, 68% yield). MS (ESI) *m/z*: 234.2 [M+H]^+^.

**(*E*)-3-(2-Ethoxyphenyl)-*N*-(3-(methylsulfonyl)allyl)isoxazole-5-carboxamide (10).** Amide coupling of 3-(2-ethoxyphenyl)isoxazole-5-carboxylic acid (**XI**) and (*E*)-3-(methylsulfonyl)prop-2-en-1-amine (**V**) following General Procedure-I furnished (*E*)-3-(2-ethoxyphenyl)-*N*-(3-(methylsulfonyl)-allyl)isoxazole-5-carboxamide (**10**) as a white solid (0.035 g, 40% yield, m.p. 170°C). ^1^H NMR (400 MHz, DMSO-*d*_6_): *δ* 8.96 (s, 1H), 7.90 (d, *J* = 8.0 Hz, 1H), 7.53 (t, *J* = 16 Hz, 1H), 7.24 (d, *J* = 8.0 Hz, 1H), 7.14 (t, *J* = 8.0 Hz, 2H), 6.85 – 6.74 (m, 2H), 4.29 (q, *J* = 20 Hz, 2H), 4.16 (t, *J* = 8 Hz, 2H), 2.99 (s, 3H), 1.47 (d, *J* = 16 Hz, 3H). ^13^C NMR (101 MHz, DMSO-*d*_6_): *δ* 167.3, 159.6, 159.2, 155.8, 142.8, 132.8, 130.9, 121.3, 115.2, 113.5, 102.9, 64.6, 42.6, 14.9. HRMS (ESI) *m/z*: [M+H]^+^ calculated for C_16_H_19_N_2_O_5_S: 351.1015, found 351.1013. HPLC purity (254 nm) >99%.

**(*E*)-5-Cyclohexyl-*N*-(3-(methylsulfonyl)allyl)-1*H*-pyrazole-3-carboxamide (4c).** 5-cyclohexyl-1*H*-pyrazole-3-carboxylic acid (**XIII**) was synthesized from 1-cyclohexylethan-1-one (**XII**) (1.0 g, 7.9 mmol) following a sequence of Claisen condensation/pyrazole formation/ester hydrolysis as white solid (1.0 g, 65% yield over three steps). MS (ESI) *m/z*: 195 [M+H]^+^. **4c** was synthesized by amide coupling of 5-cyclohexyl-1*H*-pyrazole-3-carboxylic acid (**XIII**) and (*E*)-3-(methylsulfonyl)prop-2-en-1-amine (**V**) following General Procedure-I (white solid, 0.06 g, 37% yield). ^1^H NMR (400 MHz, DMSO-*d*_6_): *δ* 12.98 –12.91 (m, 1H), 8.43 (t, *J* = 6.0 Hz, 1H), 6.77 (dt, *J* = 15.1, 4.3 Hz, 1H), 6.62 (dt, *J* = 15.2, 1.8 Hz, 1H), 6.39 (d, *J* = 2.0 Hz, 1H), 4.07 – 4.02 (m, 2H), 2.99 (s, 3H), 2.69 – 2.60 (m, 1H), 1.94 – 1.89 (m, 2H), 1.78 – 1.72 (m, 2H), 1.71 – 1.64 (m, 1H), 1.40 – 1.32 (m, 4H), 1.23 (ddd, *J* = 12.0, 8.9, 3.3 Hz, 1H). ^13^C NMR (126 MHz, DMSO-*d*_6_): *δ* 162.1, 149.8, 146.1, 143.6, 129.9, 101.6, 42.1, 34.4, 32.2, 25.5, 25.4. HRMS (ESI) *m/z*: [M+H]^+^ calculated for C_14_H_22_N_3_O_3_S: 312.1382, found 312.1378. HPLC purity (254 nm) >99%.

**(*E*)-3-Benzhydryl-*N*-(3-(methylsulfonyl)allyl)-1*H*-pyrazole-5-carboxamide (4g). 5-**benzhydryl-1-(tetrahydro-2*H*-pyran-2-yl)-1*H*-pyrazole 3-carboxylic acid (**XV**) was synthesized from 1,1-diphenylpropan-2-one (**XIV**) (5.0 g, 23.8 mmol) following a sequence of Claisen condensation/ pyrazole formation/THP protection/ ester hydrolysis as white solid (0.7 g, 15% yield over four steps). MS (ESI) *m/z*: 361.3 [M+H]^+^. To a stirring solution of 5-benzhydryl-2*H*-pyrrole-3-carboxylic acid (**XV**) (0.7 g, 1.93 mmol, 1.0 eq.) in DMF (7 mL) were added TBTU (0.74 g, 2.32 mmol, 1.2 eq.), DIPEA (1.02 mL, 5.80 mmol, 3.0 eq..) and (*E*)-3-(methyl sulfonyl) prop-2-en-1-amine (**V**) (0.27 g, 1.93 mmol, 1.0 eq.). The reaction was stirred at 25 °C for 16 h. After completion, the reaction mixture was quenched with water and extracted with EtOAc. The organic layers were washed with brine, dried over anhydrous Na_2_SO_4_, filtered, and concentrated to afford crude (*E*)-3-benzhydryl-*N*-(3-(methylsulfonyl)allyl)-1-(tetrahydro-2*H*-pyran-2-yl)-1*H*-pyrazole-5-carbox-amide (0.65 g, 70% yield). MS (ESI) *m/z*: 431.4 [M+H]^+^. To a stirring solution of (*E*)-3-benzhydryl-*N*-(3-(methylsulfonyl) allyl)-1-(tetrahydro-2*H*-pyran-2-yl)-1*H*-pyrazole-5-carboxamide (0.2 g, 0.41 mmol, 1.0 eq.) in DCM (2 mL) was added 4M HCl in dioxane (1 mL). Reaction mixture was stirred at 25 °C for 16 h. After completion, the reaction mixture was concentrated, and the resulting crude was purified by preparative HPLC to afford (*E*)-3-benzhydryl-*N*-(3-(methyl sulfonyl) allyl)-*1H*-pyrazole-5-carboxamide (**4g**) as a white solid (0.06 mg, 38% yield). ^1^H NMR (400 MHz, DMSO-*d*_6_): *δ* 11.16 (s, 1H), 7.32 (t, *J* = 7.1 Hz, 4H), 7.21 (t, *J* = 11.8 Hz, 6H), 6.83 (d, *J* = 12.2 Hz, 1H), 6.53 (d, *J* = 15.8 Hz, 1H), 6.35 (s, 1H), 5.63 (d, *J* = 3.2 Hz, 1H), 4.13 (t, *J* = 3.2 Hz, 2H), 2.89 (s, 1H); ^13^C NMR (101 MHz, CD_3_CN): *δ* 162.7, 144.9, 143.2, 130.4, 129.7, 129.5, 127.9, 118.3, 106.3, 68.3, 49.4, 42.9, 39.7. HRMS (ESI) *m/z*: [M+H]^+^ calculated for C_21_H_22_N_3_O_3_S: 396.1382, found 396.1377. HPLC purity (254 nm) >99%.

### Synthesis of Intermediates XVIIa–XIXa and XVIIb–XIXb

To a stirred solution 2-(2-ethoxyphenyl)acetic acid (**XVIa**) (10.0 g, 55.49 mmol, 1.0 eq.) in DMF (2 mL) at 0 °C were added HATU (23.2 g, 61.04 mmol, 1.1 eq.), DIPEA (25.61 mL, 138..7 mmol, 2.5 eq.) and Weinreb amine (5.41 g, 55.5 mmol, 1.0 eq.). Reaction was stirred at 25 °C for 6 h. On completion of the reaction (based on TLC and LCMS), the reaction mixture was diluted with water (100 mL), organic layer extracted with EtOAc (2 x 200 mL), washed with brine washing and concentrated. The resulting crude was purified by combiflash (eluted with 30% EtOAc in hexane) to afford 2-(2-ethoxyphenyl)-*N*-methoxy-*N*-methylacetamide (8.0 g, 65% yield). MS (ESI) *m/z*: 224.3 [M+H]^+^. Step-2: To a stirred solution of 2-(2-ethoxyphenyl)-*N*-methoxy-*N*-methylacetamide (9.5 g, 42.54 mmol, 1.0 eq.) in THF (40 mL) at 0 °C, 3M CH_3_MgBr (28.36 mL, 85.0 mmol, 2.0 eq.) was added dropwise and stirred at 0 °C for 3 h. After completion, the reaction was quenched with MeOH (50 mL) and 1N HCl solution (100 mL) was added. The organic layer was extracted with EtOAc (2 x 300 mL) and concentrated. The resulting crude was purified by combiflash (eluted with 40% EtOAc in hexane) to afford 1-(2-ethoxyphenyl)propan-2-one (**XVIIa**) (6.0 g, 79% yield). MS (ESI) *m/z*: 179.2 [M+H]^+^.

To a stirred solution 1-(2-ethoxyphenyl)propan-2-one (**XVIIa**) (6.0 g, 33.66 mmol, 1.0 eq.) and diethyl oxalate (4.59 mL, 33.66 mmol, 1.0 eq.) in THF (2 mL) at –78 °C was added 1M LiHMDS (33.6 mL, 33.66 mmol, 1.1 eq.) and stirred at 25 °C for 6 h. After completion, the reaction mixture was diluted with NH_4_Cl solution (100 mL), organic layer extracted with EtOAc (2 x 200 mL), washed with brine, dried over anhydrous Na_2_SO_4_, filtered, and concentrated. The residue was purified by combiflash (eluted with 30% EtOAc in hexane) to afford ethyl 5-(2-ethoxyphenyl)-2,4-dioxopentanoate (6.0 g, 64% yield). MS (ESI) *m/z*: 279.2 [M+H]^+^. To a stirred solution ethyl 5-(2-ethoxyphenyl)-2,4-dioxopentanoate (2.0 g, 7.18 mmol, 1.0 eq.) in EtOH (20 mL) was added hydrazine hydrate (0.352 mL, 7.18 mmol, 1.0 eq.) and the reaction was stirred at 80 °C for 3 h. After completion, the reaction mixture was concentrated, and the crude was purified by combiflash (eluted with 30 % EtOAc in hexane) to afford ethyl 5-(2-ethoxybenzyl)-1*H*-pyrazole-3-carboxylate (**XVIIIa**) (0.8 g, 41% yield). MS (ESI) *m/z*: 275.2 [M+H]^+^.

THP protection of 5-(2-ethoxybenzyl)-1*H*-pyrazole-3-carboxylate (**XVIIIa**) (0.8 g, 2.91 mmol, 1.0 eq.) according to General Procedure-IV afforded ethyl 3-(2-ethoxybenzyl)-1-(tetrahydro-2*H*-pyran-2-yl)-1*H*-pyrazole-5-carboxylate (0.4 g, 39% yield). MS (ESI) *m/z*: 359.1 [M+H]^+^. Ester hydrolysis of ethyl 3-(2-ethoxybenzyl)-1-(tetrahydro-2*H*-pyran-2-yl)-1*H*-pyrazole-5-carboxylate (0.4 g, 1.11 mmol, 1.0 eq.) according to General Procedure-III afforded 5-(2-ethoxybenzyl)-1-(tetrahydro-2*H*-pyran-2-yl)-1*H*-pyrazole-3-carboxylic acid (**XIXa**) (0.4 g, quantitative yield). MS (ESI) *m/z*: 331.1 [M+H]^+^.

**(*E*)-3-(2-Ethoxybenzyl)-*N*-(3-(methylsulfonyl)allyl)-1*H*-pyrazole-5-carboxamide (4d).** Step-1: Amide coupling between 5-(2-ethoxybenzyl)-1-(tetrahydro-2*H*-pyran-2-yl)-1*H*-pyrazole-3-carboxylic acid (**XIXa**) and (*E*)-3-(methylsulfonyl)prop-2-en-1-amine (**V**) following General Procedure-II furnished (*E*)-5-(2-ethoxybenzyl)-*N*-(3-(methylsulfonyl)allyl)-1-(tetrahydro-2*H*-pyran-2-yl)-1*H*-pyrazole-3-carboxamide (0.3 g, 56% yield). Step-2: Following General Procedure-V for THP deprotection, desired compound (*E*)-5-(2-ethoxybenzyl)-*N*-(3-(methylsulfonyl)allyl)-1*H*-pyrazole-3-carboxamide (**4d**) was obtained as a white solid (0.02 g, 10% yield, m.p. 160–162 °C). ^1^H NMR (400 MHz, DMSO-*d*_6_): *δ* 12.82 (s, 1H), 8.15 (s, 1H), 7.20 – 7.16 (m, 1H), 7.10 (d, *J* = 7.2 Hz, 1H), 6.96 (d, *J* = 8.0 Hz, 1H), 6.86 (t, *J* = 7.2 Hz, 1H), 6.79 – 6.63 (m, 2H), 6.29 (s, 1H), 4.07 – 4.02 (m, 4H), 3.92 – 3.89 (m, 2H), 2.94 (s, 3H), 1.33 (t, *J* = 7.2 Hz, 3H). ^13^C NMR (101 MHz, DMSO-*d*_6_): *δ* 162.5, 156.5, 146.9, 144.1, 143.6, 130.4, 130.0, 128.5, 127.1, 120.7, 112.2, 104.7, 63.7, 42.6, 39.1, 26.0, 15.2. HRMS (ESI) *m/z*: [M+H]^+^ calculated for C_17_H_22_N_3_O_4_S: 364.1331, found 364.1325. HPLC purity (254 nm) >98%. Intermediates **XVIIb**–**XIXb** were synthesized following the same sequence of reactions used for the synthesis of **XVIIa**–**XIXa**.

**(*E*)-5-(1-(2-Ethoxyphenyl)ethyl)-*N*-(3-(methylsulfonyl)allyl)-1*H*-pyrazole-3-carboxamide (4e)**. The compound was synthesized following a two-step reaction sequence of a) amide coupling with acid **XIXb** and amine **V** following General Procedure-II, and b) THP deprotection (white solid, 0.05 g, 19% yield over two steps, m.p. 128–130 °C). ^1^H NMR (400 MHz, DMSO-*d*_6_): *δ* 13.0 (s, 1H), 8.46 (s, 1H), 7.19 – 7.17 (m, 1H), 7.05 (d, *J* = 7.6 Hz, 1H), 6.98 (d, *J* = 8.0 Hz, 1H), 6.88 (t, *J* = 7.6 Hz, 1H), 6.79 (s, 1H), 6.77 – 6.75 (m, 1H), 6.65 (s, 1H), 4.57 – 4.54 (m, 1H), 4.04 – 4.02 (m, 4H), 3.00 (s, 3H), 1.53 (d, *J* = 6.8 Hz, 3H), 1.34 (t, *J* = 7.2 Hz, 3H); ^13^C NMR (101 MHz, DMSO-*d*_6_): *δ* 162.5, 155.9, 148.8, 146.6, 144.1, 132.7, 130.5, 128.1, 127.6, 120.9, 112.4, 103.9, 63.9, 42.6, 29.9, 20.5, 15.2. HRMS (ESI) *m/z*: [M+H]^+^ calculated for C_18_H_24_N_3_O_4_S: 378.1488, found 378.1483. HPLC purity (254 nm) >99%.

**(*E*)-*N*-(3-(Methylsulfonyl)allyl)-5-(pyridin-2-yl)-1*H*-pyrazole-3-carboxamide (4h).** The compound was synthesized following General Procedure-I using 5-(pyridin-2-yl)-1*H*-pyrazole-3-carboxylic acid (**XX**) and (*E*)-3-(methylsulfonyl)prop-2-en-1-amine (**V**) (white solid, 0.07 g, 16% yield). ^1^H NMR (500 MHz, DMSO-*d*_6_): *δ* 8.80 (t, *J* = 5.5 Hz, 1H), 8.65 (dt, *J* = 4.8, 1.4 Hz, 1H), 8.03 – 7.96 (m, 2H), 7.48 – 7.38 (m, 2H), 6.82 (dt, *J* = 15.3, 4.3 Hz, 1H), 6.72 (dt, *J* = 15.3, 1.7 Hz, 1H), 4.13 (td, *J* = 4.3, 1.8 Hz, 2H), 3.01 (s, 3H), 2.54 (s, 1H). ^13^C NMR (214 MHz, DMSO-*d*_6_): *δ* 158.5, 158.3, 158.1, 157.9, 148.6, 143.2, 138.5, 130.2, 123.5, 120.4, 117.3, 115.9, 114.6, 113.3, 104.2, 42.1, 40.4. HRMS (ESI) *m/z*: [M+H]^+^ calculated for C_13_H_15_N_4_O_3_S: 307.0865, found 307.0853. HPLC purity (254 nm) >99%.

**(*E*)-5-(1-Methyl-1*H*-pyrrol-2-yl)-*N*-(3-(methylsulfonyl)allyl)-1*H*-pyrazole-3-carboxamide (4i).** The compound was synthesized following General Procedure-I using 5-(1-methyl-1*H*-pyrrol-2-yl)-1*H*-pyrazole-3-carboxylic acid (**XXI**) and (*E*)-3-(methylsulfonyl)prop-2-en-1-amine (**V**) (white solid, 0.04 g, 10% yield). ^1^H NMR (500 MHz, DMSO-*d*_6_): *δ* 13.36 (s, 1H), 8.64 (s, 1H), 6.87 (s, 2H), 6.81 (dt, *J* = 15.3, 4.4 Hz, 1H), 6.70 (d, *J* = 15.4 Hz, 1H), 6.41 (s, 1H), 6.07 (t, *J* = 3.2 Hz, 1H), 4.14 – 4.08 (m, 2H), 3.74 (s, 3H), 3.01 (s, 3H). ^13^C NMR (214 MHz, DMSO-*d*_6_): *δ* 161.9, 146.9, 143.4, 130.1, 124.9, 108.9, 107.4, 102.8, 42.2, 38.7, 35.3. HRMS (ESI) *m/z*: [M+H]^+^ calculated for C_13_H_17_N_4_O_3_S: 309.1021, found 309.1011. HPLC purity (254 nm) >99%.

**(*E*)-*N*-(3-(Methylsulfonyl)allyl)-1*H*-indazole-3-carboxamide (5).** The compound was synthesized following General Procedure-I using 1*H*-indazole-3-carboxylic acid (**XXII**) and (*E*)-3-(methylsulfonyl)prop-2-en-1-amine (**V**) (white solid, 0.06 g, 13% yield, m.p. 191–192 °C). ^1^H NMR (500 MHz, DMSO-*d*_6_): *δ* 13.64 (s, 1H), 8.80 (t, *J* = 5.9 Hz, 1H), 8.17 (dt, *J* = 8.1, 1.0 Hz, 1H), 7.62 (dt, *J* = 8.4, 0.9 Hz, 1H), 7.42 (ddd, *J* = 8.3, 6.8, 1.1 Hz, 1H), 7.25 (ddd, *J* = 8.0, 6.9, 0.9 Hz, 1H), 6.84 (dt, *J* = 15.3, 4.4 Hz, 1H), 6.72 (dt, *J* = 15.2, 1.7 Hz, 1H), 4.16 (ddd, *J* = 6.1, 4.4, 1.8 Hz, 2H), 3.01 (s, 3H). ^13^C NMR (126 MHz, DMSO-*d*_6_): *δ* 162.4, 143.5, 141.1, 137.9, 130.1, 126.6, 122.1, 121.5(2), 121.5(0), 110.7, 42.1, 38.6. HRMS (ESI) *m/z*: [M+H]^+^ calculated for C_12_H_14_N_3_O_3_S: 280.0756, found 280.0745. HPLC purity (254 nm) >99%.

**(*E*)-*N*-(3-(Methylsulfonyl)allyl)-4,5-dihydro-1*H*-benzo[*g*]indazole-3-carboxamide (6).** The compound was synthesized following General Procedure-I using 4,5-dihydro-1*H*-benzo[*g*]indazole-3-carboxylic acid (**XXIII**) and (*E*)-3-(methylsulfonyl)prop-2-en-1-amine (V) (white solid, 0.02 g, 4% yield). ^1^H NMR (500 MHz, DMSO-*d*_6_): *δ* 8.47 (t, *J* = 6.0 Hz, 1H), 7.66 (dd, *J* = 7.6, 1.4 Hz, 1H), 7.33 – 7.27 (m, 2H), 7.24 (td, *J* = 7.4, 1.4 Hz, 1H), 6.80 (dt, *J* = 15.3, 4.4 Hz, 1H), 6.70 (dt, *J* = 15.2, 1.8 Hz, 1H), 4.09 (ddd, *J* = 6.1, 4.4, 1.8 Hz, 2H), 3.01 (s, 3H), 2.95 (ddd, *J* = 7.3, 5.8, 2.1 Hz, 2H), 2.92 – 2.88 (m, 2H). ^13^C NMR (214 MHz, DMSO-*d*_6_): *δ* 162.6, 143.6, 142.0, 139.6, 135.8, 130.0, 128.6, 127.9, 126.8, 125.9, 121.4, 116.8, 42.2, 38.5, 28.8, 19.1. HRMS (ESI) *m/z*: [M+H]^+^ calculated for C_16_H_18_N_3_O_3_S: 332.1069; found 332.1064. HPLC purity (254 nm) >99%.

**(*E*)-5-(2-Ethoxyphenyl)-1-methyl-*N*-(3-(methylsulfonyl)allyl)-1*H*-pyrazole-3-carboxamide (20).** The compound was synthesized following General Procedure-I using 4,5-dihydro-1*H*-benzo[*g*]indazole-3-carboxylic acid (**XXIV**) and (*E*)-3-(methylsulfonyl)prop-2-en-1-amine (**V**) (white sticky solid, 0.13 g, 81% yield). ^1^H NMR (400 MHz, DMSO-*d*_6_): *δ* 8.57 (br s, 1H), 7.48 (t, *J* = 8.0 Hz, 1H), 7.30 (d, *J* = 8.0 Hz, 1H), 7.18 (d, *J* = 8.0 Hz, 1H), 7.07 (t, *J* = 8 Hz, 1H), 6.81 – 6.76 (m, 1H), 6.69 – 6.63 (m, 2H), 4.12 – 4.07 (m, 4H), 3.70 (s, 3H), 3.01 (s, 3H), 1.28 (t, *J* = 4 Hz, 3H). ^13^C NMR (101 MHz, DMSO-*d*_6_): *δ* 161.9, 156.3, 145.3, 143.9, 142.3, 131.9, 131.6, 130.5, 121.0, 118.7, 112.9, 107.6, 63.9, 42.6, 38.1, 15.0. HRMS (ESI) *m/z*: [M+H]^+^ calculated for C_17_H_22_N_3_O_4_S: 364.1331, found 364.1329. HPLC purity (254 nm) >99%.

**5-(2-Ethoxyphenyl)-*N*-(3-(methylsulfonyl)propyl)-1*H*-pyrazole-3-carboxamide (2).** To a stirred solution of 5-(2-ethoxyphenyl)-1*H*-pyrazole-3-carboxylic acid (**IVa**) and 3-(methylsulfonyl)propan-1-amine (**XXVa**) in DMF (3 mL) were added HBTU (221 mg, 0.583 mmol, 0.8 eq.), HOBt (123 mg, 0.8 mmol, 1.1 eq.), and DIPEA (0.4 mL, 2.19 mmol, 3.0 eq.). The reaction was stirred at 25 °C for 16 h. After completion, the reaction was quenched with water, extracted with EtOAc, washed with brine, dried over anhydrous Na_2_SO_4_, filtered, and concentrated. The residue was purified by preparative HPLC to afford 5-(2-ethoxyphenyl)-*N*-(3-(methylsulfonyl)propyl)-1*H*-pyrazole-3-carboxamide (**2**) as a white sticky solid (0.08 g, 31% yield). ^1^H NMR (500 MHz, DMSO-*d*_6_): *δ* 8.34 (t, *J* = 6.2 Hz, 1H), 7.73 (dd, *J* = 7.6, 1.7 Hz, 1H), 7.34 (ddd, *J* = 8.7, 7.3, 1.7 Hz, 1H), 7.13 (dd, *J* = 8.4, 1.1 Hz, 1H), 7.09 – 6.99 (m, 2H), 4.16 (q, *J* = 6.9 Hz, 2H), 3.37 (q, *J* = 6.6 Hz, 2H), 3.18 – 3.12 (m, 2H), 2.98 (s, 3H), 1.99 – 1.91 (m, 2H), 1.41 (t, *J* = 6.9 Hz, 3H). ^13^C NMR (126 MHz, DMSO-*d*_6_): *δ* 155.0, 129.6, 127.8, 120.5, 112.8, 105.4, 63.7, 51.5, 37.1, 22.5, 14.5. HRMS (ESI) *m/z*: [M+H]^+^ calculated for C_16_H_22_N_3_O_4_S: 352.1331; found 352.1331. HPLC purity (254 nm) >99%.

**5-(2-Ethoxyphenyl)-*N*-methyl-1*H*-pyrazole-3-carboxamide (3)**. The compound was synthesized following General Procedure-I using 5-(2-ethoxyphenyl)-1*H*-pyrazole-3-carboxylic acid (**IVa**) and methylamine **XXVb** (yellow solid, 0.03 g, 8% yield, m.p. 121–123 °C). ^1^H NMR (400 MHz, DMSO-*d*_6_): *δ* 8.15 (d, J = 6.2 Hz, 1H), 7.74 (dd, J = 7.7, 1.7 Hz, 1H), 7.33 (ddd, J = 8.9, 7.4, 1.7 Hz, 1H), 7.13 (dd, J = 8.5, 1.1 Hz, 1H), 7.06 (s, 1H), 7.01 (td, J = 7.5, 1.1 Hz, 1H), 4.16 (q, J = 7.0 Hz, 2H), 2.77 (d, J = 4.7 Hz, 3H), 1.41 (t, J = 6.9 Hz, 3H). ^13^C NMR (126 MHz, DMSO-*d*_6_): *δ* 162.4, 154.9, 147.2, 139.9, 129.7, 120.6, 117.5, 112.8, 105.1, 63.7, 25.6, 14.5. HRMS (ESI) *m/z*: [M+H]^+^ calculated for C_13_H_16_N_3_O_2_: 246.1243; found 246.1237. HPLC purity (254 nm) >99%.

***N*-(1,1-Dioxido-2,3-dihydrothiophen-3-yl)-5-(2-ethoxyphenyl)-1*H*-pyrazole-3-carboxamide (24f).** The compound was synthesized following General Procedure-II using intermediate **IVa** and 3-amino-2,3-dihydrothiophene 1,1-dioxide (**XXVc**) (white solid, 0.03 g, 20% yield, m.p. 106 °C). ^1^H NMR (400 MHz, DMSO-*d*_6_): *δ* 8.59 (s, 1H), 7.72 (d, *J* = 4.8 Hz, 1H), 7.37 – 7.29 (m, 1H), 7.14 – 7.09 (m, 3H), 7.05 (t, *J* = 7.6 Hz, 1H), 6.91 – 6.89 (m, 1H), 5.34 (s, 1H), 4.21 (q, *J* = 7.2 Hz, 2H), 3.75 (d, *J* = 8.0 Hz, 1H), 3.36 (d, *J* = 5.6 Hz, 1H), 1.42 (t, *J* = 6.8 Hz, 3H). ^13^C NMR (101 MHz, DMSO-*d*_6_): *δ* 155.5, 142.4, 133.3, 130.2, 128.2, 121.0, 113.3, 106.2, 64.1, 53.9, 48.3, 15.0. HRMS (ESI) *m/z*: [M+H]^+^ calculated for C_16_H_18_N_3_O_4_S: 348.1018, found 348.1023. HPLC purity (254 nm) >99%.

**Methyl (*E*)-4-(5-(2-ethoxyphenyl)-1*H*-pyrazole-3-carboxamido)but-2-enoate (28).** The compound was synthesized following General Procedure-I using intermediate **IVa** and methyl (*E*)-4-aminobut-2-enoate (**XXVd**) (white solid, 0.06 g, 54% yield, m.p. 180 °C). ^1^H NMR (400 MHz, DMSO-*d*_6_): *δ* 13.09 (s, 1H), 8.23 (s, 1H), 7.68 (s, 1H), 7.34 (s, 1H), 7.14 (d, *J* = 8.0 Hz, 1H), 7.04 (d, *J* = 4.0 Hz, 1H), 6.96 (tt, *J* = 12 Hz, 1H), 5.95 (d, *J* = 16 Hz, 2H), 4.21 (q, *J* = 20 Hz, 2H), 4.07 (t, *J* = 4.0 Hz, 2H), 3.67 (s, 3H), 1.42 (t, *J* = 12 Hz, 3H). ^13^C NMR (101 MHz, DMSO-*d*_6_): *δ* 166.4, 162.4, 155.4, 147.1, 140.5, 130.6, 128.2, 121.0, 120.4, 117.9, 113.2, 105.8, 64.1, 51.8, 14.9. HRMS (ESI) m/z: [M+H]^+^ calculated for C_17_H_20_N_3_O_4_: 330.1454, found 330.1459. HPLC purity (254 nm) >99%.

**(*E*)-*N*-(4-(Dimethylamino)-4-oxobut-2-en-1-yl)-5-(2-ethoxyphenyl)-1*H*-pyrazole-3-carboxamide (29).** The compound was synthesized following General Procedure-I using intermediate **IVa** and (*E*)-4-amino-*N,N*-dimethylbut-2-enamide (**XXVe**) (yellow solid, 0.04 g, 5% yield, m.p. 70 °C). ^1^H NMR (500 MHz, DMSO-*d*_6_): *δ* 13.31 (s, 1H), 8.45 (t, *J* = 6.0 Hz, 1H), 7.70 (dd, *J* = 7.7, 1.7 Hz, 1H), 7.40 – 7.32 (m, 1H), 7.14 (d, *J* = 8.3 Hz, 1H), 7.07 – 7.00 (m, 2H), 6.63 (dt, *J* = 15.1, 5.3 Hz, 1H), 6.48 (dt, *J* = 15.2, 1.7 Hz, 1H), 4.17 (q, *J* = 6.9 Hz, 2H), 4.05 – 4.00 (m, 2H), 3.01 (s, 3H), 2.86 (s, 3H), 1.40 (t, *J* = 6.9 Hz, 3H). ^13^C NMR (126 MHz, MeOD): *δ* 159.1, 147.4, 133.1, 121.6, 119.7, 112.5, 104.3, 96.3, 55.9, 31.7, 28.4, 26.5, 5.6. HRMS (ESI) *m/z*: [M+H]^+^ calculated for C_18_H_23_N_4_O_3_: 343.1770, found 343.1768. HPLC purity (254 nm) >99%.

***N*-(Cyanomethyl)-5-(2-ethoxyphenyl)-1*H*-pyrazole-3-carboxamide (30).** The compound was synthesized following General Procedure-II using intermediate **IVa** and 2-aminoacetonitrile (**XXVf**) (white solid, 0.04 g, 17% yield, m.p. 168 °C). ^1^H NMR (400 MHz, DMSO-*d*_6_): *δ* 13.50 (s, 1H), 8.62 (s, 1H), 7.68 (d, *J* = 6.0 Hz, 1H), 7.37 (t, *J* = 7.2 Hz, 1H), 7.23 – 7.13 (m, 1H), 7.07 – 6.95 (m, 2H), 4.20 (d, *J* = 5.2 Hz, 2H), 4.21 – 4.16 (m, 2H), 1.40 (t, *J* = 6.8 Hz, 3H). ^13^C NMR (101 MHz, DMSO-*d*_6_): *δ* 162.6, 155.2, 145.9, 140.5, 130.4, 128.1, 120.9, 117.9, 117.4, 113.1, 105.8, 64.0, 27.4, 14.8. HRMS (ESI) *m/z*: [M+H]^+^ calculated for C_14_H_15_N_4_O_2_: 271.1195, found 271.1198. HPLC purity (254 nm) >99%.

***N*-(1-Cyanopyrrolidin-3-yl)-5-(2-ethoxyphenyl)-1*H*-pyrazole-3-carboxamide (31).** The compound was synthesized using intermediate **IVa** (125 mg, 0.54 mmol, 1.0 eq.), 3-aminopyrrolidine-1-carbonitrile (**XXVg**) (72 mg, 0.65 mmol, 1.2 eq.), T3P (50% by wt., 0.5 mL, 0.8 mmol, 1.5 eq.), and pyridine (0.13 mL, 1.6 mmol, 3.0 eq.) in DCM (3 mL) (yellow solid, 0.07 g, 40% yield, m.p. 143–144 °C). ^1^H NMR (500 MHz, DMSO-*d*_6_): *δ* 13.29 (s, 1H), 8.38 (d, *J* = 6.9 Hz, 1H), 7.69 (d, *J* = 7.6 Hz, 1H), 7.36 (t, *J* = 7.9 Hz, 1H), 7.14 (d, *J* = 8.3 Hz, 1H), 7.03 (d, *J* = 6.3 Hz, 2H), 4.48 (qd, *J* = 6.7, 5.4 Hz, 1H), 4.16 (q, *J* = 6.9 Hz, 2H), 3.68 – 3.51 (m, 2H), 3.43 (q, *J* = 7.8 Hz, 1H), 3.37 – 3.33 (m, 1H), 2.10 (dq, *J* = 13.4, 6.9 Hz, 1H), 2.03 – 1.95 (m, 1H), 1.40 (t, *J* = 6.9 Hz, 3H). ^13^C NMR (126 MHz, DMSO-*d*_6_): *δ* 162.1, 154.9, 146.7, 140.0, 129.9, 127.8, 120.6, 117.5, 117.4, 112.8, 105.5, 63.7, 54.2, 48.7, 48.7, 30.7, 14.5. HRMS (ESI) *m/z*: [M+H]^+^ calculated for C_17_H_20_N_5_O_2_: 326.1617, found 326.1622. HPLC purity (254 nm) >99%.

**(*E*)-*N*-(3-(Methylsulfonyl)allyl)-5-(phenylsulfonamido)-1*H*-pyrazole-3-carboxamide (7a).** Ethyl 5-amino-1*H*-pyrazole-3-carboxylate (**XXVI**) (1.0 g, 6.4 mmol, 1.0 eq.), DMAP (0.079 g, 0.64 mmol, 0.1 eq.) were dissolved in pyridine (10 mL) and benzenesulfonyl chloride (0.99 mL, 7.7 mmol, 1.2 eq.) was added at room temperature. After stirring at 85 °C for 8 h, 1 mol/L HCl was added, and the mixture was extracted with EtOAc. The extract was washed with 1 mol/L HCl, water and saturated brine, organic layers dried over anhydrous Na_2_SO_4_, filtered, and concentrated. The residue was purified by combiflash to afford ethyl 5-(phenylsulfonamido)-1*H*-pyrazole-3-carboxylate as a brown solid (1.3 g, 68% yield). Ethyl 5-(phenylsulfonamido)-1*H*-pyrazole-3-carboxylate (100 mg, 0.34 mmol, 1.0 eq.) was subjected to General Procedure-III for ester hydrolysis to afford 5-(phenylsulfonamido)-1*H*-pyrazole-3-carboxylic acid (**XXVIIa**) as a white solid (0.08 g, 88% yield). **7a** was synthesized following General Procedure-I using intermediate **XXVIIa** and (*E*)-3-(methylsulfonyl)prop-2-en-1-amine (**V**) to give **7a** as a white solid (0.125 g, 36% yield, m.p. 181 °C). ^1^H NMR (500 MHz, DMSO-*d*_6_): *δ* 13.17 (s, 1H), 10.64 (s, 1H), 8.86 (s, 1H), 7.80 – 7.77 (m, 2H), 7.63 (t, *J* = 7.3 Hz, 1H), 7.57 (t, *J* = 7.5 Hz, 2H), 6.82 – 6.69 (m, 3H), 4.07 (t, *J* = 4.7 Hz, 2H), 3.00 (s, 3H). ^13^C NMR (126 MHz, DMSO-*d*_6_): *δ* 158.4, 145.9, 142.7, 140.1, 136.9, 132.8, 130.3, 129.1, 126.6, 97.3, 42.1, 38.7. HRMS (ESI) *m/z*: [M+H]^+^ calculated for C_14_H_17_N_4_O_5_S_2_: 385.0640, found 385.0577. HPLC purity (254 nm) >99%.

**(*E*)-*N*-(3-(Methylsulfonyl)allyl)-5-(3-phenylureido)-1*H*-pyrazole-3-carboxamide (7d).** To a stirred solution of ethyl 5-amino-1*H*-pyrazole-3-carboxylate (**XXVI**) (1.5 g, 9.6 mmol, 1.0 eq.) in DMF was added phenyl isocyanate (1.1 g, 9.6 mmol, 1.0 eq.) and the reaction mixture was stirred at 70 °C for 12 h. After completion, the reaction mixture was concentrated to afford ethyl 5-(3-phenylureido)-1*H*-pyrazole-3-carboxylate (0.8 g, 30% yield). MS (ESI) *m/z*: 275.2 [M+H]^+^. Ester hydrolysis of ethyl 5-(3-phenylureido)-1*H*-pyrazole-3-carboxylate (0.9 g, 3.28 mmol, 1.0 eq.) following General Procedure-III afforded 5-(3-phenylureido)-1*H*-pyrazole-3-carboxylic acid (**XXVIIb**) as a white solid (0.5 g, 62% yield). MS (ESI) *m/z*: 247.1 [M+H]^+^. **7d** was synthesized following General Procedure-II using intermediate **XXVIIb** and (*E*)-3-(methylsulfonyl)prop-2-en-1-amine (**V**) (white solid, 0.01 g, 8% yield, m.p. 166–168 °C). ^1^H NMR (400 MHz, DMSO-*d*_6_): *δ* 13.03 (s, 1H), 9.02 (s, 1H), 8.90 – 8.84 (m, 2H), 7.45 (d, *J* = 7.6 Hz, 2H), 7.28 (t, *J* = 8.0 Hz, 2H), 7.05 – 6.95 (m, 2H), 6.82 – 6.71 (m, 2H), 4.09 (s, 2H), 3.06 (s, 3H). ^13^C NMR (101 MHz, DMSO-*d*_6_): *δ* 157.7, 151.9, 151.8, 147.9, 139.6, 139.3, 134.5, 130.2, 128.8, 122.1, 121.9, 118.3, 118.1, 96.6, 53.8, 50.2, 42.9, 42.1, 41.5. HRMS (ESI) *m/z*: [M+H]^+^ calculated for C_15_H_18_N_5_O_4_S: 364.1080, found 364.1075. HPLC purity (254 nm) >99%.

### Synthesis of Intermediates XXVIIIa-XXXa

To a stirred solution of ethyl 5-amino-1*H*-pyrazole-3-carboxylate (**XXVI**) (2.0 g, 12.9 mmol, 1.0 eq.) and benzoyl chloride (2.1 g, 15.0 mmol, 1.2 eq.) in DCM (20 mL) was added triethylamine (5.4 mL, 40.1 mmol, 3.0 eq.) and stirred at 25 °C for 12 h. After completion, the reaction mixture was diluted with water (50 mL), extracted with DCM (2 x 50 mL), washed with brine, dried over anhydrous Na_2_SO_4_, filtered, and concentrated. The crude was purified by combiflash (eluted with 40% EtOAc in hexane) to afford intermediate **XXVIIIa** as a yellow liquid (1.8 g, 55% yield).

Ethyl 5-benzamido-1-(tetrahydro-2*H*-pyran-2-yl)-1*H*-pyrazole-3-carboxylate (**XXIXa**) was synthesized from intermediate **XXVIIIa** following General Procedure-IV (0.72 g, 42% yield).

5-benzamido-1-(tetrahydro-2*H*-pyran-2-yl)-1*H*-pyrazole-3-carboxylic acid (**XXXa**) was synthesized from intermediate following general procedure of ester hydrolysis (0.4 g, 57% yield).

**(*E*)-5-Benzamido-*N*-(3-(methylsulfonyl)allyl)-1*H*-pyrazole-3-carboxamide (7b).** The compound was synthesized following a two-step reaction sequence: a) amide coupling using intermediate **XXXa** and (*E*)-3-(methylsulfonyl)prop-2-en-1-amine (**V**) by General Procedure-I, b) THP deprotection following General Procedure-V (white solid, 0.02 g, 13% yield over two steps, m.p. 208–210 °C). ^1^H NMR (400 MHz, DMSO-*d*_6_): *δ* 12.98 (s, 1H), 10.65 (s, 1H), 8.69 (s, 1H), 8.01 (d, *J* = 8.0 Hz, 2H), 7.58 – 7.51 (t, *J*_1_ = 8.0 Hz, *J*_2_ = 8.0 Hz, 3H), 7.30 (s, 1H), 6.85 – 6.71 (m, 2H), 4.12 (s, 2H), 3.13 – 2.99 (m, 3H). ^13^C NMR (101 MHz, DMSO-*d*_6_): *δ* 165.1, 159.2, 148.1, 143.3, 136.7, 134.4, 132.1, 130.7, 128.8, 128.2, 97.8, 42.6. HRMS (ESI) *m/z*: [M+H]^+^ calculated for C_15_H_17_N_4_O_4_S: 349.0971, found 349.0967. HPLC purity >99%.

### Synthesis of Intermediates XXVIIIb-XXXb

Ethyl 5-(2-ethoxybenzamido)-1*H*-pyrazole-3-carboxylate (**XXVIIIb**) was synthesized following General Procedure-I using 2-ethoxybenzoic acid and ethyl 3-amino-1*H*-pyrazole-5-carboxylate (**XXVI**) (1.4 g, 77% yield).

Ethyl 5-(2-ethoxybenzamido)-1-(tetrahydro-2*H*-pyran-2-yl)-1*H*-pyrazole-3-carboxylate (**XXIXb**) was synthesized from intermediate **XXVIIIb** following General Procedure-IV (0.7 g, 42% yield).

5-(2-ethoxybenzamido)-1-(tetrahydro-2*H*-pyran-2-yl)-1*H*-pyrazole-3-carboxylic acid (**XXXb**) was synthesized from intermediate following General Procedure-III (0.38 g, 57% yield).

**(*E*)-5-(2-Ethoxybenzamido)-*N*-(3-(methylsulfonyl)allyl)-1*H*-pyrazole-3-carboxamide (7c).** The compound was synthesized following a two-step reaction sequence: a) amide coupling using intermediate **XXXb** and (*E*)-3-(methylsulfonyl)prop-2-en-1-amine (**V**) by General Procedure-II, b) THP deprotection following General Procedure-V (white solid, 0.02 g, 16% yield over two steps, m.p. 201–203 °C). ^1^H NMR (400 MHz, DMSO-*d*_6_): *δ* 13.26 (s, 1H), 10.55 (s, 1H), 8.97 (s, 1H), 7.90 (d, *J* = 6.4 Hz, 1H), 7.56 – 7.52 (m, 1H), 7.39 (s, 1H), 7.21 (d, *J* = 8.4 Hz, 1H), 7.10 (t, *J* = 7.6 Hz, 1H), 6.84 – 6.78 (m, 2H), 4.28 (q, *J* = 6.8 Hz, 2H), 4.12 (s, 2H), 3.02 (s, 3H), 1.45 (t, *J* = 6.4 Hz, 3H). ^13^C NMR (101 MHz, DMSO-*d*_6_): *δ* 162.4, 158.7, 156.4, 147.1, 142.9, 136.5, 133.1, 130.7, 130.2, 122.0, 120.8, 113.3, 96.6, 64.7, 42.1, 38.8, 14.6. HRMS (ESI) *m/z*: [M+H]^+^ calculated for C_17_H_21_N_4_O_5_S: 393.1233, found 393.1233. HPLC purity (254 nm) 97%.

**(*E*)-5-(*N*-Methylbenzamido)-*N*-(3-(methylsulfonyl)allyl)-1*H*-pyrazole-3-carboxamide (7e).** *N*-Methylation of intermediate **XXIXa** (1.0 g, 2.91 mmol, 1.0 eq.) using methyl iodide (0.3 mL, 5.83 mmol, 2.0 eq.) and NaH (0.35 g, 8.74 mmol, 3.0 eq.) in DMF afforded ethyl 5-(*N*-methylbenzamido)-1-(tetrahydro-2*H*-pyran-2-yl)-1*H*-pyrazole-3-carboxylate (0.65 g, 63% yield). Ester hydrolysis of 5-(*N*-methylbenzamido)-1-(tetrahydro-2*H*-pyran-2-yl)-1*H*-pyrazole-3-carboxylate following General Procedure-III yielded 5-(*N*-methylbenzamido)-1-(tetrahydro-2*H*-pyran-2-yl)-1*H*-pyrazole-3-carboxylic acid (**XXXIa**) as a white solid (0.4 g, 59% yield). MS (ESI) *m/z*: 329.3 [M+H]^+^. **7e** was synthesized following a two-step reaction sequence: a) amide coupling using intermediate **XXXIa** and (*E*)-3-(methylsulfonyl)prop-2-en-1-amine (**V**) by General Procedure-I, b) THP deprotection following General Procedure-V (white solid, 0.04 g, 9% yield over two steps, m.p. 210 °C). ^1^H NMR (400 MHz, DMSO-*d*_6_): *δ* 13.41 (s, 1H), 8.82 (s, 1H), 7.36 (m, 5H), 6.80 – 6.68 (m, 3H), 4.07 (s, 2H), 3.33 (s, 3H), 3.01 (s, 3H). ^13^C NMR (214 MHz, DMSO-*d*_6_): *δ* 169.4, 158.4, 151.4, 142.6, 136.9, 136.1, 130.3, 129.8, 128.0, 127.6, 100.3, 42.1, 38.7, 36.9. HRMS (ESI) *m/z*: [M+H]^+^ calculated for C_16_H_19_N_4_O_4_S: 363.1127, found 363.1126. HPLC purity (254 nm) >98%.

**(*E*)-5-(2-Ethoxy-*N*-methylbenzamido)-*N*-(3-(methylsulfonyl)allyl)-1*H*-pyrazole-3-carboxamide (7f).** 5-(2-ethoxy-*N*-methylbenzamido)-1-(tetrahydro-2*H*-pyran-2-yl)-1*H*-pyrazole-3-carboxylic acid (**XXXIb**) was synthesized from intermediate **XXIXb** following a two-step reaction sequence of *N*-methylation/ester hydrolysis (0.36 g, 14% yield over two steps). **7f** was synthesized following a two-step reaction sequence: a) amide coupling using intermediate **XXXIb** and (*E*)-3-(methylsulfonyl)prop-2-en-1-amine (**V**) by General Procedure-I, b) THP deprotection by General Procedure-V (white solid, 0.03 g, 7% yield over two steps, m.p. 176 °C). ^1^H NMR (400 MHz, CDCl_3_): *δ* 8.08 (m, 1H), 7.47 (t, *J* = 8.4 Hz, 1H), 7.39 (d, *J* = 7.6 Hz, 1H), 7.09 (t, *J* = 7.4 Hz, 1H), 6.99 – 6.93 (m, 2H), 6.54 – 6.51 (m, 2H), 4.25 (s, 2H), 4.16 – 4.10 (m, 2H), 3.31 (s, 3H), 2.91 (s, 3H), 1.38 (t, *J* = 7.0 Hz, 3H). ^13^C NMR (101 MHz, CDCl_3_): *δ* 170.1, 162.6, 154.8, 144.7, 144.6, 144.3, 132.3, 129.9, 128.9, 124.5, 121.3, 112.1, 93.5, 64.2, 42.9, 37.1, 14.8. HRMS (ESI) *m/z*: [M+H]^+^ calculated for C_18_H_23_N_4_O_5_S: 407.1389, found 407.1386. HPLC purity (254 nm) >99%.

**(*E*)-5-(2-Ethoxyphenyl)-4-methyl-*N*-(3-(methylsulfonyl)allyl)-1*H*-pyrazole-3-carboxamide (8a).** To a stirred a solution of methyl 5-bromo-4-methyl-1*H*-pyrazole-3-carboxylate (**XXXVa**) (1 g, 4.5 mmol, 1.0 eq.) in toluene was added 2-ethoxyphenylboronic acid (1.9 g, 11.4 mmol, 2.5 eq.), NaHCO_3_ (1.9 g, 22.82 mmol, 5.0 eq.), and purged with N_2_ gas for 15 min. Then Pd(PPh_3_)_4_ (316 mg, 0.27 mmol, 0.06 eq.) was added and the reaction was stirred at 90 °C for 16 h. After completion, the reaction mixture was poured into water, extracted with EtOAc, combined organic layers washed with brine, dried over anhydrous Na_2_SO_4_, filtered, and concentrated. The residue was purified using combiflash (eluted at 12–18% EtOAc in hexane) to afford methyl 5-(2-ethoxyphenyl)-4-methyl-1*H*-pyrazole-3-carboxylate (0.7 g, 59% yield). MS (ESI) *m/z*: 261.1 [M+H]^+^. Methyl 5-(2-ethoxyphenyl)-4-methyl-1-(tetrahydro-2*H*-pyran-2-yl)-1*H*-pyrazole-3-carboxylate was synthesized from the intermediate obtained in Step-1 following General Procedure-IV (0.6 g, 65% yield). Step-3: General Procedure-III, 5-(2-ethoxyphenyl)-4-methyl-1-(tetrahydro-2*H*-pyran-2-yl)-1*H*-pyrazole-3-carboxylic acid (**XXXVI**) was synthesized from the intermediate obtained in Step-2 as a white solid (0.5 g, 87% yield). MS (ESI) *m/z*: 331.2 [M+H]^+^. **8a** was synthesized following a two-step reaction sequence: a) amide coupling using intermediate **XXXVI** and (*E*)-3-(methylsulfonyl)prop-2-en-1-amine (**V)** by General Procedure-I, b) THP deprotection following General Procedure-V (white solid, 0.02 g, 10% yield, m.p. 76 °C). ^1^H NMR (400 MHz, DMSO-*d*_6_): *δ* 13.08 (bs, 1H), 8.46 (m, 1H), 7.42 (t, *J* = 7.8 Hz, 1H), 7.29 (d, *J* = 7.6 Hz, 1H), 7.15 (d, *J* = 7.6 Hz, 1H), 7.05 (t, *J* = 7.4 Hz, 1H), 6.85 – 6.79 (m, 1H), 6.71 – 6.67 (m, 1H), 4.09 – 4.06 (m, 4H), 3.03 (s, 3H), 2.16 (s, 3H), 1.30 (d, *J* = 6.8 Hz, 3H). ^13^C NMR (101 MHz, DMSO-*d*_6_): *δ* 163.6, 156.4, 144.3, 143.0, 131.4, 130.7, 130.4, 120.7, 118.8, 115.5, 112.8, 63.9, 42.6, 14.9, 9.9. HRMS (ESI) *m/z*: [M+H]^+^ calculated for C_17_H_22_N_3_O_4_S: 364.1331, found 364.1330. HPLC purity (254 nm) >99%.

**(*E*)-3-(2-Ethoxyphenyl)-4-ethynyl-*N*-(3-(methylsulfonyl)allyl)-1*H*-pyrazole-5-carboxamide (8c)** To a stirred solution of ethyl 5-bromo-1*H*-pyrazole-3-carboxylate (**XXXVb**) (5.0 g, 22.83 mmol, 1.0 eq.) in DMF (50 mL) was added *N*-Iodosuccinimide (6.16 g, 27.40 mmol, 1.2 eq.) and the reaction was stirred at 100 °C for 8 h. After completion, the mixture was diluted with water, extracted with EtOAc (2 x 50mL), washed with brine, dried over anhydrous Na_2_SO_4_, filtered, and concentrated. The residue was purified by combiflash (eluted with 60% EtOAc in hexane) to obtain ethyl 5-bromo-4-iodo-1*H*-pyrazole-3-carboxylate as a white solid (6.1 g, 78% yield). MS (ESI) *m/z*: 344.9 [M+H]^+^. To a stirred solution of ethyl 5-bromo-4-iodo-1*H*-pyrazole-3-carboxylate (6.0 g, 17.4 mmol, 1.0 eq.) in THF (60 mL) was added 2-(trimethylsilyl)ethoxymethyl chloride (2.9 g, 17.39 mmol, 1.0 eq.) and the reaction was stirred at 25 °C for 6 h. After completion, reaction mixture was concentrated, washed with water, and extracted with EtOAc. Organic layer was dried over anhydrous Na_2_SO_4_, filtered, and concentrated. The residue was purified by combiflash (eluted with 10% EtOAc in hexane) to afford ethyl 5-bromo-4-iodo-1-((2-(trimethylsilyl)ethoxy)methyl)-1*H*-pyrazole-3-carboxylate as a liquid (6.5 g, 80% yield). MS (ESI) *m/z*: 419.0 [M+H]^+^. To a stirred solution of ethyl 5-bromo-4-iodo-1-((2-(trimethylsilyl)ethoxy)methyl)-1*H*-pyrazole-3-carboxylate (5.0 g, 10.52 mmol, 1.0 eq.) in DMF (50 mL) at 0 °C was added triethylamine (7.30 mL, 52.6 mmol, 5.0 eq.) and degassed for 10 min. CuI (1.0 g, 5.26 mmol, 0.2 eq.), PdCl_2_(PPh_3_)_2_ (1.47 g, 2.10 mmol, 0.5 eq.), and trimethylsilylacetylene (1.0 g, 5.26 mmol, 0.2 eq.) were then added and the reaction was stirred at 100 °C for 2 h under microwave irradiation. After completion, the mixture was diluted with water, extracted with EtOAc, washed with brine, dried over anhydrous Na_2_SO_4_, filtered, and concentrated. The residue was purified by combiflash (30% EtOAc in hexane) to afford ethyl 5-bromo-1-((2-(trimethylsilyl)ethoxy)methyl)-4-((trimethylsilyl)ethynyl)-1*H*-pyrazole-3-carboxylate as a colorless liquid. (3.5 g, 75% yield). MS (ESI) *m/z*: 447.2 [M+H]^+^. A stirred solution of ethyl 5-bromo-1-((2-(trimethylsilyl)ethoxy)methyl)-4-((trimethylsilyl)ethynyl)-1*H*-pyrazole-3-carboxylate (3.0 g, 6.73 mmol, 1.0 eq.) in dioxane–H_2_O (4:1) (30 mL) was degassed for 10 min. K_3_PO_4_ (3.0 g, 6.73 mmol, 1.0 eq.), Pd(dppf)Cl_2_.DCM (3.0 g, 6.73 mmol, 1.0 eq.) were then added and the reaction was stirred at 100 °C for 8 h. After completion, reaction mixture was diluted with water, extracted with EtOAc, washed with brine, dried over anhydrous Na_2_SO_4_, filtered, and concentrated. The resulting crude was purified by combiflash (30% EtOAc in hexane) to obtain ethyl 5-(2-ethoxyphenyl)-1-((2-(trimethylsilyl)ethoxy)methyl)-4-((trimethylsilyl)ethynyl)-1*H*-pyrazole-3-carboxylate as white solid (2.1 g, 65% yield). MS (ESI) *m/z*: 487.3 [M+H]^+^. Step-5: Ester hydrolysis of 5-(2-ethoxyphenyl)-1-((2-(trimethylsilyl)ethoxy)methyl)-4-((trimethylsilyl)ethynyl)-1*H*-pyrazole-3-carboxylate (2.0 g, 4.11 mmol, 1.0 eq.) following General Procedure-III afforded intermediate **XXXVII** as a brown solid (1.2 g, 75% yield). MS (ESI) *m/z*: 387.2 [M+H]^+^. **8c** was synthesized following a two-step reaction sequence: a) amide coupling using intermediate **XXXVII** and (*E*)-3-(methylsulfonyl)prop-2-en-1-amine (**V**) by General Procedure-II, b) SEM deprotection using HCl in dioxane (white solid, 0.02 g, 36% yield, m.p. 86 °C). ^1^H NMR (400 MHz, DMSO-*d*_6_): *δ* 13.47 (s, 1H), 8.55 (s, 1H), 7.57 (d, *J =* 4.4 Hz, 1H), 7.43 (t, *J =* 7.0 Hz, 1H), 7.16 (d, *J =* 8.0 Hz, 1H), 7.05 (t, *J =* 7.4 Hz, 1H), 6.83 – 6.78 (m, 1H), 6.70 (d, *J =* 15.2 Hz, 1H), 4.08 (t, *J =* 7.0 Hz, 4H), 4.02 (s, 1H), 3.02 (s, 3H), 1.32 (t, *J =* 7.0 Hz, 3H). ^13^C NMR (100 MHz, DMSO-*d*_6_): *δ* 157.0, 156.4, 155.7, 151.5, 143.5, 134.9, 130.9, 130.4, 120.6, 119.9, 112.4, 104.0, 85.4, 75.7, 63.6, 42.8, 41.2, 14.4. HRMS (ESI) *m/z*: [M+H]^+^ calculated for C_18_H_20_N_3_O_4_S: 374.1175, found 374.1174. HPLC purity (254 nm) >96%.

**(*E*)-5-(2-Ethoxyphenyl)-*N*-(3-(methylsulfonyl)allyl)-1*H*-pyrrole-3-carboxamide (12).** Step-1: To a stirring solution of methyl 5-bromo-1*H*-pyrrole-3-carboxylate (**XXXVc**) (0.5 g, 2.45 mmol, 1.0 eq.) and 2-ethoxyphenylboronic acid (0.61 g, 3.67 mmol, 1.5 eq.) in toluene:THF:H_2_O, NaHCO_3_ (0.52 g, 6.12 mmol, 2.5 eq.) was added and the reaction mixture was degassed by N_2_ gas for 15 min. Then Pd(PPh_3_)_4_ (0.056 g, 0.049 mmol, 0.02 eq.) was added and the reaction was stirred at 100 °C for 6 h. After completion, reaction mixture was washed with water, extracted with EtOAc, organic layers dried over anhydrous Na_2_SO_4_, filtered, and concentrated. Purification using combiflash (eluted at 10–15% EtOAc in hexane) afforded methyl 5-(2-ethoxyphenyl)-1*H*-pyrrole-3-carboxylate as a brown solid (0.35 g, 59% yield). MS (ESI) *m/z*: 246.2 [M+H]^+^. Step-2: Ester hydrolysis of methyl 5-(2-ethoxyphenyl)-1*H*-pyrrole-3-carboxylate (0.65 g, 2.65 mmol, 1.0 eq.) following General Procedure-III afforded 5-(2-ethoxyphenyl)-1*H*-pyrrole-3-carboxylic acid (**XXXIIa**) as a white solid (0.25 g, 41% yield). MS (ESI) *m/z*: 232.2 [M+H]^+^. Step-3: Amide coupling using intermediate **XXXIIa** and (*E*)-3-(methylsulfonyl)prop-2-en-1-amine (**V**) by General Procedure-II afforded (*E*)-5-(2-ethoxyphenyl)-*N*-(3-(methylsulfonyl)allyl)-1*H*-pyrrole-3-carboxamide (**12**) as a white solid (0.03 g, 9% yield, m.p. 58 °C). ^1^H NMR (400 MHz, DMSO-*d*_6_): *δ* 11.26 (s, 1H), 8.23 – 8.21 (m, 1H), 7.57 (d, *J* = 8.0 Hz, 1H), 7.44 (s, 1H), 7.22 (s, 1H), 7.19 (d, *J* = 8.0 Hz, 1H), 7.08 – 6.95 (m, 2H), 6.85 – 6.79 (m, 1H), 6.71 – 6.67 (m, 1H), 4.16 – 4.08 (m, 4H), 3.02 (s, 3H), 1.42 (t, *J* = 8 Hz, 3H). ^13^C NMR (101 MHz, DMSO-*d*_6_): *δ* 164.5, 154.8, 144.7, 130.3, 129.1, 127.9, 126.9, 121.7, 121.3, 121.1, 120.1, 113.2, 108.3, 64.0, 42.7, 15.1. HRMS (ESI) *m/z*: [M+H]^+^ calculated for C_17_H_21_N_2_O_4_S: 349.1222, found 349.1217. HPLC purity (254 nm) >99%.

**(*E*)-2-(2-Ethoxyphenyl)-*N*-(3-(methylsulfonyl)allyl)oxazole-5-carboxamide (13).** Step-1: Suzuki coupling between ethyl 2-bromooxazole-5-carboxylate (**XXXVd**) (0.700 g, 3.19 mmol, 1.0 eq.) and 2-ethoxyphenylboronic acid (0.63 g, 3.83 mmol, 1.2 eq.) in toluene (10 mL) and water (0.5 mL), base: K_2_CO_3_ (1.32 g, 9.58 mmol, 3.0 eq.), catalyst: Pd(PPh_3_)_4_ (0.184 g, 0.15 mmol, 0.05 eq.) at 90 °C for 12 h afforded ethyl 2-(2-ethoxyphenyl)oxazole-5-carboxylate as a white solid. (0.5 g, 60% yield). Step-2: Ester hydrolysis of ethyl 2-(2-ethoxyphenyl)oxazole-5-carboxylate (0.5 g, 1.9 mmol, 1.0 eq.) in THF:H_2_O and NaOH following General Procedure-III afforded 2-(2-ethoxyphenyl)oxazole-5-carboxylic acid (**XXXIIb**) as a white solid (0.3 g, 67% yield). Step-3: Amide coupling using intermediate **XXXIIb** and (*E*)-3-(methylsulfonyl)prop-2-en-1-amine (**V**) by General Procedure-I afforded (*E*)-2-(2-ethoxyphenyl)-*N*-(3-(methylsulfonyl)allyl)oxazole-5-carboxamide (**13**) as a white solid (0.03 g, 22% yield, m.p. 132 °C). ^1^H NMR (400 MHz, DMSO-*d*_6_): *δ* 8.93 (t, *J* = 12 Hz, 1H), 7.97 – 7.93 (m, 2H), 7.56 – 7.52 (m, 1H), 7.24 (d, *J* = 8.0 Hz, 1H), 7.10 (t, *J* = 8.0 Hz, 1H), 6.81 (t, *J* = 4.0 Hz, 2H), 4.20 – 4.13 (m, 4H), 3.01 (s, 3H), 1.39 (t, *J* = 12 Hz, 3H). ^13^C NMR (101 MHz, DMSO-*d*_6_): *δ* 161.0, 157.4, 157.2, 145.1, 143.2, 133.3, 131.6, 131.0, 130.9, 121.0, 116.1, 114.3, 64.7, 42.6,14.9. HRMS (ESI) *m/z*: [M+H]^+^ calculated for C_16_H_19_N_2_O_5_S: 351.1222, found 351.1009. HPLC purity (254 nm) >97%.

**(*E*)-2-(2-Ethoxyphenyl)-*N*-(3-(methylsulfonyl)allyl)thiazole-5-carboxamide (14).** Step-1: Suzuki coupling between 2-bromothiazole-5-carboxylic acid (**XXXVe**) (0.5 g, 2.4 mmol, 1.0 eq.) and 2-ethoxyphenylboronic acid (0.6 g, 3.6 mmol, 1.5 eq.) in dioxane (5 mL) and water (2.5 mL), base: Cs_2_CO_3_ (0.8 g, 2.4 mmol, 1.0 eq.), catalyst: Pd(dppf)Cl_2_.DCM (0.4 g, 0.48 mmol, 0.2 eq.) at 80 °C for 4 h afforded 2-(2-ethoxyphenyl)thiazole-5-carboxylic acid (**XXXIII**) (0.13 g, 61% yield). MS (ESI) *m/z*: 250.2 [M+H]^+^.

Step-2: Amide coupling using intermediate **XXXIII** and (*E*)-3-(methylsulfonyl)prop-2-en-1-amine (**V**) by General Procedure-II afforded (*E*)-2-(2-ethoxyphenyl)-*N*-(3-(methylsulfonyl)allyl)thiazole-5-carboxamide (**14**) as a white solid (0.025 g, 58% yield, m.p. 168 °C). ^1^H NMR (400 MHz, DMSO-*d*_6_): *δ* 9.04 (t, *J* = 12 Hz, 1H) 8.52 (s, 1H), 8.35 – 8.32 (m, 1H), 7.53 – 7.48 (m, 1H), 7.27 (d, *J* = 12 Hz, 1H), 7.14 – 7.10 (m, 1H), 6.82 (s, 2H), 4.35 (q, *J* = 20 Hz, 2H), 4.16 (d, *J* = 4.0 Hz, 2H), 3.03 (s, 3H), 1.53 (t, *J* = 16 Hz, 3H). ^13^C NMR (101 MHz, DMSO-*d*_6_): *δ* 164.3, 160.6, 155.6, 142.9, 142.8, 134.3, 131.9, 130.3, 127.7, 121.1, 120.9, 112.9, 64.8, 42.1, 14.7. HRMS (ESI) *m/z*: [M+H]^+^ calculated for C_16_H_19_N_2_O_4_S_2_: 367.0786, found 367.0781. HPLC purity (254 nm) >99%.

**(*E*)-2-(2-Ethoxyphenyl)-5-methyl-*N*-(3-(methylsulfonyl)allyl)-1*H*-imidazole-4-carboxamide (22).** Step-1: Suzuki coupling between ethyl 2-bromo-5-methyl-1*H*-imidazole-4-carboxylate (**XXXVf**) (0.5 g, 2.2 mmol, 1.0 eq.) and 2-ethoxyphenylboronic acid (0.7 g, 4.3 mmol, 2.0 eq.) in toluene/THF/H_2_O, base: NaHCO_3_ (0.5 g, 6.5 mmol, 3.0 eq.), catalyst: Pd(PPh_3_)_4_ (0.08 g, 0.07 mmol, 0.03 eq.) at 90 °C for 12 h afforded ethyl 2-(2-ethoxyphenyl)-5-methyl-1*H*-imidazole-4-carboxylate as colorless liquid (0.45 g, 77% yield). MS (ESI) *m/z*: 275.4 [M+H]^+^. Step-2: Ester hydrolysis of ethyl 2-(2-ethoxyphenyl)-5-methyl-1*H*-imidazole-4-carboxylate (0.45 g, 1.6 mmol, 1.0 eq.) using General Procedure-III afforded 2-(2-ethoxyphenyl)-5-methyl-1*H*-imidazole-4-carboxylic acid (**XXXIV**) as a white solid (0.3 g, 75% yield). MS (ESI) *m/z*: 247.3 [M+H]^+^. Step-3: Amide coupling using intermediate **XXXIV** and (*E*)-3-(methylsulfonyl)prop-2-en-1-amine (**V**) by General Procedure-II afforded (*E*)-2-(2-ethoxyphenyl)-5-methyl-*N*-(3-(methylsulfonyl)allyl)-1*H*-imidazole-4-carboxamide (**22**) as a white solid (0.015 g, 4% yield, m.p. 94 °C). ^1^H NMR (400 MHz, DMSO-*d*_6_): *δ* 11.74 (s, 1H), 8.26 – 8.23 (m, 1H), 8.01 – 7.99 (m, 1H), 7.38 – 7.34 (m, 1H), 7.16 (d, *J* = 8.4 Hz, 1H), 7.05 (t, *J* = 8.0 Hz, 1H), 6.83 – 6.78 (m, 1H), 6.68 – 6.64 (m, 1H), 4.27 – 4.22 (m, 2H), 4.09 (t, *J* = 4 Hz, 2H), 3.01 (s, 3H), 2.53 (s, 3H), 1.42 (t, *J* = 7.2 Hz, 3H). ^13^C NMR (101 MHz, DMSO-*d*_6_): *δ* 163.5, 159.8, 154.9, 144.1, 141.1, 133.8, 131.8, 130.2, 129.9, 129.8, 127.8, 120.7, 118.6, 112.7, 63.8, 46.2, 42.2, 38.4, 14.5. HRMS (ESI) *m/z*: [M+H]^+^ calculated for C_18_H_23_N_2_O_4_S: 364.1379, found 364.1329. HPLC purity (254 nm) >98%.

**(*E*)-4-Chloro-5-(2-ethoxyphenyl)-*N*-(3-(methylsulfonyl)allyl)-1*H*-pyrazole-3-carboxamide (8b).** Step-1: To a stirred solution of ethyl 5-(2-ethoxyphenyl)-1-(tetrahydro-2*H*-pyran-2-yl)-1*H*-pyrazole-3-carboxylate (**VI**) (0.7 g, 2.03 mmol, 1.0 eq.) in DMF (10 mL) at 0 °C was added NCS (0.27 g, 2.03 mmol, 1.0 eq.) and stirred at 70 °C for 12 h. After completion, the mixture was diluted with water (20 mL), extracted with EtOAc (2 x 20 mL), washed with brine, dried over anhydrous Na_2_SO_4_, filtered, and concentrated. The residue was purified by combiflash (eluted with 30% EtOAc in hexane) to obtain ethyl 4-chloro-5-(2-ethoxyphenyl)-1-(tetrahydro-2*H*-pyran-2-yl)-1*H*-pyrazole-3-carboxylate as a white solid (0.7 g, 91% yield). MS (ESI) *m/z*: 379.3 [M+H]^+^. Step-2: Ester hydrolysis of ethyl 4-chloro-5-(2-ethoxyphenyl)-1-(tetrahydro-2*H*-pyran-2-yl)-1*H*-pyrazole-3-carboxylate (0.7 g, 1.85 mmol, 1.0 eq.) following General Procedure-III afforded 4-chloro-5-(2-ethoxyphenyl)-1-(tetrahydro-2*H*-pyran-2-yl)-1*H*-pyrazole-3-carboxylic acid (**XXXVIII**) as a white solid (0.6 g, 93% yield). MS (ESI) *m/z*: 267.2 [M+H]^+^. Step-3: Following a two-step reaction sequence of a) amide coupling using intermediate **XXXVII**I and (*E*)-3-(methylsulfonyl)prop-2-en-1-amine (**V**) by General Procedure-II, b) THP deprotection by General Procedure-V afforded (*E*)-4-chloro-5-(2-ethoxyphenyl)-*N*-(3-(methylsulfonyl)allyl)-1*H*-pyrazole-3-carboxamide (**8b**) as white solid (0.18 g, 22% yield over two steps, m.p. 112 °C). ^1^H NMR (400 MHz, DMSO-*d*_6_): *δ* 13.63 (s, 1H), 8.64 (s, 1H), 7.49 – 7.39 (m, 2H), 7.19 – 7.06 (m, 2H), 6.83 (dd, *J* = 4.0 Hz, 2H), 4.09 – 4.08 (m, 4H), 3.02 (s, 3H), 1.32 (t, *J* = 8 Hz, 3H). ^13^C NMR (101 MHz, DMSO-*d*_6_): *δ* 161.3, 156.6, 143.9, 141.2, 138.2, 131.6, 131.3, 130.6, 120.8, 116.3, 113.0, 108.4, 64.1, 42.6, 14.9. HRMS (ESI) *m/z*: [M+H]^+^ calculated for C_16_H_18_ClN_3_O_4_S: 384.0785, found 384.0786. HPLC purity (254 nm) >99%.

**(*E*)-4-Cyano-5-(2-ethoxyphenyl)-*N*-(3-(methylsulfonyl)allyl)-1*H*-pyrazole-3-carboxamide (8d).** Step-1: To a stirred a solution of ethyl 5-(2-ethoxyphenyl)-1-(tetrahydro-2*H*-pyran-2-yl)-1*H*-pyrazole-3-carboxylate (**VI**) (1.0 g, 2.9 mmol, 1.0 eq.) in DMF was added NBS (0.51 g, 2.9 mmol, 1.0 eq.) and the reaction was stirred at 25 °C for 16 h. After completion, the reaction mixture was poured into water, extracted with EtOAc, combined organic layers washed with brine, dried over anhydrous Na_2_SO_4_, filtered, and concentrated. The residue was purified using combiflash (eluent 20–25% EtOAc in hexane) to afford ethyl 4-bromo-5-(2-ethoxyphenyl)-1-(tetrahydro-2*H*-pyran-2-yl)-1*H*-pyrazole-3-carboxylate (0.8 g, 65% yield). MS (ESI) *m/z*: 425.4 [M+H]^+^. Step-2: To a stirred a solution of ethyl 4-bromo-5-(2-ethoxyphenyl)-1-(tetrahydro-2*H*-pyran-2-yl)-1*H*-pyrazole-3-carboxylate (0.75 g, 1.77 mmol, 1.0 eq.) in DMF were added Zn(CN)_2_ (0.415 g, 3.54 mmol, 2.0 eq.), Zn dust (0.23 g, 3.54 mmol, 2.0 eq.), and purged with N_2_ gas for 15 min. This was followed by the addition of Pd_2_(dba)_3_ (0.16 g, 0.17 mmol, 0.1 eq.), dppf (0.19 g, 0.35 mmol, 0.2 eq.) and purged with N_2_ gas for 15 min. Reaction mixture was stirred at 100 °C for 16 h. After completion, the reaction mixture was quenched with water and EtOAc, then filtered through celite bad. The filtrate was extracted with EtOAc, washed with brine, dried over anhydrous Na_2_SO_4_, filtered, and concentrated. The residue was purified using combiflash (eluent 40–45% EtOAc in hexane) to obtain ethyl 4-cyano-5-(2-ethoxyphenyl)-1-(tetrahydro-2*H*-pyran-2-yl)-1*H*-pyrazole-3-carboxylate (0.16 g, 25% yield). MS (ESI) *m/z*: 370.3 [M+H]^+^. Step-3: Ester hydrolysis of ethyl 4-cyano-5-(2-ethoxyphenyl)-1-(tetrahydro-2*H*-pyran-2-yl)-1*H*-pyrazole-3-carboxylate (0.27 g, 0.73 mmol, 1.0 eq.) using General Procedure-III afforded 4-cyano-5-(2-ethoxyphenyl)-1-(tetrahydro-2*H*-pyran-2-yl)-1*H*-pyrazole-3-carboxylic acid (**XXXIX**) (0.24 g, 96% yield). MS (ESI) *m/z*: 340.2 [M+H]^+^. Step-4: A two-step reaction sequence of a) amide coupling using intermediate **XXXIX** and (*E*)-3-(methylsulfonyl)prop-2-en-1-amine (**V**) by General Procedure-II, b) THP deprotection by General Procedure-V afforded (*E*)-4-cyano-5-(2-ethoxyphenyl)-*N*-(3-(methylsulfonyl)allyl)-1*H*-pyrazole-3-carboxamide (**8d**) as a white solid (0.02 g, 8% yield over two steps, m.p. 138 °C). ^1^H NMR (400 MHz, DMSO-*d*_6_): *δ* 8.74 (s, 1H), 8.21 (s, 1H), 7.84 (d, *J* = 8.8 Hz, 1H), 7.34 (d, *J* = 8.8 Hz, 1H), 7.24 (s, 1H), 6.85-6.79 (m, 1H), 6.73 – 6.69 (m, 1H), 4.32 – 4.27 (m, 2H), 4.12 (s, 2H), 3.02 (s, 3H), 1.45 (t, *J* = 6.8 Hz, 3H). ^13^C NMR (101 MHz, DMSO-*d*_6_): *δ* 161.2, 158.7, 143.8, 134.2, 131.9, 130.6, 119.3, 114.3, 106.8, 103.4, 65.1, 42.6, 14.8. HRMS (ESI) *m/z*: [M+H]^+^ calculated for C_17_H_19_N_4_O_4_S: 375.1127, found 375.1130. HPLC purity (254 nm) >98%.

**(*E*)-5-(2-Ethoxyphenyl)-*N*-(3-(methylsulfonyl)allyl)oxazole-2-carboxamide (15).** Step-1: To a mixture of 1-(2-ethoxyphenyl)ethan-1-one (**Ia**) (1.5 g, 0.009 mmol, 1.0 eq.) and ethyl 2-isocyanoacetate (1.03 g, 0.0091 mmol, 1.2 eq.) in DMSO (15 mL) was added I_2_ (2.3 g, 0.009 mmol, 2.0 eq.) and the reaction was stirred at 130 °C for 2 h. After completion, the reaction mixture was quenched with water (30 mL), extracted with EtOAc (2 x 50 mL), and washed with Na_2_S_2_O_3_ solution. The combined organic layers were dried over anhydrous Na_2_SO_4_, filtered, and concentrated. The residue was purified by combiflash (eluent at 10–15% EtOAc in hexane) to obtain ethyl 5-(2-ethoxyphenyl)oxazole-2-carboxylate as a yellow solid (0.4 g, 18% yield). MS (ESI) *m/z*: 262.2 [M+H]^+^. Step-2: Ester hydrolysis of ethyl 5-(2-ethoxyphenyl)oxazole-2-carboxylate (0.35 g, 1.339 mmol, 1.0 eq.) following General Procedure-III afforded 5-(2-ethoxyphenyl)oxazole-2-carboxylic acid (XL) as a white solid (0.2 g, 64% yield). MS (ESI) *m/z*: 234.1 [M+H]^+^. Step-3: Amide coupling using intermediate **XL** and (*E*)-3-(methylsulfonyl)prop-2-en-1-amine (**V**) by General Procedure-II afforded (*E*)-5-(2-ethoxyphenyl)-*N*-(3-(methylsulfonyl)allyl)oxazole-2-carboxamide (**15**) as a white solid (0.02 g, 9% yield, m.p. 86 °C). ^1^H NMR (400 MHz, DMSO-*d*_6_): *δ* 9.32 (s, 1H), 7.88 (d, *J* = 8.0 Hz, 1H), 7.68 (s, 1H), 7.43 (t, *J* = 14.0 Hz, 1H), 7.20 (d, *J* = 8.4 Hz, 1H), 7.13 (t, *J* = 15.2 Hz, 1H), 6.80 (s, 2H), 4.25 (dd, *J*_1_ = 6.8 Hz, *J*_2_ = 6.8 Hz, 2H), 4.13 (d, *J* = 4.0 Hz, 2H), 3.00 (s, 3H), 1.49 (t, *J* = 14.0 Hz, 3H). ^13^C NMR (101 MHz, DMSO-*d*_6_): *δ* 155.6, 155.3, 153.4, 149.7, 142.8, 131.1, 130.9, 126.9, 126.6, 121.3, 115.8, 112.9, 64.4, 42.6, 15.0. HRMS (ESI) *m/z*: [M+H]^+^ calculated for C_16_H_19_N_2_O_5_S: 351.1015, found 351.1016. HPLC purity (254 nm) >98%.

**(*E*)-5-(2-Ethoxyphenyl)-2-methyl-*N*-(3-(methylsulfonyl)allyl)-1*H*-pyrrole-3-carboxamide (21).** Step-1: To a stirred solution of 1-(2-ethoxyphenyl) ethan-1-one (**Ia**) (1.0 g, 6.09 mmol, 1.0 eq.) in EtOAc (10 mL), CuBr_2_ (2.03 g, 9.13 mmol, 1.5 eq.) was added and reaction stirred at 90 °C for 16 h. After completion, reaction mixture was filtered through celite pad, filtrate diluted with water (25 mL), and extracted with EtOAc (2 × 50 mL). The organic layers were combined, washed with brine, dried over anhydrous Na_2_SO_4_, filtered, and concentrated. The residue was purified by combiflash (eluent 3% EtOAc in hexane) to afford 2-bromo-1-(2-ethoxyphenyl) ethan-1-one as a white solid (0.65 g, 44% yield). Step-2: To a stirred solution of 2-bromo-1-(2-ethoxyphenyl) ethan-1-one (0.65 g, 2.67 mmol, 1.0 eq.) in THF (6.5 mL) were added NaH (83.4 mg, 3.47 mmol, 1.3 eq.) and ethyl 3-oxobutanoate (521.9 mg, 4.01 mmol, 1.5 eq.) at 0 °C. The reaction was stirred at 25 °C for 5 h. After completion, the reaction mixture was diluted with water (50 mL), extracted with EtOAc (2 × 100 mL), washed with brine (50 mL), dried over anhydrous Na_2_SO_4_, filtered, and concentrated. The crude was purified by combiflash (eluent 20% EtOAc in hexane) to afford ethyl 2-acetyl-4-(2-ethoxyphenyl)-4-oxobutanoate as a white solid (0.45 g, 57% yield). MS (ESI) *m/z*: 293.4 [M+H]^+^. Step-3: To a stirred solution of ethyl 2-acetyl-4-(2-ethoxyphenyl)-4-oxobutanoate (0.45 g, 1.53 mmol, 1.0 eq.) in EtOH (4.5 mL) was added ammonium acetate (0.24 g, 3.07 mmol, 2.0 eq.) and the reaction was stirred at 90 °C for 4 h. After completion, the reaction mixture was concentrated, diluted with water, and extracted with EtOAc (3 x 50 mL). The combined organic layers were washed with brine (50 mL), dried over anhydrous Na_2_SO_4_, filtered, and concentrated. The residue was purified by combiflash (eluent 15% EtOAc in hexane) to obtain ethyl 5-(2-ethoxyphenyl)-2-methyl-1*H*-pyrrole-3-carboxylate as a white solid (0.35 g, 83% yield). MS (ESI) *m/z*: 274.4 [M+H]^+^. Step-4: Ester hydrolysis of ethyl 5-(2-ethoxyphenyl)-2-methyl-1*H*-pyrrole-3-carboxylate (0.4 g, 1.46 mmol, 1.0 eq.) following General Procedure-III afforded 5-(2-ethoxyphenyl)-2-methyl-1*H*-pyrrole-3-carboxylic acid (**XLI**) as a white solid (0.22 g, 61% yield). MS (ESI) *m/z*: 246.1 [M+H]^+^. Step-5: Amide coupling using intermediate **XLI** and (*E*)-3-(methylsulfonyl)prop-2-en-1-amine (**V**) by General Procedure-II afforded (*E*)-5-(2-ethoxyphenyl)-2-methyl-*N*-(3-(methylsulfonyl)allyl)-1*H*-pyrrole-3-carboxamide (**21**) as a white solid (0.06 g, 10% yield, m.p. 121 °C). ^1^H NMR (400 MHz, DMSO-*d*_6_): *δ* 10.98 (s, 1H), 7.98 (s, 1H), 7.53 (d, *J* = 7.6 Hz, 1H), 7.18 (t, *J* = 7.2 Hz, 1H), 7.05 (d, *J* = 8.0 Hz, 1H), 6.96 – 6.94 (m, 2H), 6.82 – 6.79 (m, 1H), 6.67 (d, *J* = 15.2 Hz, 1H), 4.15 – 4.10 (m, 2H), 4.04 (s, 2H), 3.01 (s, 3H), 2.47 (s, 3H), 1.44 (t, *J* = 13.6 Hz, 3H). ^13^C NMR (101 MHz, DMSO-*d*_6_): *δ* 165.5, 154.7, 144.9, 133.6, 130.2, 127.4, 126.6, 126.1, 121.6, 120.9, 114.9, 113.2, 108.4, 63.9, 42.7, 15.0, 13.2. HRMS (ESI) *m/z*: [M+H]^+^ calculated for C_18_H_23_N_2_O_4_S: 364.1379, found 364.1329. HPLC purity (254 nm) >99%.

**(*E*)-3-(2-Ethoxyphenyl)-*N*-(3-(methylsulfonyl)allyl)isothiazole-5-carboxamide (11).** Step-1: To a stirred solution of 2-ethoxybenzamide (**XLIIa**) (2.0 g, 12.1 mmol, 1.0 eq.) in dioxane (40 mL) was added chlorocarbonylsulfenyl chloride (1.22 mL, 14.53 mmol, 1.2 eq.) and the reaction was stirred at 90 °C for 3 h. After completion, the reaction mixture was concentrated and the residue was purified by combiflash (eluent 20% EtOAc in hexane) to give 5-(2-ethoxyphenyl)-1,3,4-oxathiazol-2-one as a white solid (2.0 g, 74% yield). Step-2: To a stirred solution of 5-(2-ethoxyphenyl)-1,3,4-oxathiazol-2-one (2.0 g, 8.96 mmol, 1.0 eq.) in CHCl_3_ (9 mL) was added ethyl propiolate (8.78 g, 89.6 mmol, 10.0 eq.) and the reaction was stirred at 160 °C for 1 h under microwave irradiation. After completion, the reaction mixture was concentrated and the residue was purified by combiflash to afford ethyl 3-(2-ethoxyphenyl) isothiazole-5-carboxylate as a white solid (0.53 g, 21% yield). MS (ESI) *m/z*: 278.3 [M+H]^+^. Step-3: Ester hydrolysis of ethyl 3-(2-ethoxyphenyl) isothiazole-5-carboxylate (0.5 g, 1.8 mmol, 1.0 eq.) following General Procedure-III afforded 3-(2-ethoxyphenyl)isothiazole-5-carboxylic acid (**XLIII**) as a white solid (0.3 g, 67% yield). MS (ESI) *m/z*: 250.3 [M+H]^+^. Step-4: Amide coupling between intermediate **XLIII** and (*E*)-3-(methylsulfonyl)prop-2-en-1-amine (**V**) following General Procedure-II afforded (*E*)-3-(2-ethoxyphenyl)-*N*-(3-(methylsulfonyl)allyl)isothiazole-5-carboxamide (**11**) as a white solid (0.04 g, 14% yield, m.p. 104 °C). ^1^H NMR (400 MHz, DMSO-*d*_6_): *δ* 9.32 (t, *J* = 5.6 Hz, 1H), 8.40 (s, 1H), 7.87 – 7.85 (m, 1H), 7.46 – 7.42 (m, 1H), 7.18 (d, *J* = 8.4 Hz, 1H), 7.06 (d, *J* = 7.2 Hz, 1H), 6.81 (s, 2H), 4.22 – 4.17 (m, 4H), 3.01 (s, 3H), 1.41 (t, *J* = 6.8 Hz, 3H). ^13^C NMR (101 MHz, DMSO-*d*_6_): *δ* 165.9, 162.6, 159.8, 155.9, 142.9, 131.6, 130.9, 130.4, 125.3, 123.7, 121.2, 113.5, 64.4, 42.6, 14.9. HRMS (ESI) *m/z*: [M+H]^+^ calculated for C_16_H_19_N_2_O_4_S_2_: 367.0786, found 367.0781. HPLC purity (254 nm) >95%.

**(*E*)-5-(2-Ethoxyphenyl)-*N*-(3-(methylsulfonyl)allyl)-1,3,4-oxadiazole-2-carboxamide (18).** Step-1: To a stirred solution of 2-ethoxybenzohydrazide (**XLIIb**) (3.4 g, 18.89 mmol, 1.0 eq.) in DCM (40 mL) at 0 °C were added triethylamine (7.6 mL, 56.7 mmol, 3.0 eq.), ethyl 2-chloro-2-oxoacetate (2.5 mL, 22.7 mmol, 1.2 eq.) and the reaction was stirred 25 °C for 4 h. After completion, the reaction mixture was quenched with NaHCO_3_ (100 mL), extracted with DCM (100 mL x 3), dried over anhydrous Na_2_SO_4_, filtered, and concentrated. The residue was purified by combiflash (eluent 0–80% EtOAc in hexane) to furnish ethyl 2-(2-(2-ethoxybenzoyl)hydrazineyl)-2-oxoacetate (4.0 g, 76% yield). MS (ESI) *m/z*: 281.2 [M+H]^+^. Step-2: A stirred solution of ethyl 2-(2-(2-ethoxybenzoyl)hydrazineyl)-2-oxoacetate (1.8 g, 6.53 mmol, 1.0 eq.) and P_2_O_5_ (6.49 g, 45.7 mmol, 7.0 eq.) in toluene (20 mL) was heated at 100 °C for 12 h. After completion, the reaction mixture was extracted with EtOAc (25 mL x 3), dried over anhydrous Na_2_SO_4_, filtered, and concentrated. The residue was purified by combiflash (eluent 0–100% EtOAc in hexane) to afford ethyl 5-(2-ethoxyphenyl)-1,3,4-oxadiazole-2-carboxylate (0.4 g, 24% yield). Step-3: Ester hydrolysis of ethyl 5-(2-ethoxyphenyl)-1,3,4-oxadiazole-2-carboxylate (1.0 g, 3.9 mmol, 1.0 eq.) following General Procedure-III afforded 5-(2-ethoxyphenyl)-1,3,4-oxadiazole-2-carboxylic acid (**XLIV**) as a white solid (0.7 g, 72% yield). Step-4: Amide coupling of intermediate **XLIV** and (*E*)-3-(methylsulfonyl)prop-2-en-1-amine (**V**) by General Procedure-II afforded (*E*)-5-(2-ethoxyphenyl)-*N*-(3-(methylsulfonyl)allyl)-1,3,4-oxadiazole-2-carboxamide (**18**) as a white solid (0.05 g, 12% yield, m.p. 125°C). ^1^H NMR (400 MHz, DMSO-*d*_6_): *δ* 9.62 (t, *J* = 5.6 Hz, 1H), 7.93 (d, *J* = 8.0 Hz, 1H), 7.64 (dd, *J* = 8.8 Hz 1H), 7.29 (d, *J* = 8.4 Hz 1H), 7.17 (t, *J* = 7.6 Hz, 1H), 6.86 – 6.78 (m, 2H), 4.22 (q, *J* = 6.8 Hz, 4H), 3.01 (s, 3H), 1.38 (t, *J* = 7.2 Hz, 3H). ^13^C NMR (101 MHz, DMSO-*d*_6_): *δ* 164.4, 158.7, 157.5, 153.8, 142.3, 134.6, 131.1, 131.0, 121.3, 114.3, 112.4, 64.8, 42.6, 14.9. HRMS (ESI) *m/z*: [M+H]^+^ calculated for C_15_H_18_N_3_O_5_S: 352.0967, found 352.0968. HPLC purity (254 nm) >95%.

**(*E*)-5-(2-Ethoxyphenyl)-*N*-(3-(methylsulfonyl)allyl)furan-2-carboxamide (16).** Step-1: To a stirred solution of 2-ethoxyphenylboronic acid (**XLV**) (0.5 g, 3.03 mmol, 1.0 eq.) in 1,4-dioxane:H_2_O were added methyl 5-bromofuran-2-carboxylate (0.62 g, 3.03 mmol, 1.0 eq.) and K_2_CO_3_ (1.3 g, 9.09 mmol, 3.0 eq.) and purged with N_2_ gas for 10 min. Then PdCl_2_(PPh_3_)_2_ (0.12 g, 0.15 mmol, 0.05 eq.) was added and the reaction was stirred at 90 °C for 6 h. After completion, the reaction mixture was poured into water (20 mL), extracted with EtOAc (3 x 40 mL), dried over anhydrous Na_2_SO_4_, filtered, and concentrated. The residue was purified by combiflash (eluent 20% EtOAc in hexane) to afford methyl 5-(2-ethoxyphenyl) furan-2-carboxylate as a white solid (0.5 g, 68% yield). MS (ESI) *m/z*: 247.1 [M+H]^+^. Step-2: Ester hydrolysis of methyl 5-(2-ethoxyphenyl) furan-2-carboxylate (0.3 g, 1.2 mmol, 1.0 eq.) following General Procedure-III afforded 5-(2-ethoxyphenyl)furan-2-carboxylic acid (**XLVI**) as a white solid (0.2 g, 72% yield). MS (ESI) *m/z*: 233.2 [M+H]^+^. Step-3: Amide coupling of intermediate (**XLVI**) and (*E*)-3-(methylsulfonyl)prop-2-en-1-amine (**V**) by General Procedure-II afforded (*E*)-5-(2-ethoxyphenyl)-*N*-(3-(methylsulfonyl)allyl)furan-2-carboxamide (**16**) as a white solid (0.1 g, 53% yield, m.p. 153 °C). ^1^H NMR (400 MHz, DMSO-*d*_6_): *δ* 7.89 (d, *J* = 6.4 Hz, 1H), 7.34 (q, *J* = 27.6 Hz, 1H), 7.12 – 7.11 (d, *J* = 3.6 Hz, 1H), 7.07 – 6.98 (m, 3H), 6.73 (s, 1H), 6.61 – 6.57 (d, *J* =15.2 Hz, 1H), 4.35 (s, 2H), 4.22 – 4.16 (q, *J* = 20.8 Hz, 2H), 2.97 (s,3H), 1.61 (q, *J* = 27.6 Hz, 3H). ^13^C NMR (101 MHz, DMSO-*d*_6_): *δ* 158.6, 155.7, 152.7, 144.8, 144.2, 130.1, 129.7, 126.4, 120.6, 118.3, 117.5, 112.3, 112.0, 64.0, 42.8, 39.0, 14.9. HRMS (ESI) *m/z*: [M+H]^+^ calculated for C_17_H_20_NO_5_S: 351.1062, found 351.1061. HPLC purity (254 nm) >97%.

**(*E*)-5-(2-Ethoxyphenyl)-*N*-(3-(methylsulfonyl)allyl)thiophene-2-carboxamide (17).** Step-1: To a stirred solution of 2-ethoxyphenylboronic acid (**XLV**) (0.25 g, 1.15 mmol, 1.0 eq.) in 1,4-dioxane:H_2_O were added 5-bromothiophene-2-carboxylic acid (313.6 mg, 1.151 mmol, 1.0 eq.), K_2_CO_3_ (627.3 mg, 4.545 mmol, 3.0 eq.) and purged with N_2_ gas for 10 min. Then PdCl_2_(PPh_3_)_2_ (0.06 g, 0.08 mmol, 0.05 eq.) was added and the reaction was stirred at 90 °C for 6 h. After completion, the reaction mixture was poured into cold water (20 mL), extracted with EtOAc (30 ml x 3), organic layer dried over anhydrous Na_2_SO_4_, filtered, and concentrated. The residue was purified by combiflash to obtain 5-(2-ethoxyphenyl) thiophene-2-carboxylic acid (**XLVII**) as a white solid (0.24 g, 64% yield). MS (ESI) *m/z*: 249.1 [M+H]^+^. Step-2: Amide coupling of intermediate (**XLVII**) and (*E*)-3-(methylsulfonyl)prop-2-en-1-amine (**V**) by General Procedure-II afforded (*E*)-5-(2-ethoxyphenyl)-*N*-(3-(methylsulfonyl)allyl)thiophene-2-carboxamide (**17**) as a white solid (0.07 g, 32% yield, m.p. 118 °C). ^1^H NMR (400 MHz, DMSO-*d*_6_): *δ* 7.71 (d, *J* = 6.4 Hz, 1H), 7.57 – 7.48 (m, 3H), 7.34 (d, *J* = 8.4 Hz, 1H), 7.04 (q, *J* = 17.2 Hz, 1H), 6.60 – 6.55 (m, 1H), 6.37 (s, 1H), 4.33 (d, *J* = 4.0 Hz, 2H), 4.25 (q, *J* = 20.8 Hz, 2H), 2.97 (s, 3H), 1.60 (q, *J* = 20.4 Hz, 3H). ^13^C NMR (101 MHz, DMSO-*d*_6_): *δ* 162.5, 155.1, 145.0, 144.6, 136.1, 132.0, 129.9, 129.6, 128.5, 128.3, 125.3, 122.2, 120.9, 112.5, 64.5, 42.9, 39.7, 14.9. HRMS (ESI) *m/z*: [M+H]^+^ calculated for C_17_H_20_NO_4_S_2_: 366.0834, found 366.0828. HPLC purity (254 nm) >99%.

**(*E*)-5-(2-Ethoxyphenyl)-*N*-(3-(methylsulfonyl)allyl)-1,2,4-oxadiazole-3-carboxamide (19).** Step-1: To a stirred solution of ethyl carbonocyanidate (**XLVIII**) (2.5 g, 25.22 mmol, 1.0 eq.) in DMF (25 mL) were added hydroxylamine hydrochloride (2.6 g, 37.84 mmol, 1.5 eq.) and K_3_PO_4_ (10.69 g, 50.45 mmol, 2.0 eq.). The reaction was stirred at 90 °C for 1 h. After completion, the reaction mixture was poured into water (30 mL), extracted with EtOAc (2 x 50 mL), dried over anhydrous Na_2_SO_4_, filtered, and concentrated to afford ethyl 2-amino-2-(hydroxyimino)acetate as a yellow solid (2.6 g, 78% yield). Step-2: To a stirred solution of ethyl 2-amino-2-(hydroxyimino)acetate (2.6 g, 19.7 mmol, 1.0 eq.) in DMF (12 mL) was added 2-ethoxybenzoyl chloride (4.8 g, 19.7 mmol, 1.0 eq.) at 0 °C. Reaction mixture was stirred at 110 °C for 16 h. After completion, the reaction was diluted with water (30 mL), extracted with EtOAc (30 x 3 mL), dried over anhydrous Na_2_SO_4_, filtered, and concentrated. The residue was purified using combiflash (eluent 10% EtOAc in hexane) to furnish ethyl 5-(2-ethoxyphenyl)-1,2,4-oxadiazole-3-carboxylate as a white solid (0.4 g, 8% yield). Step-3: Ester hydrolysis of ethyl 5-(2-ethoxyphenyl)-1,2,4-oxadiazole-3-carboxylate (0.3 g, 1.15 mmol, 1.0 eq.) following General Procedure-III afforded 5-(2-ethoxyphenyl)-1,2,4-oxadiazole-3-carboxylic acid (**XLIX**) as a white solid (0.25 g, 94% yield). MS (ESI) *m/z*: 235.1 [M+H]^+^. Step-4: Amide coupling of intermediate (**XLIX**) and (*E*)-3-(methylsulfonyl)prop-2-en-1-amine (**V**) by General Procedure-I afforded (*E*)-5-(2-ethoxyphenyl)-*N*-(3-(methylsulfonyl)allyl)-1,2,4-oxadiazole-3-carboxamide (**19**) as a white solid (0.03 g, 15% yield, m.p. 56 °C). ^1^H NMR (400 MHz, DMSO-*d*_6_): *δ* 9.38 (s, 1H), 8.05 (d, *J* = 7.6 Hz, 1H), 7.70 (t, *J* = 15.6 Hz, 1H), 7.33 (d, *J* = 7.6 Hz, 1H), 7.19 (t, *J* = 13.2 Hz,1H) 6.80 (s, 2H), 4.25 (d, *J* =5.2 Hz, 2H), 4.15 (s, 2H), 3.02 (d, *J* = 2 Hz, 3H), 1.41 (q, *J* = 11.6 Hz, 3H). ^13^C NMR (101 MHz, DMSO-*d*_6_): *δ* 176.3, 163.7, 158.1, 156.9, 142.5, 135.7, 131.9, 131.0, 121.3, 114.4, 112.5, 64.9, 42.1, 14.9. HRMS (ESI) *m/z*: [M+H]^+^ calculated for C_15_H_18_N_3_O_5_S: 352.0967, found 352.0965. HPLC purity (254 nm) >96%.

**(*E*)-5-(2-Ethoxyphenyl)-*N*-(3-(methylsulfonyl)allyl)isoxazole-3-carboxamide (9).** Step-1: To a stirred solution of diethyl 2-oxosuccinate (**L**) (3.0 g, 14.24 mmol, 1.0 eq.) and hydroxylamine hydrochloride (1.1 g, 14.24 mmol, 1.0 eq.) in water (30 mL) was added NaHCO_3_ (2.67 g, 28.48 mmol, 2.0 eq.) and the reaction was stirred at 25 °C for 16 h. After completion, the reaction mixture was poured into water (30 mL), pH maintained with 1N HCl solution (pH = 5), extracted with EtOAc (2 x 50 mL), and washed with Na₂S₂O₃ solution. The combined organic layers were dried over anhydrous Na_2_SO_4_, filtered, and concentrated to afford ethyl 5-oxo-4,5-dihydroisoxazole-3-carboxylate as colorless liquid (1.5 g, 68% yield). MS (ESI) *m/z*: 157.9 [M+H]^+^. Step-2: To a stirred solution of ethyl 5-oxo-4,5-dihydroisoxazole-3-carboxylate (1.2 g, 7.64 mmol, 1.0 eq.) in DCM (12 mL) were added triethylamine (0.32 mL, 22.929 mmol, 3.0 eq.), POBr_3_ (4.3 g, 15.3 mmol, 2.0 eq.), and the reaction was stirred at 25 °C for 6 h. After completion, the reaction was diluted with water (30 mL), extracted with EtOAc (30 x 3 mL), dried over anhydrous Na_2_SO_4_, filtered, and concentrated to afford ethyl 5-bromoisoxazole-3-carboxylate (**LI**) as a white solid (0.95 g, 57% yield). Step-3: To a stirred solution of ethyl 5-bromoisoxazole-3-carboxylate (0.7 g, 3.18 mmol, 1.0 eq.) in 1,4-dioxane:water were added 2-ethoxyphenylboronic acid (0.53 g, 3.18 mmol, 1.0 eq.), K_2_CO_3_ (1.32 g, 9.55 mmol, 3.0 eq.), and purged with N_2_ gas for 10 min. Then PdCl_2_(PPh_3_)_2_ (0.11 g, 0.16 mmol, 0.05 eq.) was added and the reaction was stirred at 100 °C for 4 h. After completion, the reaction was diluted with water (20 mL), extracted with EtOAc (20 x 3 mL), dried over anhydrous Na_2_SO_4_, filtered, and concentrated. The residue was purified by combiflash (eluent 50% EtOAc in hexane) to obtain ethyl 5-(2-ethoxyphenyl)isoxazole-3-carboxylate (0.35 g, 42% yield). Step-4: Ester hydrolysis of ethyl 5-(2-ethoxyphenyl)isoxazole-3-carboxylate (0.3 g, 1.29 mmol, 1.0 eq.) following General Procedure-III afforded 5-(2-ethoxyphenyl)isoxazole-3-carboxylic acid (**LI**) as a white solid (0.2 g, 75% yield). MS (ESI) *m/z*: 234.2 [M+H]^+^. Step-5: To a stirred solution of 5-(2-ethoxyphenyl)isoxazole-3-carboxylic acid (**LII**) (0.12 g, 0.52 mmol, 1.0 eq.) in DMF (3 mL) were added pyridine (0.2 mL, 2.58 mmol, 5.0 eq.), (*E*)-3-(methylsulfonyl)prop-2-en-1-amine (**V**) (0.07 g, 0.52 mmol, 1.0 eq.), and POCl_3_ (0.18 mL, 0.772 mmol, 1.5 eq.). The reaction was stirred at 25 °C for 1 h. After completion, the reaction mixture was poured into water (20 mL), extracted with EtOAc (3 x 50 mL), dried over anhydrous Na_2_SO_4_, filtered, and concentrated. The residue was purified by combiflash (eluent 80% EtOAc in hexane) to furnish (*E*)-5-(2-ethoxyphenyl)-*N*-(3-(methylsulfonyl)allyl)isoxazole-3-carboxamide (**9**) as a white solid (0.06 g, 34% yield, m.p. 149 °C). ^1^H NMR (400 MHz, DMSO-*d*_6_): *δ* 9.37 (s, 1H), 7.82 (dd, *J* = 1.6Hz, 1H), 7.53 – 7.47 (m, 2H), 7.23 (d, *J* = 8.4 Hz, 1H), 7.10 – 7.06 (m, 1H), 6.82 (t, *J* = 4.4 Hz, 1H), 4.20 – 4.15 (m, 4H), 3.02 (s, 3H), 1.42 (t, *J* = 13.6 Hz, 3H). ^13^C NMR (101 MHz, DMSO-*d*_6_): *δ* 163.2, 160.7, 156.7, 156.3, 142.6, 132.6, 130.9, 129.4, 121.2, 116.8, 113.6, 107.9, 64.4, 42.6, 14.9. HRMS (ESI) *m/z*: [M+H]^+^ calculated for C_16_H_19_N_2_O_5_S: 351.1127, found 351.1008. HPLC purity (254 nm) >96%.

**(*E*)-*N*-(3-(Cyclopropylsulfonyl)allyl)-5-(2-ethoxyphenyl)-1*H*-pyrazole-3-carboxamide (25a).** Step-1: Boc-protection of propargylamine (**LIIIa**) following the literature procedure^20^ furnished *N*-Boc propargylamine in quantitative yield. Step-2: To a solution of *N*-Boc propargylamine (5.0 g, 32.2 mmol, 1.0 eq.) in toluene (60 mL) were added tributyl tin hydride (10.3 g, 35.4 mmol, 1.1 eq.) and AIBN (264.4 mg, 1.61 mmol, 0.05 eq.). The reaction was stirred at 120 °C for 2 h. After completion, the reaction mixture was concentrated and the crude was purified by combiflash (eluent 3–5% EtOAc in hexane) to afford *tert*-butyl (*E*)-(3-(tributylstannyl)allyl)carbamate (8.0 g, 56% yield). Step-3: To a solution of *tert*-butyl (*E*)-(3-(tributylstannyl)allyl)carbamate (8.0 g, 17.92 mmol, 1.0 eq.) in DCM (80 mL) was added iodine (4.6 g, 17.92 mmol, 1.0 eq.) and the resulting suspension was stirred 25 °C for 2 h. After completion, the reaction mixture was diluted with water (200 mL), extracted with DCM (200 mL x 3), organic layers dried over anhydrous Na_2_SO_4_, filtered, and concentrated. The crude was purified by combiflash (eluent 4–5% EtOAc in hexane) to afford *ter*t-butyl (*E*)-(3-iodoallyl)carbamate (**LIV**) (2.0 g, 40% yield). Step-4: To stirred a solution of *tert*-butyl (*E*)-(3-iodoallyl)carbamate (**LIV**) (1.0 g, 3.53 mmol, 1.0 eq.), in DMSO (10 mL) were added sodium cyclopropanesulfinate (0.68 g, 5.3 mmol, 1.5 eq.) and copper iodide (0.34 g, 1.77 mmol, 0.5 eq.). The reaction mixture was stirred at 80 °C for 5 h. After completion, the reaction mixture was poured into water, extracted with EtOAc (2 x 50 mL), organic layer dried over anhydrous Na_2_SO_4_, filtered, and concentrated. The crude was purified by combiflash (eluent 10% EtOAc in hexane) to furnish *tert*-butyl (*E*)-(3-(cyclopropylsulfonyl)allyl)carbamate as white solid (0.65 g, 71% yield). MS (ESI) *m/z*: 279.2 [M+H]^+^. Step-5: Boc deprotection of *tert-*butyl (*E*)-(3-(cyclopropylsulfonyl) allyl) carbamate (0.5 g, 1.9 mmol, 1.0 eq.) following General Procedure-V afforded (*E*)-3-(cyclopentylsulfonyl) prop-2-en-1-amine (**LV**) (0.3 g, 97% yield). Step-6: Amide coupling of 5-(2-ethoxyphenyl)-1-(tetrahydro-2*H*-pyran-2-yl)-1*H*-pyrazole-3-carboxylic acid (**VII**) and (*E*)-3-(cyclopentylsulfonyl) prop-2-en-1-amine (**LV**) by General Procedure-II afforded (*E*)-*N*-(3-(cyclopropylsulfonyl)allyl)-5-(2-ethoxyphenyl)-1-(tetrahydro-2*H*-pyran-2-yl)-1*H*-pyrazole-3-carboxamide as a white solid (4.25 g, 59% yield). Subsequent THP deprotection following General Procedure-V furnished (*E*)-*N*-(3-(cyclopropylsulfonyl)allyl)-5-(2-ethoxyphenyl)-1*H*-pyrazole-3-carboxamide (**25a**) as a white solid (0.07 g, 44% yield, m.p. 105 °C). ^1^H NMR (400 MHz, DMSO-*d*_6_): *δ* 11.76 (s, 1H), 7.75 (d, *J* = 7.2 Hz, 1H), 7.57 (s, 1H), 7.40 (t, *J* = 7.2 Hz, 1H), 7.16 – 7.06 (m, 3H), 6.89 – 6.84 (m, 1H), 6.58 (d, *J* = 15.2 Hz, 1H), 4.30 – 4.20 (m, 4H), 2.49 – 2.45 (m, 1H), 1.49 (t, *J* = 7.2 Hz, 3H), 1.09 – 1.02 (m, 4H). ^13^C NMR (101 MHz, DMSO-*d*_6_): *δ* 162.1, 155.3, 143.7, 130.1, 129.1, 128.3, 121.0, 117.3, 112.8, 108.7, 64.3, 38.8, 30.4, 13.9, 4.4. HRMS (ESI) *m/z*: [M+H]^+^ calculated for C_18_H_22_N_3_O_4_S: 376.1331, found 376.1340. HPLC purity (254 nm) >99%.

**(*E*)-5-(2-Ethoxyphenyl)-*N*-(3-(pyridin-2-ylsulfonyl)allyl)-1*H*-pyrazole-3-carboxamide (25g).** Step-1: To a solution *tert*-butyl (*E*)-(3-iodoallyl)carbamate (**LIV**) (1.0 g, 3.53 mmol, 1.0 eq.) in toluene (10 mL) was added pyridine-2-thiol (0.39 g, 3.53 mmol, 1.0 eq.) and degassed for 10 min. Then CuI (0.13 g, 0.7 mmol, 0.2 eq.) and DBU (1.07 g, 7.06 mmol, 2.0 eq.) were added and the reaction was stirred 110 °C for 2 h. After completion, the reaction mixture was diluted with water (100 mL), extracted with EtOAc (100 mL x 3), dried over anhydrous Na_2_SO_4_, filtered, and concentrated. The residue was purified by combiflash (eluent 15% EtOAc in hexane) to afford *tert*-butyl (*E*)-(3-(pyridin-2-ylthio)allyl)carbamate (0.4 g, 43% yield). MS (ESI) *m/z*: 267.1 [M+H]^+^. Step-2: To a stirred solution of *tert*-butyl (*E*)-(3-(pyridin-2-ylthio)allyl)carbamate (0.4 g, 1.5 mmol, 1.0 eq.) in DCM (10 mL) was added m-CPBA (1.03 g, 6.00 mmol, 4.0 eq.) at 0 °C and the reaction was stirred at 25 °C for 12 h. After completion, the reaction was quenched with NaHCO_3_ (100 mL), extracted with DCM (100 mL x 3), organic layer dried over anhydrous Na_2_SO_4_, filtered, and concentrated. The residue was purified by combiflash (eluent 60% EtOAc in hexane) to furnish *tert*-butyl (*E*)-(3-(pyridin-2-ylsulfonyl)allyl)carbamate (0.3 g, 67% yield). MS (ESI) *m/z*: 299.3 [M+H]^+^. Step-3: Boc deprotection of *tert*-butyl (*E*)-(3-(pyridin-2-ylsulfonyl)allyl)carbamate (0.3 g, 1.0 mmol, 1.0 eq.) following General Procedure-V afforded (*E*)-3-(pyridin-2-ylsulfonyl)prop-2-en-1-amine (**LVI**) (0.2 g, quantitative yield). MS (ESI) *m/z*: 199.3 [M+H]^+^. Step-4: A two-step reaction sequence of a) amide coupling following General Procedure-II with acid **VII** and amine **LVI**, b) THP deprotection following General Procedure-V furnished (*E*)-5-(2-ethoxyphenyl)-*N*-(3-(pyridin-2-ylsulfonyl)allyl)-1*H*-pyrazole-3-carboxamide (**25g**) as a white solid (0.025 g, 12% yield over two steps, m.p. 161 °C). ^1^H NMR (400 MHz, DMSO-*d*_6_): *δ* 8.79 (d, *J* = 3.2 Hz, 1H), 8.68 (d, *J* = 5.2 Hz, 1H), 8.18 – 8.14 (m, 1H), 8.09 (d, *J* = 7.2 Hz, 1H), 7.76 – 7.74 (m, 2H), 7.38 (t, *J* = 8.4 Hz, 1H), 7.16 – 7.02 (m, 4H), 6.81 (d, *J* = 16.0 Hz, 1H), 4.02 – 4.15 (m, 4H), 1.43 (t, *J* = 8.0 Hz, 3H). ^13^C NMR (101 MHz, DMSO-*d*_6_): *δ* 161.9, 158.0, 155.5, 150.9, 147.4, 145.2, 141.9, 139.6, 130.1, 128.4, 128.3, 128.1, 122.2, 121.0, 118.7, 113.3, 106.1, 64.2, 15.0. MS (ESI) *m/z*: [M+H]^+^ calculated for C_20_H_21_N_4_O_4_S: 413.13, found 413.2. HPLC purity (254 nm) >98%.

**(*E*)-3-(2-Ethoxyphenyl)-*N*-(3-(methylsulfinyl)allyl)-1*H*-pyrazole-5-carboxamide (26).** Step-1: To a stirred solution of *ter*t-butyl (*E*)-(3-iodoallyl)carbamate (**LIV**) (8.0 g, 28.3 mmol, 1.0 eq.) in toluene (160 mL) were added CH_3_SNa (2.96 g, 42.4 mmol, 1.5 eq.), CuI (2.7 g, 14.13 mmol, 0.5 eq.), and DBU (8.65 g, 56.54 mmol, 2.0 eq.). Reaction mixture was stirred at 80 °C for 4 h. After completion, the reaction mixture was diluted with water (200 mL), extracted with EtoAc (200 mL x 2), dried over anhydrous Na_2_SO_4_, filtered, and concentrated. The crude was purified by combiflash (eluent 30% EtOAc in hexane) to obtain *tert*-butyl (*E*)-(3-(methylthio)allyl)carbamate (3.8 g, 66% yield). Step-2: m-CPBA mediated oxidation (0.93 g, 5.42 mmol, 1.0 eq.) of (*E*)-(3-(methylthio)allyl)carbamate (1.1 g, 5.42 mmol, 1.0 eq.) in DCM (20 mL) furnished *tert*-butyl (*E*)-(3-(methylsulfinyl)allyl)carbamate (0.8 g, 67% yield). Step-3: Boc deprotection of *tert*-butyl (*E*)-(3-(methylsulfinyl)allyl)carbamate (0.4 g, 1.83 mmol, 1.0 eq.) following General Procedure-V afforded (*E*)-3-(methylsulfinyl)prop-2-en-1-amine (**LVII**) (0.2 g, 92% yield). Step-4: A two-step reaction sequence of a) amide coupling of 5-(2-ethoxyphenyl)-1-(tetrahydro-2*H*-pyran-2-yl)-1*H*-pyrazole-3-carboxylic acid (**VII**) and (*E*)-3-(methylsulfinyl)prop-2-en-1-amine (**LVII**) by General Procedure-II, b) THP deprotection following General Procedure-V afforded (*E*)-3-(2-ethoxyphenyl)-*N*-(3-(methylsulfinyl)allyl)-1*H*-pyrazole-5-carboxamide (**26**) as a white solid (0.03 g, 28% yield over two steps). ^1^H NMR (400 MHz, DMSO-*d*_6_): *δ* 12.94 (br, 1H) 8.27 (br, 1H), 7.73 (d, *J* = 6.4 Hz, 1H), 7.36 (t, *J* = 8.4 Hz, 1H), 7.14 (d, *J* = 8.4 Hz, 1H), 7.08 – 7.00 (m, 2H), 6.68 (d, *J* = 14.2 Hz, 1H), 6.39 – 6.33 (m, 1H), 4.21 (q, *J* = 6.93 Hz, 2H), 4.10 – 4.08 (m, 2H), 2.59 (s, 3H), 1.42 (t, *J* = 3.6 Hz, 3H). ^13^C NMR (101 MHz, DMSO-*d*_6_): *δ* 154.9, 137.5, 136.7, 135.3, 134.1, 129.6, 127.7, 120.5, 112.8, 105.4, 63.6, 14.5. HRMS (ESI) *m/z*: [M+H]^+^ calculated for C_16_H_20_N_3_O_3_S: 334.1225, found 334.1219. HPLC purity (254 nm) >99%.

**(*E*)-3-(2-Ethoxyphenyl)-*N*-(3-(*S*-methylsulfonimidoyl)allyl)isoxazole-5-carboxamide (27).** Step-1: Amide coupling of 3-(2-ethoxyphenyl)isoxazole-5-carboxylic acid (**XI**) and (*E*)-3-(methylsulfinyl)prop-2-en-1-amine (**LVII**) following General Procedure-I afforded (*E*)-3-(2-ethoxyphenyl)-*N*-(3-(methylsulfinyl)allyl)isoxazole-5-carboxamide (0.09 g, 31% yield). MS (ESI) *m/z*: 335.4 [M+H]^+^. Step-2: To a stirred solution of (*E*)-3-(2-ethoxyphenyl)-*N*-(3-(methylsulfinyl)allyl)isoxazole-5-carboxamide (0.07 g, 0.21 mmol, 1.0 eq.) in MeOH (2 mL) were added ammonium carbamate (0.08 mg, 1.05 mmol, 5.0 eq.) and iodobenzene diacetate (0.2 mg, 0.63 mmol, 3.0 eq.). The reaction was stirred at 25 °C for 1 h. After completion, the reaction mixture was diluted water (5 mL), extracted with EtOAc (5 mL x 2), organic layer dried over anhydrous Na_2_SO_4_, filtered, and concentrated. The residue was purified by preparative HPLC to afford (*E*)-3-(2-ethoxyphenyl)-*N*-(3-(*S*-methylsulfonimidoyl)allyl)isoxazole-5-carboxamide (**27**) as a white solid (0.015 g, 20% yield, m.p. 123**–**124 °C). ^1^H NMR (400 MHz, DMSO-*d*_6_): *δ* 9.38 (t, *J*_1_= 5.6, *J*_2_= 6, 1H), 7.82 – 7.79 (m, 1H), 7.53 – 7.47 (m, 2H), 7.23 – 7.21 (d, *J* = 8 Hz, 1H), 7.10 – 7.06 (m, 1H), 6.79 – 6.66 (m, 2H), 4.21 – 4.11 (m, 4H), 3.87 (s, 1H), 2.92 (s, 3H), 1.40 (t, *J*_1_= 6.8, *J*_2_= 6.8 Hz, 3H). ^13^C NMR (101 MHz, DMSO-*d*_6_): *δ* 163.2, 160.7, 156.7, 156.2, 139.7, 134.5, 132.6, 129.4, 121.2, 116.8, 113.6, 107.9, 64.4, 44.6, 14.9. HRMS (ESI) *m/z*: [M+H]^+^ calculated for C_16_H_20_N_3_O_4_S: 350.1175, found 350.1169. HPLC purity (254 nm) >99%.

**(*E*)-5-(2-Ethoxyphenyl)-*N*-(2-methyl-4-(methylsulfonyl)but-3-en-2-yl)-1*H*-pyrazole-3-carboxamide (24b).** Step-1: *tert*-butyl (*E*)-(4-iodo-2-methylbut-3-en-2-yl)carbamate (**LVIII**) was synthesized from 2-methylbut-3-yn-2-amine (**LIIIb**) over three steps (5.0 g, 28% yield over three steps), following a similar route as used for the synthesis of intermediate (**LIV**). Step-2: To a stirred solution of *tert*-butyl (*E*)-(4-iodo-2-methylbut-3-en-2-yl)carbamate (**LVIII**) (5.0 g, 16.1 mmol, 1.0 eq.) in DMSO (50 mL) were added sodium methane sulfinate (2.45 g, 24.1 mmol, 1.5 eq.) and CuI (4.5 g, 24.1 mmol, 1.5 eq.). The reaction was stirred at 80 °C for 4 h. After completion, the reaction was diluted with water (100 mL), extracted in EtOAc (2 x 100 mL), washed with brine, dried over anhydrous Na_2_SO_4_, filtered, and concentrated. The residue was purified by combiflash (eluent 70% EtOAc in hexane) to give *tert-*butyl (*E*)-(2-methyl-4-(methylsulfonyl)but-3-en-2-yl)carbamate as a white solid (1.8 g, 43% yield). Step-3: Boc deprotection of *tert-*butyl (*E*)-(2-methyl-4-(methylsulfonyl)but-3-en-2-yl)carbamate (1.5 g, 5.7 mmol, 1.0 eq.) following General Procedure-V afforded (*E*)-2-methyl-4-(methylsulfonyl)but-3-en-2-amine (**LIX**) as a white solid (0.9 g, 91% yield). Step-4: A two-step reaction sequence of a) amide coupling of 5-(2-ethoxyphenyl)-1-(tetrahydro-2*H*-pyran-2-yl)-1*H*-pyrazole-3-carboxylic acid VII and (*E*)-2-methyl-4-(methylsulfonyl)but-3-en-2-amine (**LIX**) by General Procedure-I, b) THP deprotection following General Procedure-V afforded (*E*)-5-(2-ethoxyphenyl)-*N*-(2-methyl-4-(methylsulfonyl)but-3-en-2-yl)-1*H*-pyrazole-3-carboxamide (**24b**) as a white solid (0.08 g, 41% yield over two steps, m.p. 128 °C). ^1^H NMR (400 MHz, DMSO-*d*_6_): *δ* 7.85 (brs, 1H), 7.73 (d, *J* = 8.0 Hz, 1H), 7.36 (dd, *J* = 16 Hz, 1H), 7.14 (d, *J* = 8.0 Hz, 1H), 7.08 (s, 1H), 7.04 (t, *J* = 8.0 Hz, 1H), 6.97 (s, 1H), 6.72 (s, 1H), 4.18 (q, *J* = 8.0 Hz, 2H), 2.99 (s, 3H), 1.51 (s, 6H), 1.43 (q, *J* = 8.0 Hz, 3H). ^13^C NMR (101 MHz, DMSO-*d*_6_): *δ* 161.4, 155.5, 151.7, 130.2, 129.8, 128.3, 128.1, 120.0, 118.7, 113.3, 108.4, 106.0, 64.1, 42.8, 26.1, 15.2. HRMS (ESI) *m/z*: [M+H]^+^ calculated for C_18_H_24_N_3_O_4_S: 378.1488, found 378.1483. HPLC purity (254 nm) >99%.

**5-(2-Ethoxyphenyl)-*N*-(2-(methylsulfonyl)allyl)-1*H*-pyrazole-3-carboxamide (24e).** Step-1: To a solution of propargylamine (**LIIIa**) (4.0 g, 25.8 mmol, 1.0 eq.) in THF (40 mL) were added sodium thiomethoxide (1.8 g, 25.8 mmol, 1.0 eq.), Pd(OAc)_2_ (115 mg, 0.52 mmol, 0.02 eq.), and the reaction was stirred at 80 °C for 24 h. After completion, the reaction mixture was diluted with water (200 mL), extracted with EtOAc (200 mL x 2), organic layer dried over anhydrous Na_2_SO_4_, filtered, and concentrated. The residue was purified by combiflash (eluent 25% EtOAc in hexane) to furnish *tert*-butyl (2-(methylthio)allyl)carbamate (1.3 g, 24% yield). MS (ESI) *m/z*: 148.1 [M+H]^+^. Step-2: m-CPBA oxidation (4.4 g, 25.6 mmol, 4.0 eq.) of (*E*)-(3-(methylthio)allyl)carbamate (1.3 g, 6.4 mmol, 1.0 eq.) in DCM (15 mL) furnished *tert*-butyl (2-(methylsulfonyl)allyl)carbamate (0.65 g, 56% yield). Step-3: Boc deprotection of *tert*-butyl (2-(methylsulfonyl)allyl)carbamate (0.65 g, 2.8 mmol, 1.0 eq.) following General Procedure-V afforded 2-(methylsulfonyl)prop-2-en-1-amine (**LXI)** as a white solid (0.35 g, 93% yield). Step-4: A two-step reaction sequence of a) amide coupling of 5-(2-ethoxyphenyl)-1-(tetrahydro-2*H*-pyran-2-yl)-1*H*-pyrazole-3-carboxylic acid (**VII**) and 2-(methylsulfonyl)prop-2-en-1-amine (**LXI**) by General Procedure-I, b) THP deprotection following General Procedure-V afforded 5-(2-ethoxyphenyl)-*N*-(2-(methylsulfonyl)allyl)-1*H*-pyrazole-3-carboxamide (**24e**) as a white solid (0.05 g, 23% yield over two steps, m.p. 152 °C). ^1^H NMR (400 MHz, DMSO-*d*_6_): *δ* 8.73 (s, 1H), 7.75 (d, *J* = 8.0 Hz, 1H), 7.38 – 7.33 (m, 1H), 7.15 – 7.11 (m, 2H), 7.05 – 7.01 (m, 1H), 6.16 (s, 1H), 5.93 (s, 1H), 4.26 (d, *J* = 4.0 Hz, 2H), 4.20 – 4.14 (m, 2H), 3.13 (s, 3H), 1.41 (t, *J* = 6.8 Hz, 3H). ^13^C NMR (101 MHz, DMSO-*d*_6_): *δ* 162.0, 155.5, 148.3, 145.5, 141.9, 130.2, 128.3, 124.5, 121.0, 119.6, 118.6, 113.3, 106.1, 64.2, 41.6, 37.9, 15.0. HRMS (ESI) *m/z*: [M+H]^+^ calculated for C_16_H_20_N_3_O_4_S: 350.1175, found 350.1170. HPLC purity (254 nm) >97%.

### Synthesis of 25b–f

Step-1: To a solution of *N*-Boc propargylamine (obtained by Boc-protection of propargylamine **LIIIa**) (1.0 eq.) in benzene were added AIBN (3.0 eq.) and corresponding alkyl/ aryl thiols (1.0 eq.). The reaction was stirred at 80 °C for 2–5 h. After completion, the reaction mixture was concentrated, and the residue was purified by combiflash to obtain intermediates **LXIIa–e** as solids (30– 79% yields). Step-2: A two-step reaction sequence of a) m-CPBA or oxone-mediated oxidation, and b) Boc deprotection following General Procedure-V afforded intermediates **LXIIIa–e**. <u>Method A</u>: To stirred solution of **LXIIb**, **LXIIc**, or **LXIIf** (1.0 eq.) in DCM, m-CPBA (2.0–4.0 eq.) was added and the reaction was stirred at 25 °C for 16 h. After completion, reaction mixture was filtered through celite pad and washed with DCM. The filtrate was diluted with 0.1N NaOH (100 mL) and extracted with DCM (3 x 100 mL). Combined layers were dried over anhydrous Na_2_SO_4_, filtered, and concentrated. The residue was purified by combiflash to obtain desired products as solids (20–48% yields). Subsequent Boc deprotection following General Procedure-V furnished intermediates **LXIIIb**, **LXIIIc**, and **LXIIIf** respectively. <u>Method B</u>: To a stirred solution of **LXIId** or **LXIIe** (1.0 eq.) in THF:H_2_O was added oxone (10.0 eq.) and reaction stirred at 25 °C for 4–16 h. After completion, the reaction mixture was diluted with water (30 mL), extracted with EtOAc (200 mL x 2), organic layer dried over anhydrous Na_2_SO_4_, filtered, and concentrated. The residue was purified by combiflash to furnish desired products as solids (27–45% yields). Subsequent Boc deprotection following General Procedure-V furnished intermediates **LXIIId** and **LXIIIe** respectively. Step-3: A two-step reaction sequence: a) amide coupling of 5-(2-ethoxyphenyl)-1-(tetrahydro-2*H*-pyran-2-yl)-1*H*-pyrazole-3-carboxylic acid (**VII**) and intermediates **LXIIIa–e** by General Procedure-I, b) THP deprotection following General Procedure-V afforded final compounds **25b–f** as solids.

**(*E*)-5-(2-Ethoxyphenyl)-*N*-(3-(isopropylsulfonyl)allyl)-1*H*-pyrazole-3-carboxamide (25b)**. White solid (61% yield), m.p. 58 °C, ^1^H NMR (400 MHz, DMSO-*d*_6_): *δ* 13.31 (s, 1H), 8.65 (s, 1H), 7.73 (s, 1H), 7.34 – 7.32 (m, 1H), 7.14 – 7.00 (m, 3H), 6.81 (d, *J* = 14.8 Hz, 1H), 6.54 (d, *J* = 15.2 Hz, 1H), 4.17 – 4.12 (m, 4H), 3.20 (t, *J* = 18.4 Hz, 1H), 1.42 (t, *J* = 13.2 Hz, 3H), 1.19 (d, *J* = 6.4 Hz, 6H). ^13^C NMR (101 MHz, DMSO-*d*_6_): *δ* 161.9, 155.5, 146.9, 130.1, 128.2, 126.3, 121.0, 113.2, 106.0, 64.1, 53.3, 15.4, 15.0. HRMS (ESI) *m/z*: [M+H]^+^ calculated for C_18_H_24_N_3_O_4_S: 378.1488, found 378.1493. HPLC purity (254 nm) >99%.

**(*E*)-*N*-(3-(Eyclopentylsulfonyl)allyl)-5-(2-ethoxyphenyl)-1*H*-pyrazole-3-carboxamide (25c).** White solid (2% yield), m.p. 132 °C, ^1^H NMR (400 MHz, DMSO-*d*_6_): *δ* 13.12 (s, 1H), 8.34 (s, 1H), 7.69 (s, 1H), 7.36 – 7.32 (m, 1H), 7.14 – 7.12 (d, J = 8.4 Hz, 1H), 7.05 – 7.01 (t, J = 7.2 Hz, 2H), 6.84 – 6.78 (m, 1H), 6.59 – 6.55 (d, J = 15.2 Hz, 1H), 4.20 – 4.13 (m, 4H), 3.51 – 3.45 (m, 1H), 1.93 – 1.70 (m, 4H), 1.70 – 1.55 (m, 4H), 1.42 – 1.39 (m, 3H). ^13^C NMR (101 MHz, DMSO-*d*_6_): *δ* 162.5, 155.4, 147.4, 146.1, 140.5, 130.3, 128.3, 127.7, 121.0, 117.9, 113.3, 105.9, 64.2, 61.8, 26.9, 25.9, 14.9. HRMS (ESI) *m/z*: [M+H]^+^ calculated for C_20_H_26_N_3_O_4_S: 404.1644, found 404.1639. HPLC purity (254 nm) >99%.

**(*E*)-5-(2-Ethoxyphenyl)-*N*-(3-(phenylsulfonyl)allyl)-1*H*-pyrazole-3-carboxamide (25d).** White solid (3% yield), ^1^H NMR (400 MHz, DMSO-*d*_6_): *δ* 13.32 (s, 1H), 8.60 (s, 1H), 7.89 – 7.85 (m, 2H), 7.76 – 7.69 (m, 2H), 7.67 – 7.61 (m, 2H), 7.34 (ddd, *J* = 8.8, 7.4, 1.7 Hz, 1H), 7.16 – 6.92 (m, 4H), 6.72 (dt, *J* = 15.1, 1.8 Hz, 1H), 4.19 – 4.08 (m, 4H), 1.39 (t, *J* = 6.9 Hz, 3H). ^13^C NMR (126 MHz, DMSO-*d*_6_): *δ* 161.4, 155.0, 144.6, 140.3, 133.7, 129.9, 129.7, 129.6, 127.8, 127.2, 120.5, 112.8, 105.6, 63.7, 14.5. HRMS (ESI) *m/z*: [M+H]^+^ calculated for C_21_H_22_N_3_O_4_S: 412.1331, found 412.1328. HPLC purity (254 nm) >99%.

**(*E*)-3-(2-Ethoxyphenyl)-*N*-(3-((4-methoxyphenyl)sulfonyl)allyl)-1*H*-pyrazole-5-carboxamide (25e).** White solid (30% yield), m.p. 72 °C, ^1^H NMR (400 MHz, DMSO-*d*_6_): *δ* 13.01 (s, 1H), 8.28 (s, 1H), 7.77 (d, *J* = 4 Hz, 1H), 7.67 (s, 1H), 7.32 (s, 1H), 7.13 (d, *J* = 3.8 Hz, 3H), 7.02 (t, *J* = 14 Hz, 2H), 6.87 – 6.81 (m, 1H), 6.65 (d, *J* = 14.4 Hz, 1H), 4.19 – 4.09 (m, 4H), 3.88 (d, *J* = 14.8 Hz, 3H), 1.40 (t, *J* = 13.2 Hz, 3H). ^13^C NMR (101 MHz, DMSO-*d*_6_): *δ* 163.6, 162.4, 159.0, 155.4, 147.0, 143.6, 140.5, 132.2, 131.1, 130.3, 130.1, 128.2, 121.0, 117.9, 115.3, 113.2, 107.6, 105.9, 64.1, 56.3, 14.9. HRMS (ESI) *m/z*: [M+H]^+^ calculated for C_22_H_24_N_3_O_5_S: 442.1437, found 442.1446. HPLC purity (254 nm) >99%.

**(*E*)-5-(2-Ethoxyphenyl)-*N*-(3-((2-fluorophenyl)sulfonyl)allyl)-1*H*-pyrazole-3-carboxamide (25f).** White solid (30% yield), m.p. 85 °C, ^1^H NMR (500 MHz, MeOD): *δ* 7.93 (td, *J* = 7.5, 1.8 Hz, 1H), 7.76 – 7.69 (m, 1H), 7.64 (dd, *J* = 7.7, 1.7 Hz, 1H), 7.40 (td, *J* = 7.7, 1.0 Hz, 1H), 7.37 – 7.29 (m, 2H), 7.16 – 7.10 (m, 3H), 7.03 (td, *J* = 7.5, 1.0 Hz, 1H), 6.75 (dq, *J* = 15.0, 1.6 Hz, 1H), 4.26 (dd, *J* = 4.5, 1.9 Hz, 2H), 4.21 (q, *J* = 7.0 Hz, 2H), 1.48 (t, *J* = 7.0 Hz, 3H). ^13^C NMR (101 MHz, DMSO-*d*_6_): *δ* 161.5, 159.9, 157.4, 154.9, 146.7, 136.9, 136.8, 129.6, 129.3, 129.1, 127.9, 125.4, 120.5, 118.2, 117.6, 112.8, 105.5, 63.7, 14.5. HRMS (ESI) *m/z*: [M+H]^+^ calculated for C_21_H_21_FN_3_O_4_S: 430.1237, found 430.1234. HPLC purity (254 nm) >99%.

**(*E*)-3-(2-Ethoxyphenyl)-*N*-methyl-*N*-(3-(methylsulfonyl)allyl)isoxazole-5-carboxamide (23a).** Step-1: To a stirred solution of *tert*-butyl (2-hydroxyethyl)(methyl)carbamate (**LXIV**) (5.0 g, 28.6 mmol, 1.0 eq.) in DCM (50 mL) was added Dess-Martin periodinane (13.3 g, 31.42 mmol, 1.1 eq.) and the reaction was stirred at 25 °C for 1 h. After completion, the reaction mixture was diluted with NaHCO_3_ solution (200 mL), extracted with *tert*-Butyl methyl ether (2 x 400 mL), combined organic layers dried over anhydrous Na_2_SO_4_, filtered, and concentrated to afford *tert-*butyl methyl(2-oxoethyl)carbamate (**LXV**) as a colorless liquid (5.0 g, quantitative). Step-2: To a stirred solution of diethyl ((methylsulfonyl) methyl) phosphonate (4.7 g, 20.2 mmol, 0.7 eq.) in ACN (150 mL), were added LiCl (1.3 g, 31.8 mmol, 1.1 eq.), DIPEA (4.9 mL, 28.9 mmol, 1.0 eq.) and stirred at 0 °C for 10 min. Then *tert*-butyl methyl(2-oxoethyl) carbamate (**LXV**) (5.0 g, 28.9 mmol, 1.0 eq.) was added and the reaction was stirred at 25 °C for 16 h. After completion, the reaction mixture was poured into water (150 mL), extracted with EtOAc (2 x 300 mL), combined organic layers dried over anhydrous Na_2_SO_4_, filtered, and concentrated. The residue was purified by combiflash (eluent 30% EtOAc in hexane) to afford *tert*-butyl (*E*)-methyl(3-(methylsulfonyl)allyl) carbamate as a yellow liquid (2.4 g, 33% yield). Subsequent Boc deprotection following General Procedure-V furnished (*E*)-*N*-methyl-3-(methylsulfonyl) prop-2-en-1-amine (**LXVI**) as a white solid (1.5 g). MS (ESI) *m/z*: 150.1 [M+H]^+^. Step-3: To a stirred solution of 3-(2-ethoxyphenyl)isoxazole-5-carboxylic acid (**XI**) (0.1 g, 0.42 mmol, 1.0 eq.) and (*E*)-N-methyl-3-(methylsulfonyl) prop-2-en-1-amine (**LXVI**) (0.06 g, 0.42 mmol, 1.0 eq.) in THF (1.0 mL) were added triethylamine (0.18 mL, 1.25 mmol, 3.0 eq.), T3P (50% in EtOAc, 0.4 mL, 0.64 mmol, 1.5 eq.) and the reaction was stirred at 25 °C for 15 min. After completion, the reaction was diluted with water (50 mL), extracted with EtOAc (50 mL x 2), organic layers dried over anhydrous Na_2_SO_4_, filtered, and concentrated. The residue was purified by preparative HPLC to afford (*E*)-3-(2-ethoxyphenyl)-*N*-Methyl-*N*-(3-(methylsulfonyl)allyl) isoxazole-5-carboxamide (**23a**) as a white solid (0.025 g, 16% yield, m.p. 120 °C). ^1^H NMR (400 MHz, DMSO-*d*_6_): *δ* 7.91 (t, *J* = 7.2 Hz, 1H), 7.51 (t, *J* = 6.4 Hz, 1H), 7.25 – 7.22 (m, 1H), 7.14 – 7.07 (m, 1H), 7.00 (s, 1H), 6.88 – 6.85 (m, 1H), 6.78 – 6.72 (m, 1H), 4.38 – 4.34 (m, 2H), 4.24 – 4.21 (m, 2H), 3.17 (s, 2H), 3.04 – 3.02 (m, 4H), 1.46 – 1.42 (m, 3H). ^13^C NMR (101 MHz, DMSO-*d*_6_): *δ* 166.3, 166.2, 161.3, 161.2, 159.6, 159.4, 155.8, 141.5, 140.6, 132.8, 132.2, 131.8, 127.6, 127.5, 121.3, 115.2, 115.1, 113.5, 104.2, 103.8, 64.6, 50.9, 48.0, 34.2, 14.9. HRMS (ESI) *m/z*: [M+H]^+^ calculated for C_17_H_21_N_2_O_5_S: 365.1171, found 365.1166. HPLC purity (254 nm) >99%.

**(*E*)-5-(2-Ethoxyphenyl)-*N*-(4-(methylsulfonyl)but-3-en-2-yl)-1*H*-pyrazole-3-carboxamide (24a).** (*E*)-4-(methylsulfonyl)but-3-en-2-amine (**LXIX**) was synthesized from *tert*-butyl (1-hydroxypropan-2-yl)carbamate (**LXVII**) over a three-step sequence following the same protocol as described for the synthesis of **LXVI** (0.55 g, 12% yield over three steps). MS (ESI) *m/z*: 150.2 [M+H]^+^. Amide coupling of 5-(2-ethoxyphenyl)-1-((2-(trimethylsilyl)ethoxy)methyl)-1*H*-pyrazole-3-carboxylic acid (**IX**) and (*E*)-4-(methylsulfonyl)but-3-en-2-amine (**LXIX**) by General Procedure-II afforded (*E*)-5-(2-ethoxyphenyl)-*N*-(4-(methylsulfonyl)but-3-en-2-yl)-1-((2-(trimethylsilyl)ethoxy)methyl)-1*H*-pyrazole-3-carboxamide (**LXX**) (0.25 g, 37% yield). SEM deprotection using 4M HCl in dioxane furnished (*E*)-5-(2-ethoxyphenyl)-*N*-(4-(methylsulfonyl)but-3-en-2-yl)-1*H*-pyrazole-3-carboxamide (**24a**) as a white solid (0.17 g, 70% yield, m.p. 160 °C). Chiral SFC separation of intermediate **LXX** afforded (*R)-(E*)-5-(2-ethoxyphenyl)-*N*-(4-(methylsulfonyl)but-3-en-2-yl)-1-((2-(trimethylsilyl)ethoxy)methyl)-1*H*-pyrazole-3-carboxamide (*t*_ret_ 4.32 min, 0.04 g, 16% yield) and (*S)-(E*)-5-(2-ethoxyphenyl)-*N*-(4-(methylsulfonyl)but-3-en-2-yl)-1-((2-(trimethylsilyl)ethoxy)methyl)-1*H*-pyrazole-3-carboxamide (*t*_ret_ 4.62 min, 0.038 g, 15% yield). SEM deprotection of each enantiomer yielded (*R*)-**24a** (0.014 g, 48% yield,) and (*S*)-**24a** (0.013 g, 47% yield,). **24a**: ^1^H NMR (500 MHz, DMSO-*d*_6_): *δ* 8.44 (d, *J* = 8.3 Hz, 1H), 7.77 – 7.69 (m, 1H), 7.35 (ddd, *J* = 8.8, 7.3, 1.7 Hz, 1H), 7.17 – 7.07 (m, 2H), 7.03 (td, *J* = 7.5, 1.0 Hz, 1H), 6.84 (dd, *J* = 15.3, 5.0 Hz, 1H), 6.74 (dd, *J* = 15.3, 1.4 Hz, 1H), 4.87 – 4.79 (m, 1H), 4.16 (q, *J* = 6.9 Hz, 2H), 1.40 (t, *J* = 6.9 Hz, 3H), 1.34 (d, *J* = 7.0 Hz, 3H). ^13^C NMR (126 MHz, DMSO-*d*_6_): *δ* 155.0, 147.0, 147.0, 129.7, 129.6, 127.9, 120.5, 112.8, 105.7, 63.7, 44.5, 42.2, 19.4, 14.6. HRMS (ESI) *m/z*: [M+H]^+^ calculated for C_17_H_22_N_3_O_4_S: 364.1331, found 364.1324; HPLC purity (254 nm) >99%.

> (*R*)-**24a**: Chiral SFC: *t*_ret_ 5.17 min. [α]_D_^20^ −11.9 deg cm^3^ g^−1^ dm^−1^ (*c* 0.005 g cm^−3^, MeOH).

> (*S*)-**24a**: Chiral SFC: *t*_ret_ 6.35 min. [α]_D_^20^ +12.2 deg cm^3^ g^−1^ dm^−1^ (*c* 0.005 g cm^−3^, MeOH).

**(*E*)-5-(2-Ethoxyphenyl)-*N*-(4-(methylsulfonyl)-1-phenylbut-3-en-2-yl)-1*H*-pyrazole-3-carboxamide (24c).** Step-1: Amide coupling of Boc-*L*-phenylalanine (**LXXI**) (4.0 g, 1.5 mmol, 1.0 eq.) and *N*,*O*-dimethyl hydroxylamine hydrochloride (2.2 g, 22.6 mmol, 1.0 eq.) following General Procedure-II afforded *tert*-butyl (1-(methoxy(methyl)amino)-1-oxo-3-phenylpropan-2-yl)carbamate as a white solid (4.0 g, 86% yield). MS (ESI) *m/z*: 308.4 [M+H]^+^. This was followed by LAH reduction (1.5 eq.) in THF at 0 °C for 1 h to obtain *tert*-butyl (1-oxo-3-phenylpropan-2-yl)carbamate (**LXXII**) as a colorless liquid (1.1 g, 91% yield). Step-2: To a solution of diethyl ((methylsulfonyl)methyl)phosphonate (3.6 g, 15.7 mmol, 1.0 eq.) in THF (40 mL) at –78 °C under N_2_ atmosphere was added *n*-BuLi (13.6 mL, 31.3 mmol, 2.0 eq.) and stirred for 20 min. Then *tert*-butyl (1-oxo-3-phenylpropan-2-yl)carbamate (**LXXII**) (3.9 g, 15.7 mmol, 1.0 eq.) was added and the reaction was stirred at –78 °C for 2 h. After completion, the reaction was quenched with NH_4_Cl solution (30 mL), extracted with EtOAc (100 mL x 3), combined layers washed with water (100 mL), dried over anhydrous Na_2_SO_4_, filtered, and concentrated. The residue was purified by combiflash (eluent 30–40% EtOAc in hexane) to afford *tert*-butyl (*E*)-(4-(methylsulfonyl)-1-phenylbut-3-en-2-yl)carbamate as a yellow solid (1.0 g, 20% yield). MS (ESI) *m/z*: 325.4 [M+H]^+^. Subsequent Boc deprotection following General Procedure-V furnished (*E*)-4-(methylsulfonyl)-1-phenylbut-3-en-2-amine (**LXXIII**) as a white solid (0.46 g, 90% yield). MS (ESI) *m/z*: 225.3 [M+H]^+^. Step-3: A two-step reaction sequence: a) amide coupling of 5-(2-ethoxyphenyl)-1-(tetrahydro-2*H*-pyran-2-yl)-1*H*-pyrazole-3-carboxylic acid (**VII**) and (*E*)-4-(methylsulfonyl)-1-phenylbut-3-en-2-amine (**LXXIII**) by General Procedure-I, b) THP deprotection following General Procedure-V afforded (*E*)-5-(2-ethoxyphenyl)-*N*-(4-(methylsulfonyl)-1-phenylbut-3-en-2-yl)-1*H*-pyrazole-3-carboxamide (**24c**) as a white solid (0.005 g, 14% yield). ^1^H NMR (500 MHz, MeOD): *δ* 7.63 (dd, *J* = 7.7, 1.7 Hz, 1H), 7.35 (ddd, *J* = 8.4, 7.4, 1.7 Hz, 1H), 7.32 – 7.27 (m, 4H), 7.24 – 7.19 (m, 1H), 7.12 (dd, *J* = 8.4, 1.0 Hz, 1H), 7.09 (s, 1H), 7.02 (td, *J* = 7.5, 1.1 Hz, 1H), 6.98 (dd, *J* = 15.2, 5.3 Hz, 1H), 6.67 (dd, *J* = 15.2, 1.6 Hz, 1H), 5.10 (dddd, *J* = 8.3, 6.9, 5.3, 1.6 Hz, 1H), 4.22 (q, *J* = 7.0 Hz, 2H), 3.13 – 3.04 (m, 2H), 2.93 (s, 3H), 1.48 (t, *J* = 7.0 Hz, 3H). ^13^C NMR (126 MHz, MeOD): *δ* 154.9, 147.4, 137.9, 137.3, 133.8, 128.9, 122.1, 121.6, 121.0, 120.1, 119.7, 118.4, 112.5, 109.6, 104.3, 96.4, 55.9, 42.9, 33.3, 31.2, 5.6. HRMS (ESI) *m/z*: [M+H]^+^ calculated for C_23_H_26_N_3_O_4_S: 440.1644, found 440.1641. HPLC purity (254 nm) >99%.

**(*E*)-5-(2-Ethoxyphenyl)-*N*-(2-methyl-3-(methylsulfonyl)allyl)-1*H*-pyrazole-3-carboxamide (24d).** Step-1: To a stirred a solution of potassium phthalimide (**LXXIV**) (40.0 g, 216.2 mmol, 1.0 eq.) in ACN was added 1-bromopropan-2-one (26.0 g, 281.0 mmol, 1.0 eq.) and stirred at 60 °C for 16 h. After completion, the reaction was poured into water, extracted with EtOAc, washed with brine, dried over anhydrous Na_2_SO_4_, filtered, and concentrated. The crude was purified by combiflash (eluent 30-40% EtOAc in hexane) to afford 2-(2-oxopropyl)isoindoline-1,3-dione (25% yield). Step-2: To a stirred a solution of diethyl ((methylsulfonyl)methyl)phosphonate (2.2 g, 9.84 mmol, 0.8 eq.) in THF was added LiHMDS (24.6 mL, 2.46 mmol, 2.0 eq.) at 0 °C and stirred for 30 min at 25 °C. Then 2-(2-oxopropyl)isoindoline-1,3-dione (obtained from Step-1) (2.5 g, 12.3 mmol, 1.0 eq.) was added and the reaction was stirred at 25 °C for 2 h. After completion, the reaction was diluted with EtOAc, filtered through celite bad, and the filtrate was concentrated. The crude was purified by combiflash (eluent 70–80% EtOAc in hexane) to afford (*Z*)-2-(2-methyl-3-(methylsulfonyl)allyl)isoindoline-1,3-dione (**LXXV**) (11% yield). MS (ESI) *m/z*: 280.2 [M+H]^+^. Step-3: To a solution of intermediate (**LXXV**) (2.0 g, 7.1 mmol, 1.0 eq.) in EtOH was added hydrazine (0.36 g, 7.1 mmol, 1.0 eq.) and the reaction was stirred at 80 °C for 2 h. After completion, the reaction was concentrated to afford (*Z*)-2-methyl-3-(methylsulfonyl)prop-2-en-1-amine (**LXXVI**) (2.0 g crude). Step-4: Owing to difficulties in purification and isolation of (**LXXVI**) from Step-3, it was Boc-protected using (Boc)_2_O (1.5 eq.) and TEA (3.0 eq.) in DCM to afford *tert*-butyl (*Z*)-(2-methyl-3 (methylsulfonyl)allyl)carbamate (**LXXVII**) (0.6 g, 18% yield). Step-5: Boc deprotection of intermediate (**LXXVII**) (0.2 g, 0.80 mmol, 1.0 eq.) following General Procedure-V furnished (*Z*)-2-methyl-3-(methylsulfonyl)prop-2-en-1-amine (**LXXVI**) (0.15 g, 90% yield). Step-6: A two-step reaction sequence: a) amide coupling of 5-(2-ethoxyphenyl)-1-(tetrahydro-2*H*-pyran-2-yl)-1*H*-pyrazole-3-carboxylic acid (**VII**) and (*Z*)-2-methyl-3-(methylsulfonyl)prop-2-en-1-amine (**LXXVI**) by General Procedure-II, b) THP deprotection following General Procedure-V afforded (*E*)-5-(2-ethoxyphenyl)-*N*-(2-methyl-3-(methylsulfonyl)allyl)-1*H*-pyrazole-3-carboxamide (**24d**) as a mixture of regioisomers (allyl:vinyl = 1:3) (0.011 g, 3% yield over two steps, m.p. 152 °C). ^1^H NMR (400 MHz, DMSO-*d*_6_): *δ* 13.34 (s, 1H, NH), 8.60 – 8.49 (m, 1.3H), 7.73 (s, 1.4H), 7.33 (s, 1.5H), 7.22 – 7.02 (m, 4.4H), 6.24 (s, 0.48H, allyl <u>(minor)</u>), 5.27 (d, *J* = 7.6 Hz, 1.6H, vinyl <u>(major)</u>), 4.15 (d, *J* = 6.4 Hz, 2.9H), 3.99 (s, 4.2H), 3.29 (s, 1H), 3.01 (s, 4.3H), 2.09 (s, 1.3H), 1.04 (s, 4.5H). ^13^C NMR (101 MHz, DMSO-*d*_6_): *δ* 196.8, 155.5, 154.5, 136.1, 130.1, 128.2, 128.1, 124.7, 118.7, 113.2, 105.9, 64.1, 58.9, 45.3, 43.9, 42.9, 38.8, 15.9, 15.0. HRMS (ESI) *m/z*: [M+H]^+^ calculated for C_17_H_22_N_3_O_4_S: 364.1331, found 364.1330. HPLC purity (254 nm) >99%.

### Synthesis of 23b–f

Step-1: To a stirred solution of 3-(methylsulfonyl)prop-1-ene (**LXXVIII**) (0.5 g, 4.16 mmol, 1.0 eq.) in chloroform, Br_2_ (2.6 g, 16.64 mmol, 4.0 eq.) was added at 0 °C and the reaction was stirred at 25 °C for 2 h. After completion, the reaction was quenched with Na_2_S_2_O_3_ solution, extracted with EtOAc (2 x 200 mL), washed with brine, dried over anhydrous Na_2_SO_4_, filtered, and concentrated. The crude was purified by combiflash (eluent 50% EtOAc in hexane) to afford 1,2-dibromo-3-(methylsulfonyl) propane (**LXXIX**) as a white solid (1.0 g, 86% yield). Step-2: To a stirred solution of 1,2-dibromo-3-(methylsulfonyl) propane (**LXXIX**) (1.0 g, 3.58 mmol, 1.0 eq.) in acetone was added Na_2_CO_3_ (1.13 g, 10.7 mmol, 3 eq.) and the reaction stirred at 60 °C for 2 h. After completion, the reaction mixture was poured into water (50 mL), extracted with EtOAc (3 x 50 mL), washed with brine (30 mL), anhydrous Na_2_SO_4_, filtered, and concentrated. The crude was purified by combiflash (eluent 30% EtOAc in hexane) to afford (*E*)-3-bromo-1-(methylsulfonyl) prop-1-ene (**LXXX**) as colorless liquid (0.6 g, 85% yield). Step-3: To a stirred solution of (*E*)-3-bromo-1-(methylsulfonyl)prop-1-ene (**LXXX**) (1.0 eq.) in THF/DMF/DCM, were added TEA/K_2_CO_3_/Cs_2_CO_3_ (1.5–2.0 eq.) and corresponding amines (1.0 eq.). Reaction mixture was stirred at 25 °C for 2–3 h. After completion, the reaction was poured into water, extracted with DCM, washed with brine, dried over anhydrous Na_2_SO_4_, filtered, and concentrated (temp. < 30 °C). The resulting crude was purified by combiflash to afford intermediates **LXXXIa–e** (26–93% yields). Step-4: (*E*)-*N*-cyclopropyl-3-(2-ethoxyphenyl)-*N*-(3-(methylsulfonyl)allyl)isoxazole-5-carboxamide **23b** was furnished by amide coupling of 3-(2-ethoxyphenyl)isoxazole-5-carboxylic acid (**XI**) and (*E*)-*N*-(3-(methylsulfonyl)allyl)cyclopropanamine (**LXXXIa**) following General Procedure-II as white solid (0.01 g, 4% yield). ^1^H NMR (400 MHz, DMSO-*d*_6_): *δ* 7.93 (d, *J* = 6.4 Hz, 1H), 7.54 (t, *J* = 8.0 Hz, 1H), 7.25 (d, *J* = 8.4 Hz, 1H), 7.15 (t, *J* = 7.6 Hz, 2H), 6.88 (s, 2H), 4.36 (d, *J* = 2.0 Hz, 2H), 4.26 – 4.21 (m, 2H), 3.06 – 3.00 (m, 4H), 1.46 (t, *J* = 6.8 Hz, 3H), 0.68 (s, 4H). ^13^C NMR (101 MHz, DMSO-*d*_6_): *δ* 165.8, 163.5, 160.3, 155.8, 141.9, 132.7, 131.5, 127.6, 121.3, 115.3, 113.5, 103.6, 64.6, 47.7, 42.6, 31.9, 14.9, 9.5. HRMS (ESI) *m/z*: [M+H]^+^ calculated for C_19_H_23_N_2_O_5_S: 391.1328, found 391.1323. HPLC purity (254 nm) >99%.

**(*E*)-3-(2-Ethoxyphenyl)-*N*-isopropyl-*N*-(3-(methylsulfonyl)allyl)isoxazole-5-carboxamide (23c)**. The compound was synthesized by amide coupling of 3-(2-ethoxyphenyl)isoxazole-5-carboxylic acid **XI** and (*E*)-*N*-isopropyl-3-(methylsulfonyl)prop-2-en-1-amine (**LXXXIb**) following General Procedure-I as a white sticky solid (0.05 g, 12% yield). ^1^H NMR (400 MHz, DMSO-*d*_6_): *δ* 7.89 (d, *J* = 7.6 Hz, 1H), 7.50 (t, *J* = 7.0 Hz, 1H), 7.22 (d, *J* = 8.4 Hz, 1H), 7.12 (t, *J* = 7.6 Hz, 1H), 7.03 (s, 1H), 6.80 (s, 2H), 4.28 – 4.23 (m, 5H), 2.99 (s, 3H), 1.42 (t, *J* = 7.0 Hz, 3H), 1.22 (d, 6H). ^13^C NMR (100 MHz, DMSO-*d*_6_): *δ* 166.3, 161.4, 159.9, 159.7, 143.9, 142.9, 132.8, 131.9, 131.5, 127.6, 115.1, 113.4, 103.7, 69.9, 50.6, 46.8, 42.6, 21.2, 20.1, 14.9. HRMS (ESI) *m/z*: [M+H]^+^ calculated for C_19_H_25_N_2_O_5_S: 393.1484, found 393.1481. HPLC purity (254 nm) >99%.

**(*E*)-*N*-Cyclopentyl-3-(2-ethoxyphenyl)-*N*-(3-(methylsulfonyl)allyl)isoxazole-5-carboxamide (23d).** The compound was synthesized as a white solid (0.012 g, 4% yield, m.p. 120 °C) by amide coupling of 3-(2-ethoxyphenyl)isoxazole-5-carboxylic acid (**XI**) and (*E*)-*N*-(3-(methylsulfonyl)allyl)cyclopentanamine (**LXXXIc**) using EDC/HOBt/DIPEA in DMF at 25 °C for 2 h. ^1^H NMR (400 MHz, DMSO-*d*_6_): *δ* 7.90 – 7.88 (m, 1H), 7.52 – 7.48 (m, 1H), 7.23 (d, *J* = 8.0 Hz, 1H), 7.12 (t, *J* = 7.2 Hz, 1H), 7.02 (s, 1H), 6.78 (s, 2H), 4.49 (s, 1H), 4.29 (d, *J* = 4.8 Hz, 1H), 4.26 – 4.23 (m, 2H), 2.98 (s, 3H), 1.86 (s, 2H), 1.70 (s, 2H), 1.62 (d, *J* = 3.6 Hz, 2H), 1.50 (d, *J* = 8.8 Hz, 2H) 1.44 (t, *J* = 7.2 Hz, 3H). ^13^C NMR (101 MHz, DMSO-*d*_6_): *δ* 166.3, 161.5, 159.7, 155.8, 142.8, 132.8, 131.8, 131.2, 127.6, 121.3, 115.1, 113.4, 103.8, 64.6, 60.0, 42.6, 29.9, 28.9, 23.8, 23.6, 14.9. HRMS (ESI) *m/z*: [M+H]^+^ calculated for C_21_H_27_N_2_O_5_S: 419.1641, found 419.1637. HPLC purity (254 nm) >99%.

**(*E*)-3-(2-Ethoxyphenyl)-*N*-(3-(methylsulfonyl)allyl)-*N*-pentylisoxazole-5-carboxamide (23e).** The compound was synthesized by amide coupling of 3-(2-ethoxyphenyl)isoxazole-5-carboxylic acid (**XI**) and (*E*)-*N*-(3-(methylsulfonyl)allyl)pentan-1-amine (**LXXXId**) following General Procedure-II as yellow liquid (0.05 g, 12% yield). ^1^H NMR (400 MHz, DMSO-*d*_6_): *δ* 7.79 (d, *J* = 7.6 Hz, 1H), 7.49 (t, *J* = 7.8 Hz, 2H), 7.24 (s, 1H), 7.19 (d, *J* = 8.4 Hz, 1H), 7.07 (t, *J* = 6.8 Hz, 1H), 6.88 – 6.75 (m, 2H), 4.35 (s, 2H), 4.21 – 4.16 (m, 2H), 3.48 (t, *J* = 7.2 Hz, 2H), 3.00 (s, 3H), 1.64 – 1.59 (m, 2H), 1.43 – 1.36 (m, 3H), 1.28 (s, 4H), 0.88 (s, 3H). ^13^C NMR (101 MHz, DMSO-*d*_6_): *δ* 163.1, 160.0, 158.6, 156.7, 141.8, 141.0, 132.6, 132.3, 131.9, 129.4, 121.4, 116.7, 113.6, 108.5, 108.1, 64.4, 49.2, 48.8, 46.9, 46.4, 42.6, 28.9, 28.7, 26.8, 22.3, 22.2, 14.9, 14.3. HRMS (ESI) *m/z*: [M+H]^+^ calculated for C_21_H_29_N_2_O_5_S: 421.1797, found 421.1799. HPLC purity (254 nm) >99%.

**(*E*)-*N*-Benzyl-3-(2-ethoxyphenyl)-*N*-(3-(methylsulfonyl)allyl)isoxazole-5-carboxamide (23f).** The compound was synthesized by amide coupling of 3-(2-ethoxyphenyl)isoxazole-5-carboxylic acid (**XI**) and (*E*)-*N*-benzyl-3-(methylsulfonyl)prop-2-en-1-amine (**LXXXIe**) following General Procedure-II as white solid (0.03 g, 12% yield, m.p. 52 °C). ^1^H NMR (400 MHz, DMSO-*d*_6_): *δ* 7.79 (d, *J* = 7.6 Hz, 1H), 7.50 (t, *J* =15.6 Hz 1H), 7.41 (q, *J* = 28.8 Hz, 5H), 7.25 (s, 1H), 7.18 (d, *J* = 8 Hz, 1H), 7.08 (t, *J* = 15.2 Hz, 1H), 6.82 (q, *J* = 34.4 Hz, 2H), 4.77 (s, 2H), 4.32 (s, 2H), 4.14 (d, *J* = 6.0 Hz, 2H), 3.06 (dd, *J* = 1.6 Hz, 3H), 1.30 (d, *J* = 14.4 Hz, 3H). ^13^C NMR (101 MHz, DMSO-*d*_6_): *δ* 162.7, 162.5, 159.9, 158.9, 158.9, 156.7, 156.6, 141.4, 140.4, 136.8, 136.7, 132.6, 132.4, 132.1, 129.4, 129.3, 129.1, 128.5, 128.2, 128.0, 127.3, 121.2, 116.5, 113.5, 64.4, 52.3, 49.7, 48.8, 46.6, 42.6. HRMS (ESI) *m/z*: [M+H]^+^ calculated for C_23_H_25_N_2_O_5_S: 441.1484, found 441.1481; HPLC purity (254 nm) >99%.

### Biological Assays

The CHIKV nsP2 protease, CHIKV-nLuc viral replication, and VEEV-nLuc viral replication assays were performed as described in reference 7.

### Kinetic Solubility Assay

Kinetic solubility of final compounds was determined in triplicate in pH 7.4 phosphate buffer using 10 mM stock solutions at 2.0 % DMSO and CLND detection at Analiza, Inc.

### Associated Content

Supplemental Information. Table S1. Binding Kinetics of Covalent nsP2 Protease Inhibitors; Table S2. VEEV-nLuc viral replication assay data; Figure S1. Compound aggregation data for **1a**; Analytical spectra: ^1^H and ^13^C NMR spectra for new compounds, HPLC analysis of pyrazoles, chiral SFC of (*R*)-**24a** and (*S*)-**24a**.

## Acknowledgements

The Structural Genomics Consortium (SGC) is a registered charity (no: 1097737) that receives funds from Bayer AG, Boehringer Ingelheim, Bristol Myers Squibb, Genentech, Genome Canada, through Ontario Genomics Institute [OGI-196], EU/EFPIA/OICR/McGill/KTH/Diamond Innovative Medicines Initiative 2 Joint Under-taking [EUbOPEN grant 875510], Janssen, Merck KGaA (also known as EMD in Canada and the US), Pfizer, and Takeda. We gratefully acknowledge the support of Piramal Pharma Solutions for assistance in the synthesis of intermediates and final compounds. The research reported in this publication was supported by NIH grant 1U19AI171292-01 (READDI-AViDD Center) and in part by the NC Biotech Center Institutional Support Grant 2018-IDG-1030 and by NIH grant S10OD032476 for upgrading the 500 MHz NMR spectrometer in the UNC Eshelman School of Pharmacy NMR Facility.

