## Supporting Information for "Structure Activity of β-Amidomethyl Vinyl Sulfones as Covalent Inhibitors of *Chikungunya* nsP2 Cysteine Protease with Anti-alphavirus Activity"



| Table of Contents | | Pages |
|---|---|---|
| Table S1 | Binding kinetics of covalent nsP2 protease inhibitors | S2 |
| Table S2 | VEEV-nLuc $pEC_{50}$ for compounds with $pEC_{50}$ >6 on CHIKV-nLuc | S3 |
| Figure S1 | DLS data for compound **1a** | S4 |
| Figures S2–S153 | NMR spectra of final compounds | S5–S80 |
| Figures S154–S206 | HPLC analysis of pyrazole analogs | S81–S95 |
| Figures S207–S208 | Chiral SFC of (*R*)-**24a** and (*S*)-**24a** | S96–S97 |

**Table S1.** Binding kinetics of covalent nsP2 protease inhibitors

**Binding Kinetics Assay.** CHICKV nsP2 protease activity was measured using an internally quenched peptide substrate continuously over 45 min as previously described.[1] 2-fold serial-dilutions of inhibitor were used to measure $k_{obs}$ values and the data were fit to the equation ($k_{obs}$ = ($k_{inact}$ x [I]) / ($K_i$ + [I]) to determine the potency of the initial binding event ($K_i$) and maximum potential rate of covalent bond formation ($k_{inact}$).

| **Compound** | **$IC_{50}$ (μM)** | **$K_i$ (μM)** | **$k_{inact}/K_i$ ($M^{-1}s^{-1}$)** |
|---|---|---|---|
| **1a** | 0.06 | 0.28 | 6400 |
| **1b** | 0.13 | 1.9 | 1500 |
| **1c** | 0.08 | 0.64 | 2600 |
| **1e** | 0.07 | 0.30 | 3600 |
| **1g** | 0.06 | 1.5 | 2000 |
| **1h** | 0.04 | 0.20 | 4500 |
| **1k** | 0.18 | 0.29 | 3300 |
| **1l** | 0.17 | 0.79 | 1400 |
| **1o** | 0.03 | 0.11 | 9100 |
| **4c** | 0.10 | 0.80 | 1500 |
| **4d** | 0.05 | 0.25 | 5900 |
| **4f** | 0.10 | 1.7 | 1000 |
| **8d** | 0.02 | 0.06 | 12000 |
| **10** | 0.04 | 0.29 | 6000 |
| **14** | 0.15 | 0.46 | 1900 |
| **20** | 1.4 | 16 | 100 |
| **24a** | 0.13 | 0.30 | 1500 |
| **25d** | 0.15 | 0.86 | 1900 |

(1) Merten, E. M.; Sears, J. D.; Leisner, T. M.; Hardy, P. B.; Ghoshal, A.; Hossain, M. A.; Asressu, K. H.; Brown, P. J.; Stashko, M. A.; Herring, L. E.; et al. Discovery of a cell-active chikungunya virus nsP2 protease inhibitor using a covalent fragment-based screening approach. *bioRxiv* **2024**. DOI: 10.1101/2024.03.22.586341.

**Table S2.** VEEV-nLuc $pEC_{50}$ for compounds with $pEC_{50}$ >6 on CHIKV-nLuc

| Compound | CHIKV-nLuc ($pEC_{50}$) | VEEV-nLuc ($pEC_{50}$) |
|---|---|---|
| **1a** | 7.4 | 6.4 |
| **1b** | 7.3 | 5.5 |
| **1c** | 7.3 | 5.9 |
| **1d** | 6.1 | 5.1 |
| **1e** | 7.1 | 6.1 |
| **1f** | 6.3 | <5 |
| **1g** | 7.1 | 6.0 |
| **1h** | 7.0 | 6.2 |
| **1j** | 6.6 | 6.0 |
| **1k** | 6.7 | 5.7 |
| **1l** | 7.4 | 6.2 |
| **1n** | 7.2 | 6.5 |
| **1o** | 7.4 | 6.5 |
| **4c** | 7.1 | 6.2 |
| **4d** | 6.7 | 6.2 |
| **4e** | 7.0 | 6.2 |
| **4f** | 7.3 | 5.9 |
| **4g** | 6.9 | 6.0 |
| **4i** | 6.7 | 6.0 |
| **5** | 6.6 | 5.1 |
| **6** | 6.2 | 5.6 |
| **8d** | 6.5 | 5.8 |
| **9** | 7.1 | 6.9 |
| **10** | 7.3 | 6.3 |
| **11** | 6.6 | 5.8 |
| **12** | 6.5 | 5.6 |
| **13** | 6.7 | 5.7 |
| **14** | 7.2 | 6.1 |
| **15** | 7.2 | 6.5 |
| **16** | 6.9 | 6.1 |
| **17** | 6.8 | 6.0 |
| **18** | 6.4 | 5.8 |
| **19** | 7.0 | 6.0 |
| **20** | 6.2 | 5.7 |
| **21** | 6.0 | 5.8 |
| **23b** | 6.5 | 5.6 |
| **24a** | 6.4 | 5.7 |
| **(*R*)-24a** | 6.3 | 6.1 |
| **24e** | 6.5 | 5.6 |
| **25a** | 7.1 | 6.1 |
| **25b** | 6.5 | 5.5 |
| **25c** | 6.5 | 5.2 |
| **25e** | 6.6 | 5.7 |
| **27** | 6.3 | 5.6 |
| **28** | 6.0 | <5 |
| **31** | 6.2 | 5.4 |

**Figure S1.** DLS data for **1a**

**Dynamic Light Scattering (DLS) Assay**. Aggregation behavior of **1a** was determined by DLS in duplicate at 25 ºC in 25mM HEPES buffer (pH 7.4) containing 0.003% Tween-20 and 1mM DTT, using a 10 mM stock solution contianing 2% DMSO. Light scattering was measured using a DynaPro Plate Reader III. Buffer with 2% DMSO control produced an average laser intensity of 858 kCnts/s.

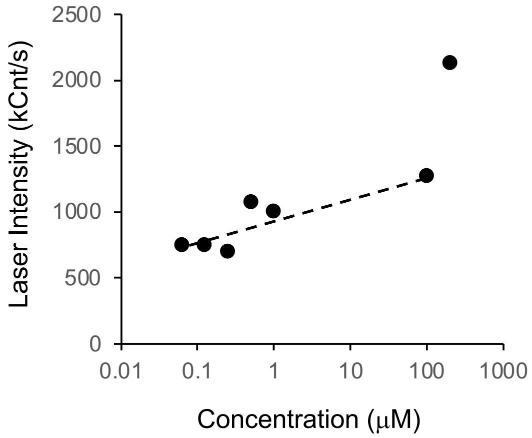

### NMR spectra of final compounds

**Figure S2**: $^{1}$H NMR (500 MHz, DMSO-$d_6$) for **1a**

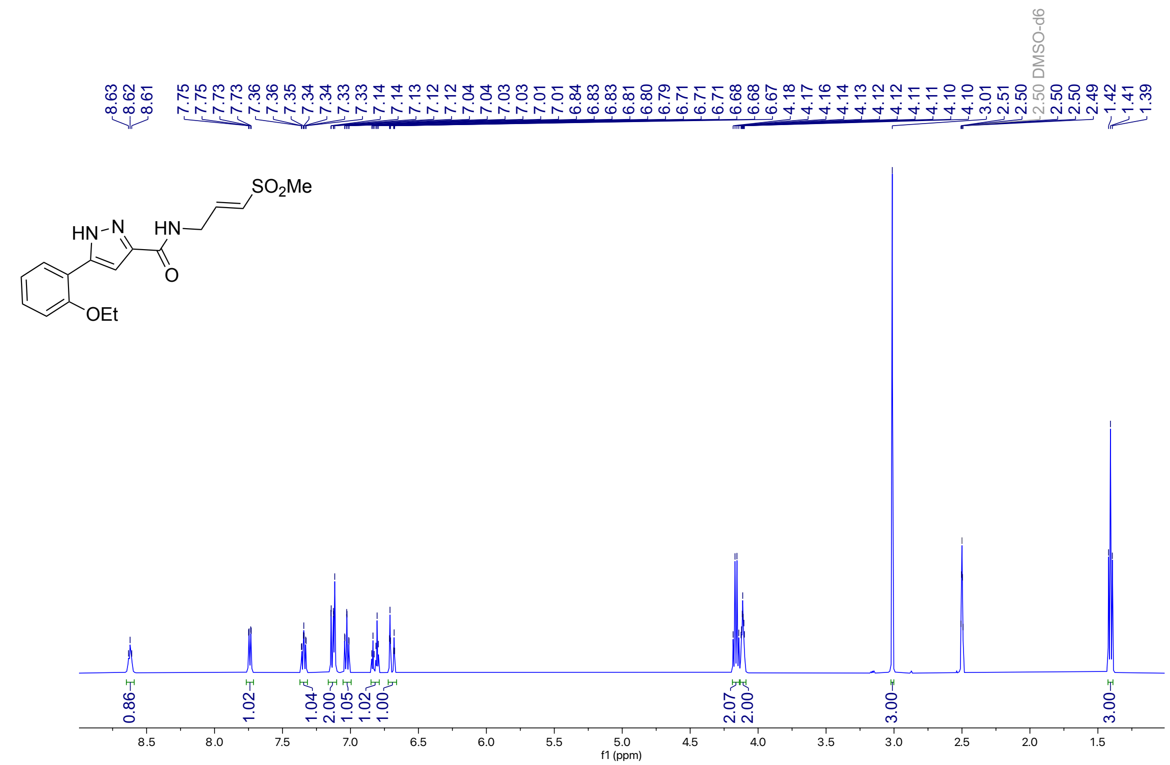


**Figure S3**: $^{13}$C NMR (126 MHz, DMSO-$d_6$) for **1a**

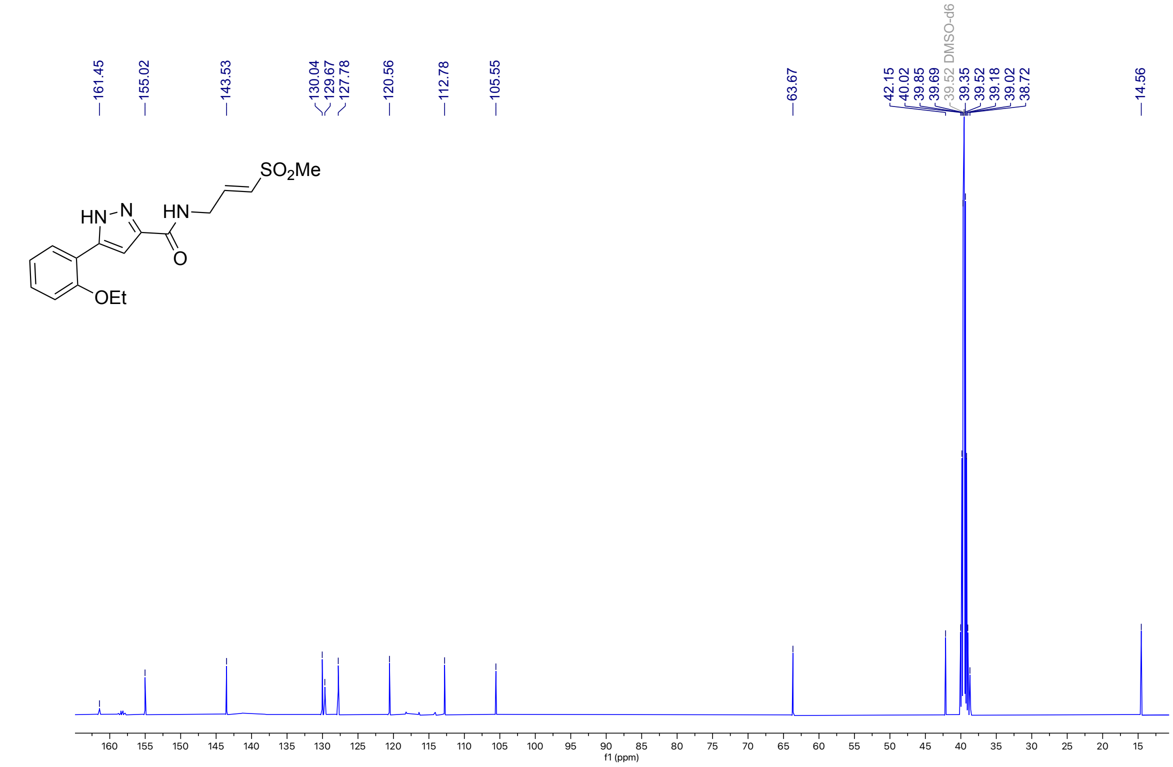

**Figure S4**: $^{1}$H NMR (500 MHz, DMSO-$d_6$) for **2**

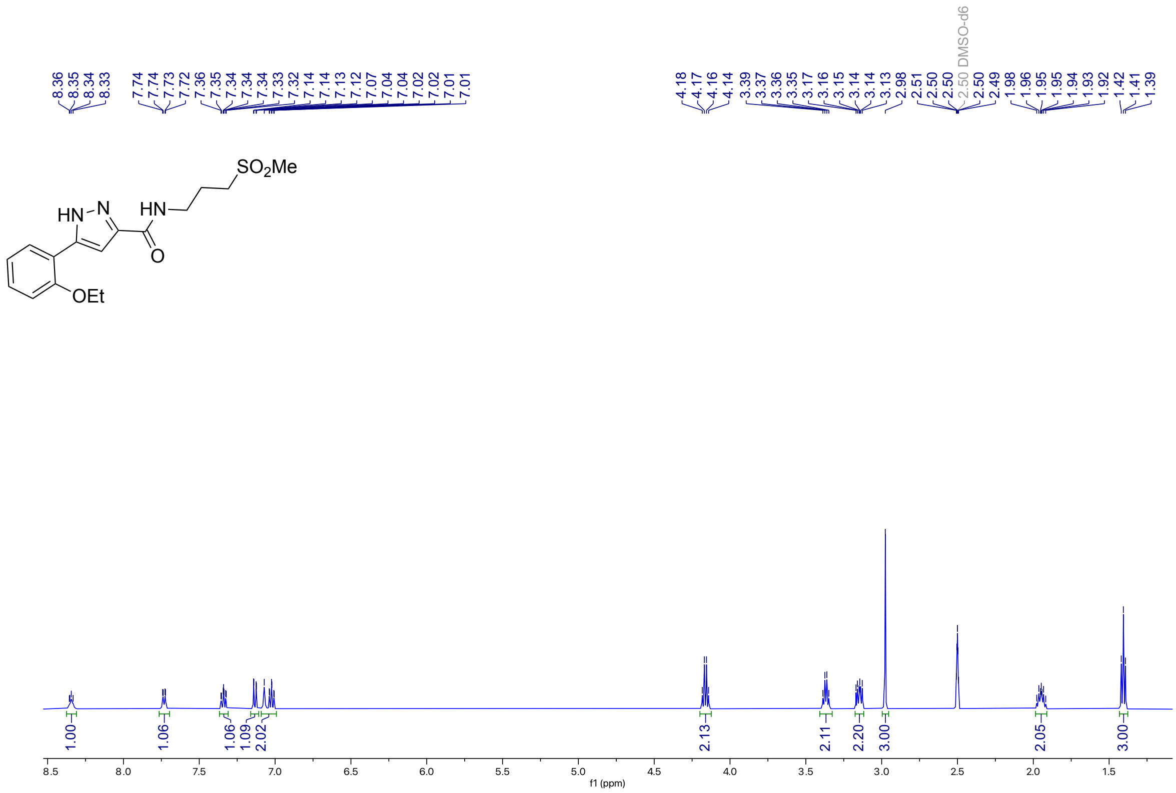

**Figure S5**: $^{13}$C NMR (126 MHz, DMSO-$d_6$) for **2**

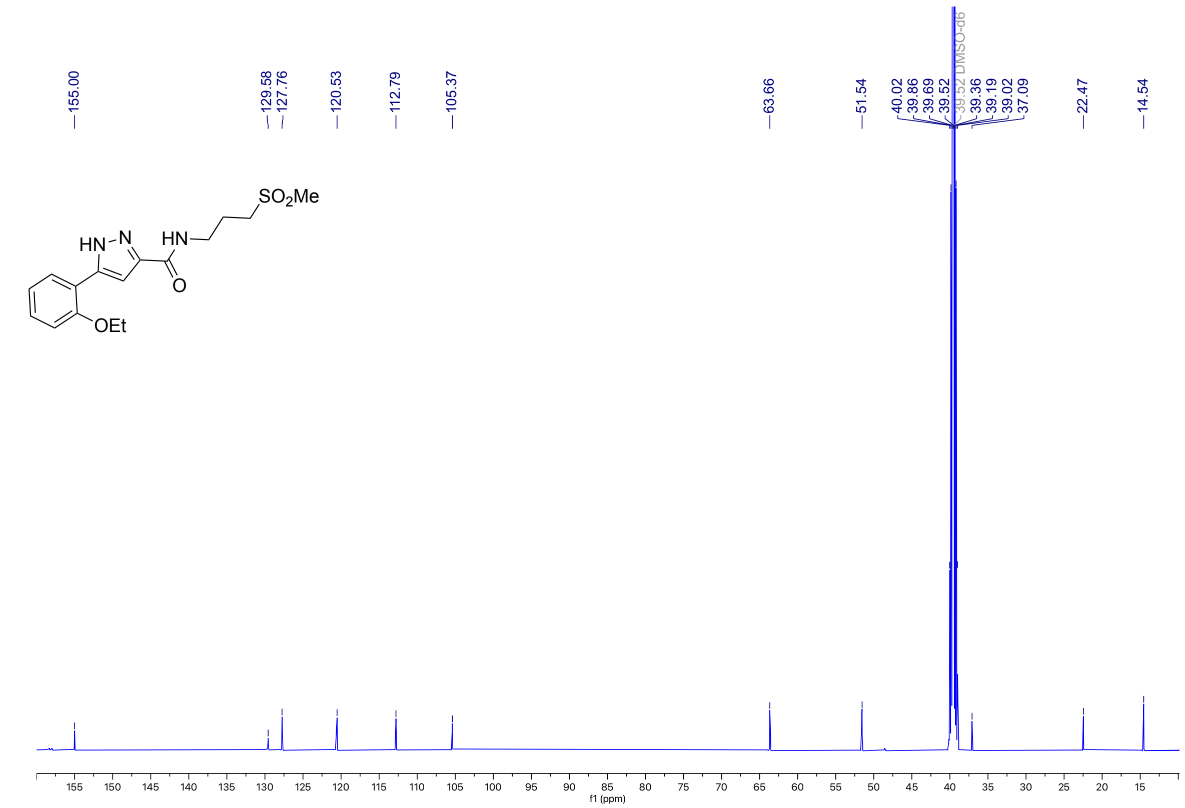

**Figure S6**: $^{1}$H NMR (400 MHz, DMSO-$d_6$) for RA-0002993-01 **3**

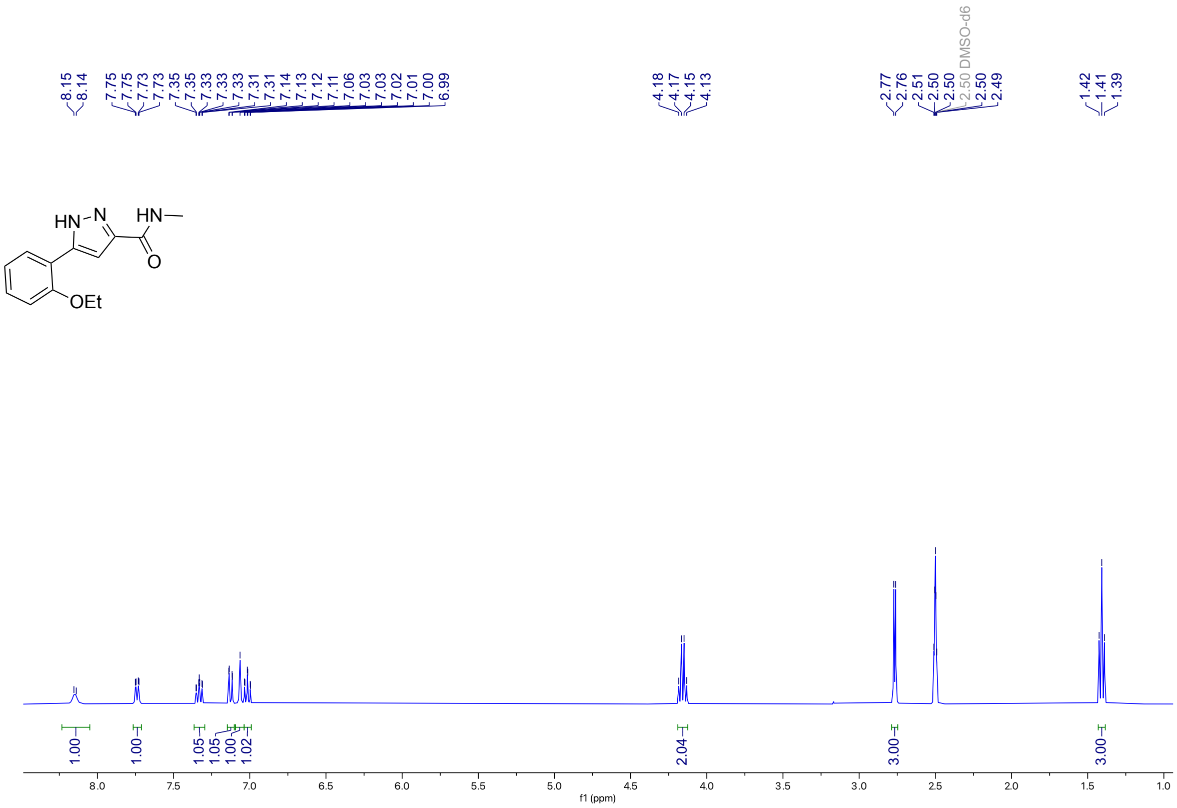

**Figure S7**: $^{13}$C NMR (126 MHz, DMSO-$d_6$) for **3**

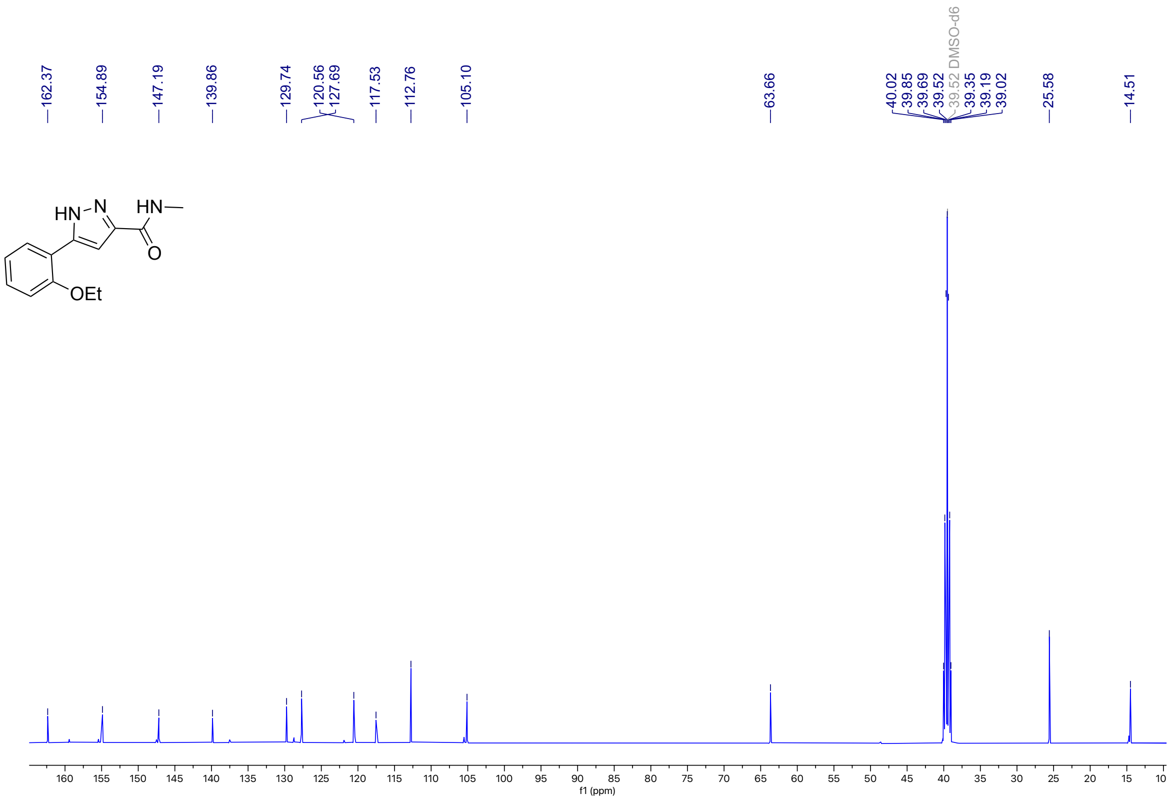

**Figure S8**: $^{1}$H NMR (400 MHz, DMSO-$d_6$) for **1b**

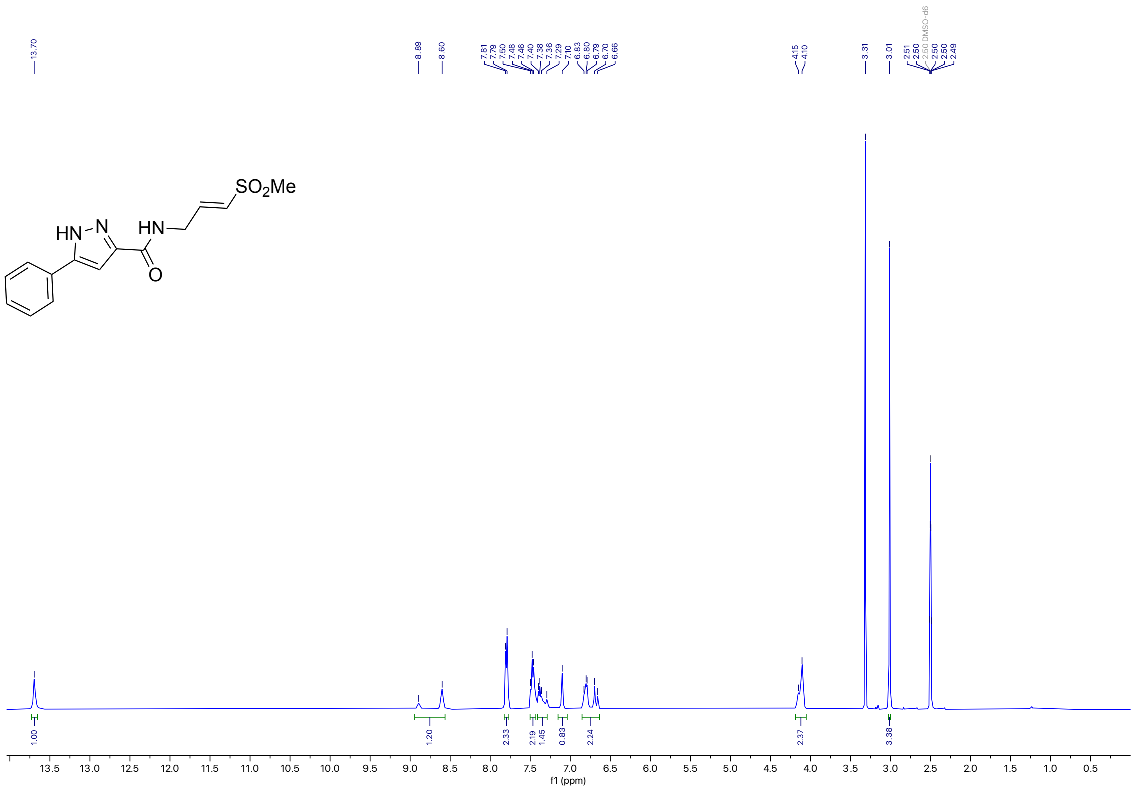


**Figure S9**: $^{13}$C NMR (100 MHz, DMSO-$d_6$) for **1b**

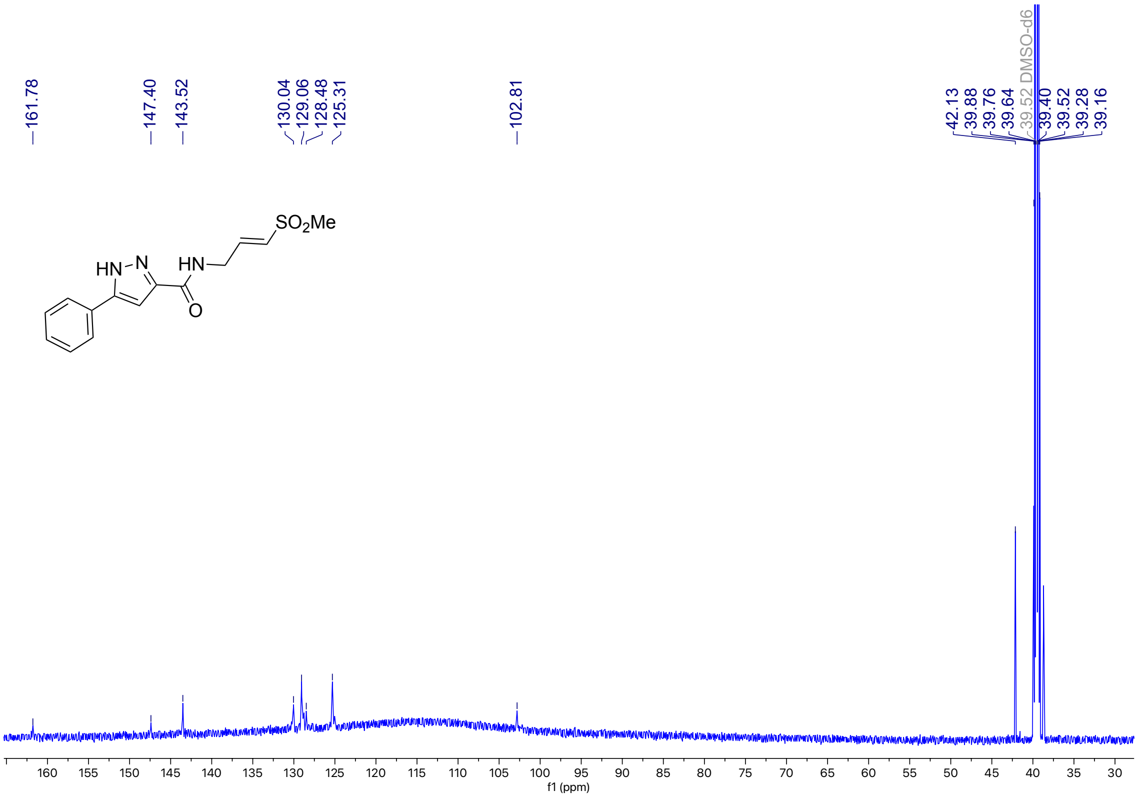

**Figure S10**: $^{1}$H NMR (500 MHz, DMSO-$d_6$) for **1c**

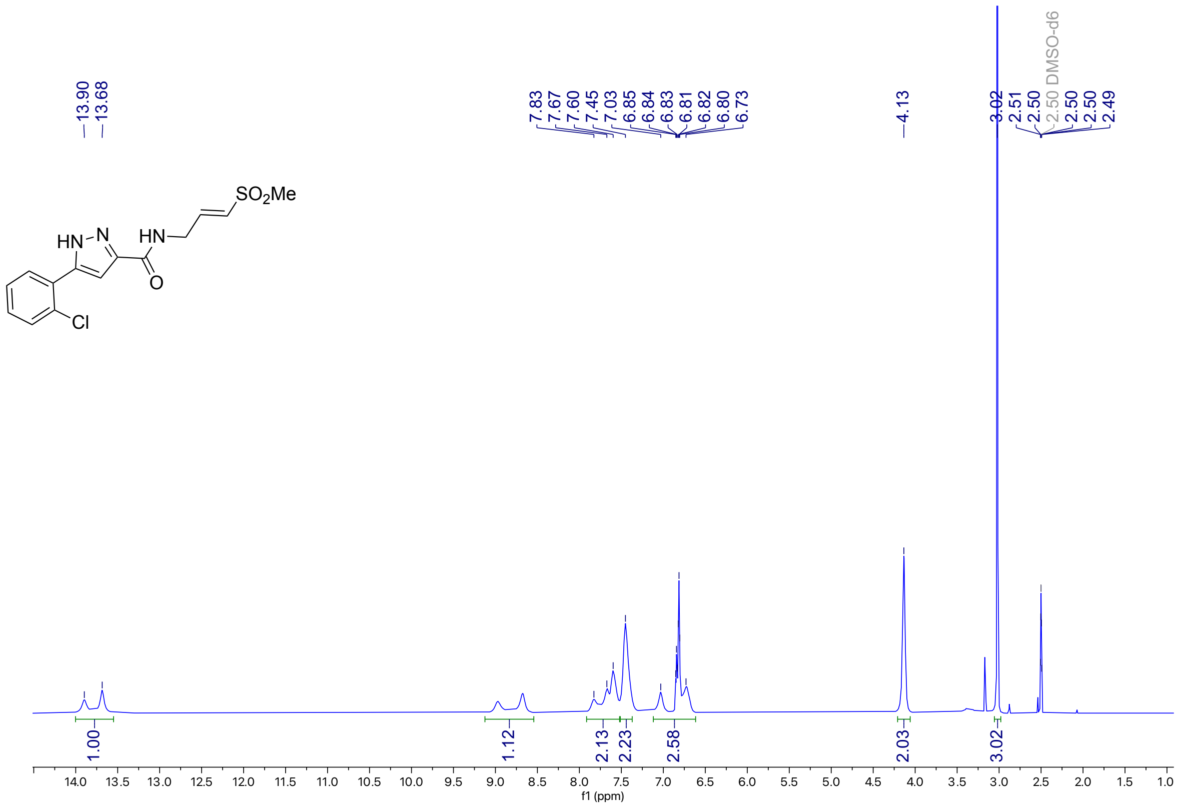

**Figure S11**: $^{13}$C NMR (176 MHz, DMSO-$d_6$) for **1c**

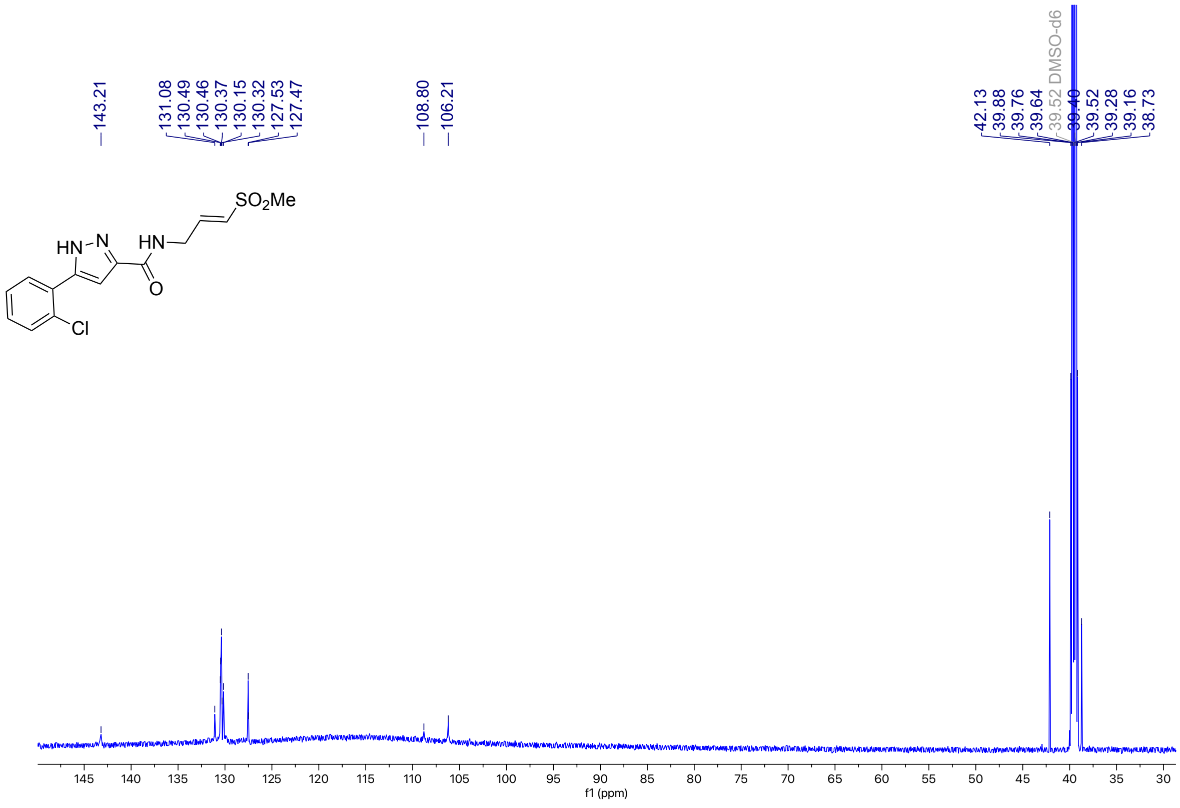

**Figure S12**: $^1$H NMR (500 MHz, DMSO-$d_6$) for **1d**

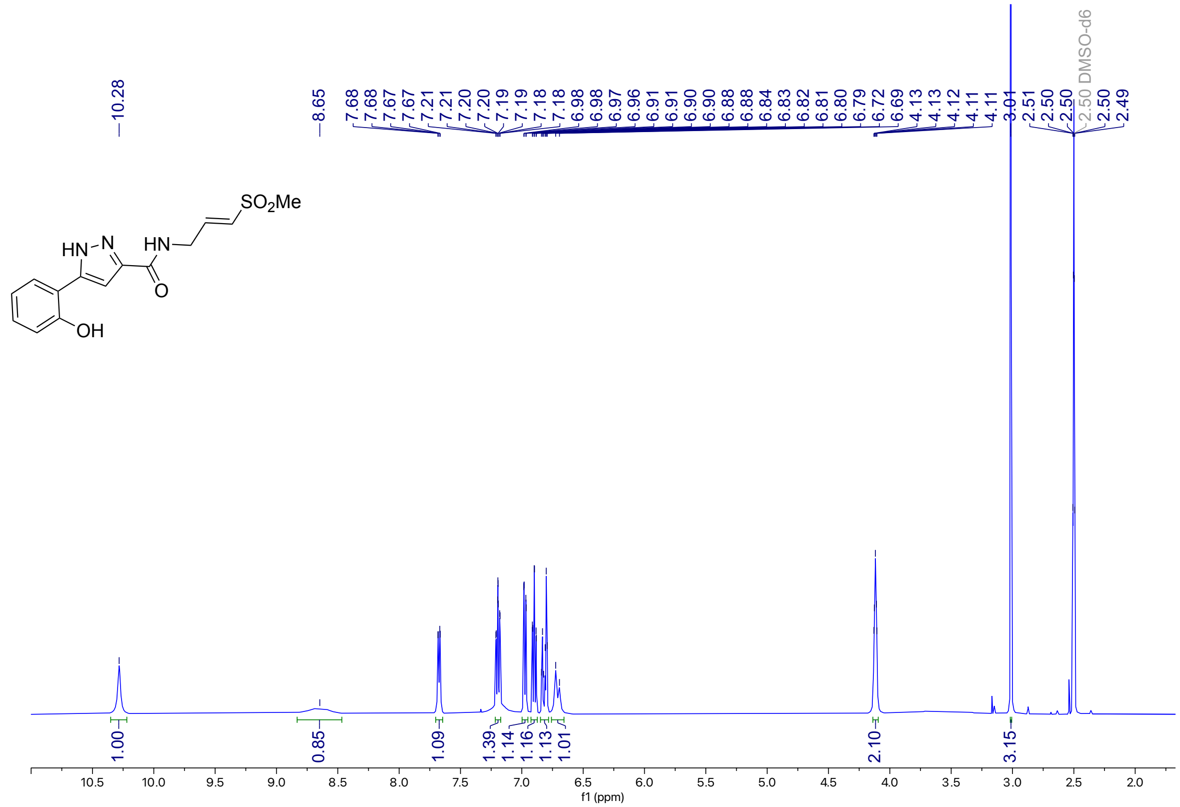

**Figure S13**: $^{13}$C NMR (126 MHz, DMSO-$d_6$) for **1d**

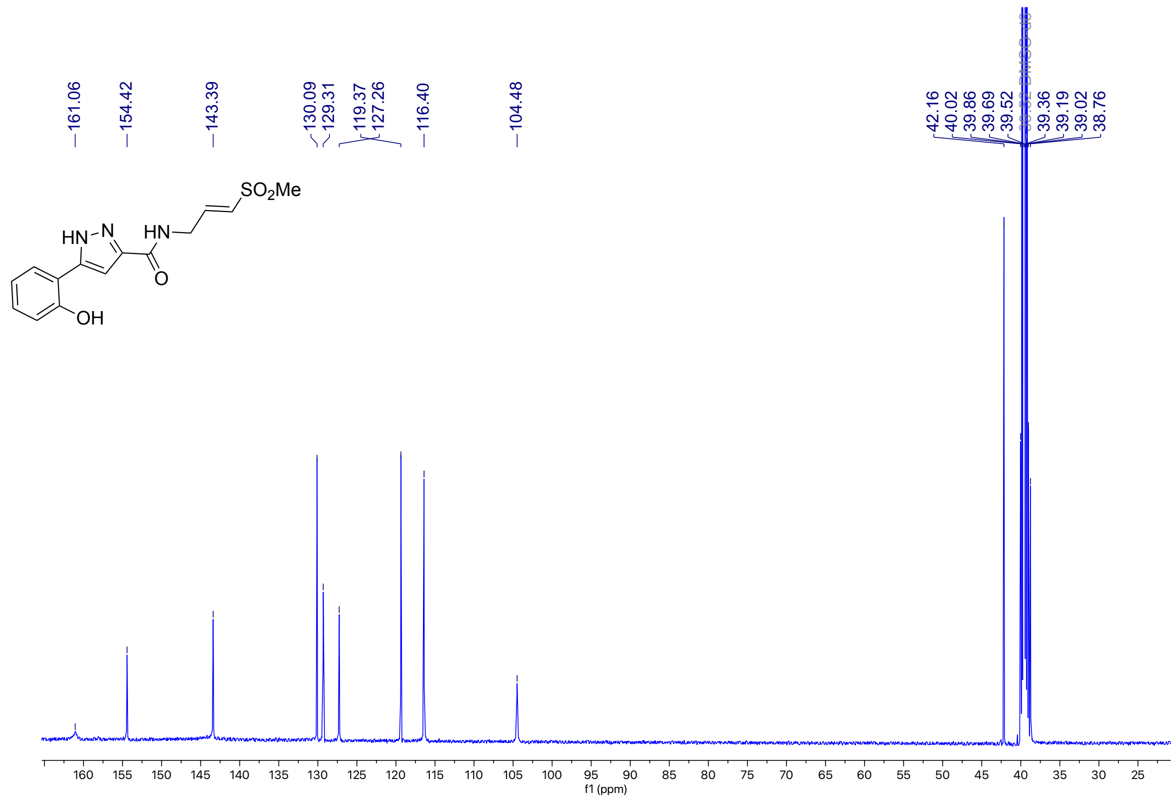

**Figure S14**: $^{1}$H NMR (500 MHz, DMSO-$d_6$) for **1e**

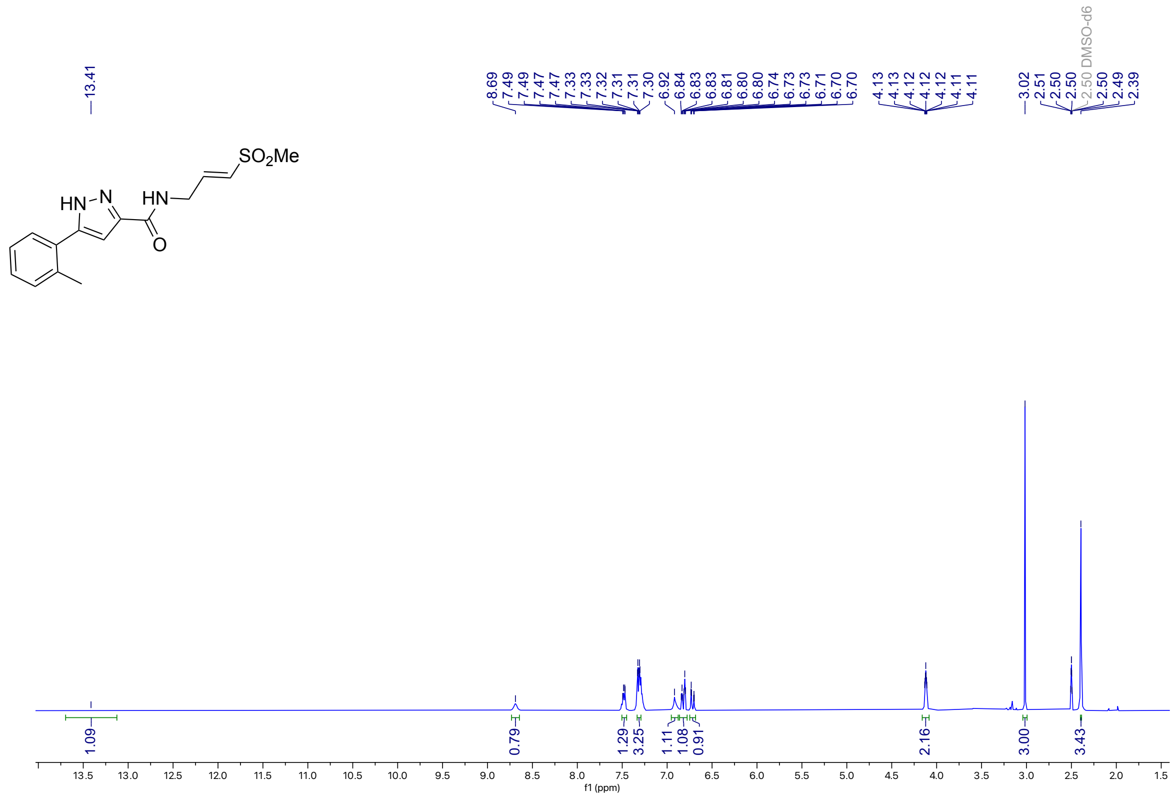


**Figure S15**: $^{13}$C NMR (214 MHz, DMSO-$d_6$) for **1e**

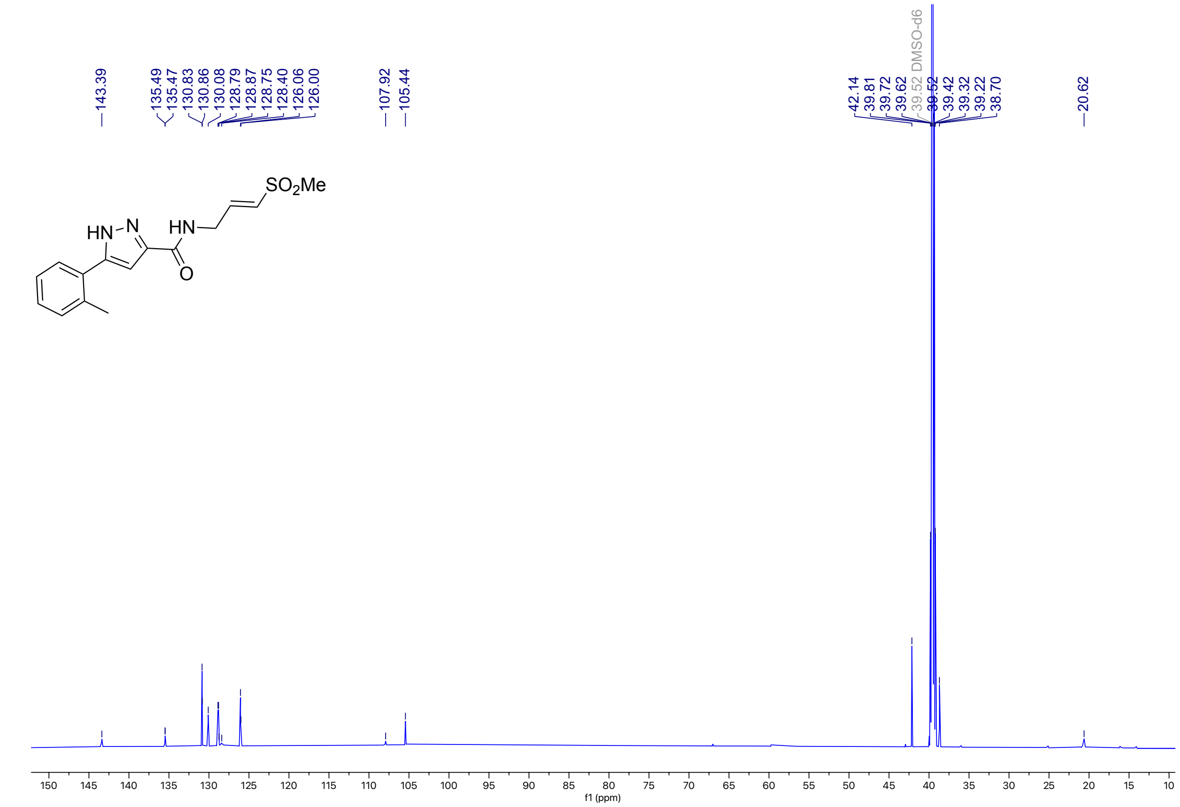

**Figure S16**: $^{1}H$ NMR (400 MHz, DMSO-$d_6$) for **1f**

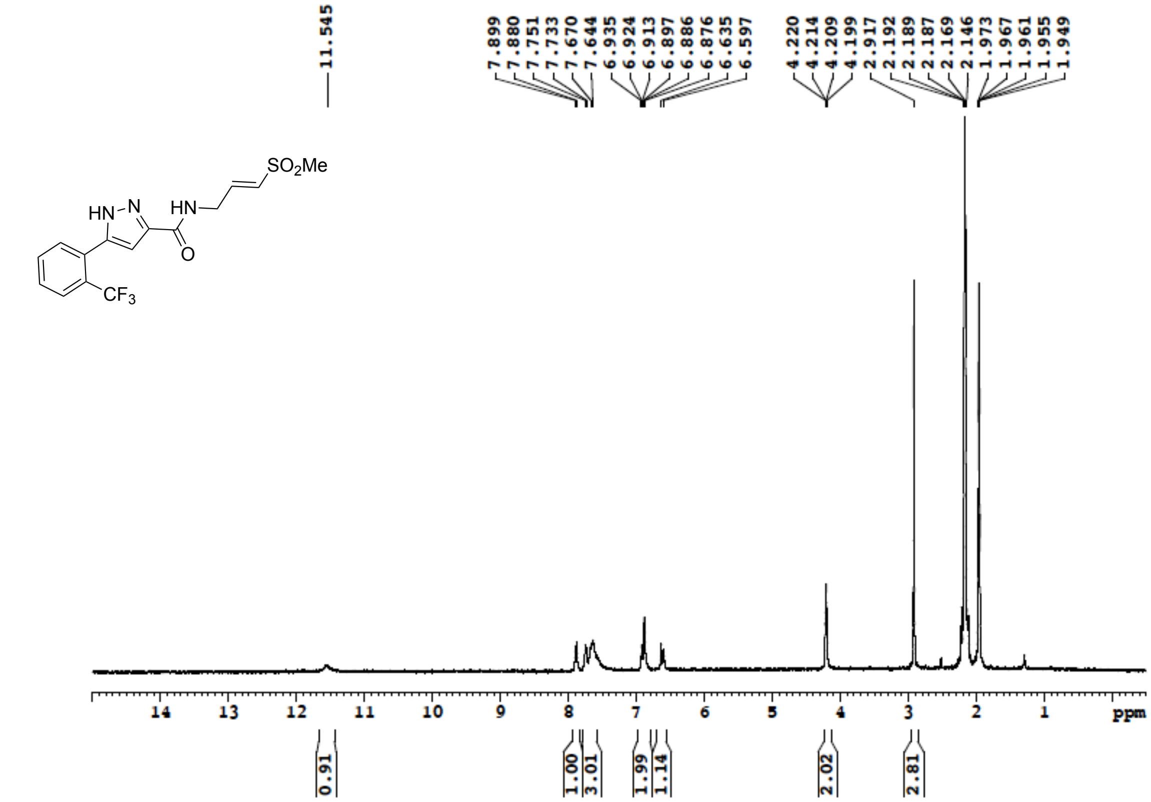


**Figure S17**: $^{13}C$ NMR (100 MHz, DMSO-$d_6$) for **1f**

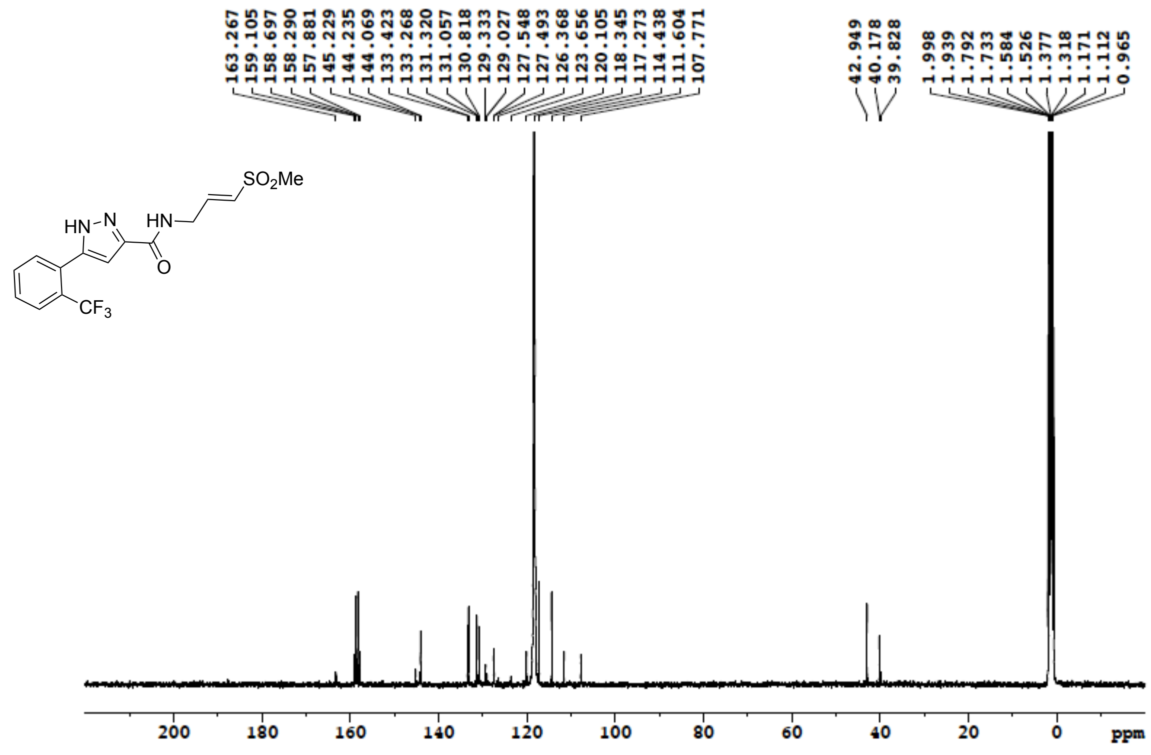

**Figure S18**: $^{1}$H NMR (500 MHz, DMSO-$d_6$) for **1g**

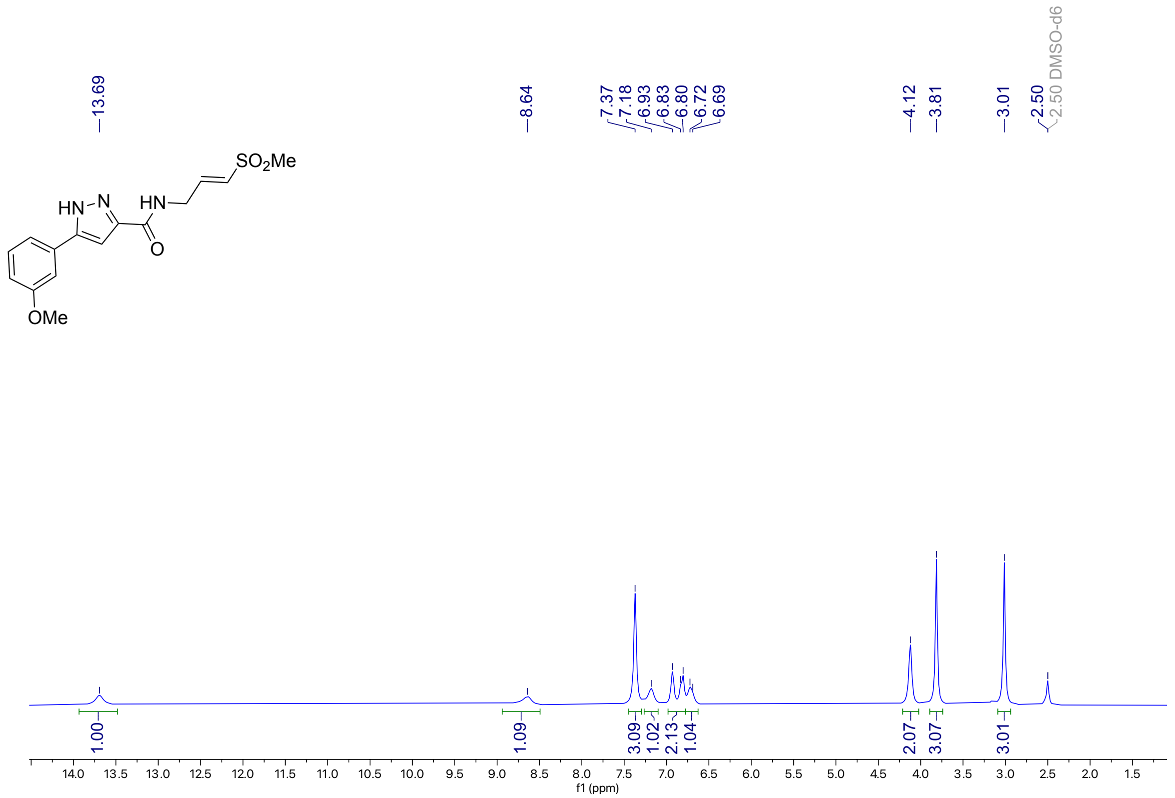


**Figure S19**: $^{13}$C NMR (126 MHz, DMSO-$d_6$) for **1g**

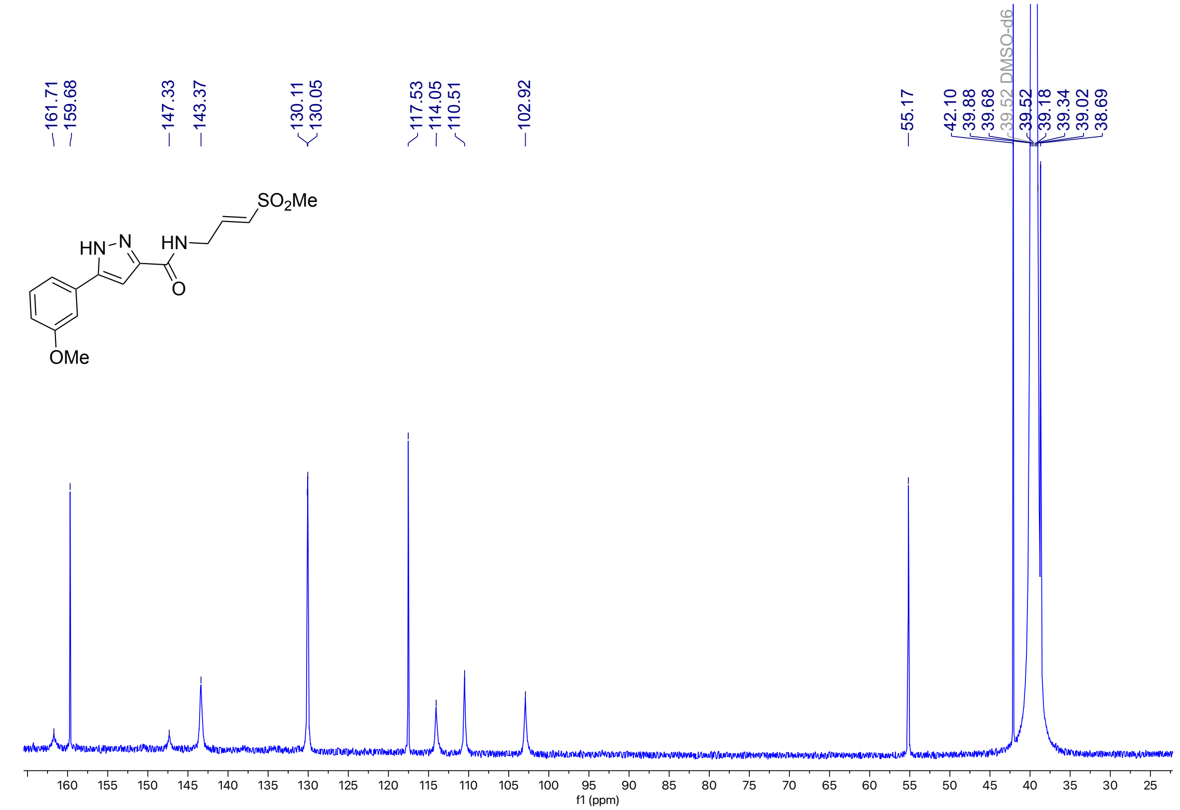

**Figure S20**: [1]H NMR (400 MHz, DMSO-$d_6$) for **1h**

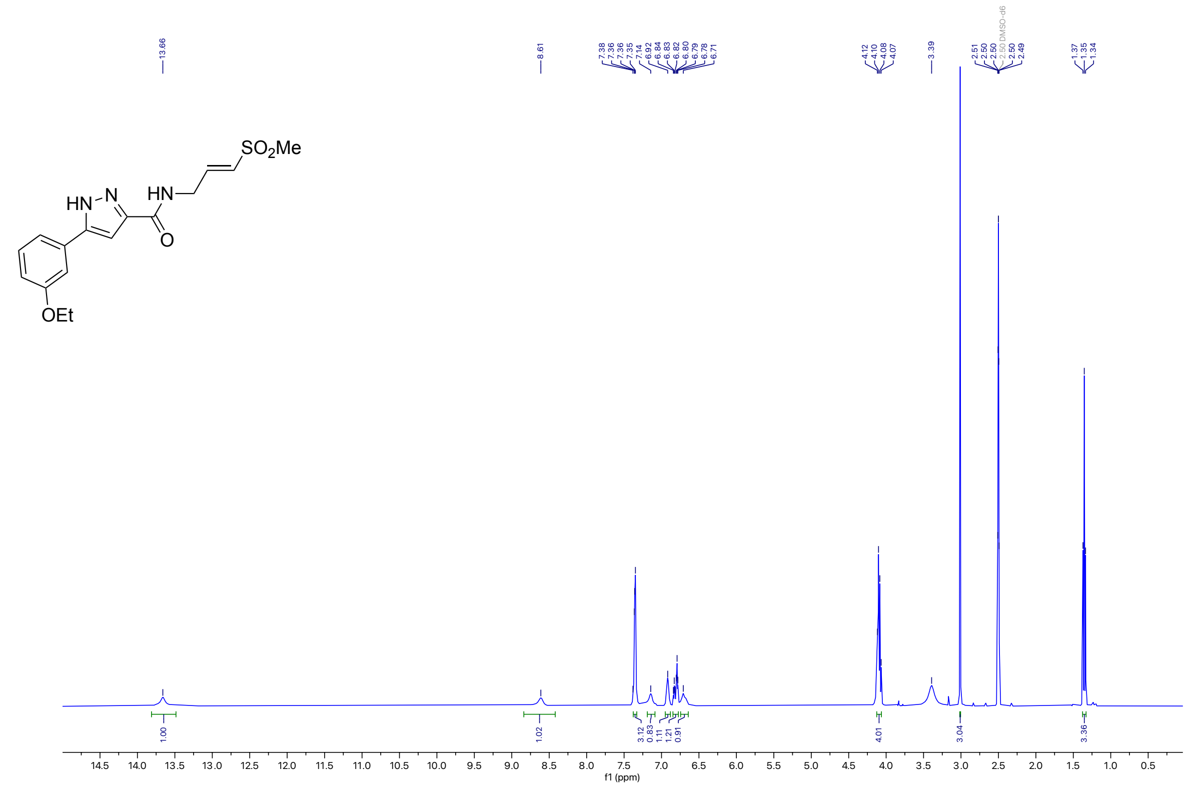

**Figure S21**: [13]C NMR (214 MHz, DMSO-$d_6$) for **1h**

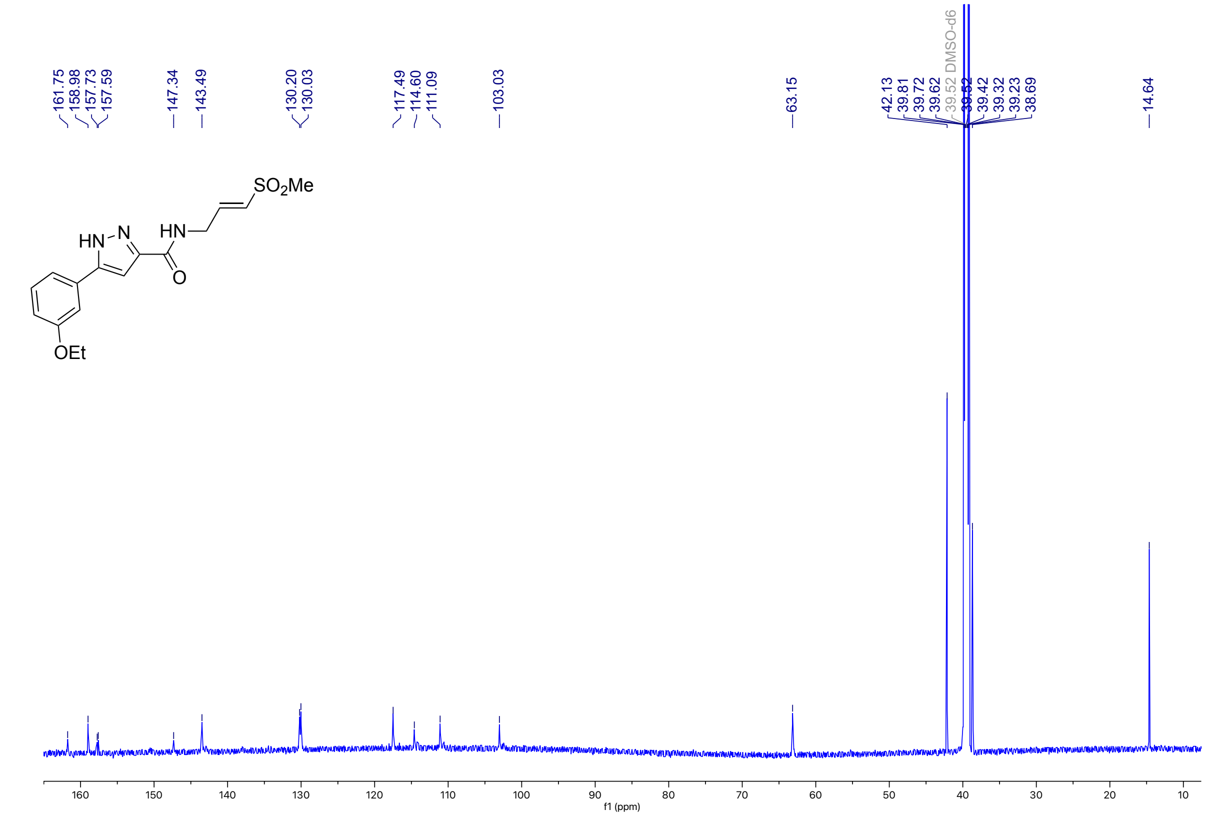

**Figure S22**: $^{1}$H NMR (400 MHz, DMSO-$d_6$) for **1i**

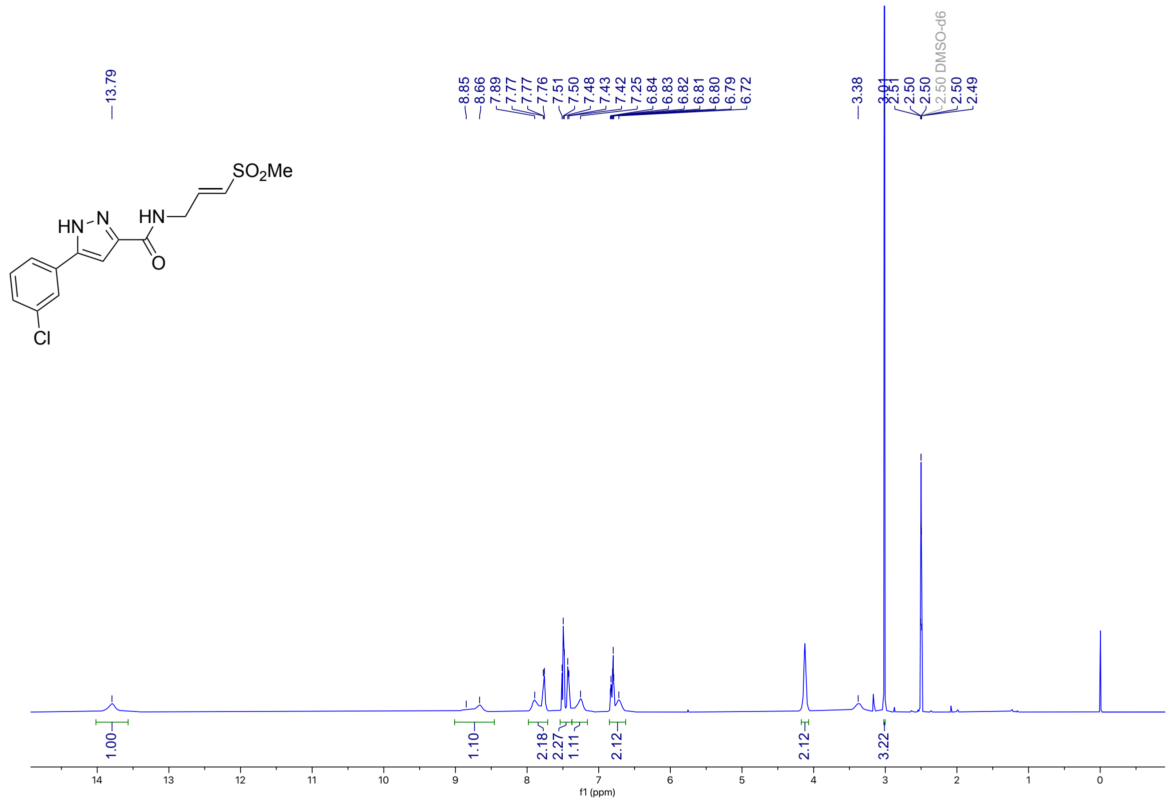

**Figure S23**: $^{13}$C NMR (100 MHz, DMSO-$d_6$) for **1i**

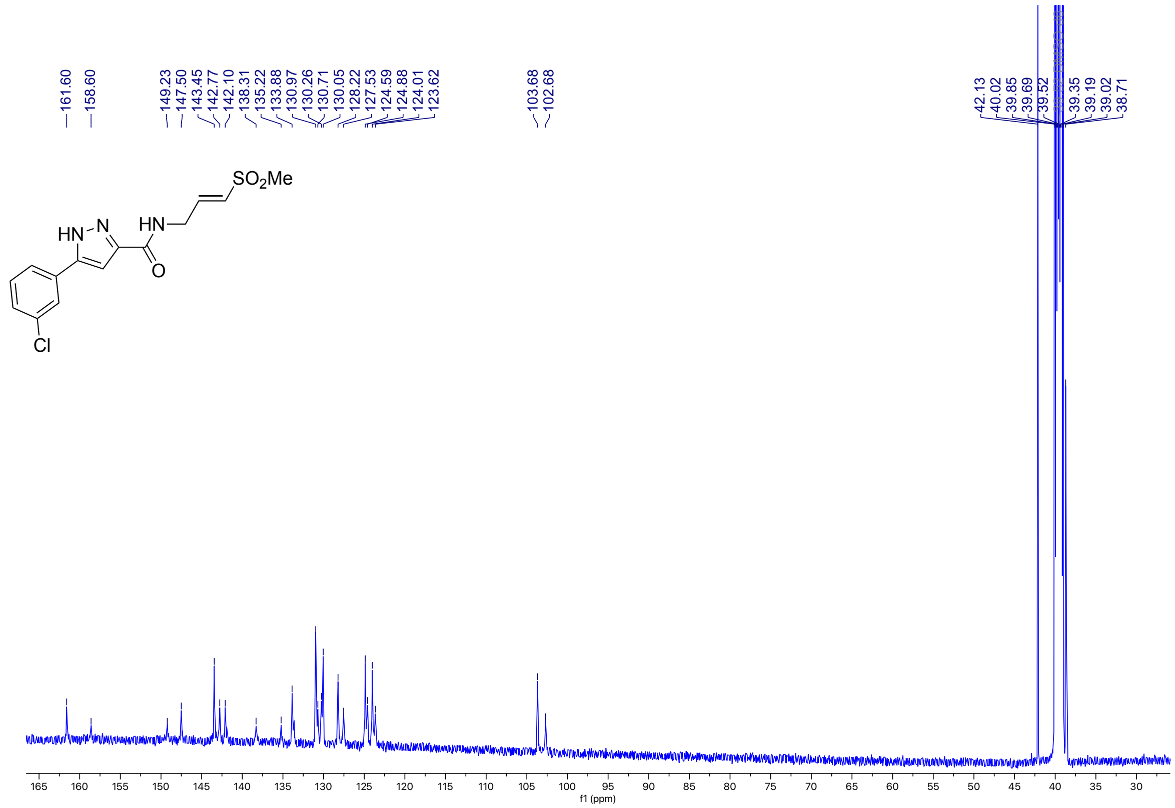

**Figure S24**: $^{1}$H NMR (500 MHz, MeOD) for **1j**

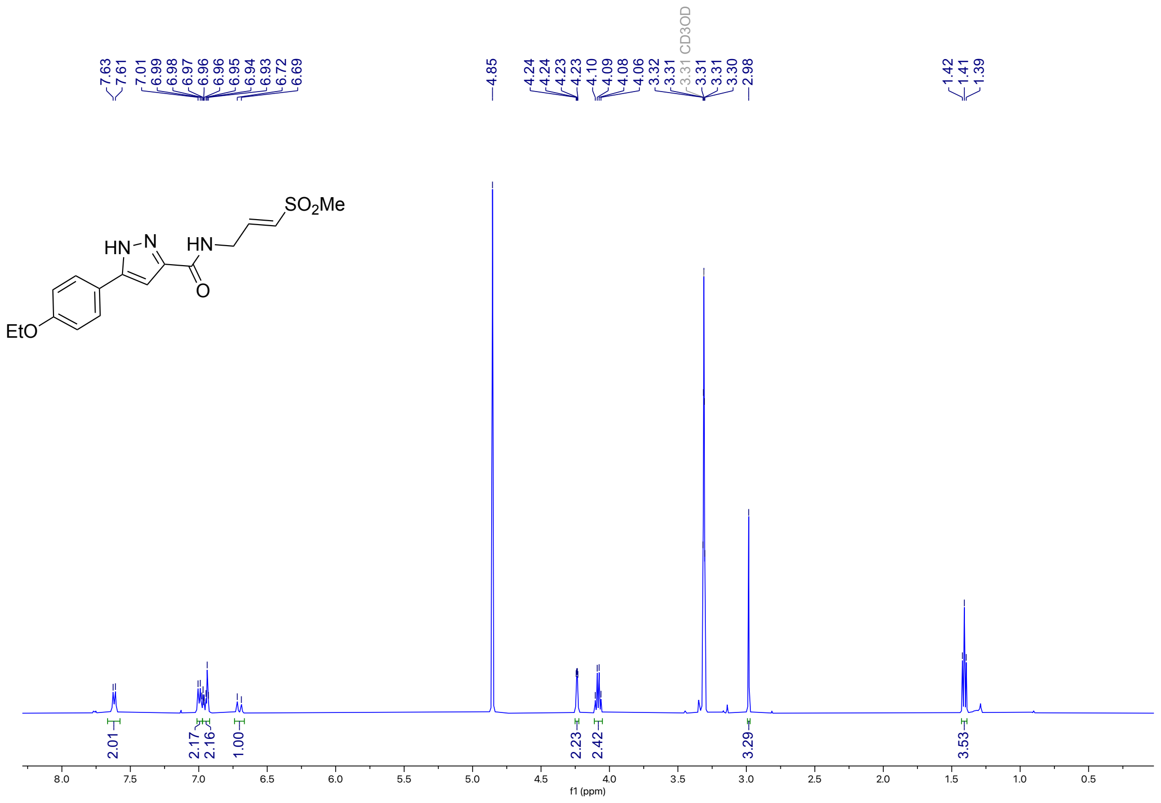

**Figure S25**: $^{13}$C NMR (100 MHz, DMSO-$d_6$) for **1j**

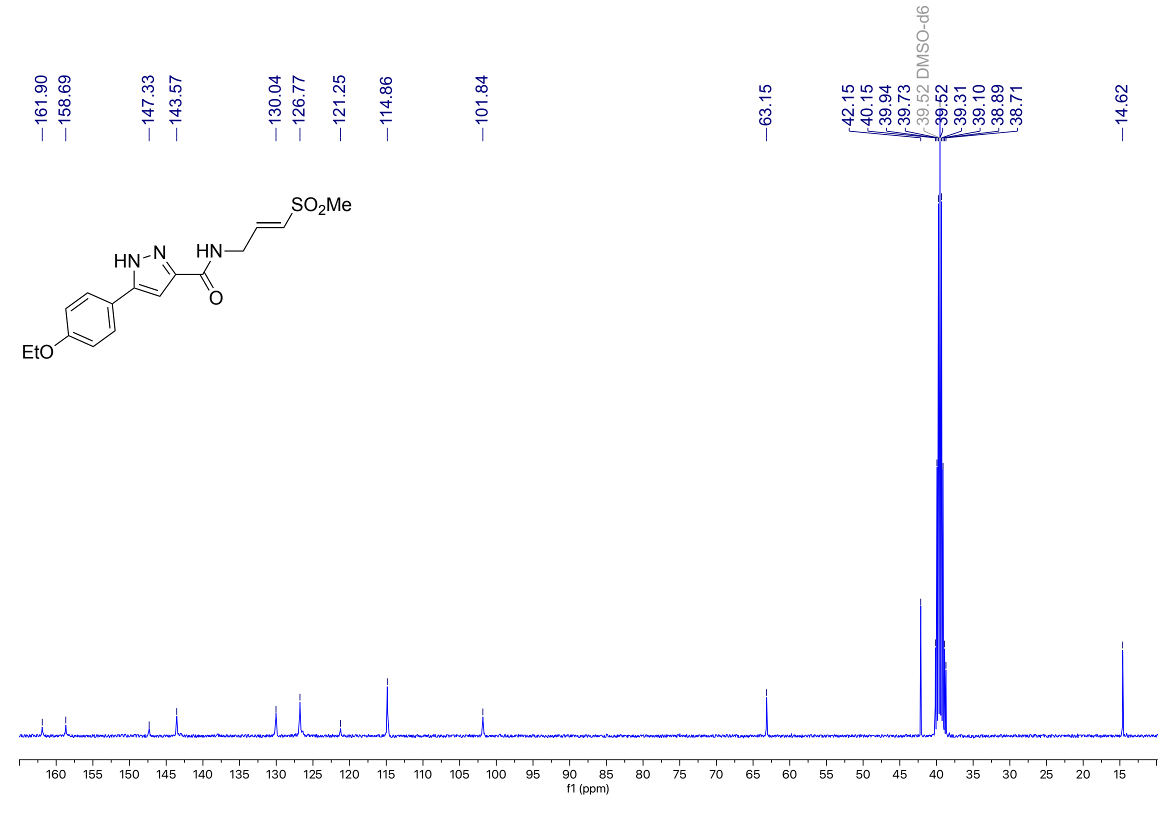

**Figure S26**: $^{1}$H NMR (500 MHz, MeOD) for **1k**

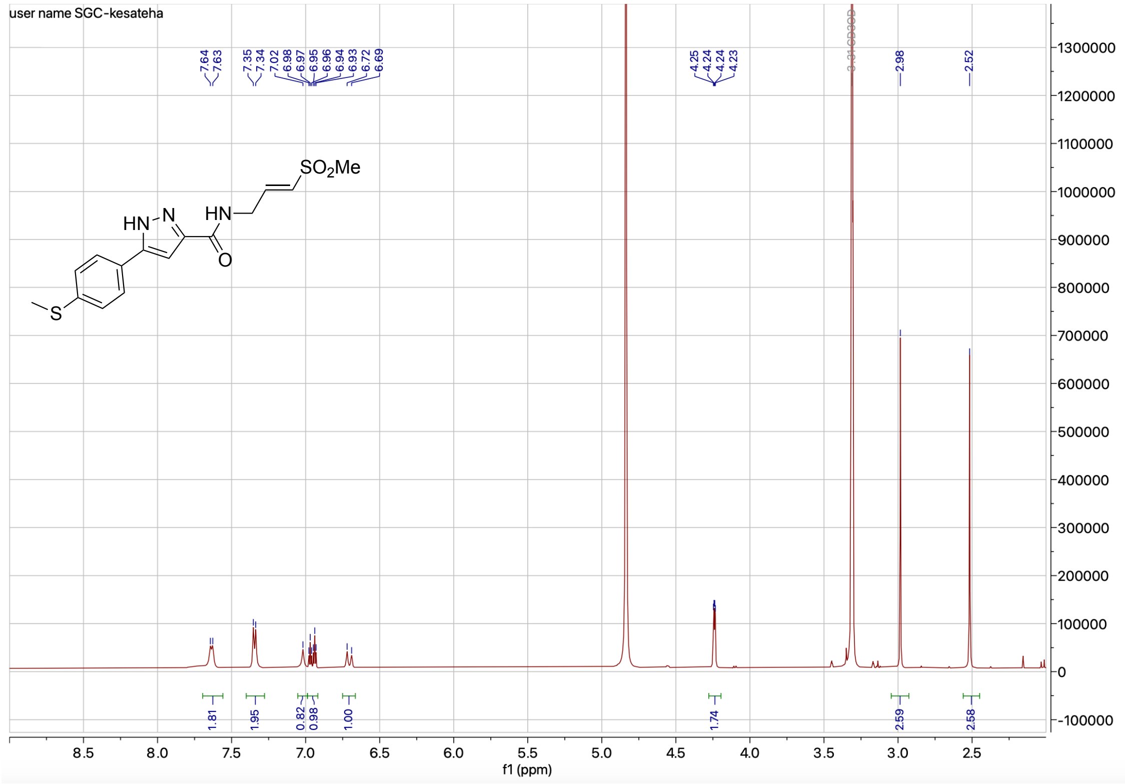


**Figure S27**: $^{13}$C NMR (100 MHz, DMSO-$d_6$) for **1k**

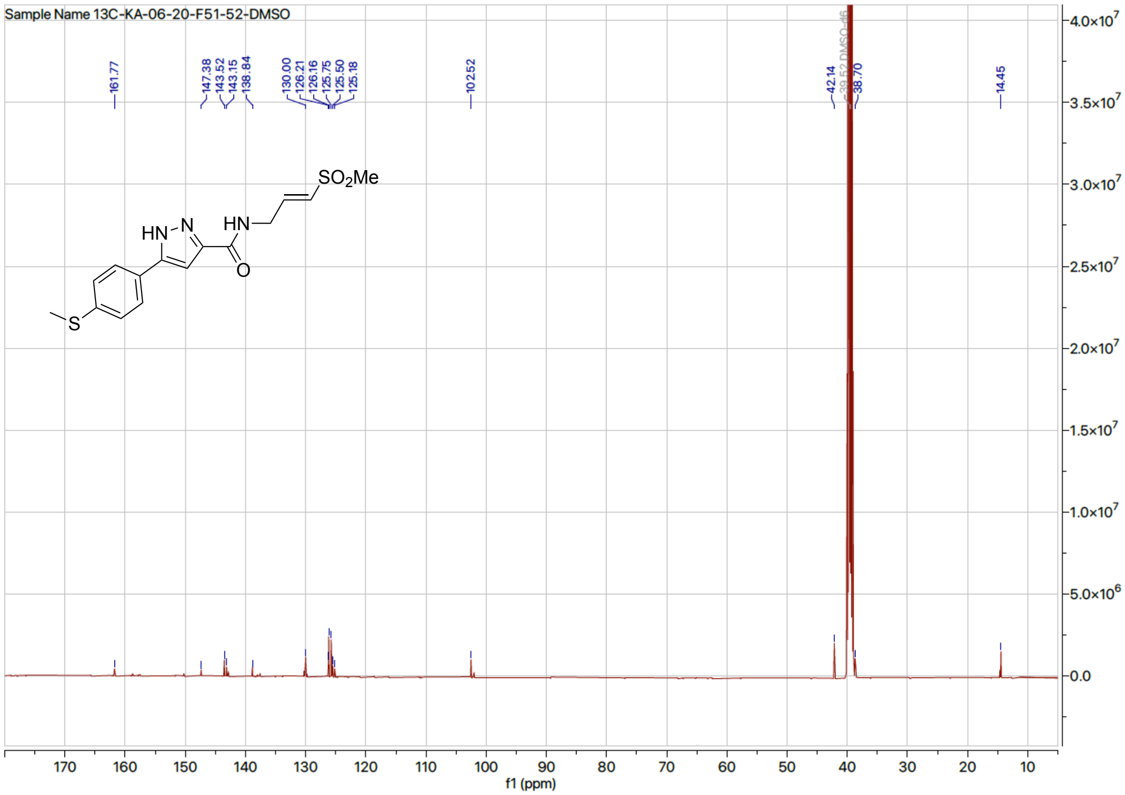

**Figure S28**: $^{1}$H NMR (500 MHz, MeOD) for **1l**

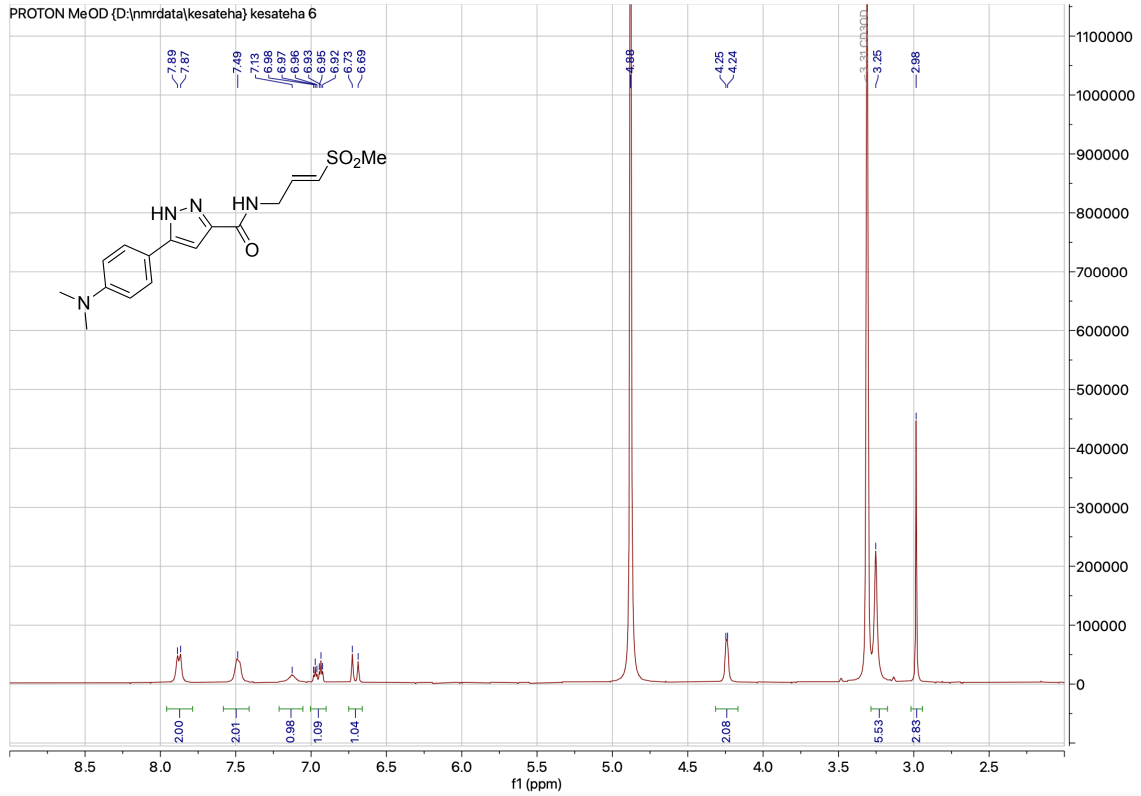


**Figure S29**: $^{13}$C NMR (126 MHz, DMSO-$d_6$) for **1l**

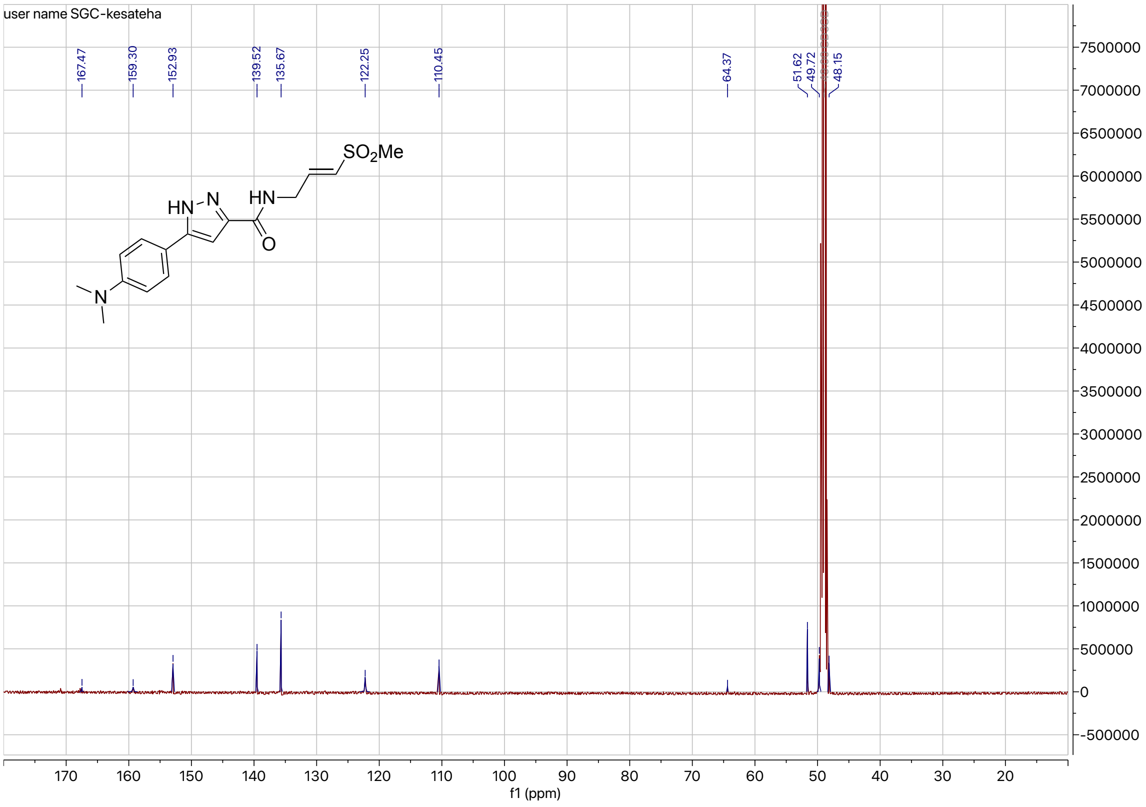

**Figure S30**: $^1$H NMR (400 MHz, DMSO-$d_6$) for **1m**

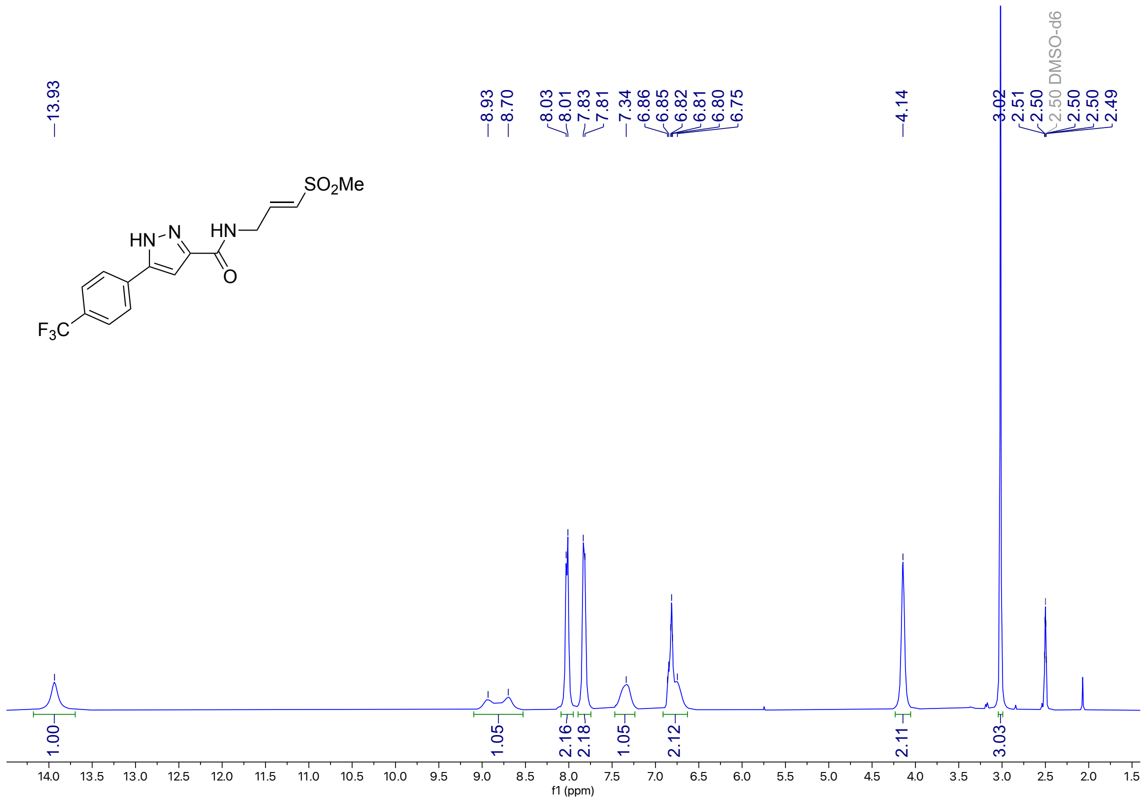

**Figure S31**: $^{13}$C NMR (214 MHz, DMSO-$d_6$) for **1m**

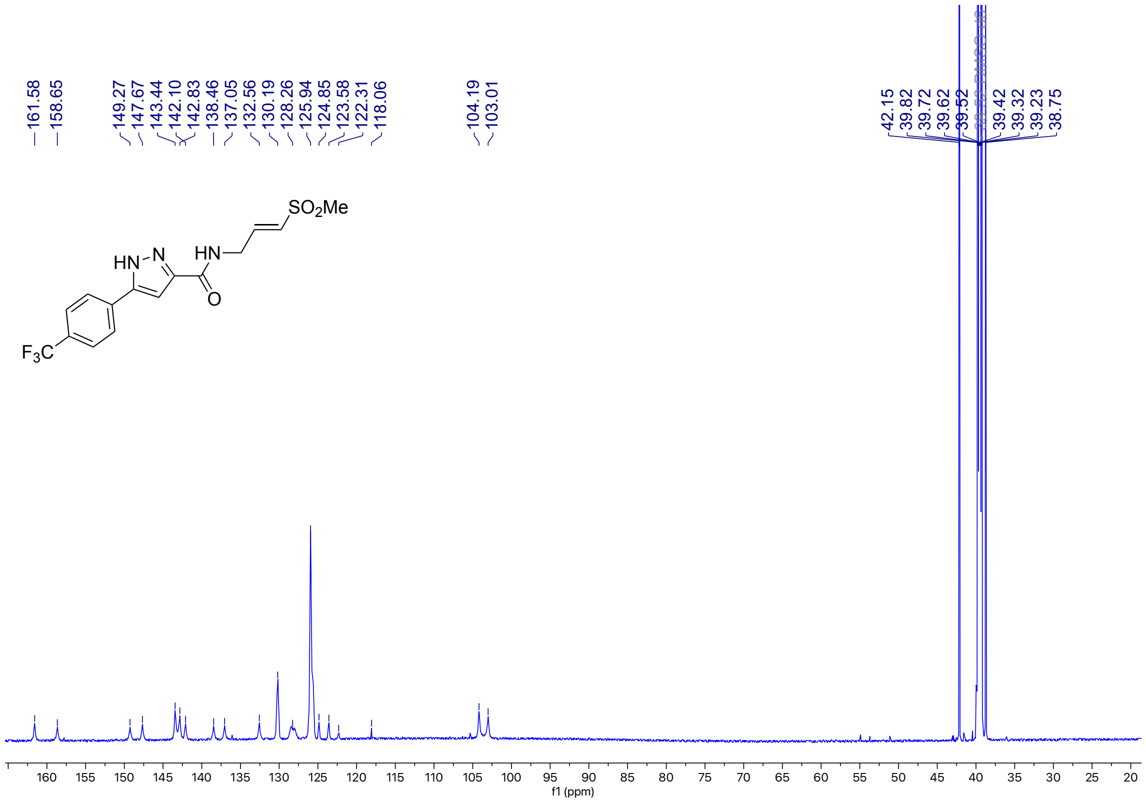

**Figure S32**: $^{1}$H NMR (500 MHz, DMSO-$d_6$) for **1n**

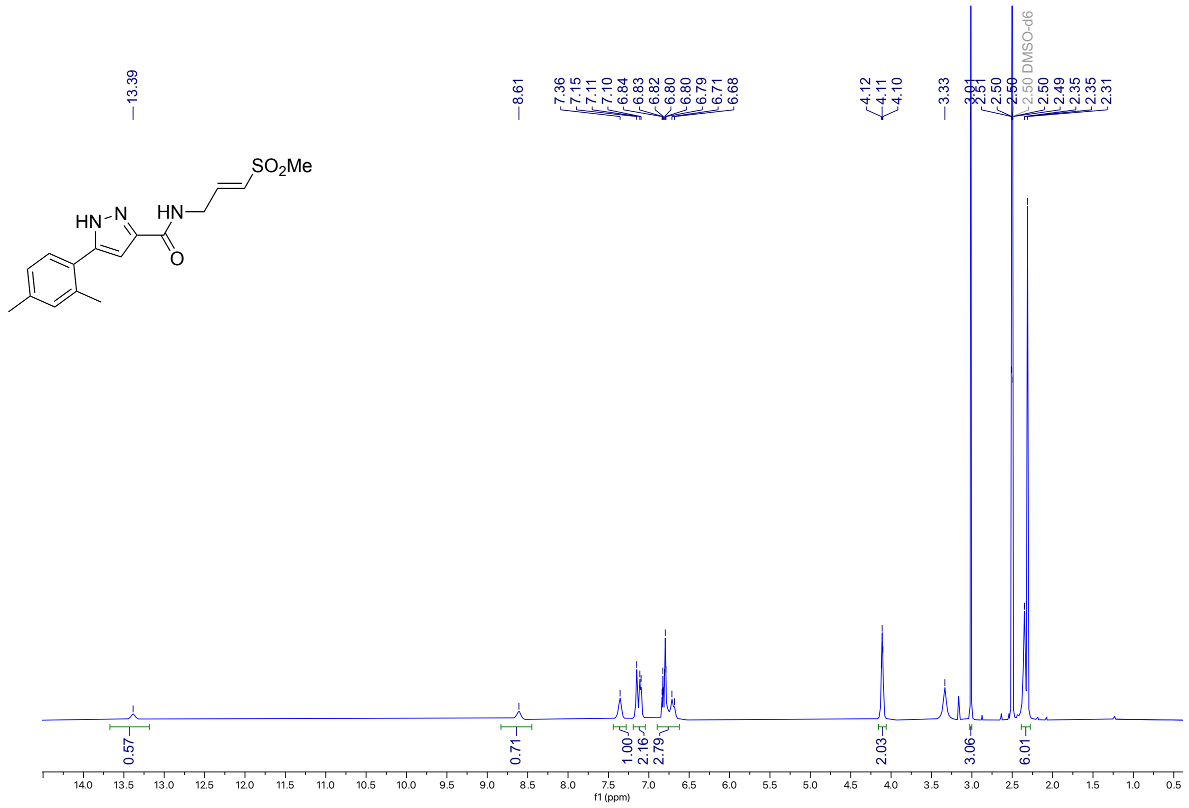

**Figure S33**: $^{13}$C NMR (126 MHz, DMSO-$d_6$) for **1n**

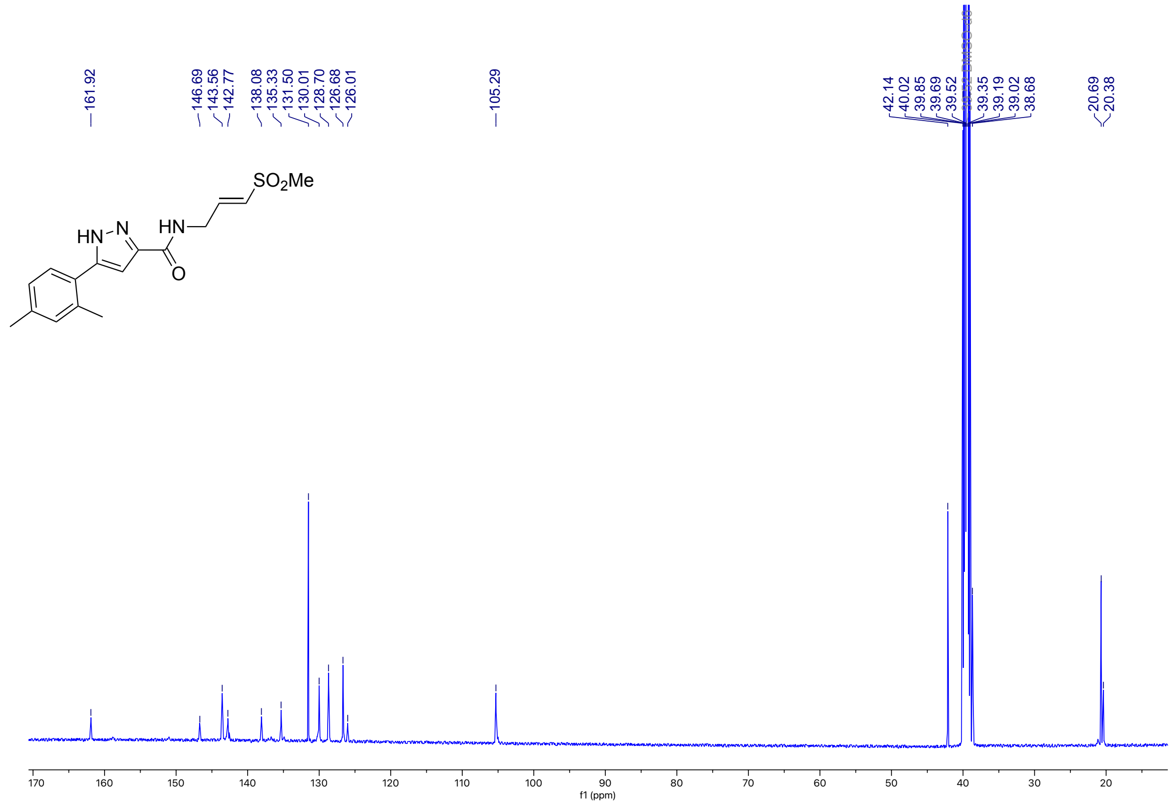

**Figure S34**: $^{1}$H NMR (500 MHz, DMSO-$d_6$) for **1o**

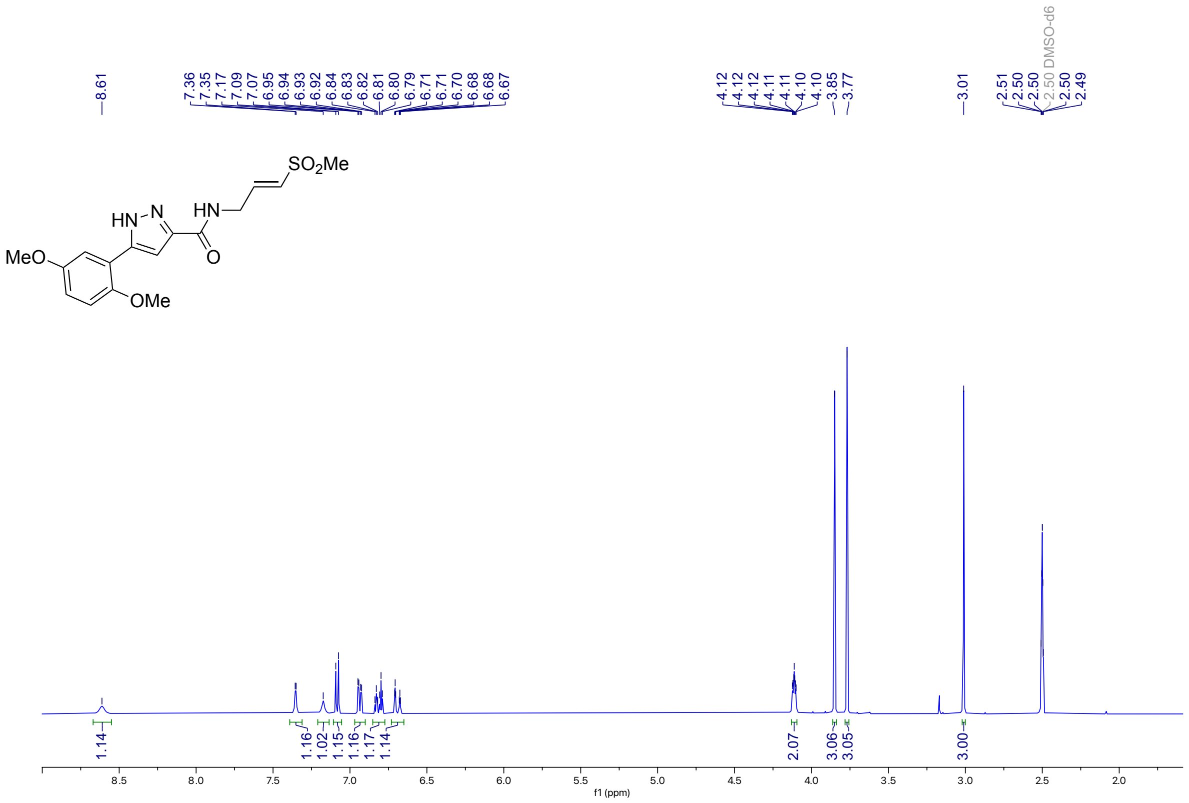

**Figure S35**: $^{13}$C NMR (126 MHz, DMSO-$d_6$) for **1o**

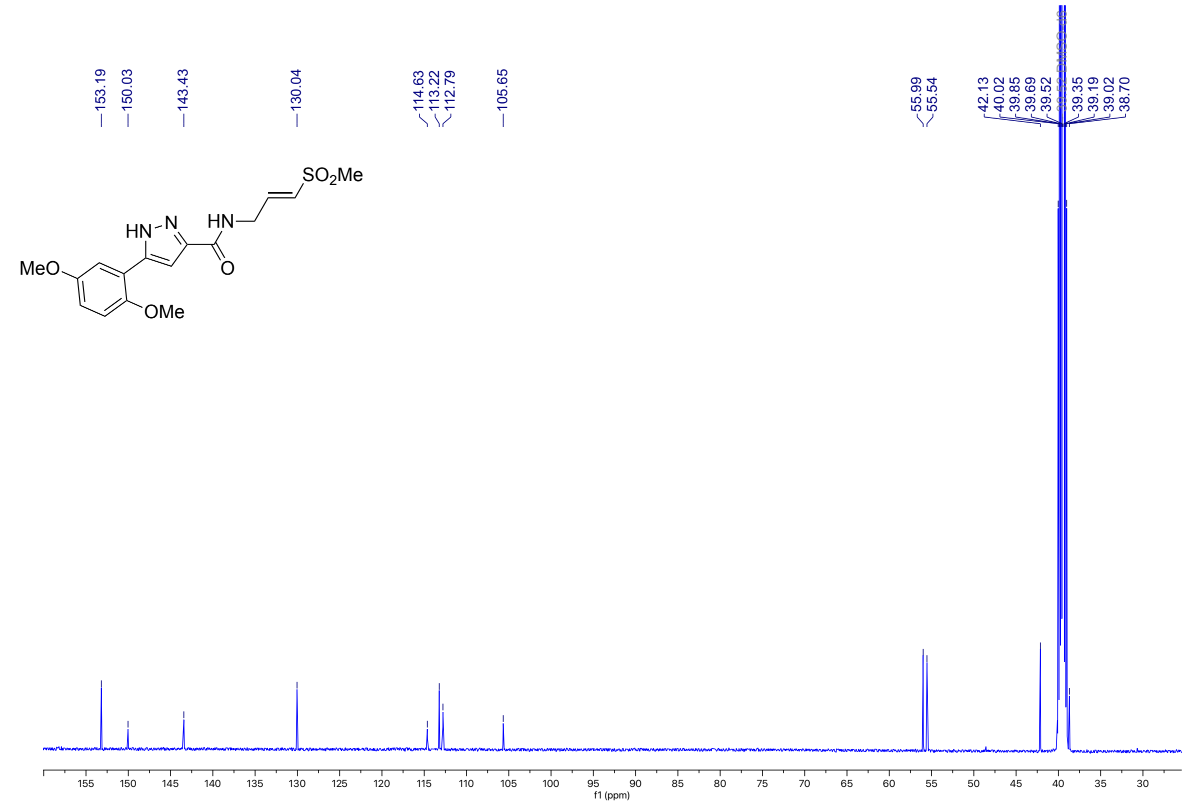

**Figure S36**: $^{1}$H NMR (500 MHz, DMSO-$d_6$) for **1p**

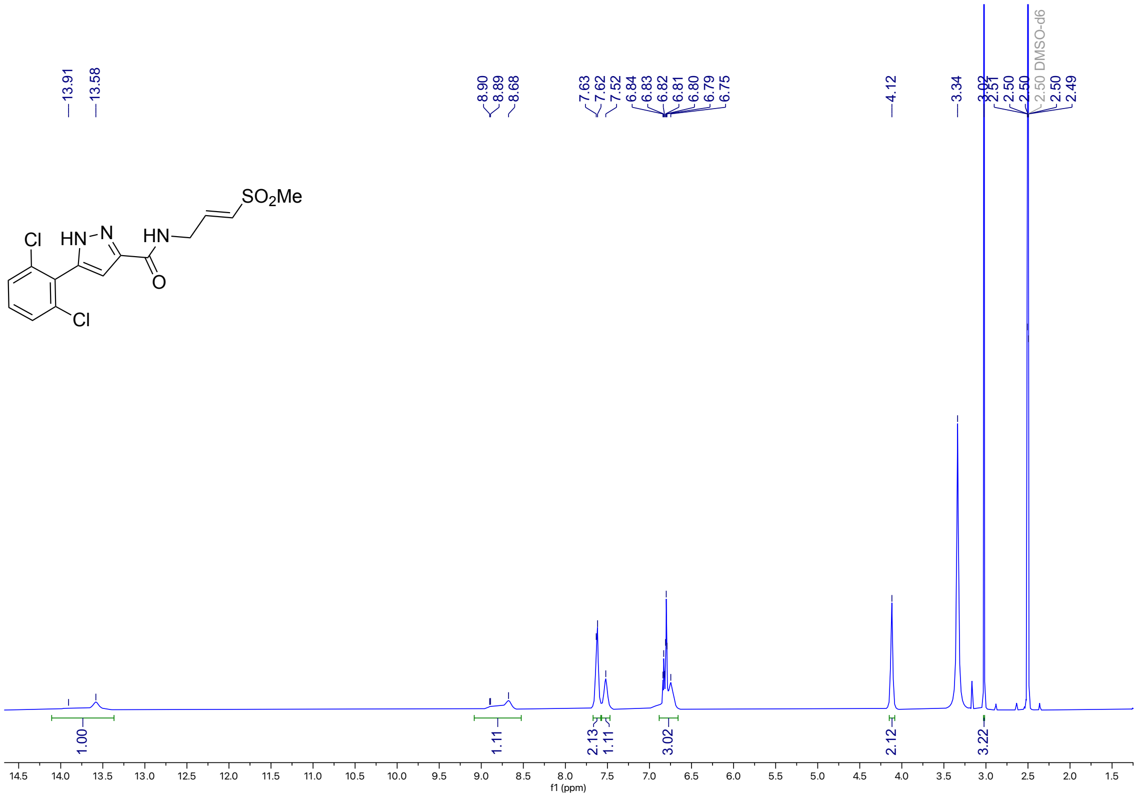

**Figure S37**: $^{13}$C NMR (126 MHz, DMSO-$d_6$) for **1p**

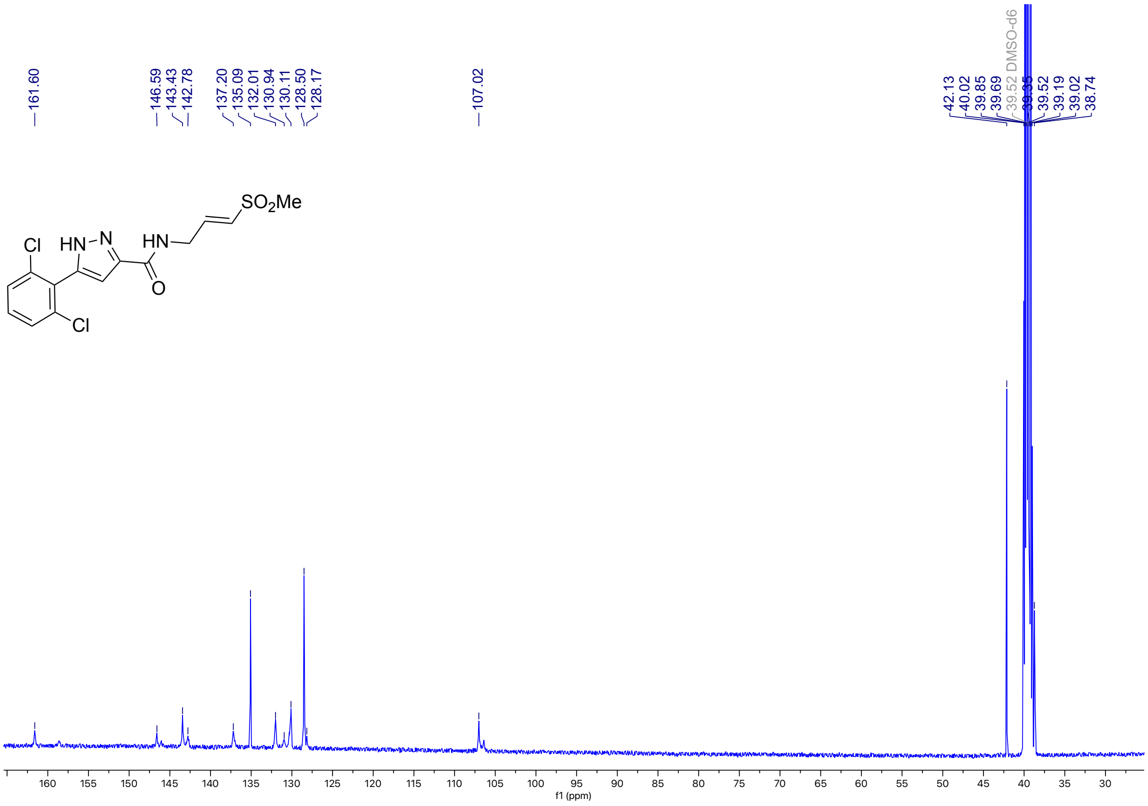

**Figure S38**: $^{1}$H NMR (500 MHz, DMSO-$d_6$) for **1q**

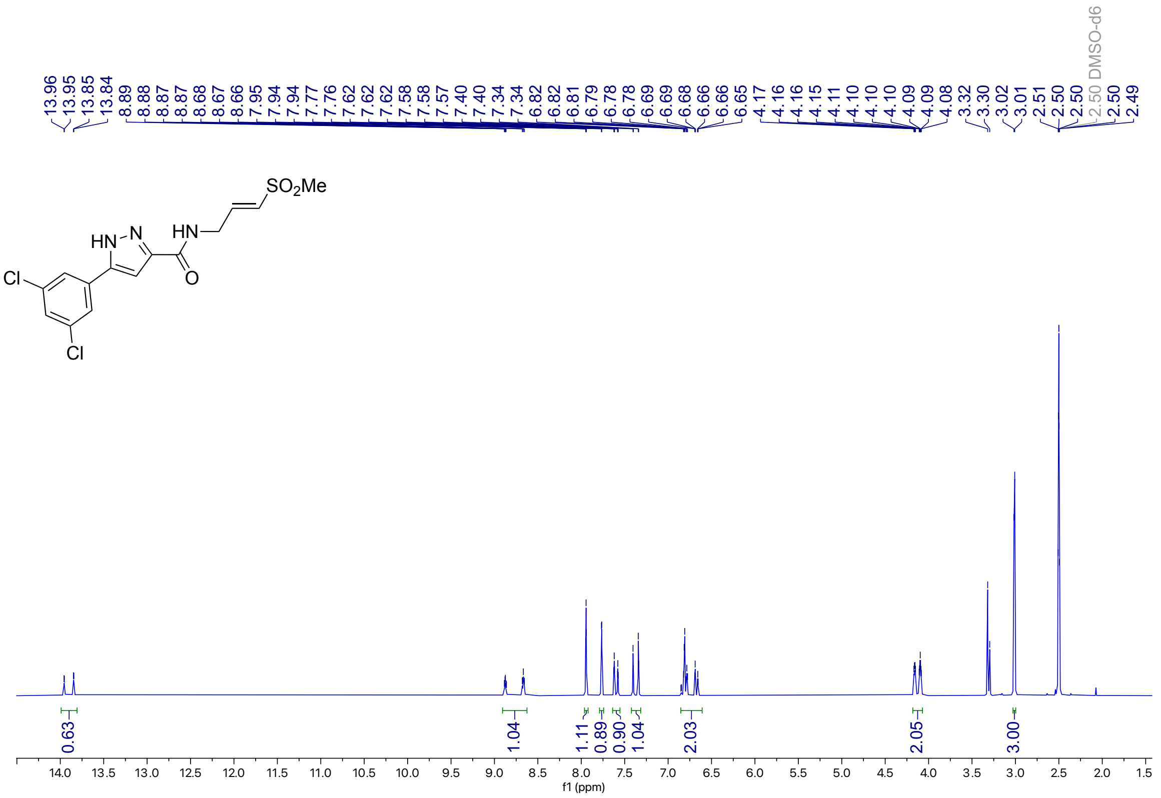

**Figure S39**: $^{13}$C NMR (126 MHz, DMSO-$d_6$) for **1q**

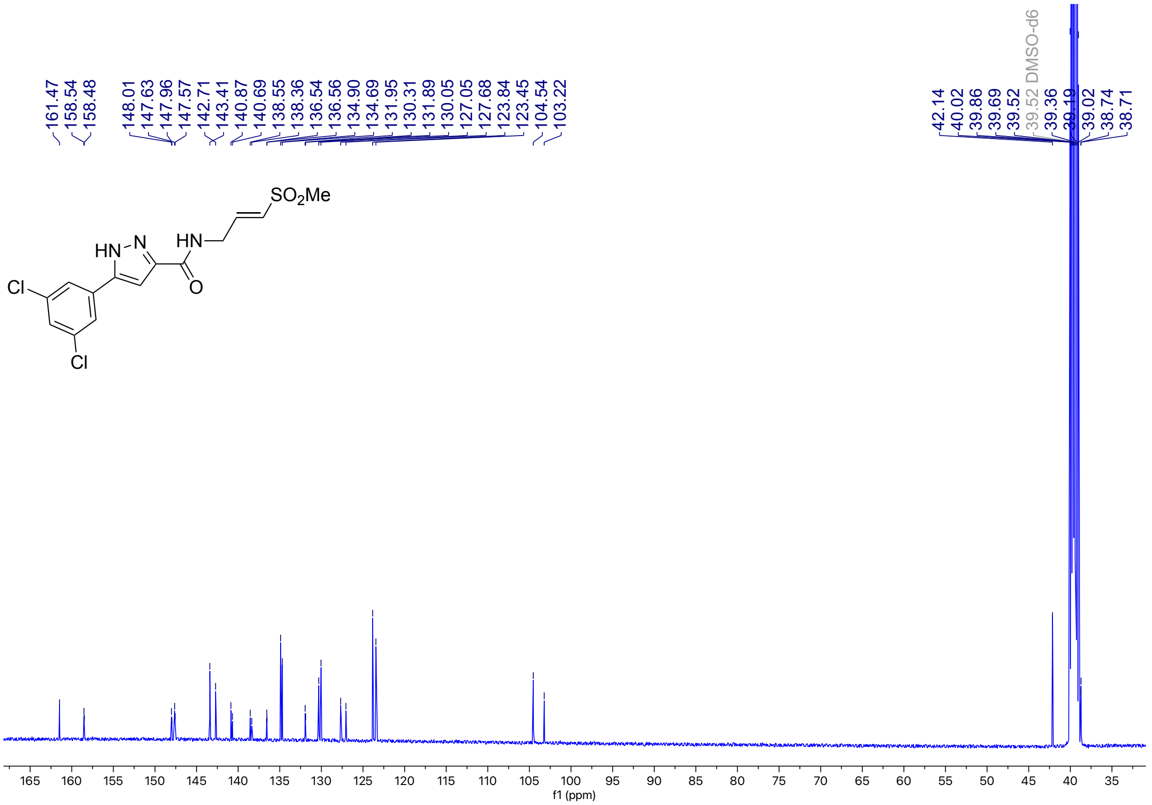

**Figure S40**: $^{1}$H NMR (500 MHz, DMSO-$d_6$) for **4c**

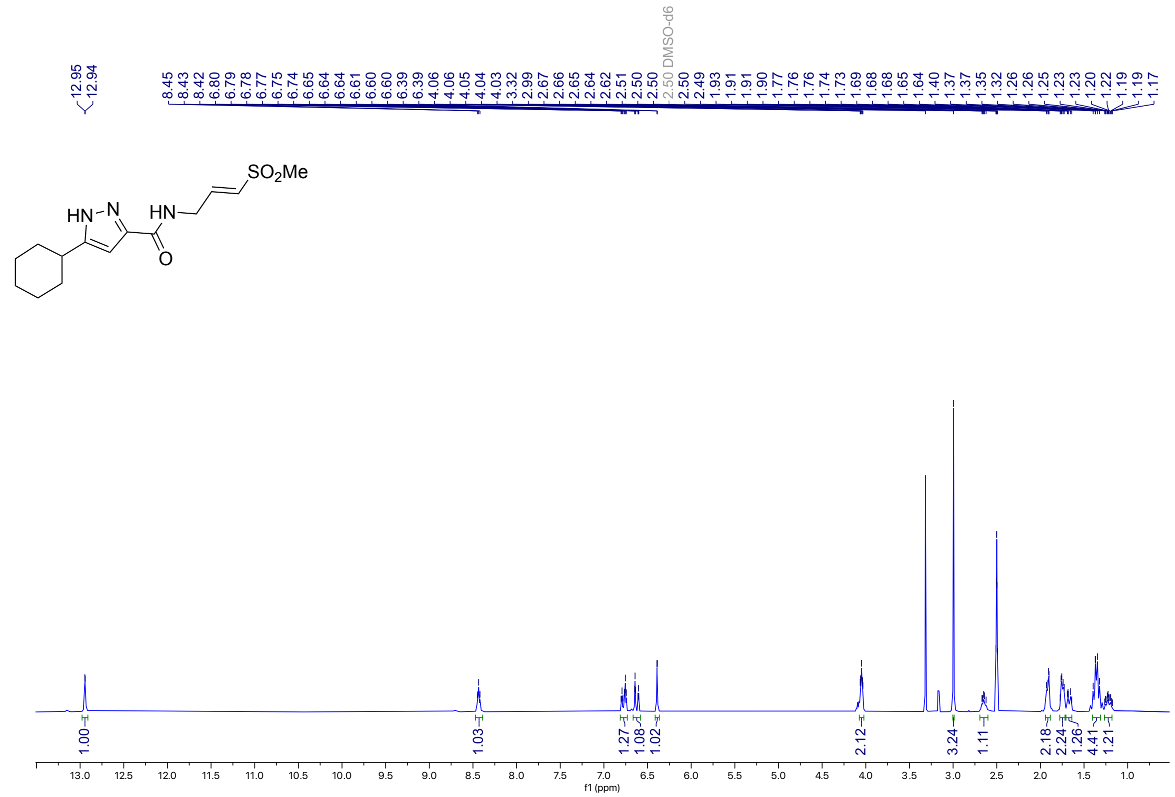


**Figure S41**: $^{13}$C NMR (126 MHz, DMSO-$d_6$) for **4c**

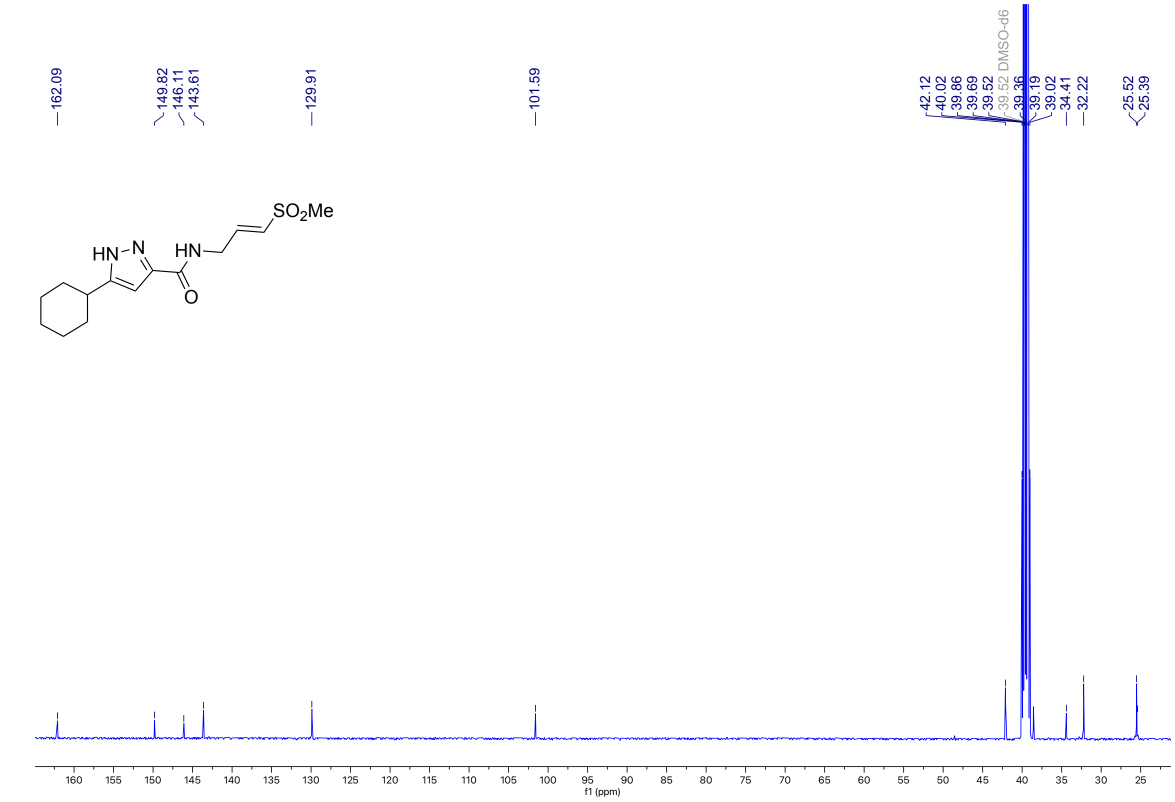

**Figure S42**: $^{1}$H NMR (400 MHz, DMSO-$d_6$) for **4d**

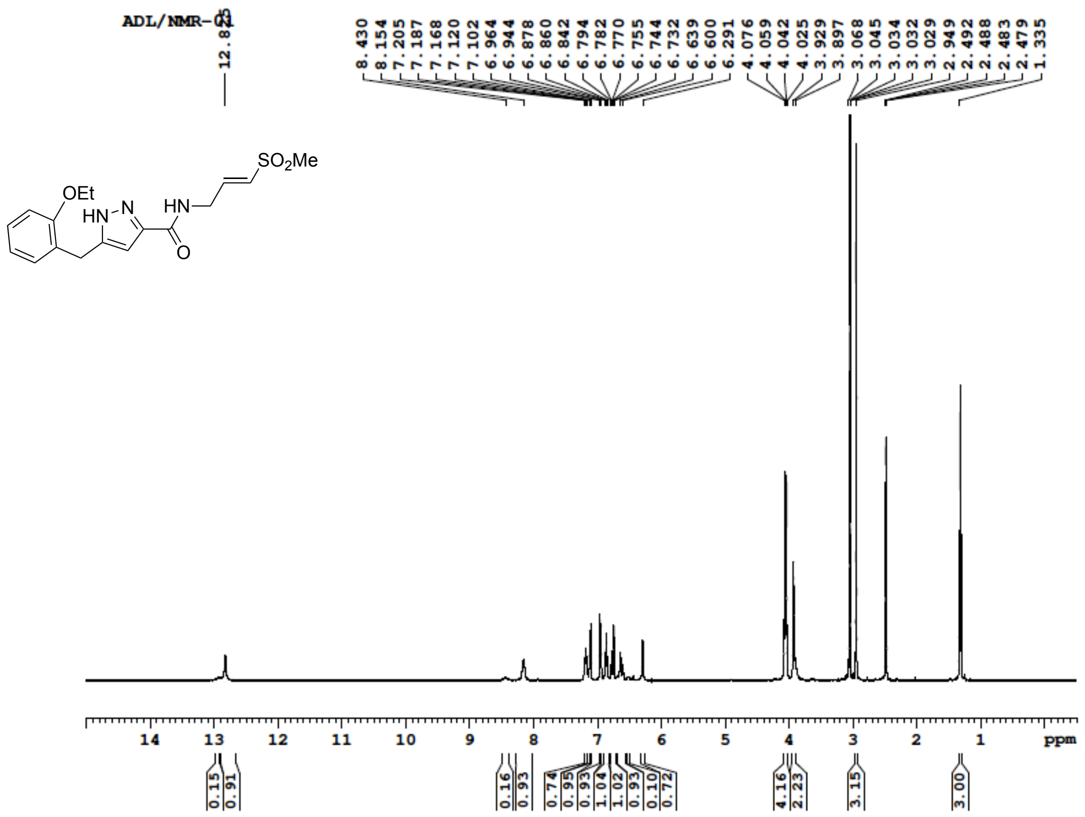


**Figure S43**: $^{13}$C NMR (100 MHz, DMSO-$d_6$) for **4d**

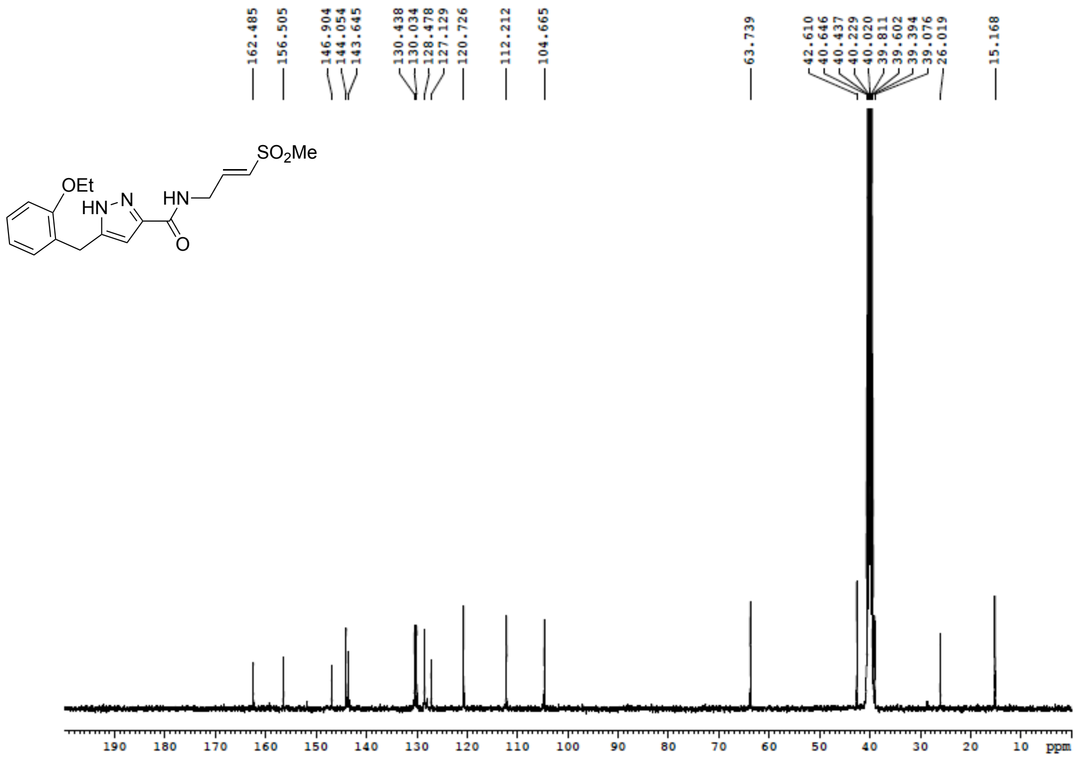

**Figure S44**: $^{1}$H NMR (400 MHz, DMSO-$d_6$) for **4e**

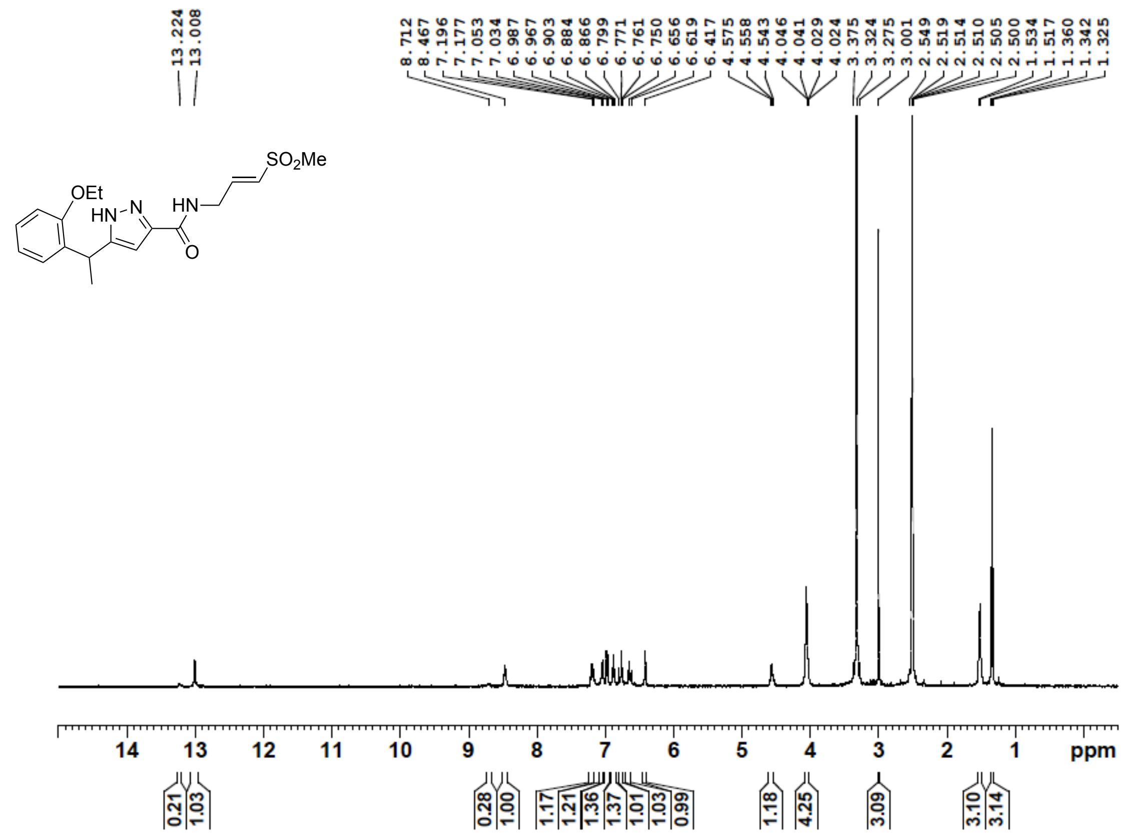

**Figure S45**: $^{13}$C NMR (100 MHz, DMSO-$d_6$) for **4e**

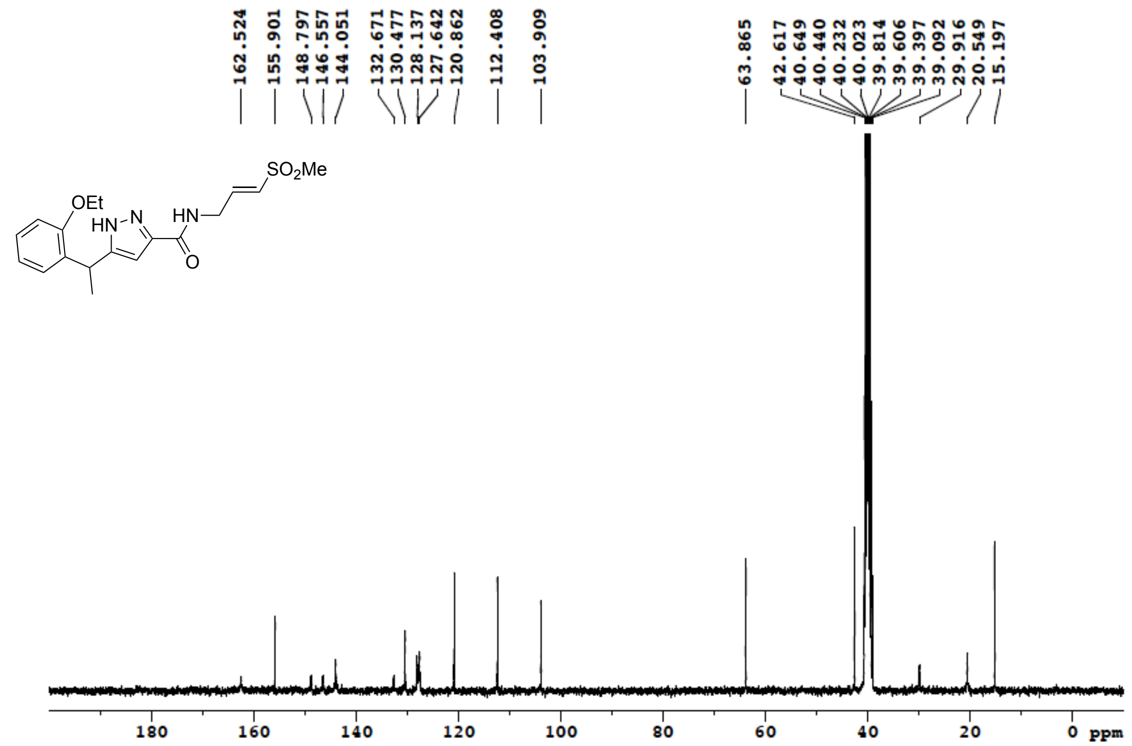

**Figure S46**: $^{1}H$ NMR (400 MHz, DMSO-$d_6$) for **4g**

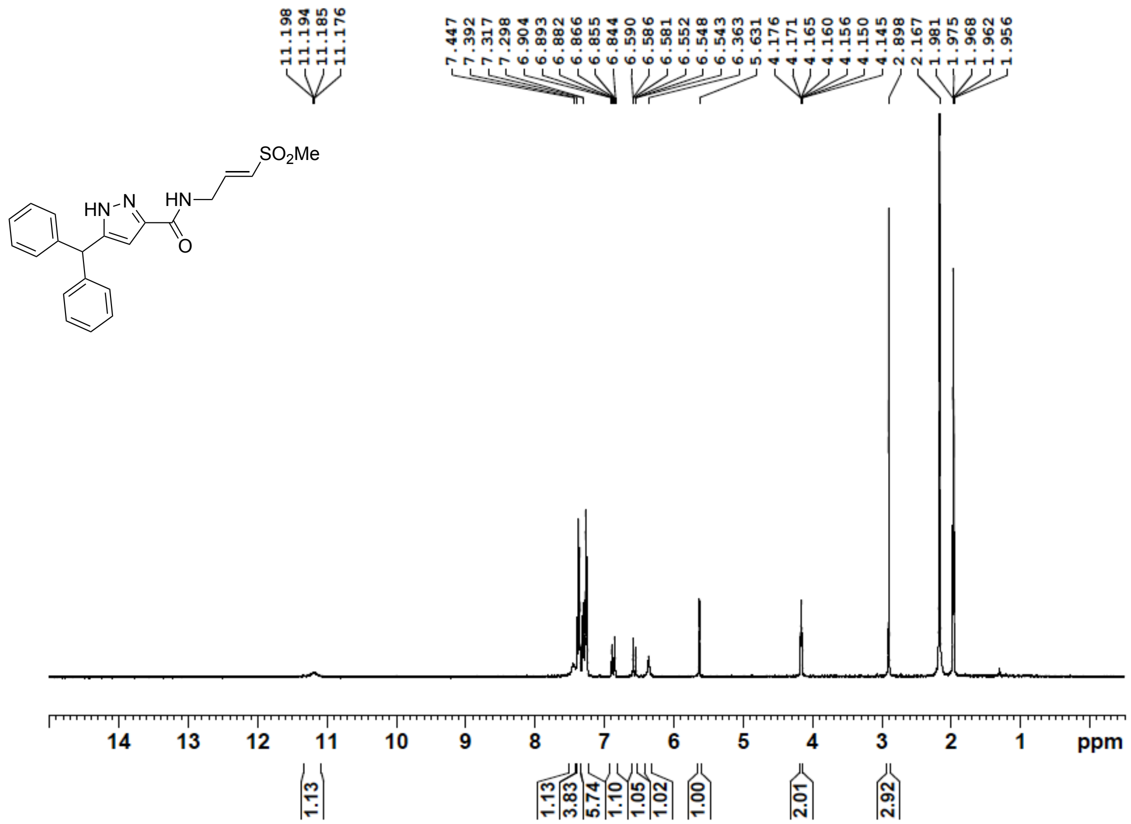


**Figure S47**: $^{13}C$ NMR (100 MHz, DMSO-$d_6$) for **4g**

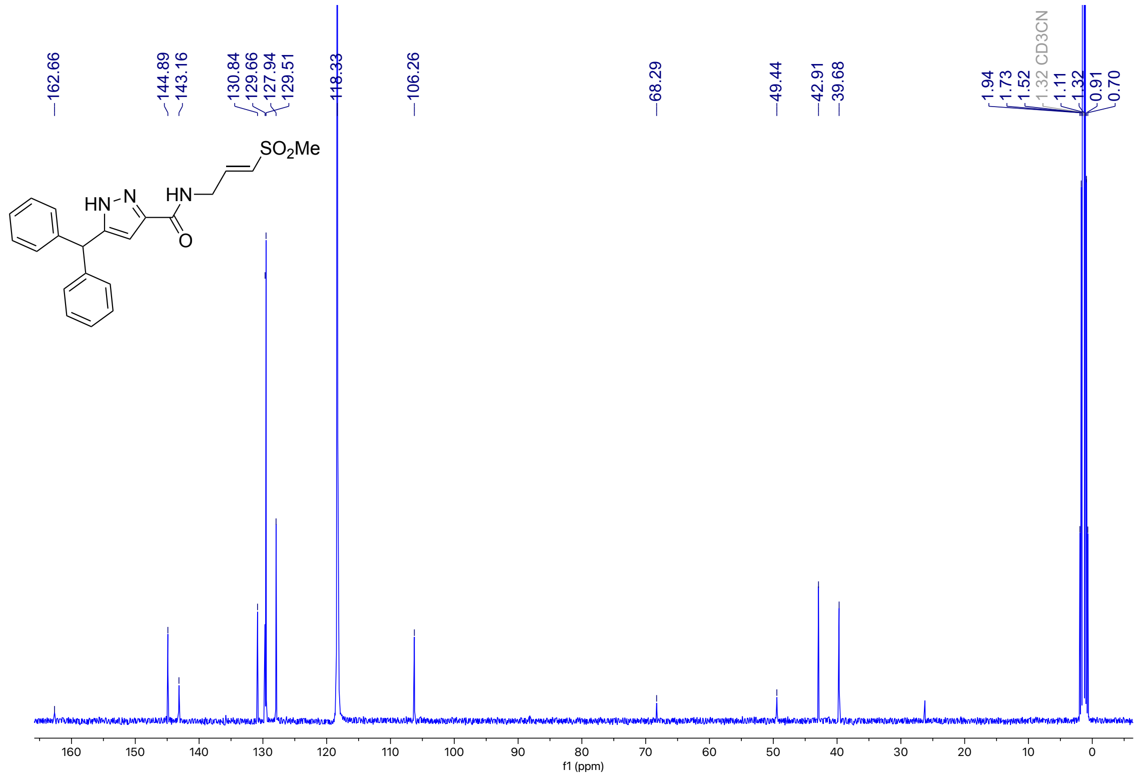

**Figure S48**: $^{1}$H NMR (500 MHz, DMSO-$d_6$) for **4h**

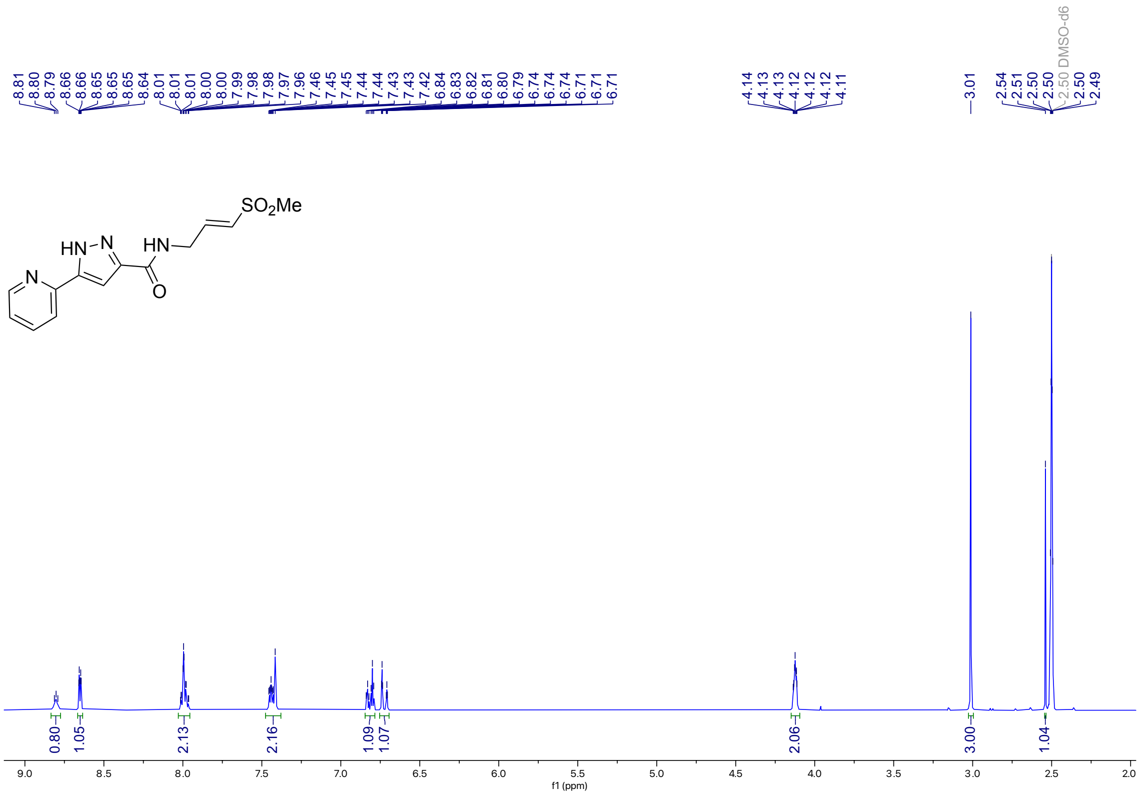

**Figure S49**: $^{13}$C NMR (126 MHz, DMSO-$d_6$) for **4h**

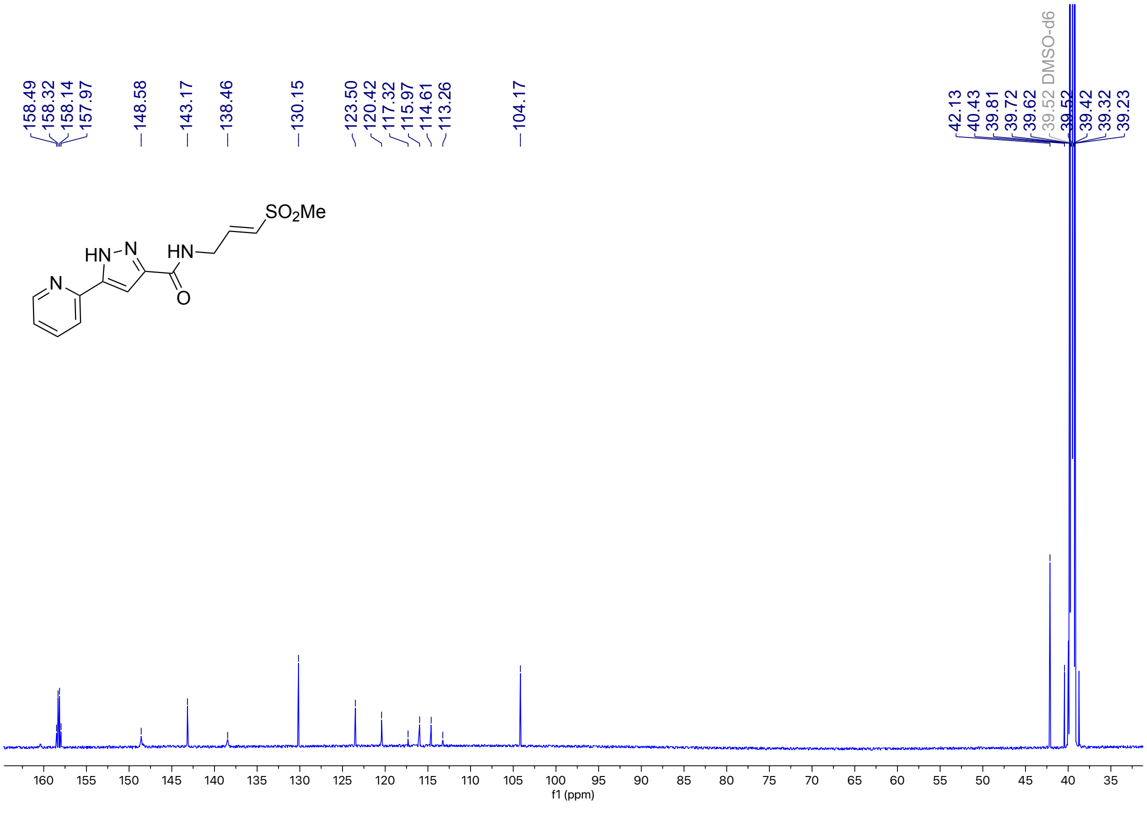

**Figure S50**: [1]H NMR (500 MHz, DMSO-$d_6$) for **4i**

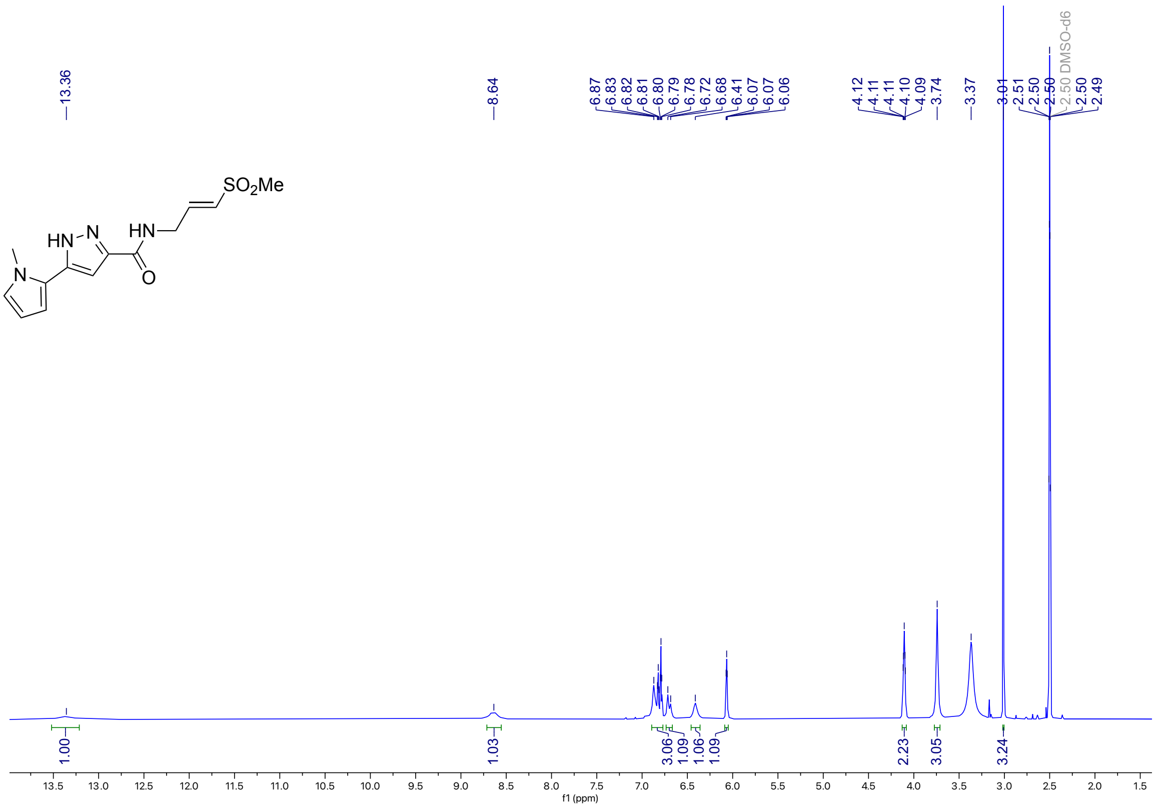


**Figure S51**: [13]C NMR (214 MHz, DMSO-$d_6$) for **4i**

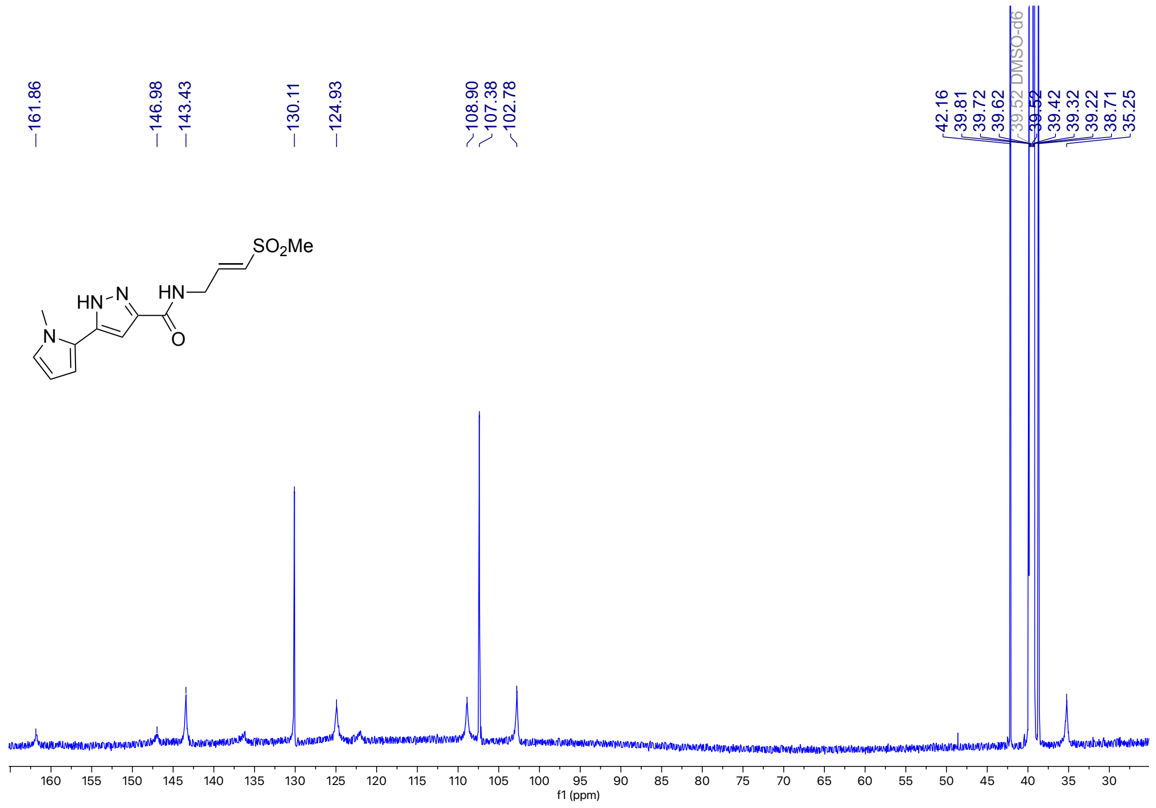

**Figure S52**: $^{1}$H NMR (400 MHz, DMSO-$d_6$) for **5**

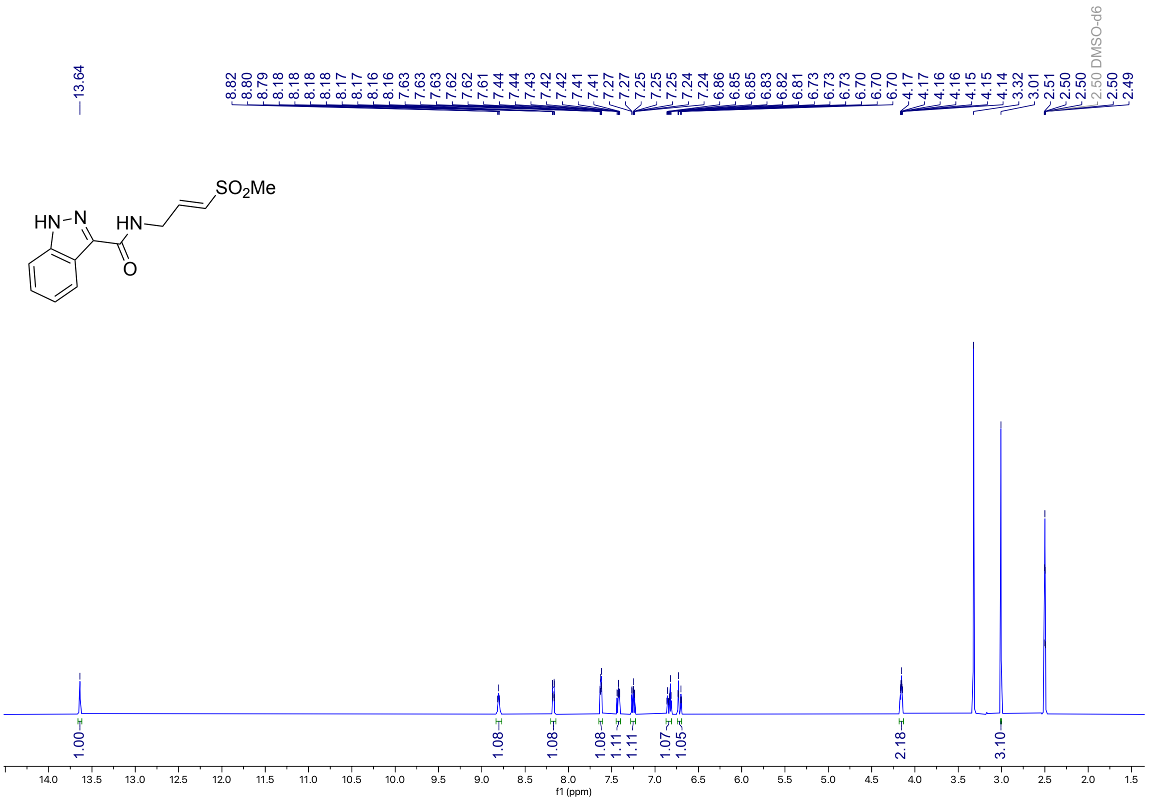

**Figure S53**: $^{13}$C NMR (100 MHz, DMSO-$d_6$) for **5**

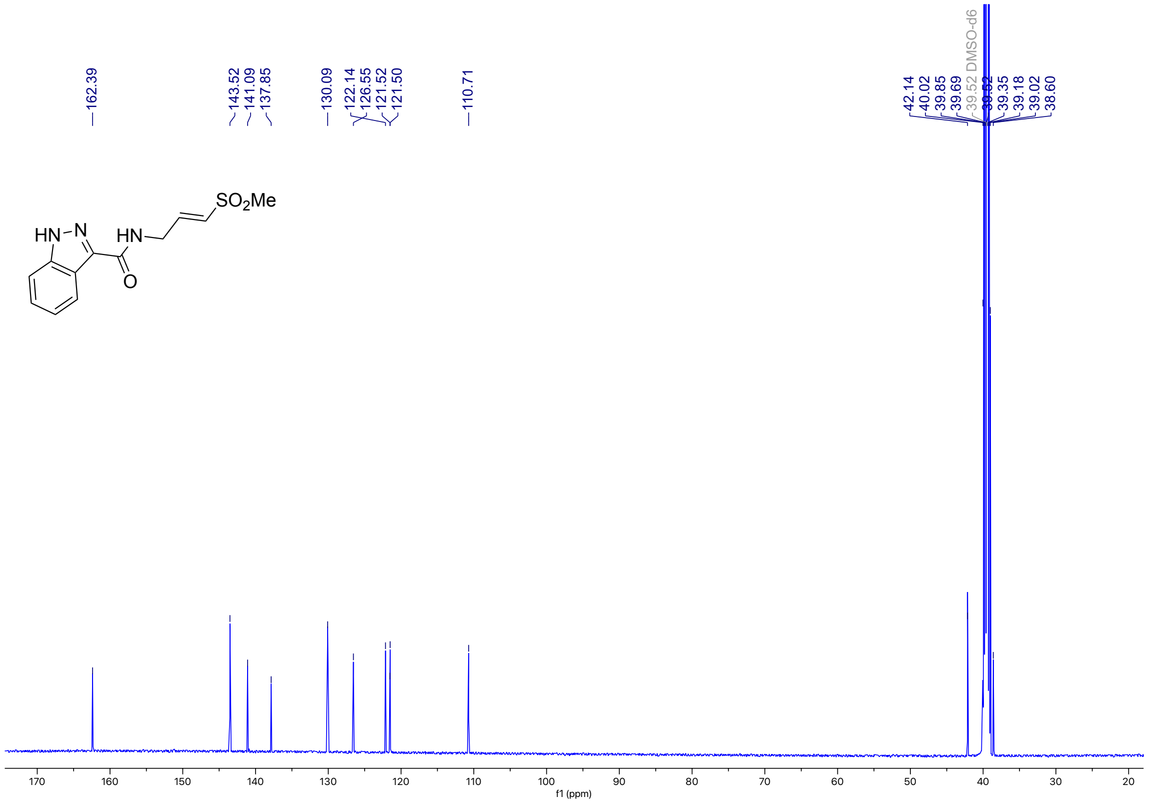

**Figure S54**: [1]H NMR (500 MHz, DMSO-$d_6$) for **6**

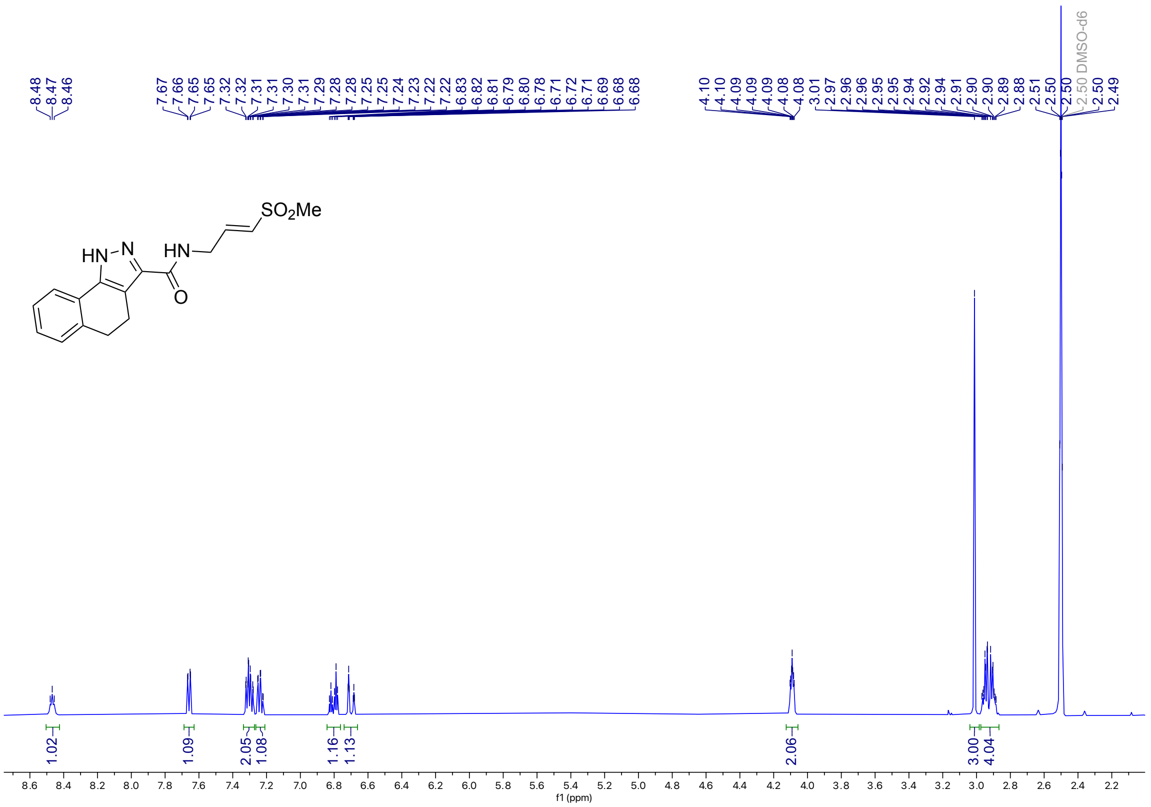

**Figure S55**: [13]C NMR (214 MHz, DMSO-$d_6$) for **6**

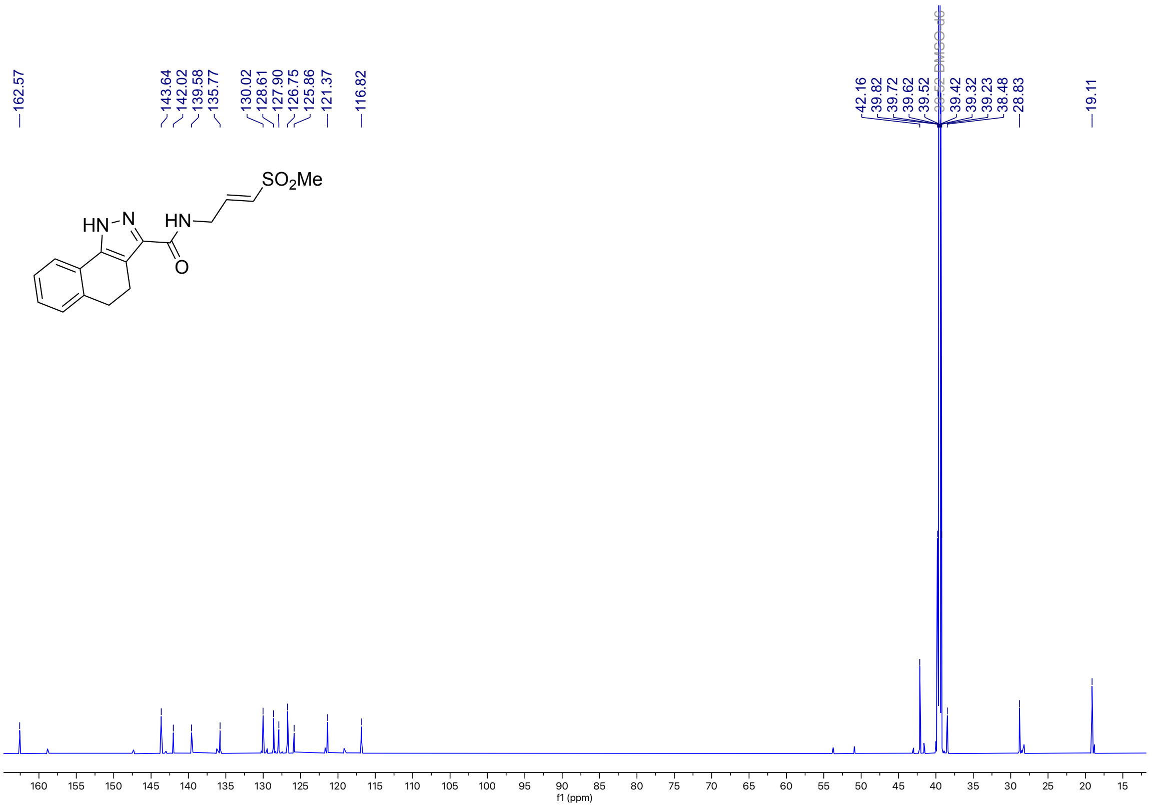

**Figure S56**: $^{1}$H NMR (500 MHz, DMSO-$d_6$) for **7a**

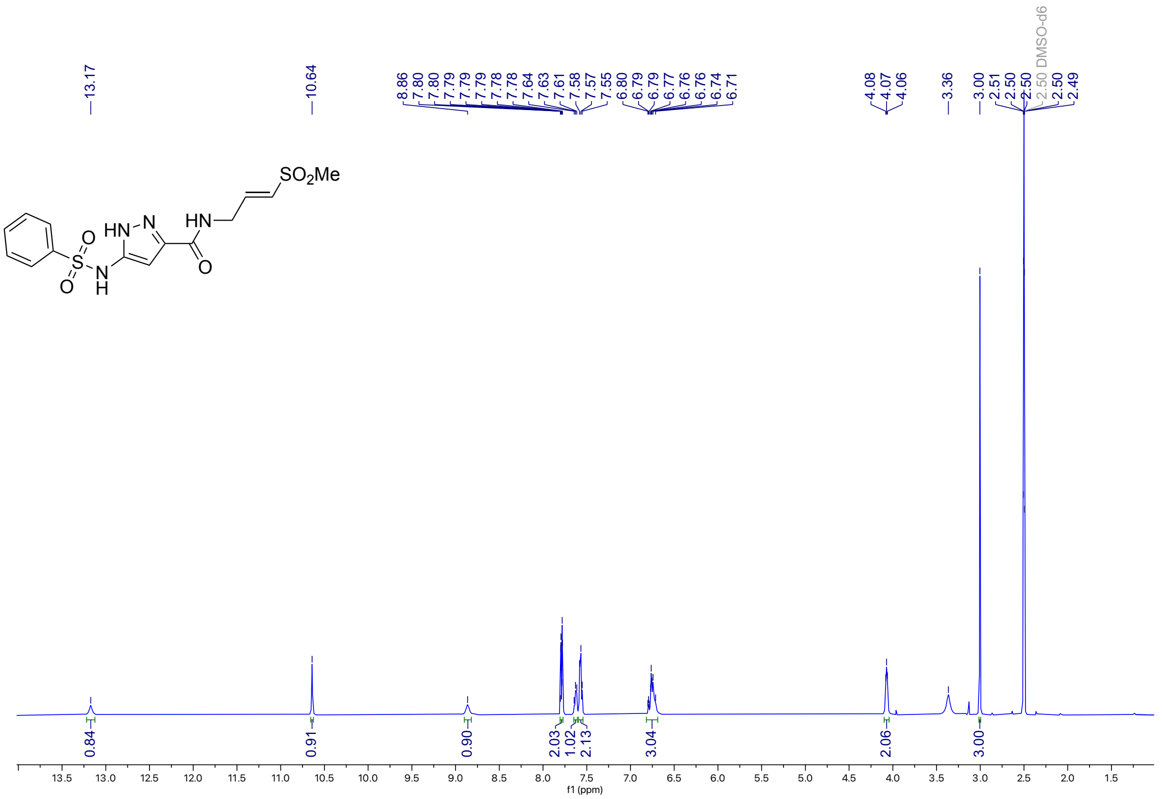

**Figure S57**: $^{13}$C NMR (126 MHz, DMSO-$d_6$) for **7a**

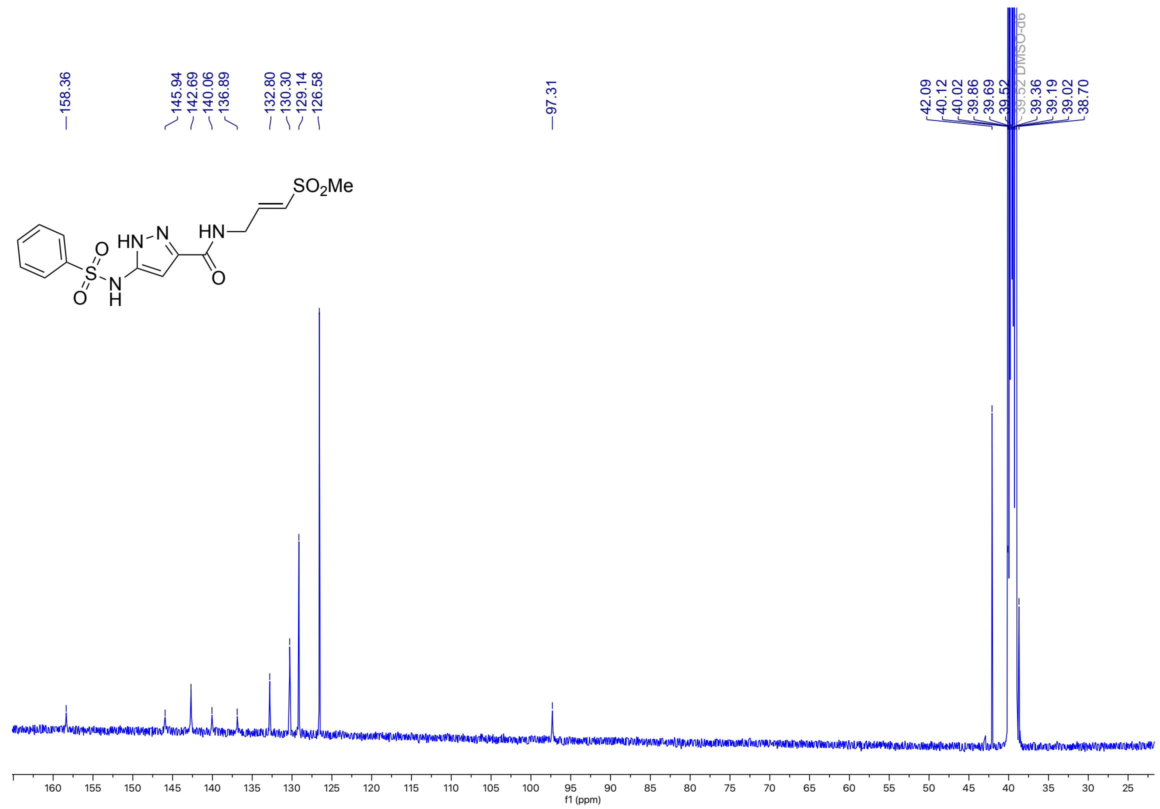

**Figure S58**: $^{1}$H NMR (400 MHz, DMSO-$d_6$) for **7b**

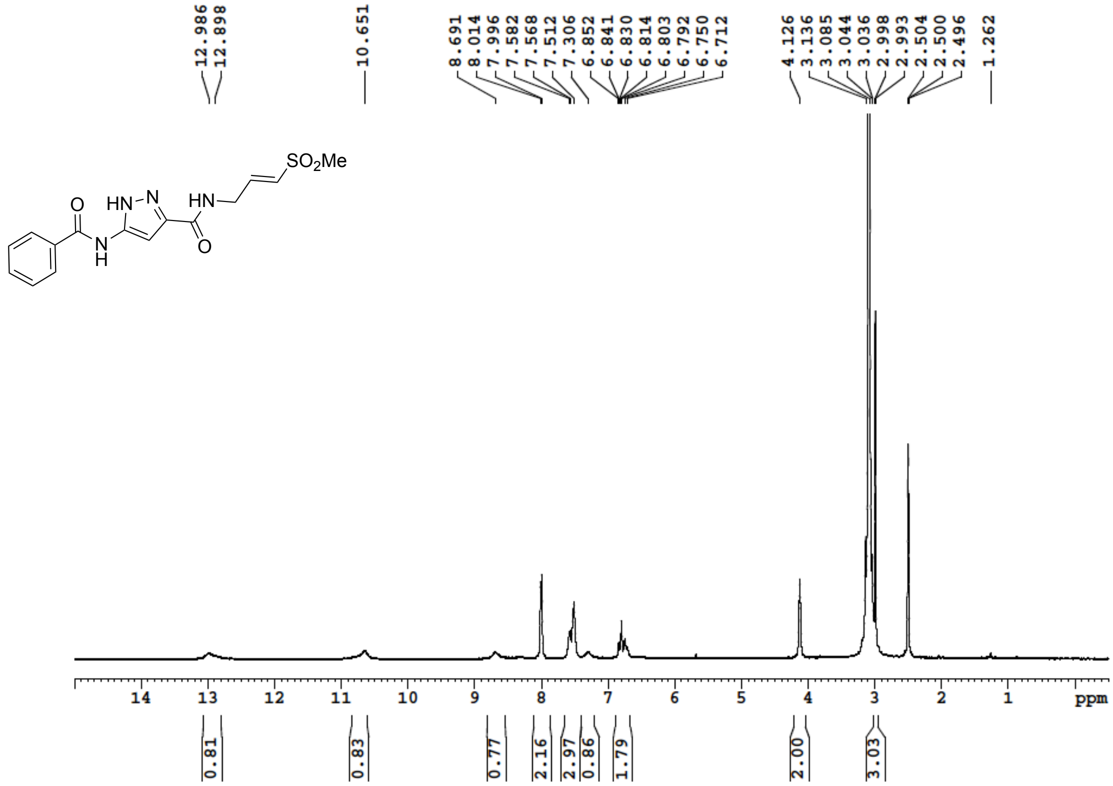


**Figure S59**: $^{13}$C NMR (100 MHz, DMSO-$d_6$) for **7b**

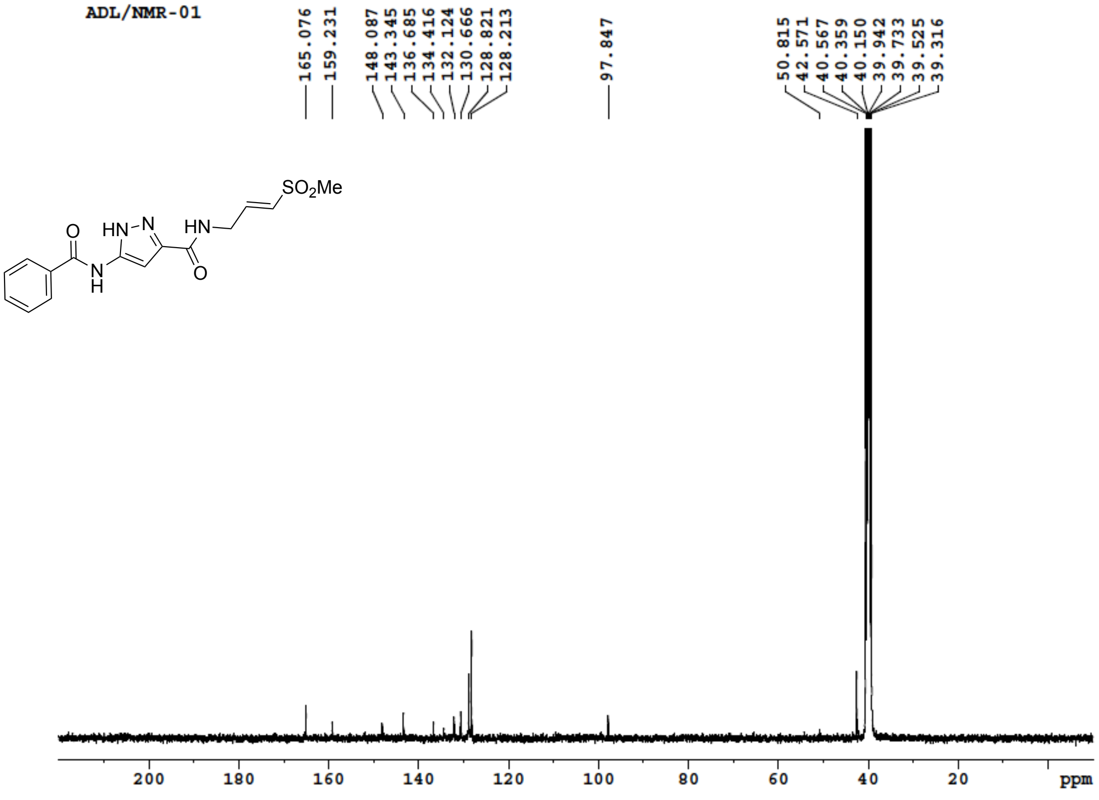

**Figure S60**: $^{1}$H NMR (400 MHz, DMSO-$d_6$) for **7c**

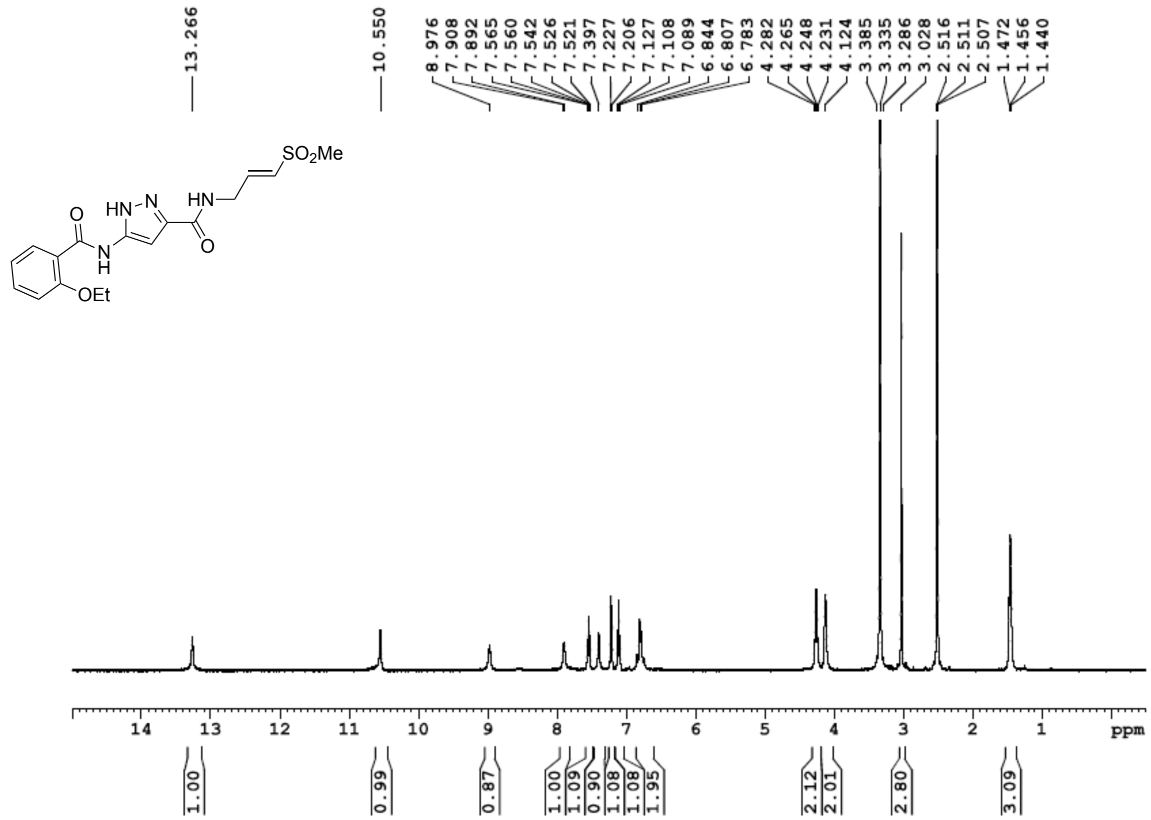

**Figure S61**: $^{13}$C NMR (100 MHz, DMSO-$d_6$) for **7c**

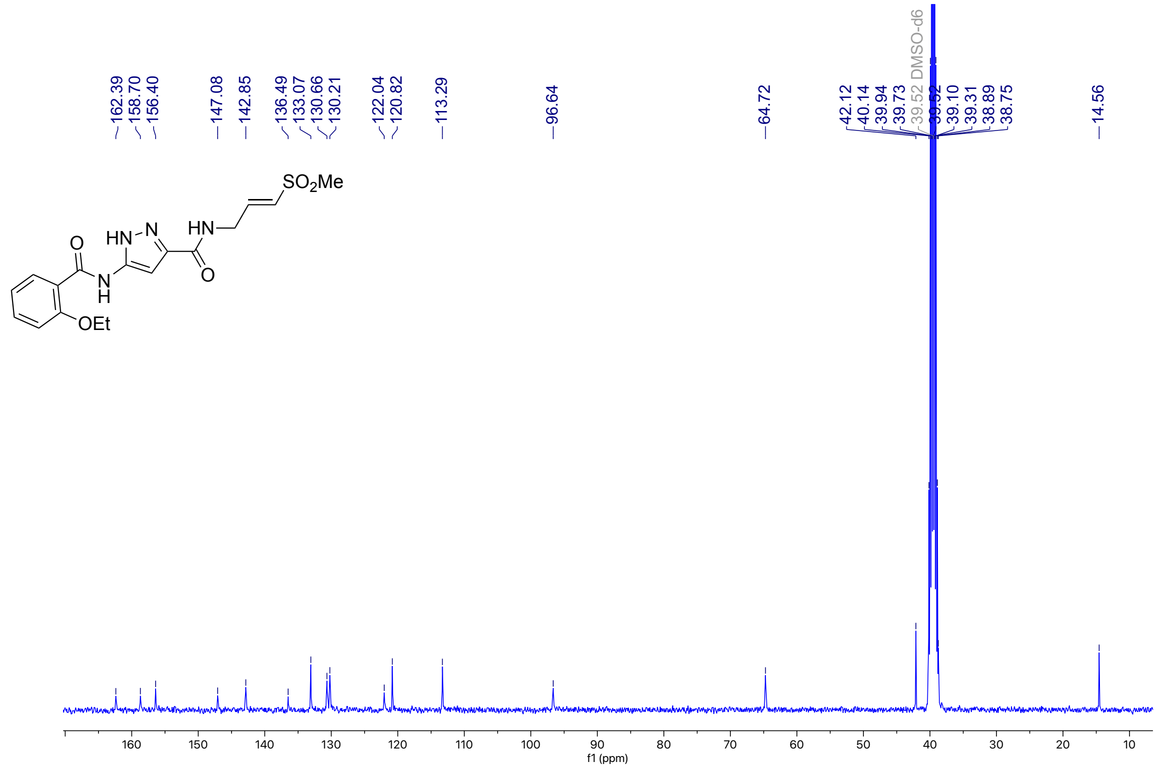

**Figure S62**: $^1$H NMR (400 MHz, DMSO-$d_6$) for **7d**

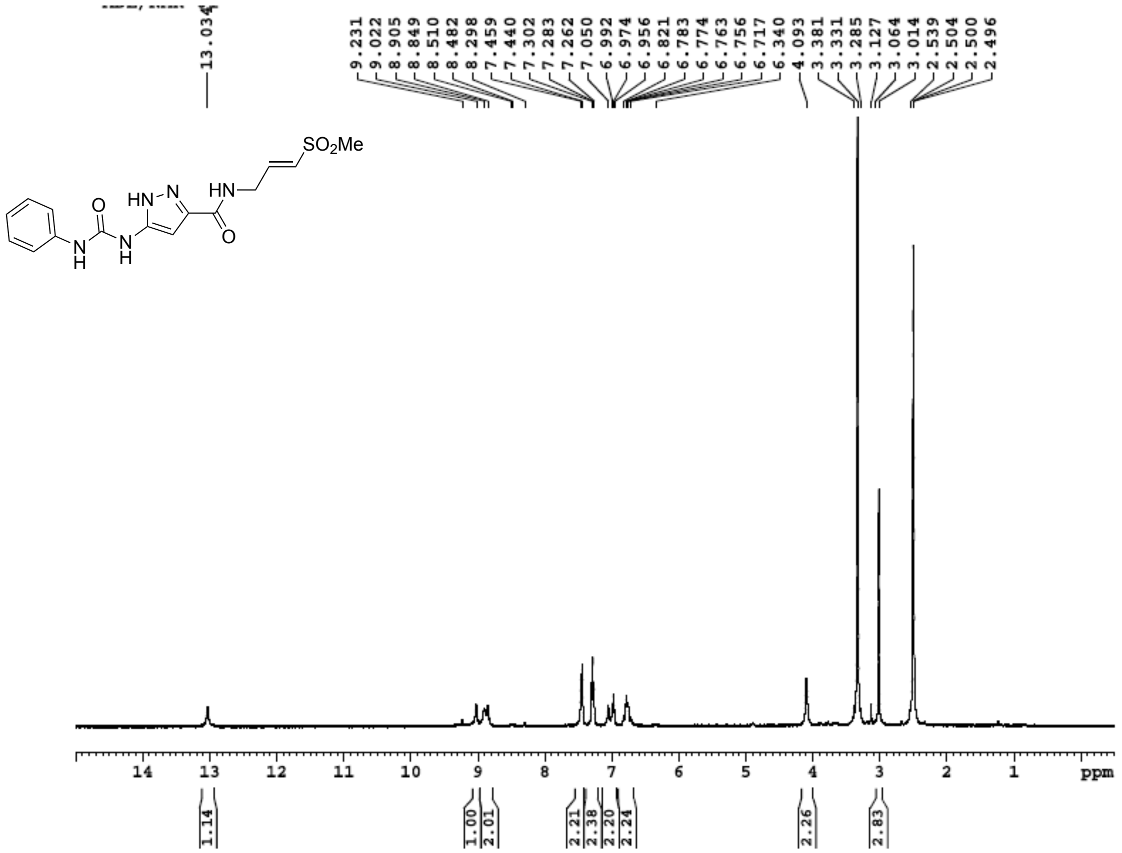

**Figure S63**: $^1$H NMR (101 MHz, DMSO-$d_6$) for **7d**

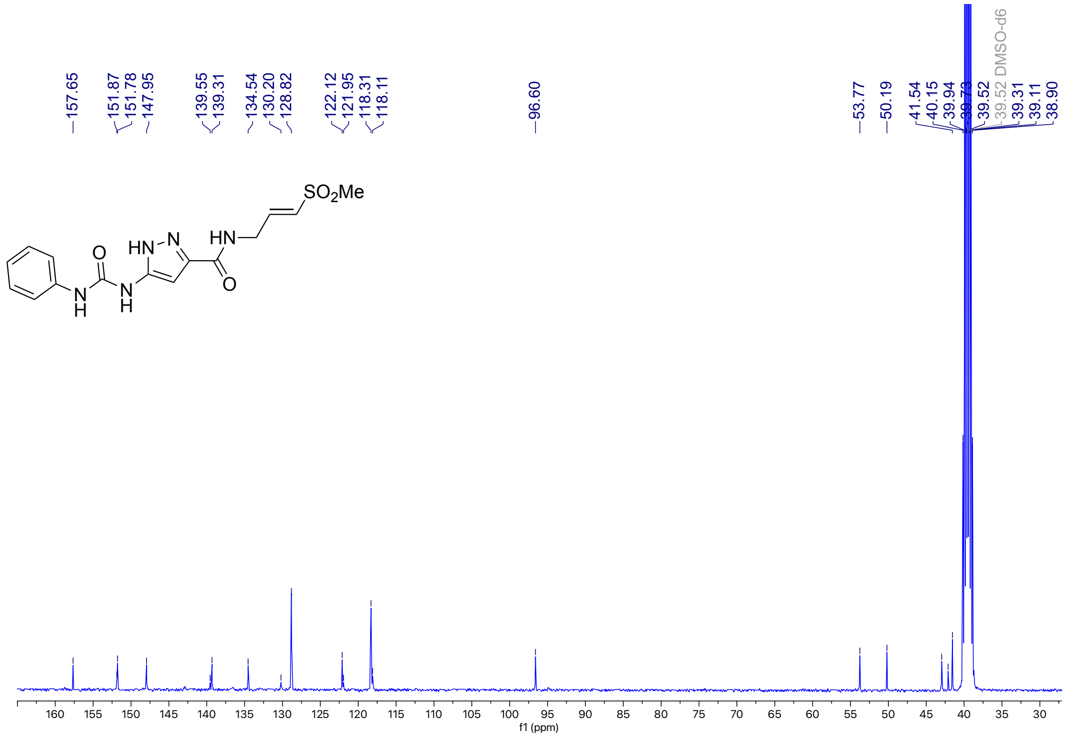

**Figure S64**: $^{1}$H NMR (400 MHz, DMSO-$d_6$) for **7e**

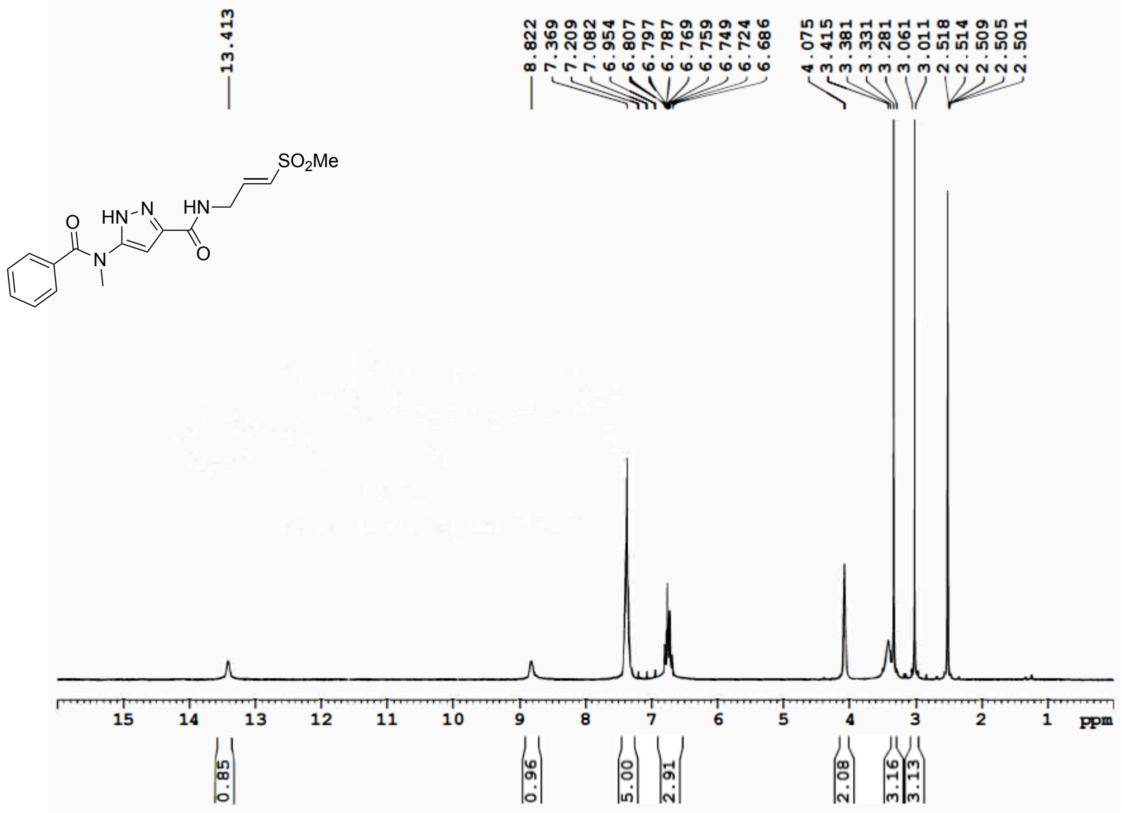


**Figure S65**: $^{13}$C NMR (100 MHz, DMSO-$d_6$) for **7e**

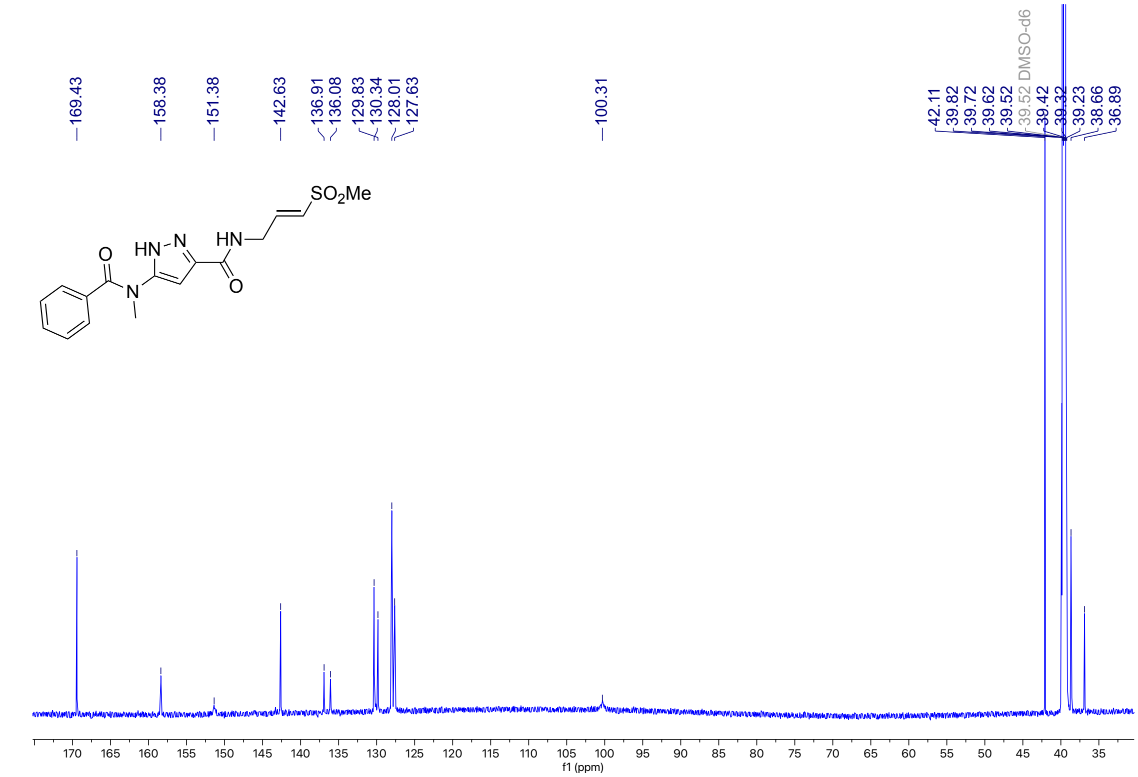

**Figure S66**: $^{1}$H NMR (400 MHz, DMSO-$d_6$) for **7f**

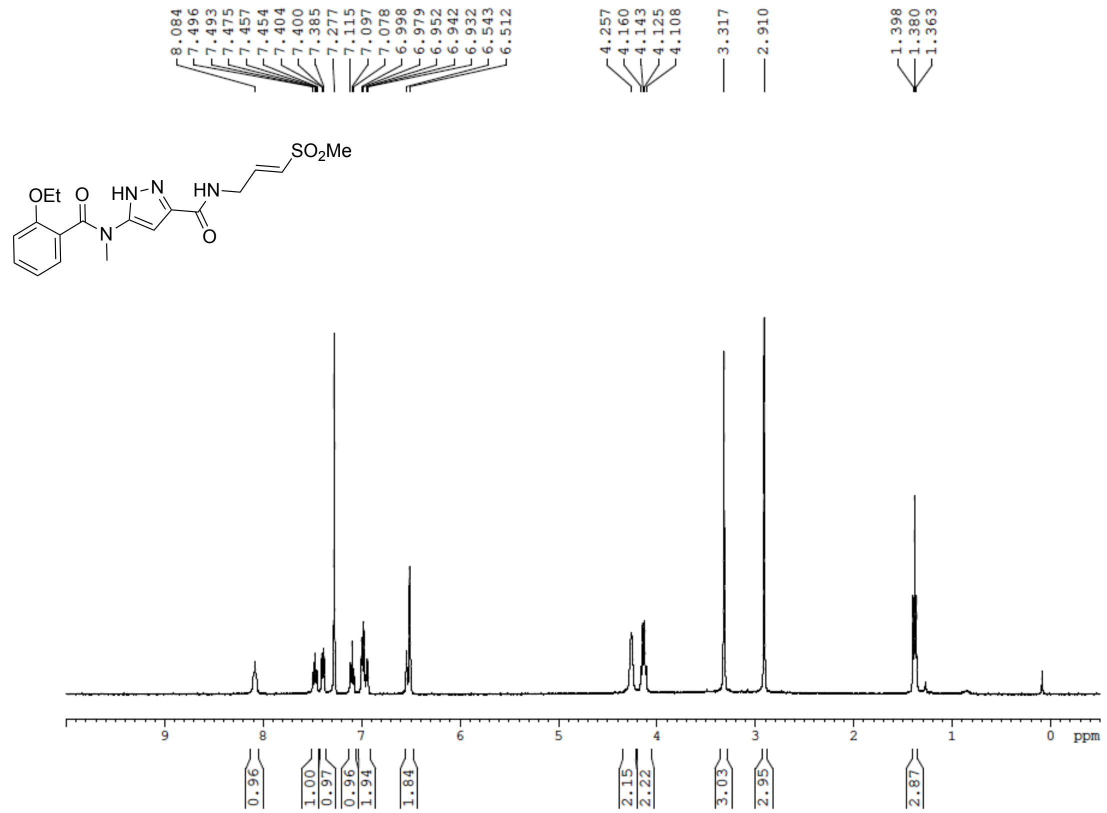


**Figure S67**: $^{13}$C NMR (100 MHz, DMSO-$d_6$) for **7f**

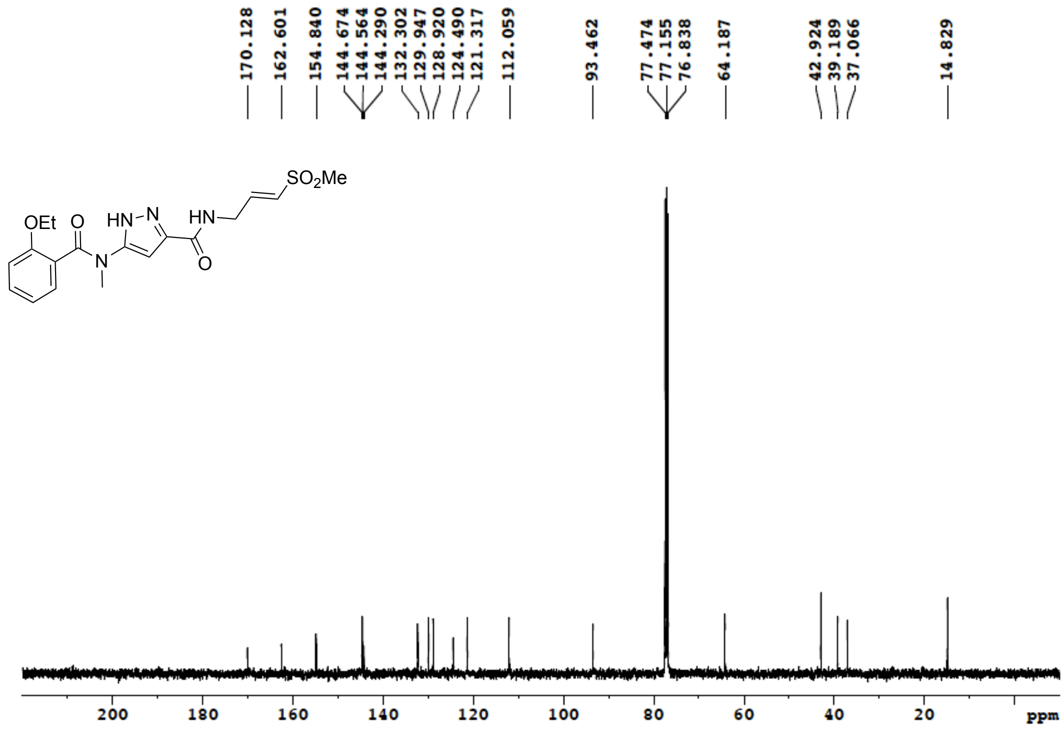

**Figure S68**: $^{1}H$ NMR (400 MHz, DMSO-$d_6$) for **8a**

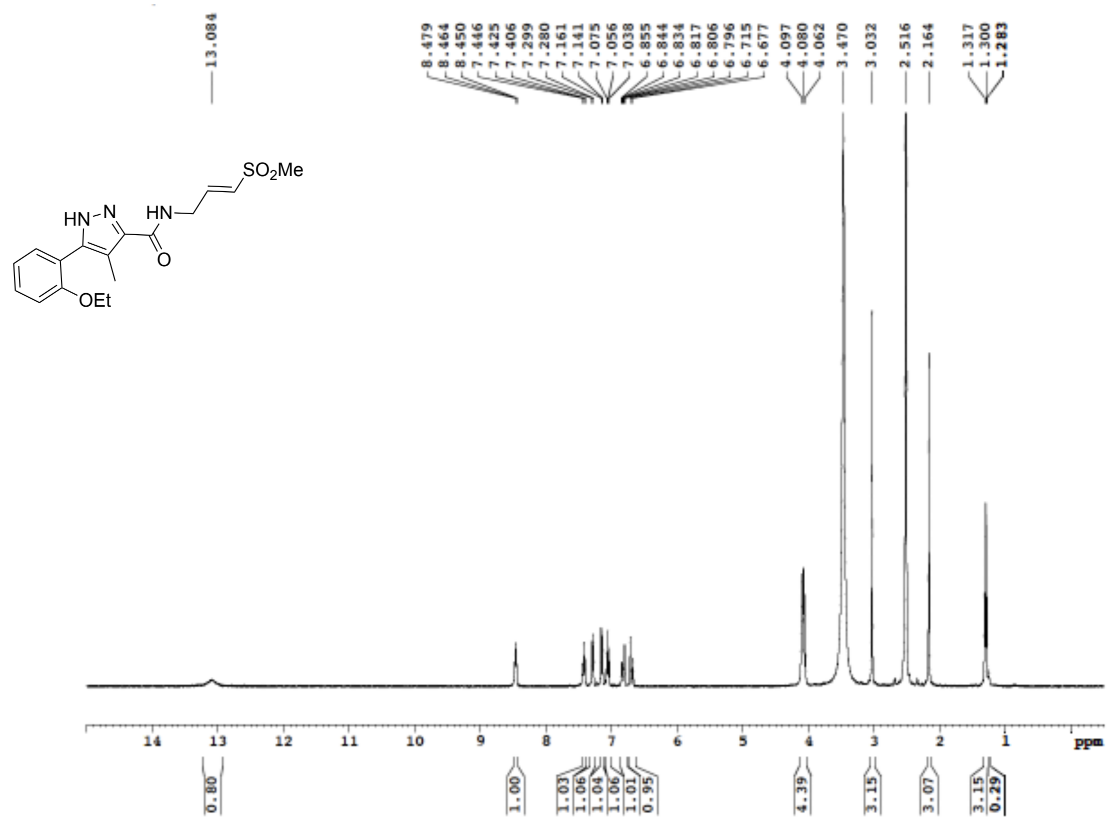

**Figure S69**: $^{13}C$ NMR (100 MHz, DMSO-$d_6$) for **8a**

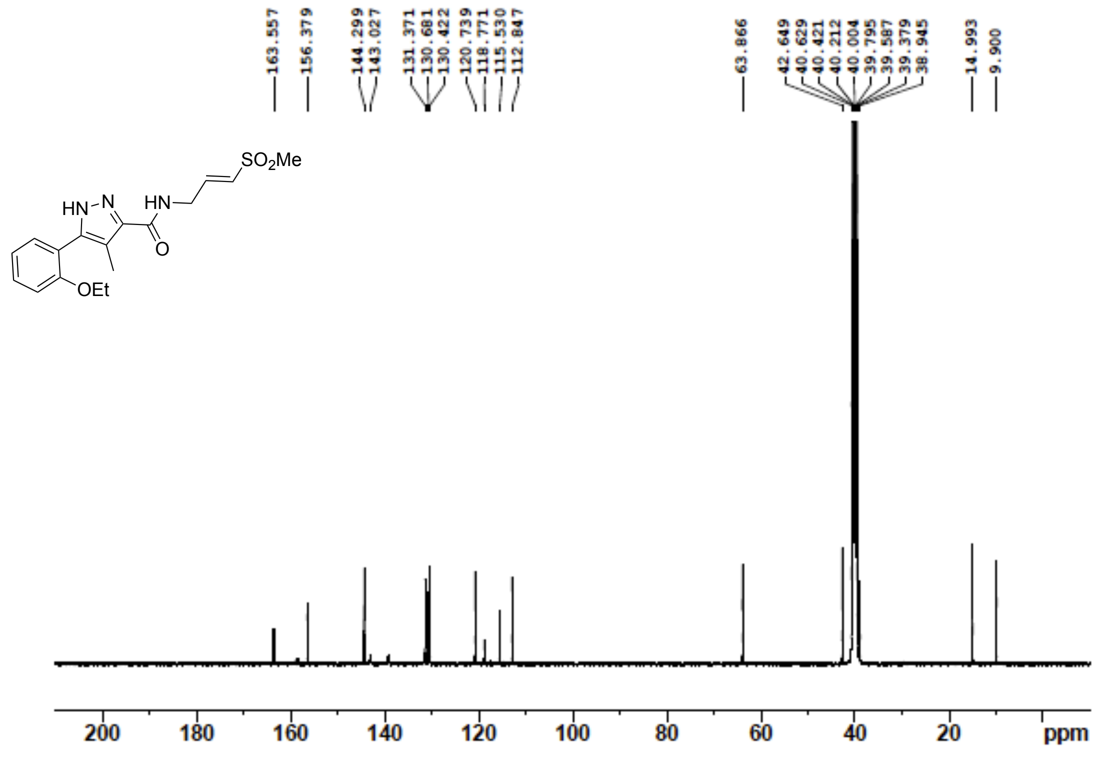

**Figure S70**: $^{1}$H NMR (400 MHz, DMSO-$d_6$) for **8b**

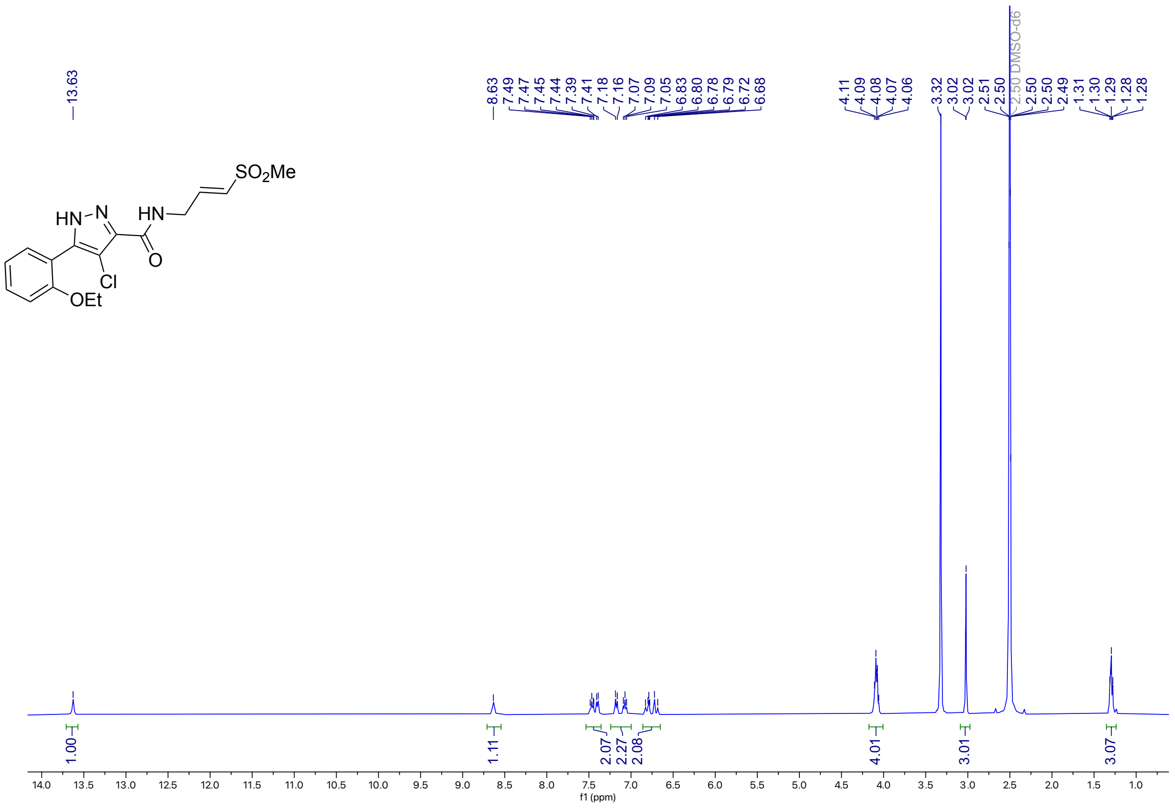


**Figure S71**: $^{13}$C NMR (100 MHz, DMSO-$d_6$) for **8b**

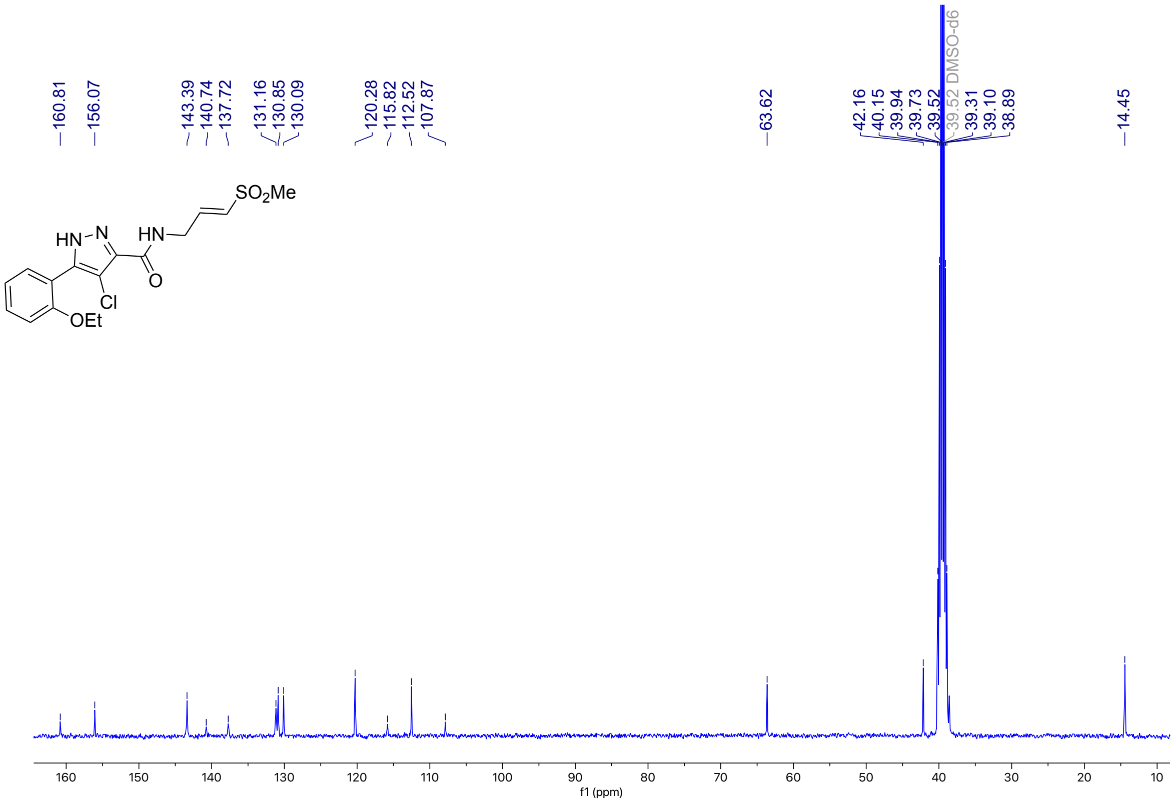

**Figure S72**: $^{1}$H NMR (400 MHz, DMSO-$d_6$) for **8c**

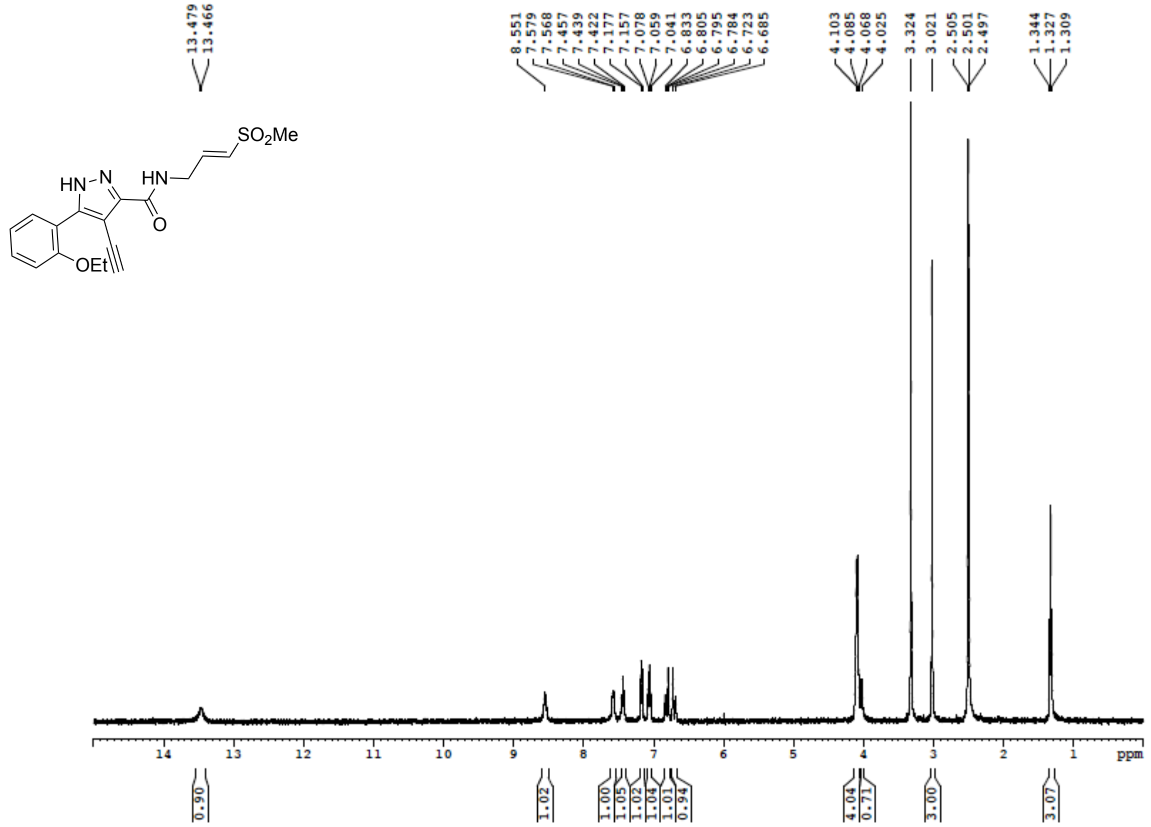


**Figure S73**: $^{13}$C NMR (100 MHz, DMSO-$d_6$) for **8c**

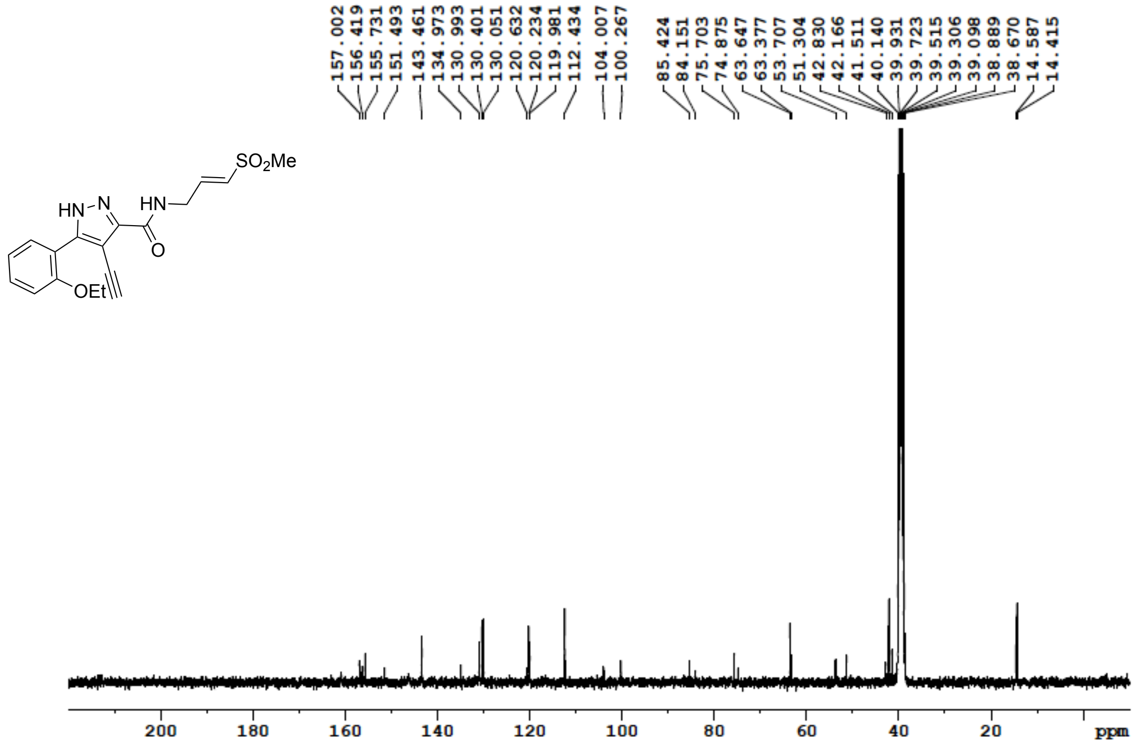

**Figure S74**: $^{1}H$ NMR (400 MHz, DMSO-$d_6$) for **8d**

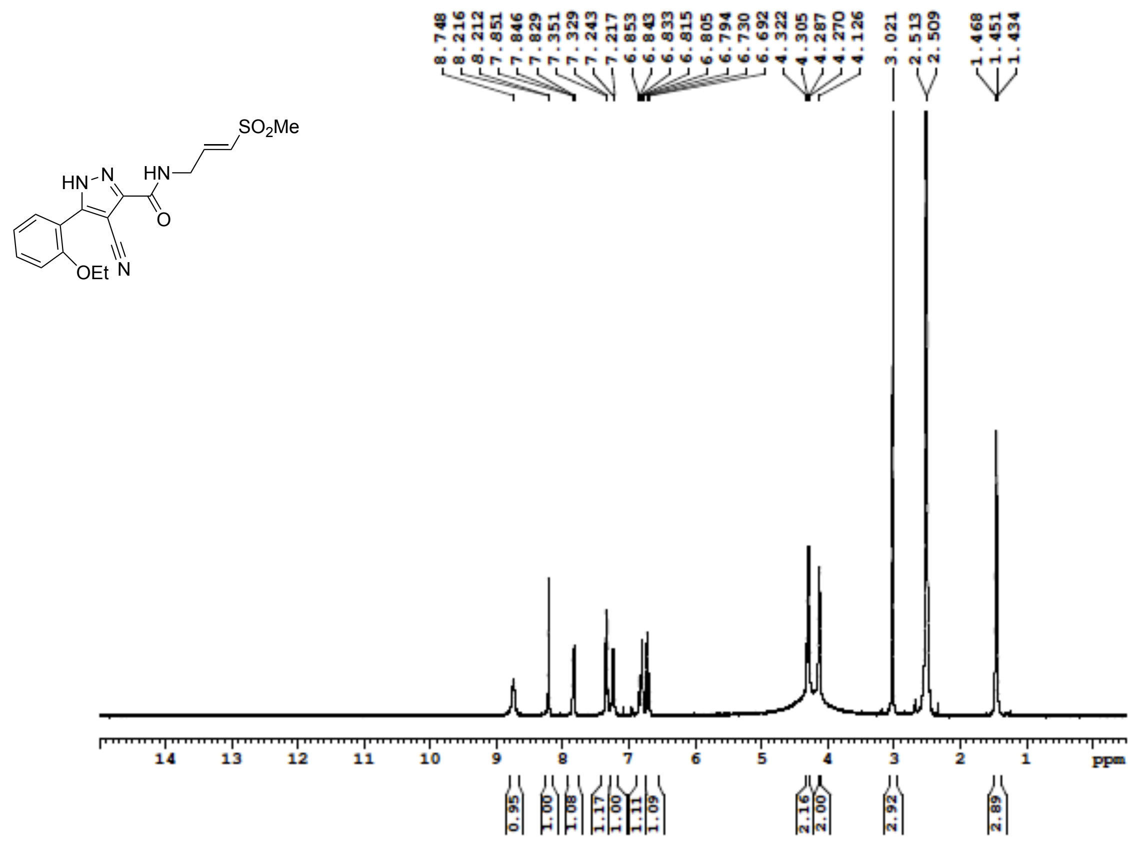

**Figure S75**: $^{13}C$ NMR (100 MHz, DMSO-$d_6$) for **8d**

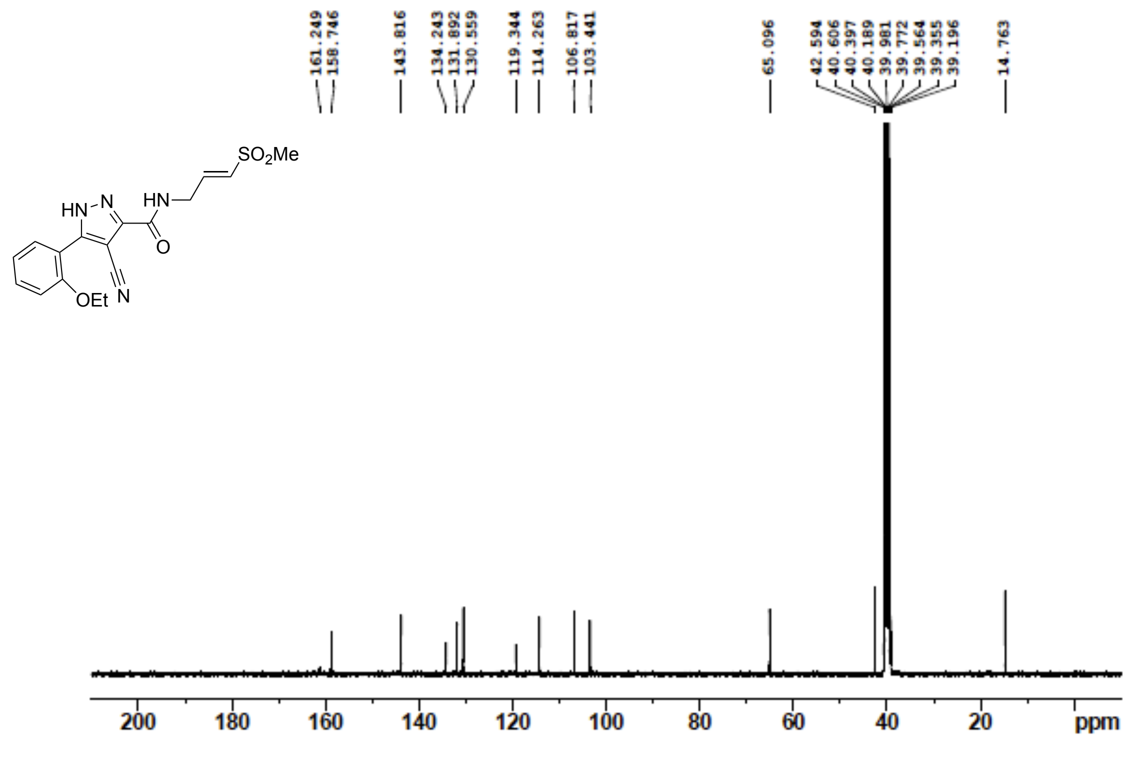

**Figure S76**: $^{1}H$ NMR (400 MHz, DMSO-$d_6$) for **9**

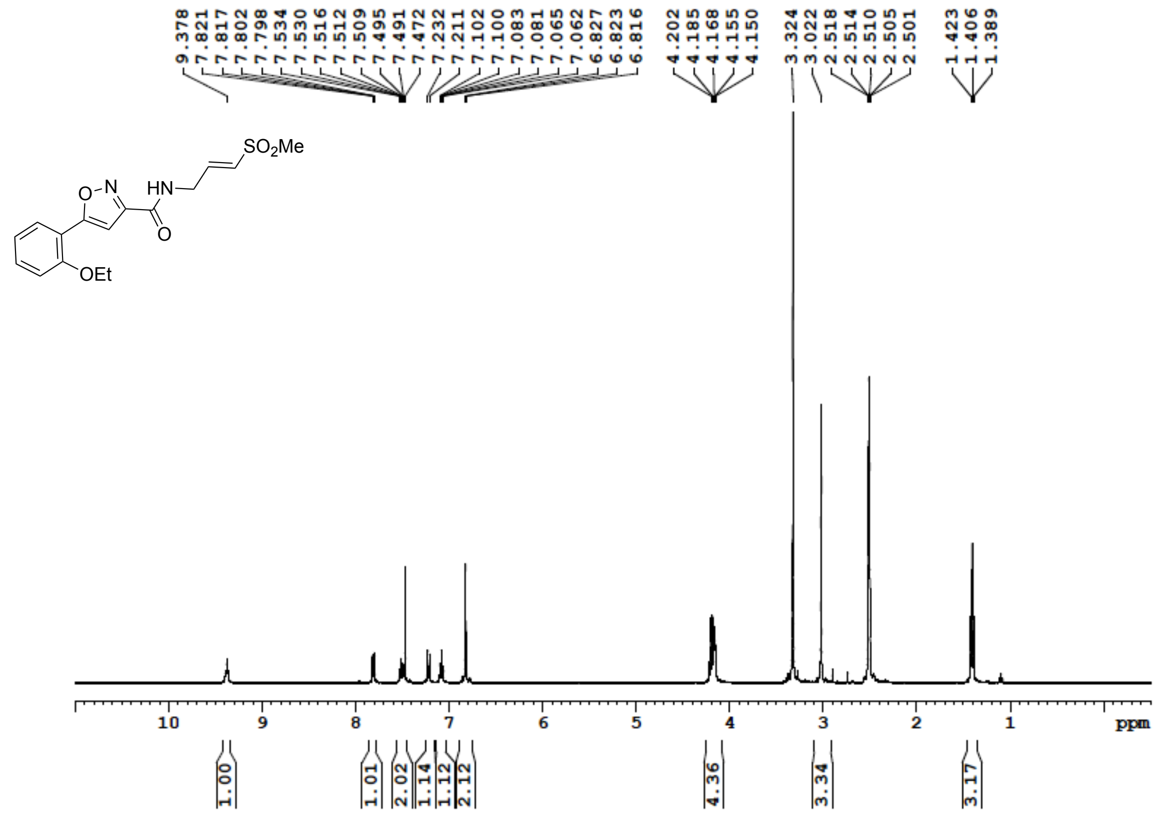


**Figure S77**: $^{13}C$ NMR (100 MHz, DMSO-$d_6$) for **9**

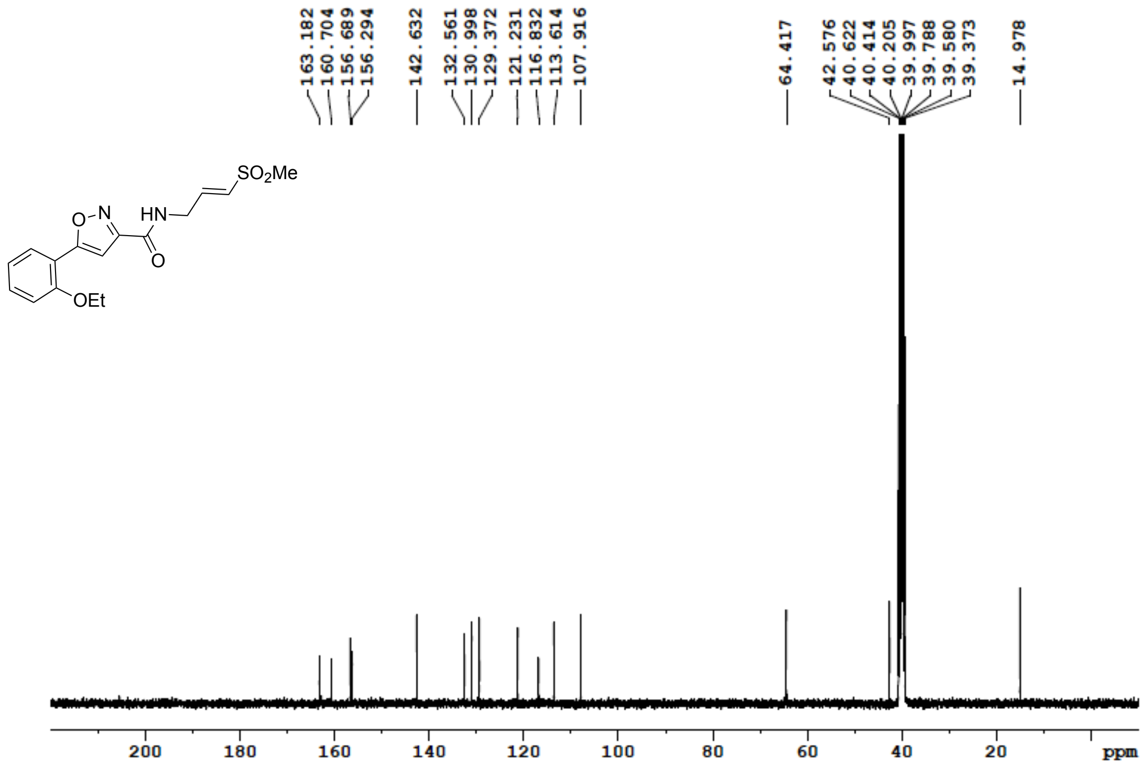

**Figure S78**: $^{1}$H NMR (400 MHz, DMSO-$d_6$) for **10**

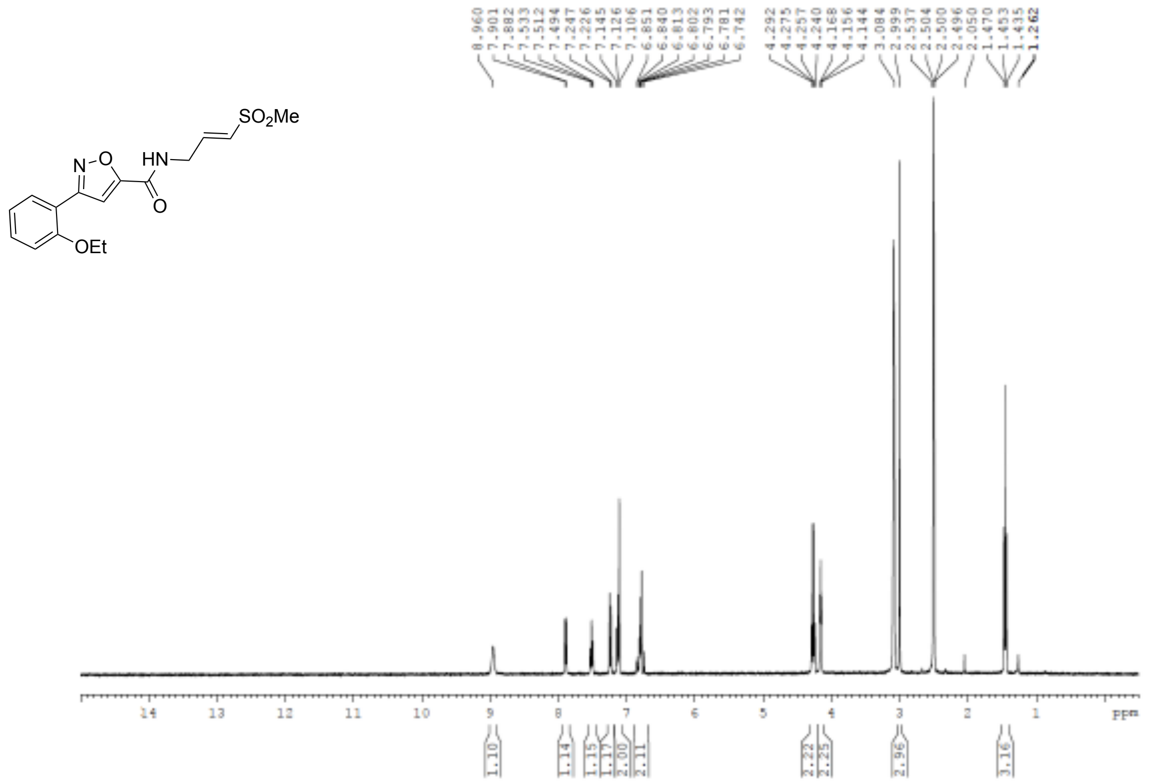


**Figure S79**: $^{13}$C NMR (100 MHz, DMSO-$d_6$) for **10**

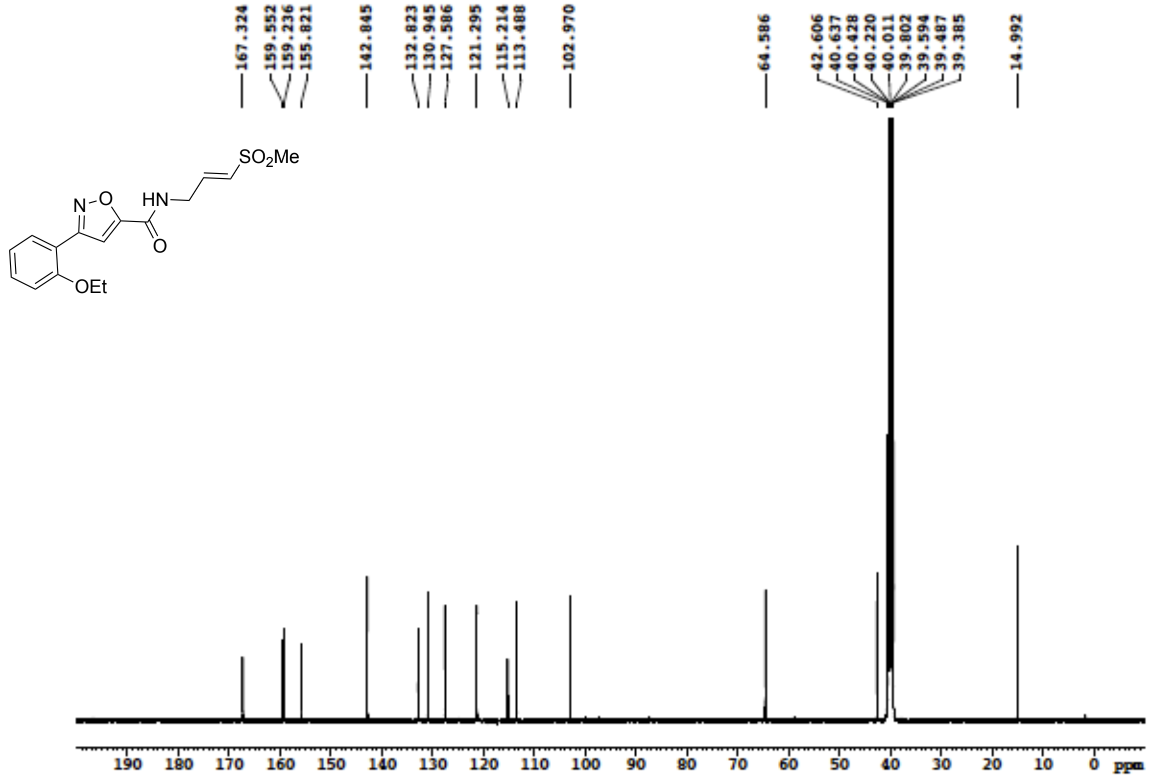

**Figure S80**: $^{1}$H NMR (400 MHz, DMSO-$d_6$) for **11**

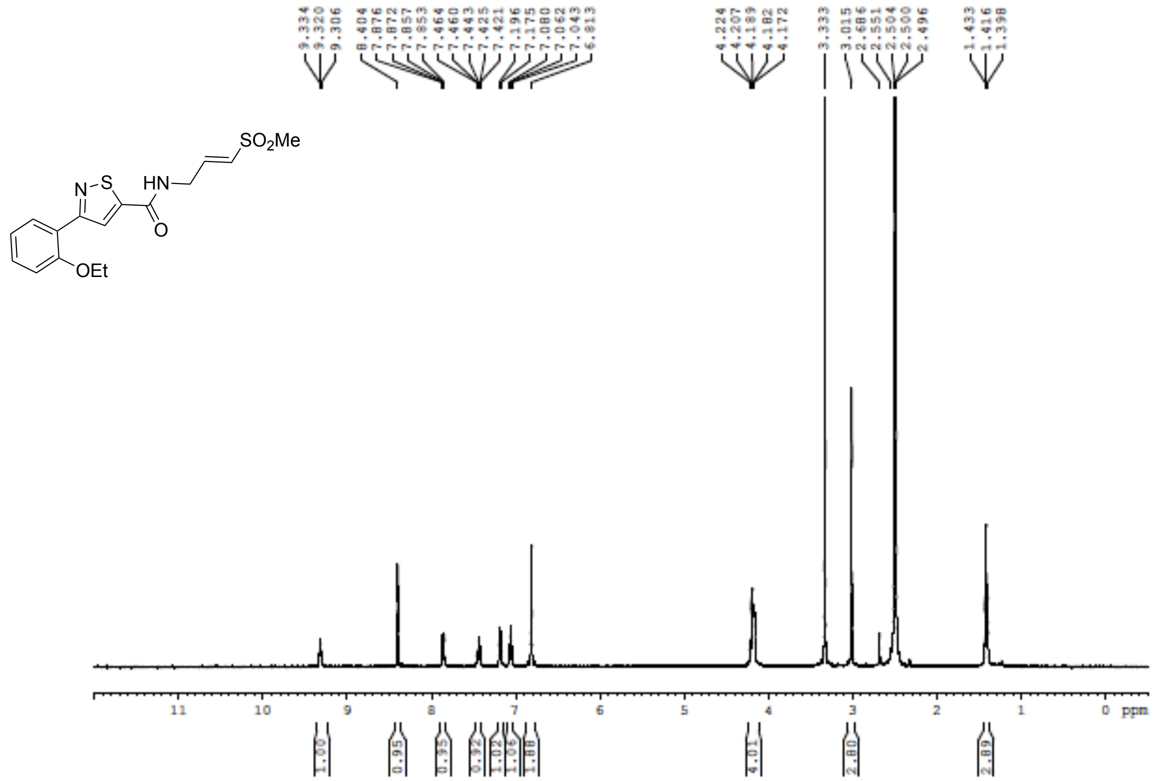

**Figure S81**: $^{13}$C NMR (100 MHz, DMSO-$d_6$) for **11**

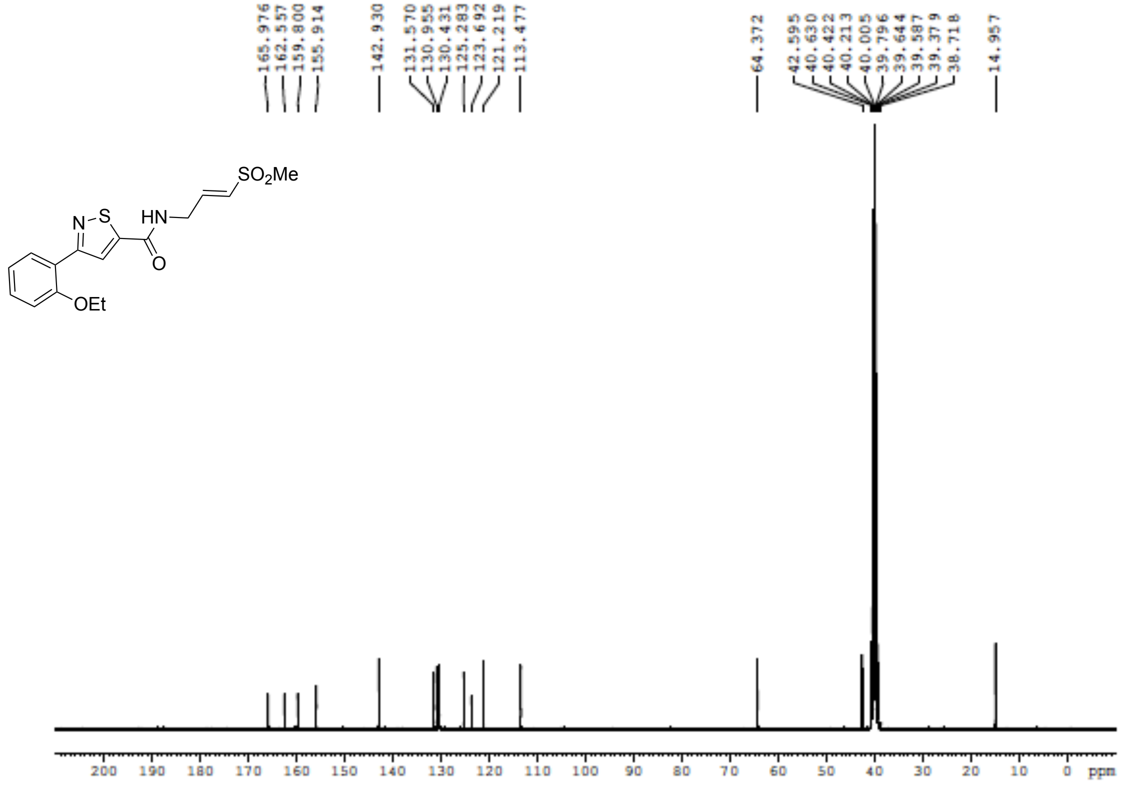

**Figure S82**: $^{1}$H NMR (400 MHz, DMSO-$d_6$) for **12**

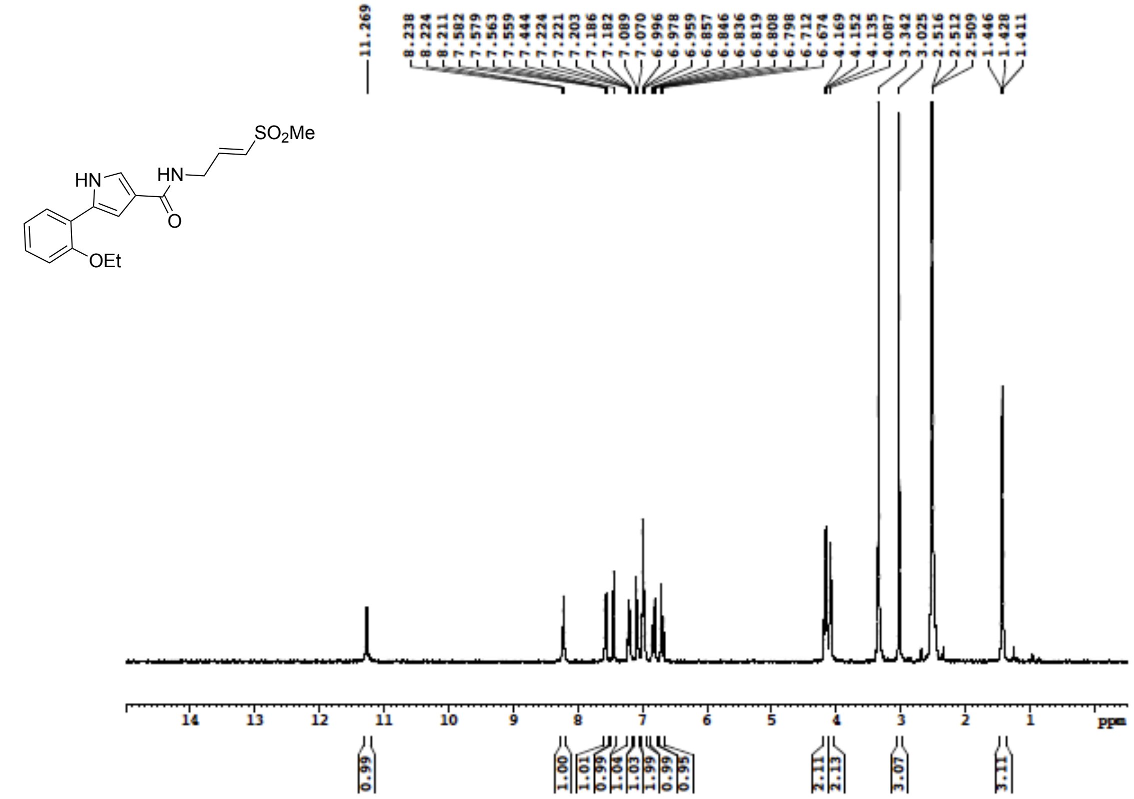


**Figure S83**: $^{13}$C NMR (100 MHz, DMSO-$d_6$) for **12**

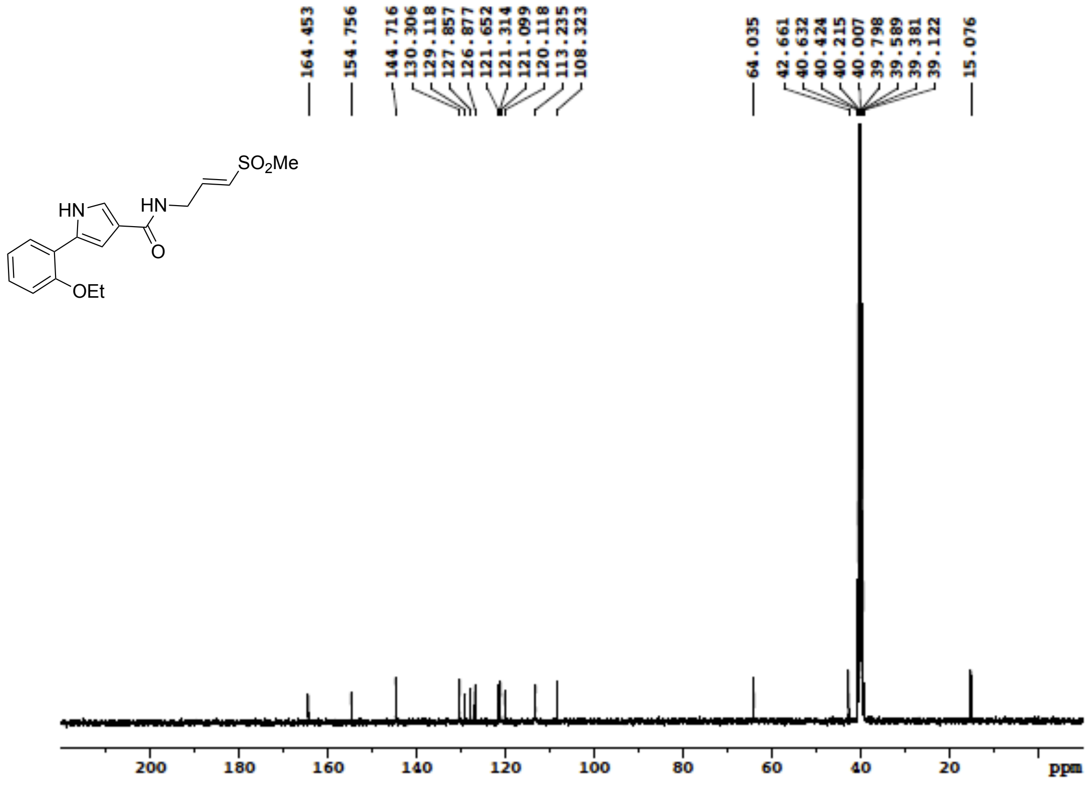

**Figure S84**: $^{1}$H NMR (400 MHz, DMSO-$d_6$) for **13**

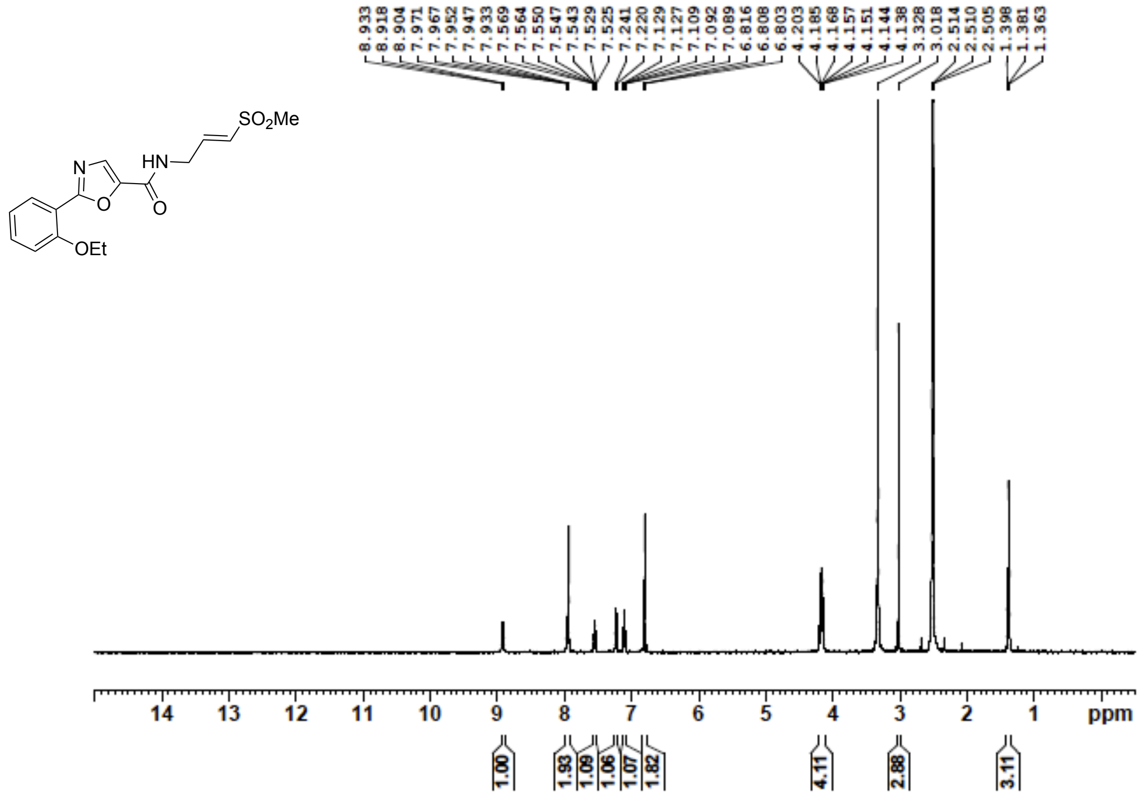


**Figure S85**: $^{13}$C NMR (100 MHz, DMSO-$d_6$) for **13**

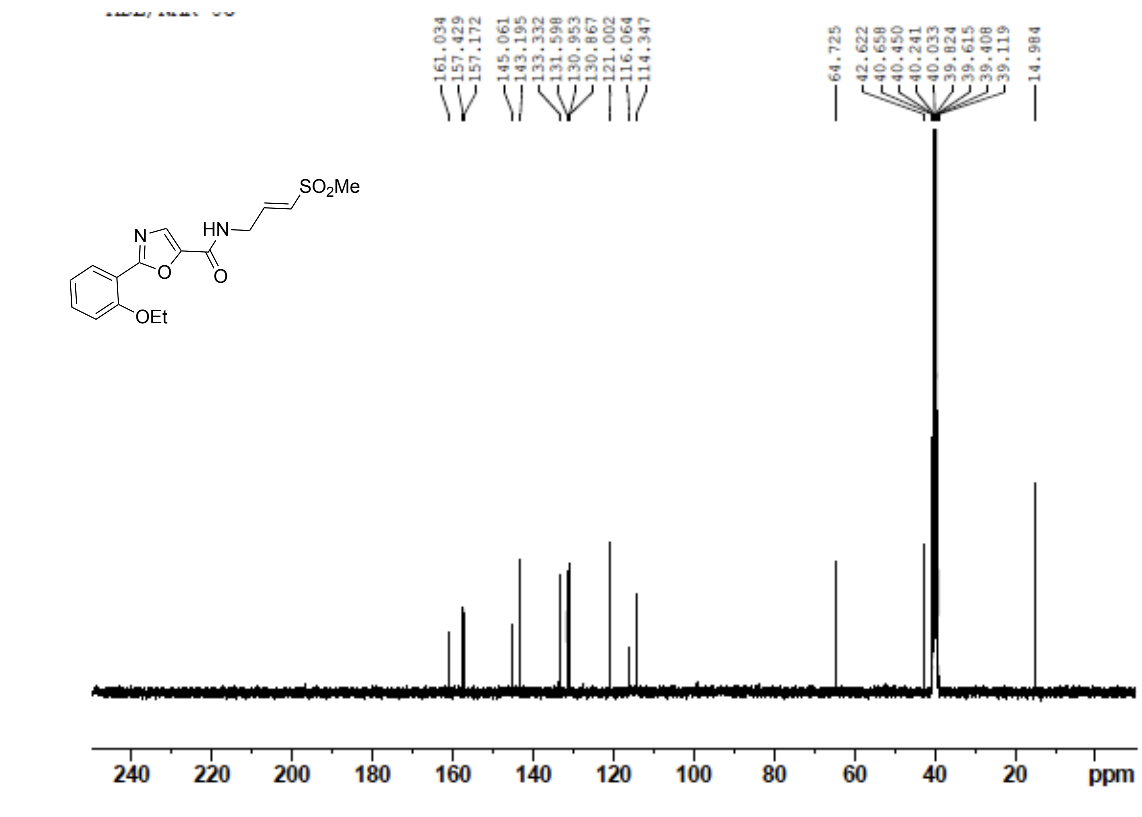

**Figure S86**: $^{1}$H NMR (400 MHz, DMSO-$d_6$) for **14**

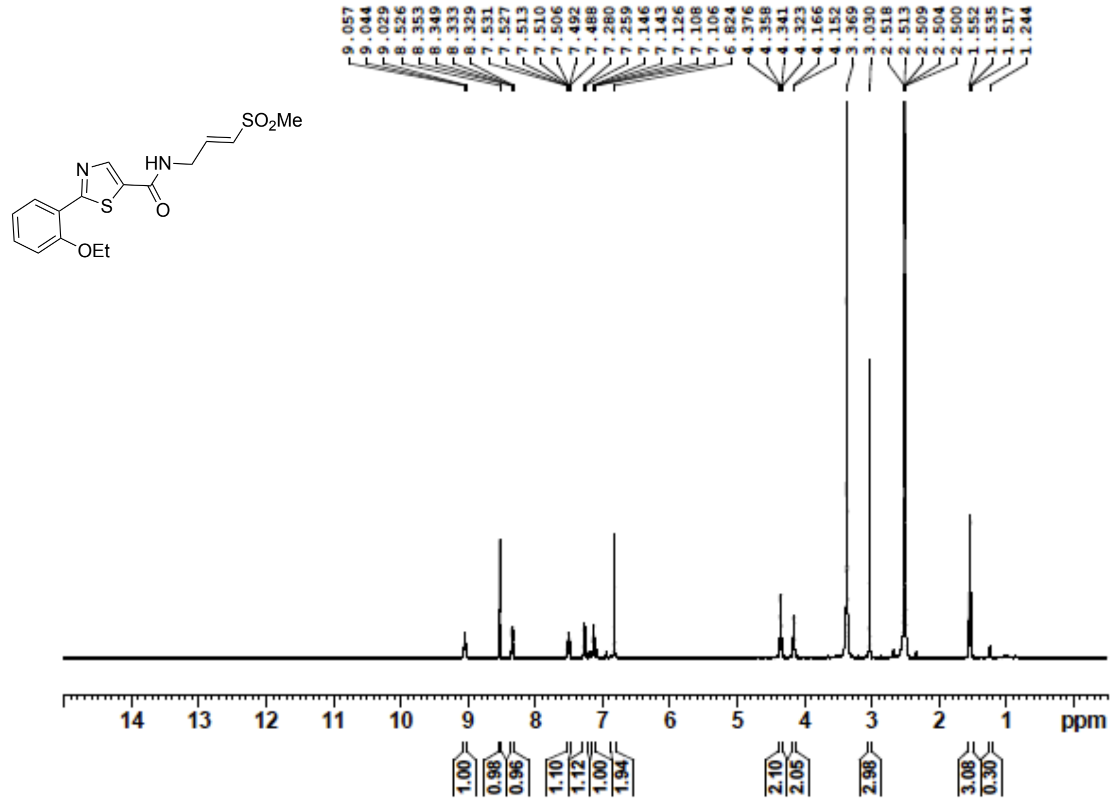


**Figure S87**: $^{13}$C NMR (100 MHz, DMSO-$d_6$) for **14**

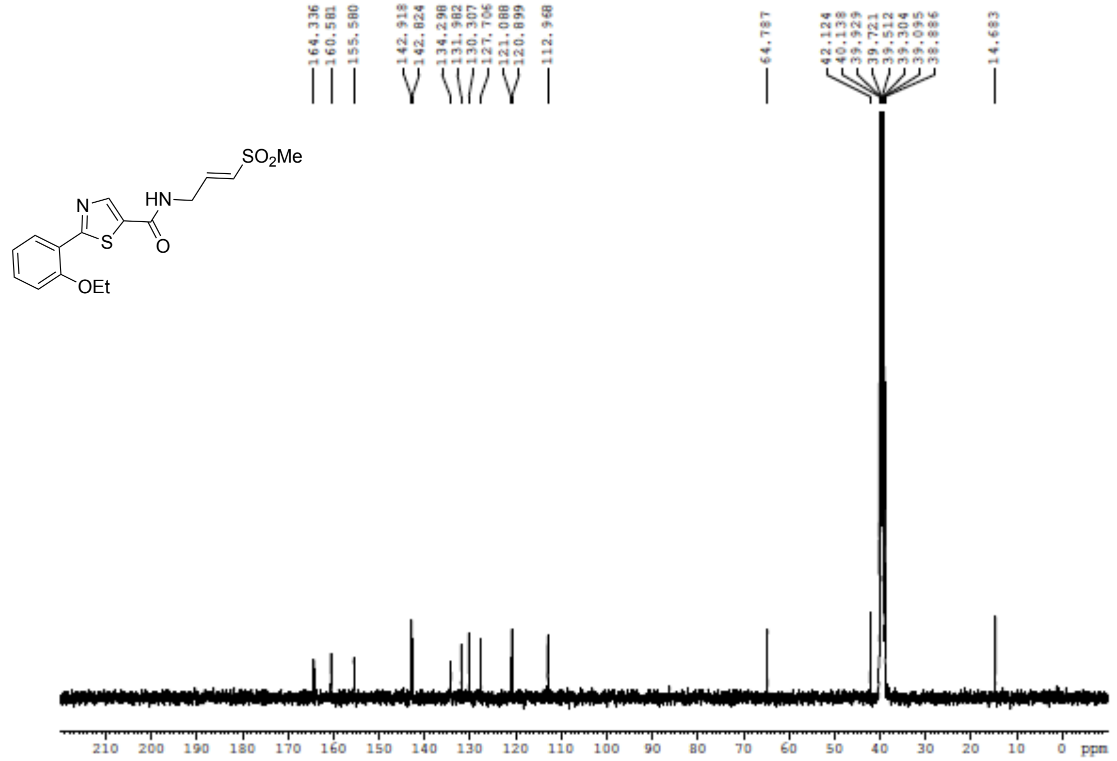

**Figure S88**: $^{1}$H NMR (400 MHz, DMSO-$d_6$) for **15**

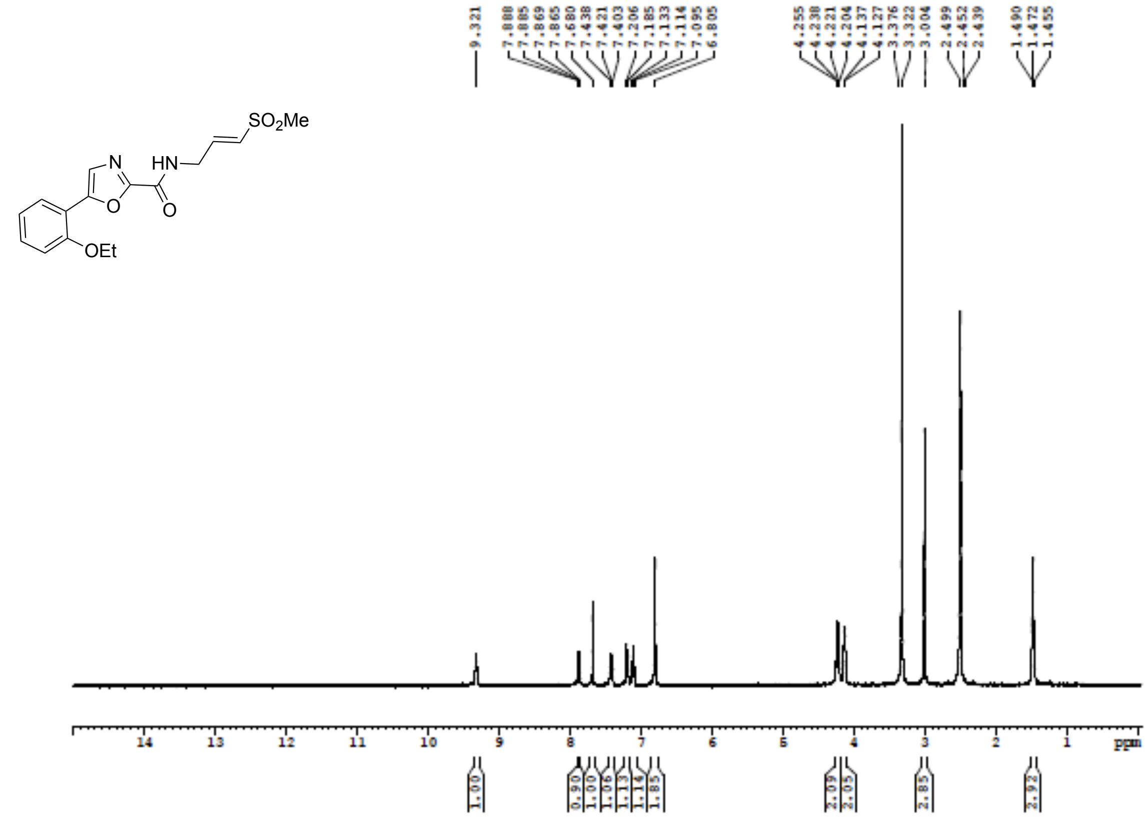


**Figure S89**: $^{13}$C NMR (100 MHz, DMSO-$d_6$) for **15**

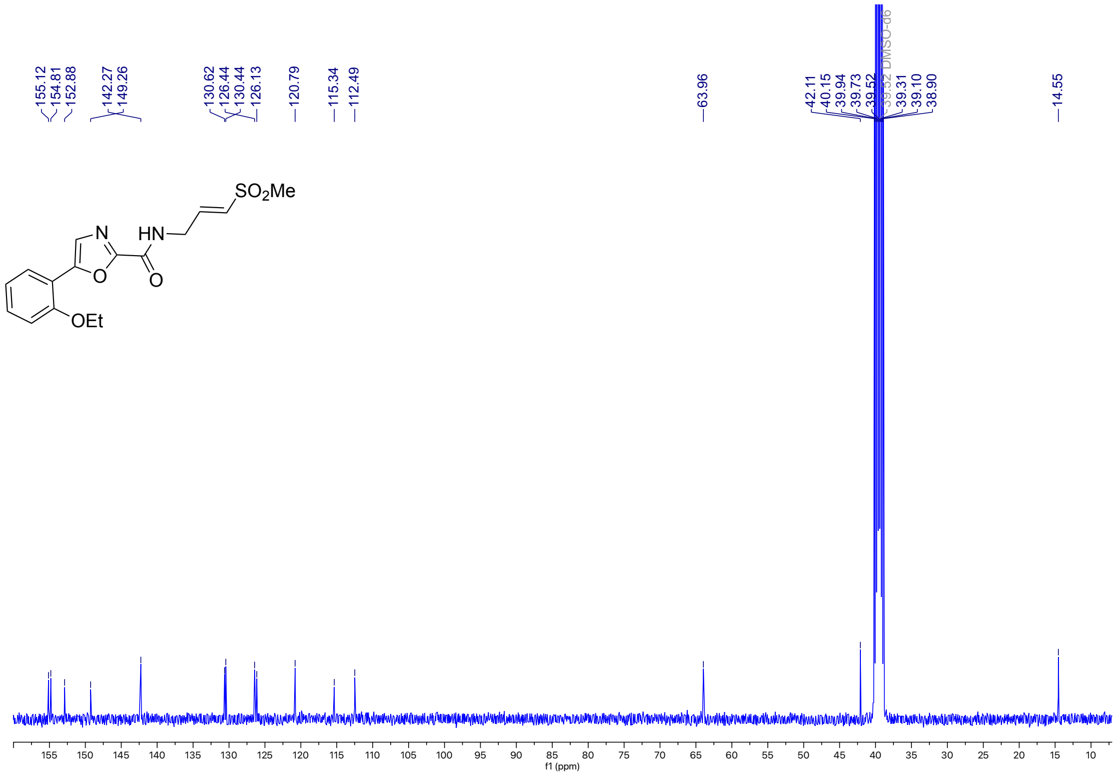

**Figure S90**: $^{1}$H NMR (400 MHz, DMSO-$d_6$) for **16**

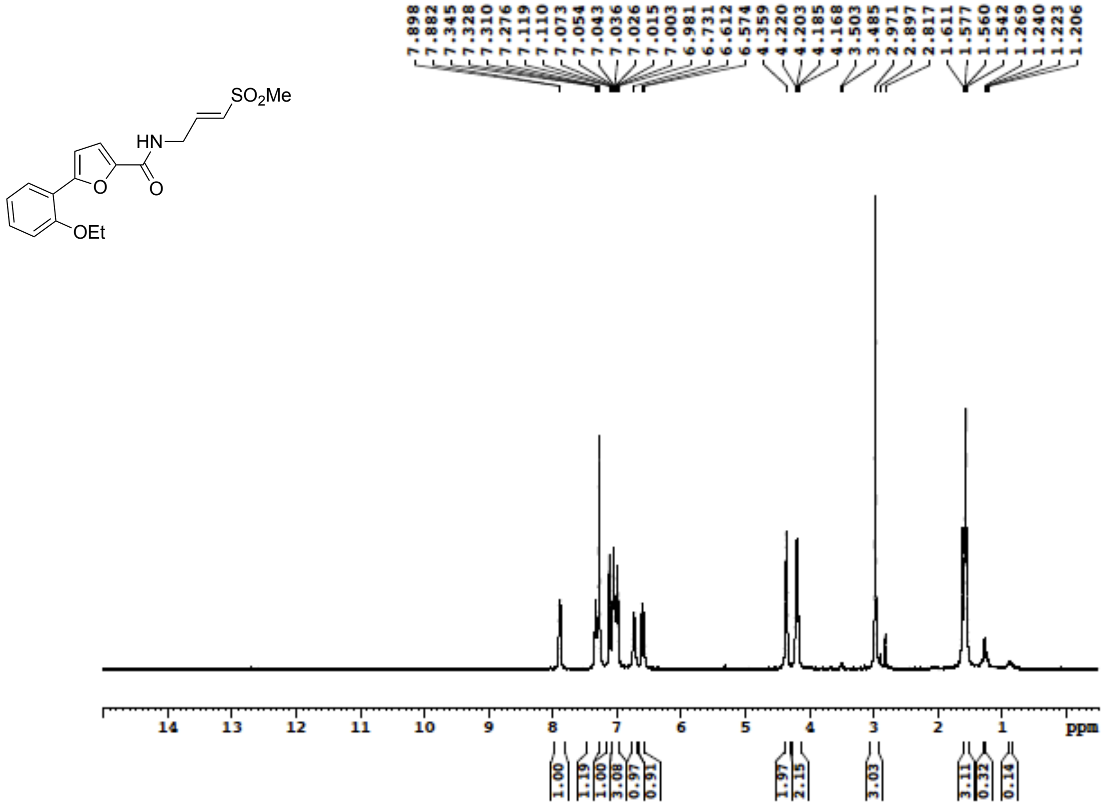


**Figure S91**: $^{13}$C NMR (100 MHz, DMSO-$d_6$) for **16**

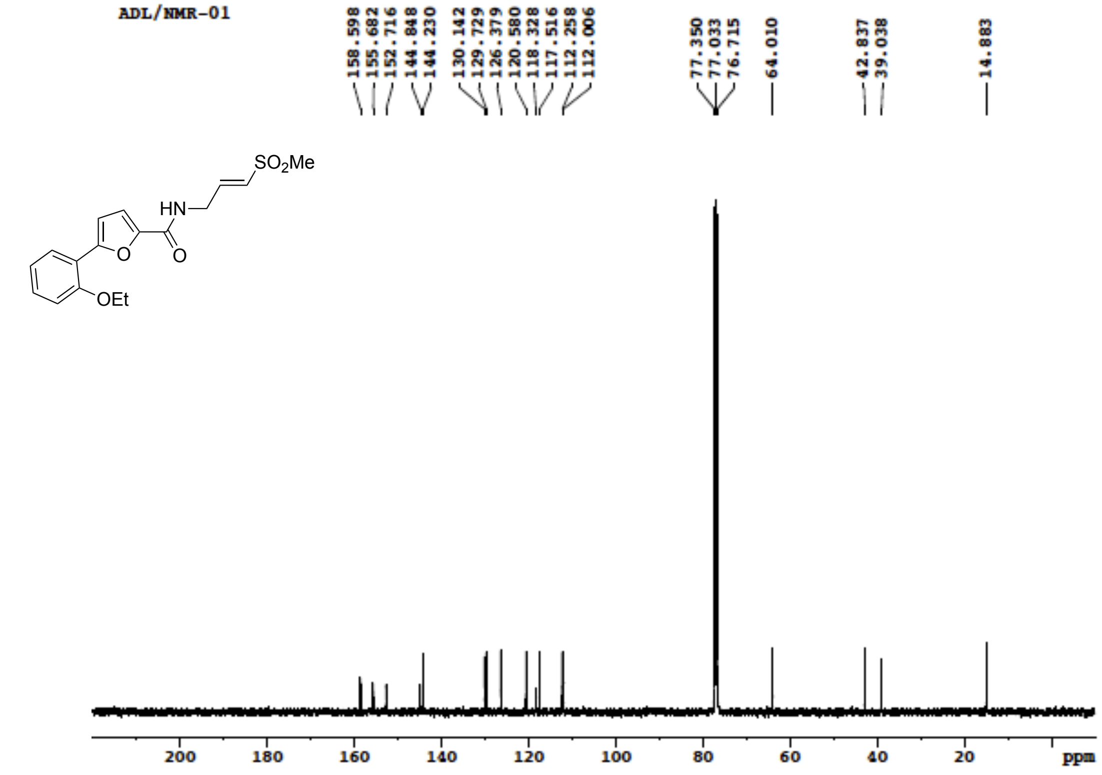

**Figure S92**: $^{1}H$ NMR (400 MHz, DMSO-$d_6$) for **17**

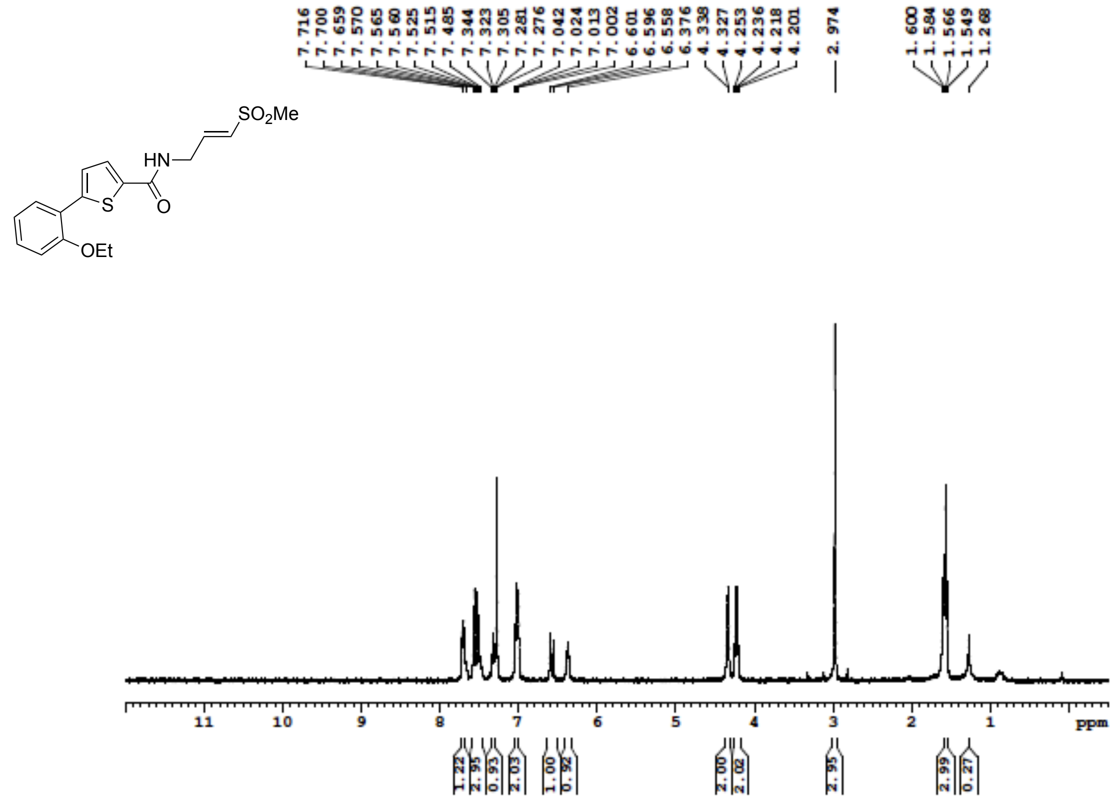


**Figure S93**: $^{13}C$ NMR (100 MHz, DMSO-$d_6$) for **17**

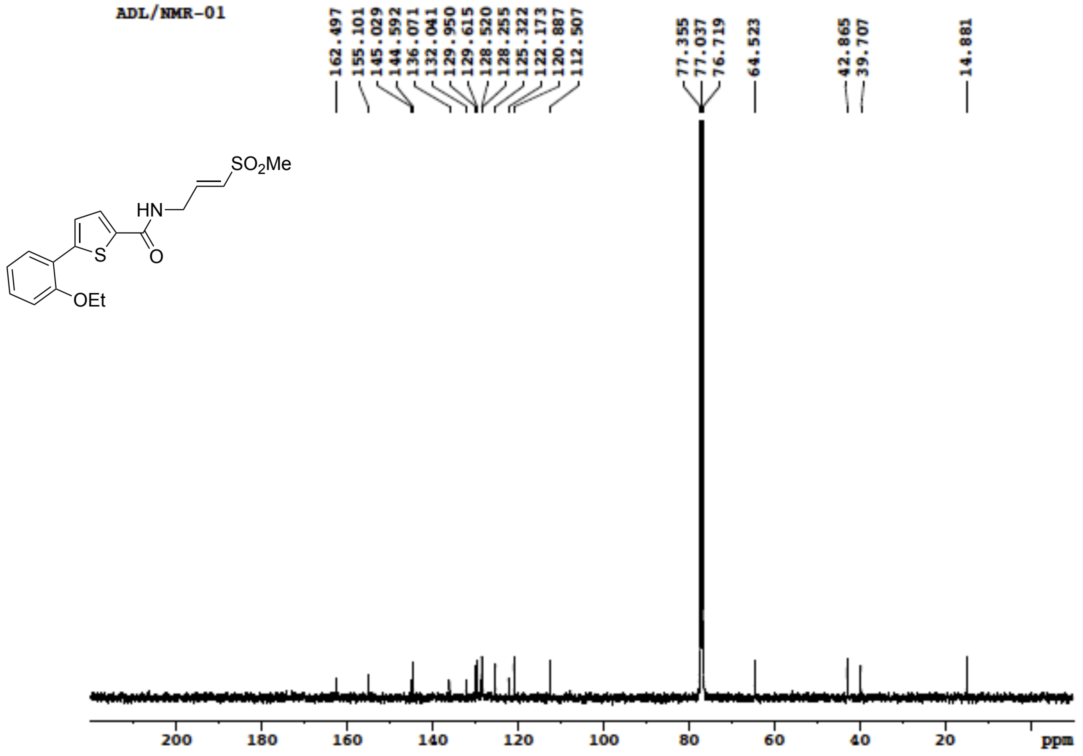

**Figure S94**: $^{1}$H NMR (400 MHz, DMSO-$d_6$) for **18**

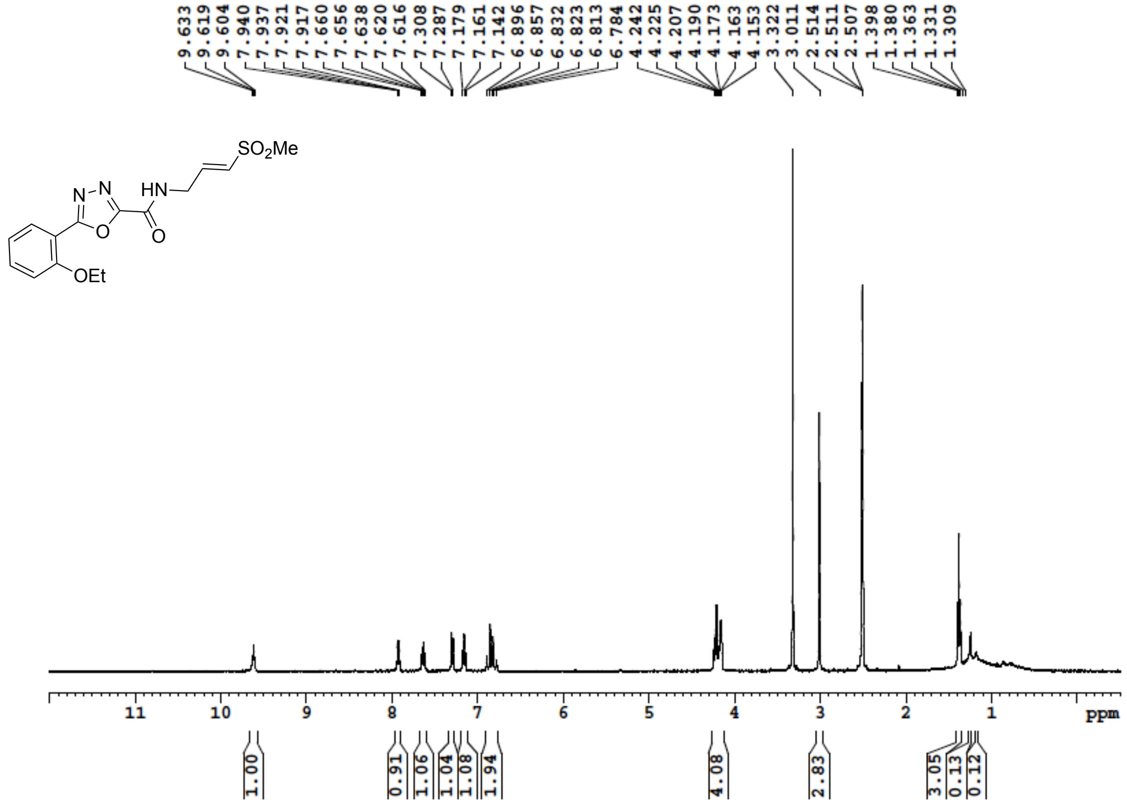

**Figure S95**: $^{13}$C NMR (101 MHz, DMSO-$d_6$) for **18**

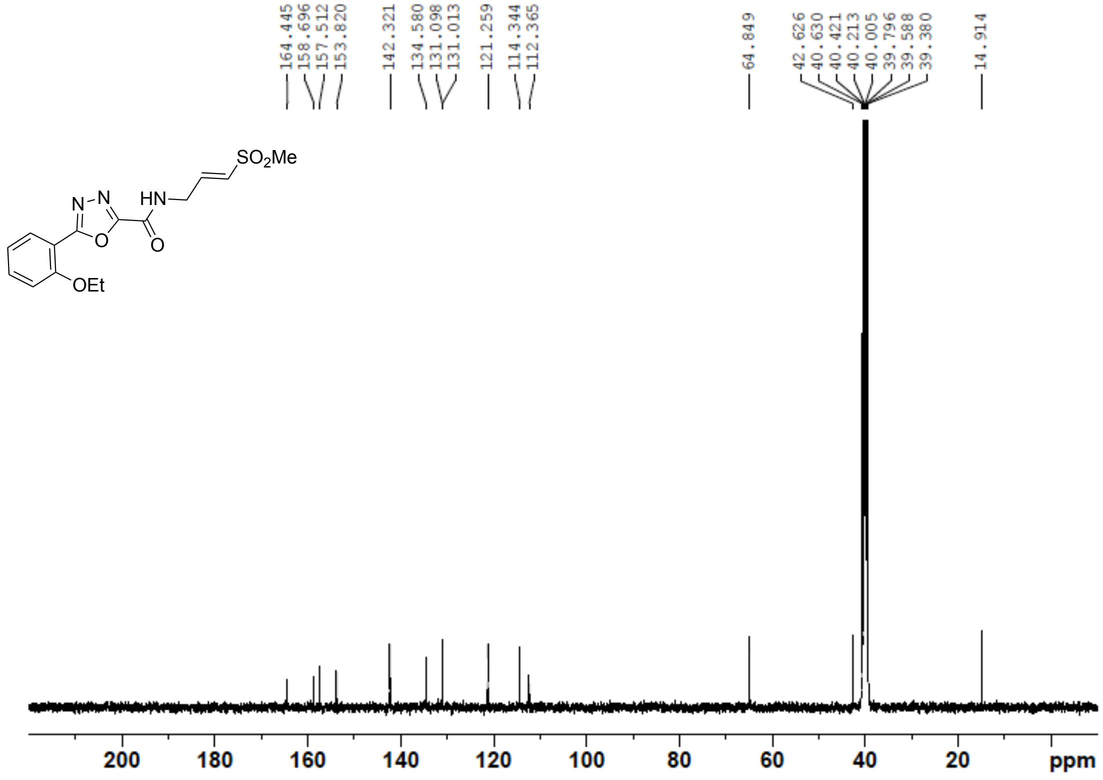

**Figure S96**: $^{1}$H NMR (400 MHz, DMSO-$d_6$) for **19**

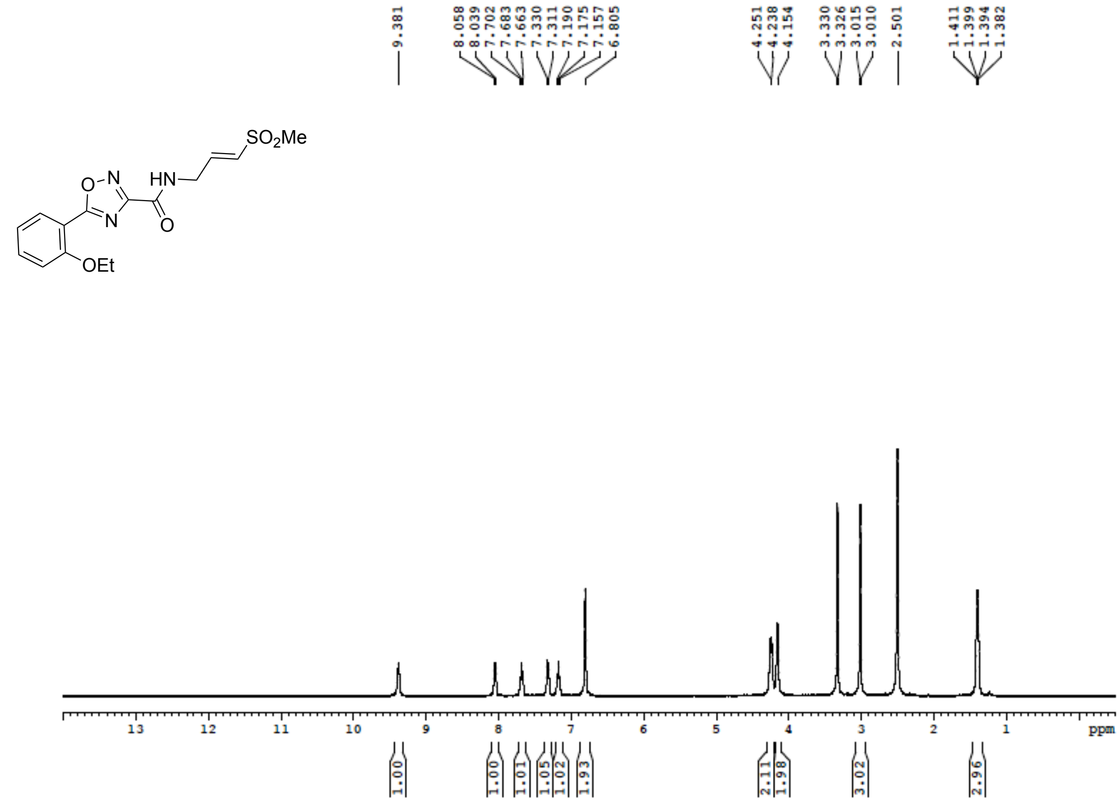


**Figure S97**: $^{13}$C NMR (100 MHz, DMSO-$d_6$) for **19**

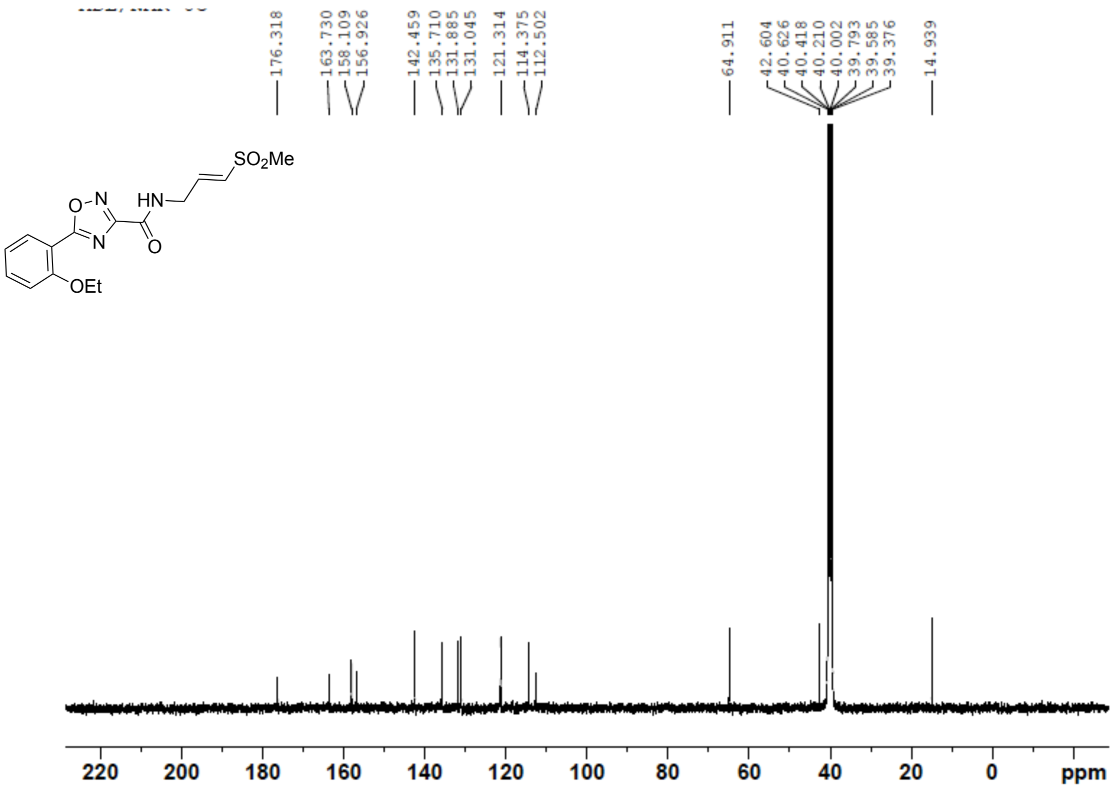

**Figure S98**: $^{1}H$ NMR (400 MHz, DMSO-$d_6$) for **20**

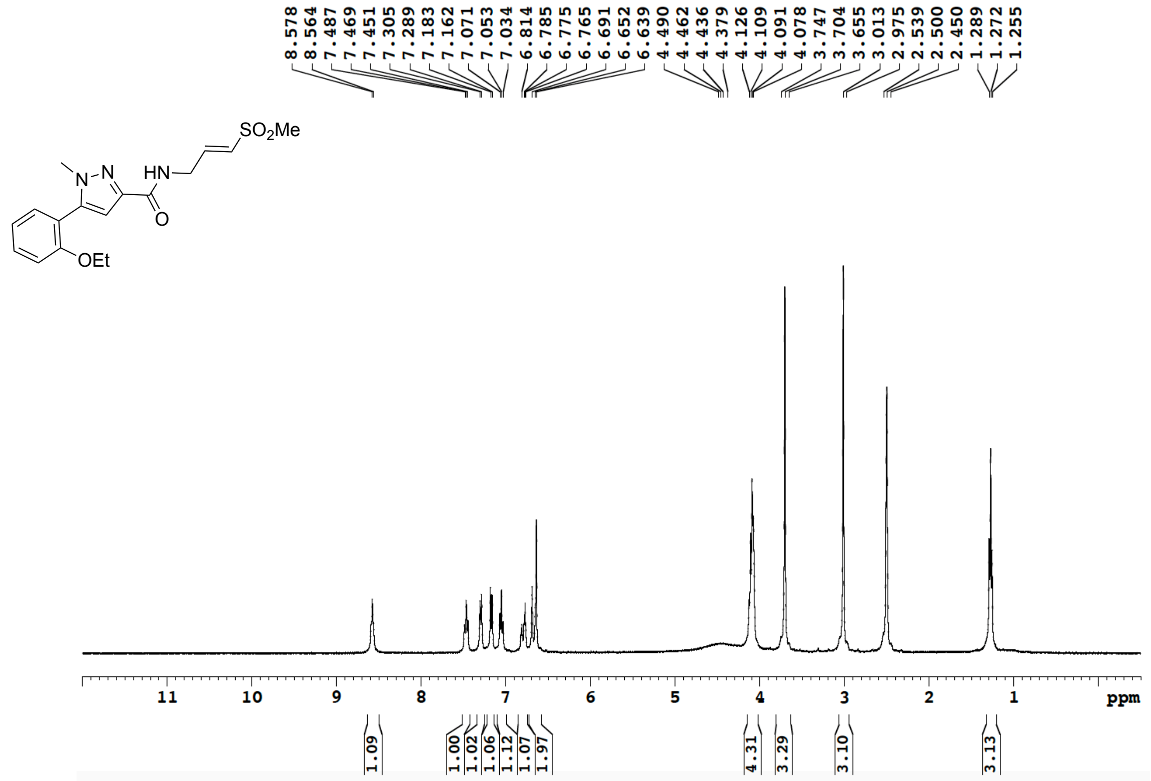

**Figure S99**: $^{13}C$ NMR (100 MHz, DMSO-$d_6$) for **20**

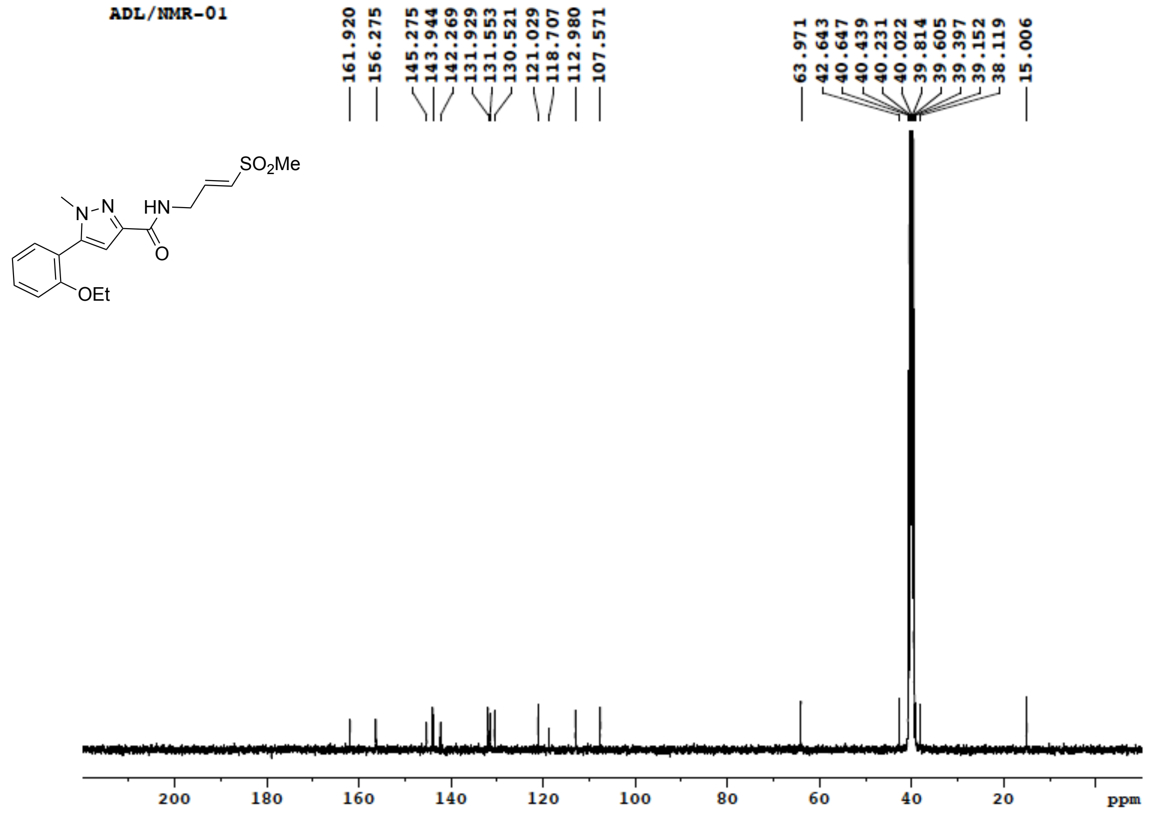

**Figure S100**: $^{1}$H NMR (400 MHz, DMSO-$d_6$) for **21**

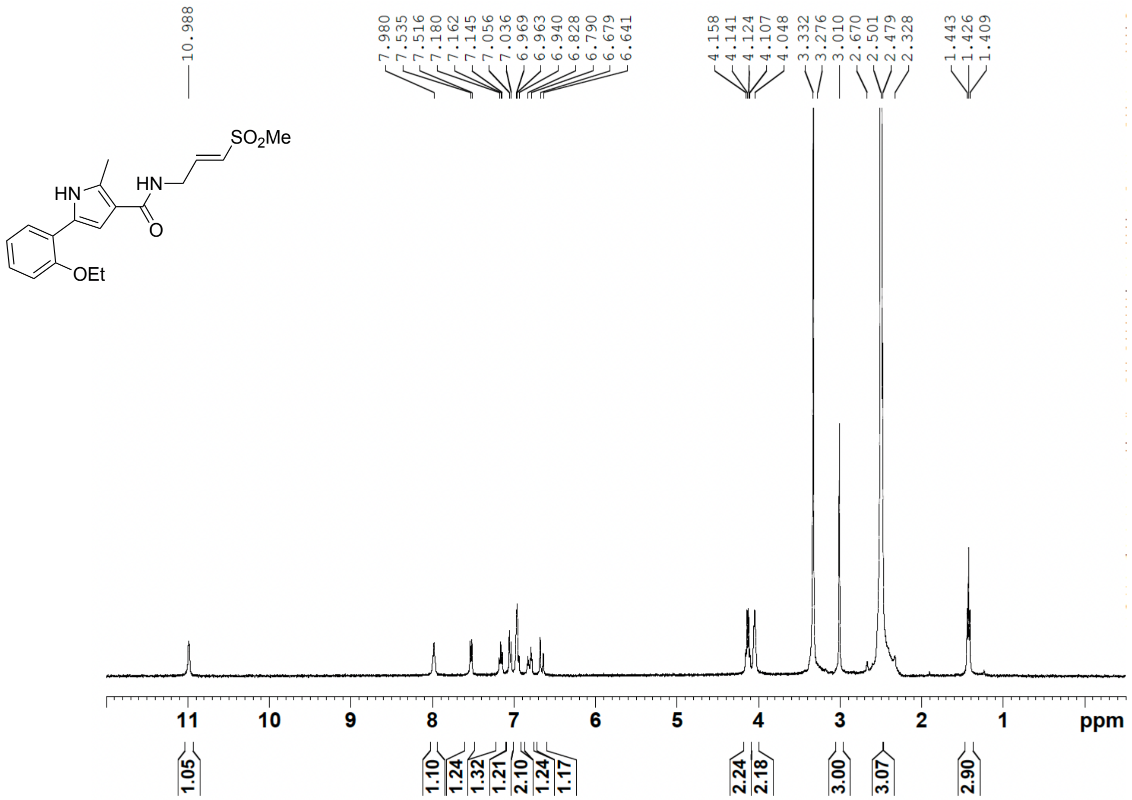

**Figure S101**: $^{13}$C NMR (100 MHz, DMSO-$d_6$) for **21**

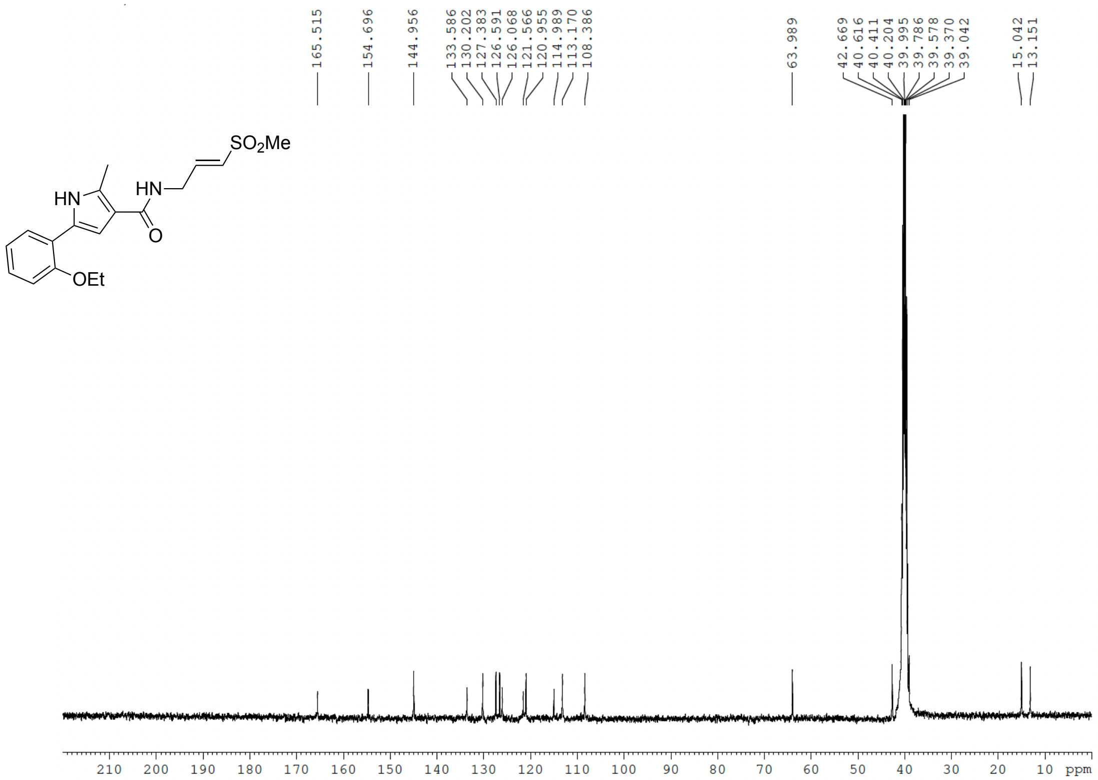

**Figure S102**: $^{1}$H NMR (400 MHz, DMSO-$d_6$) for **22**

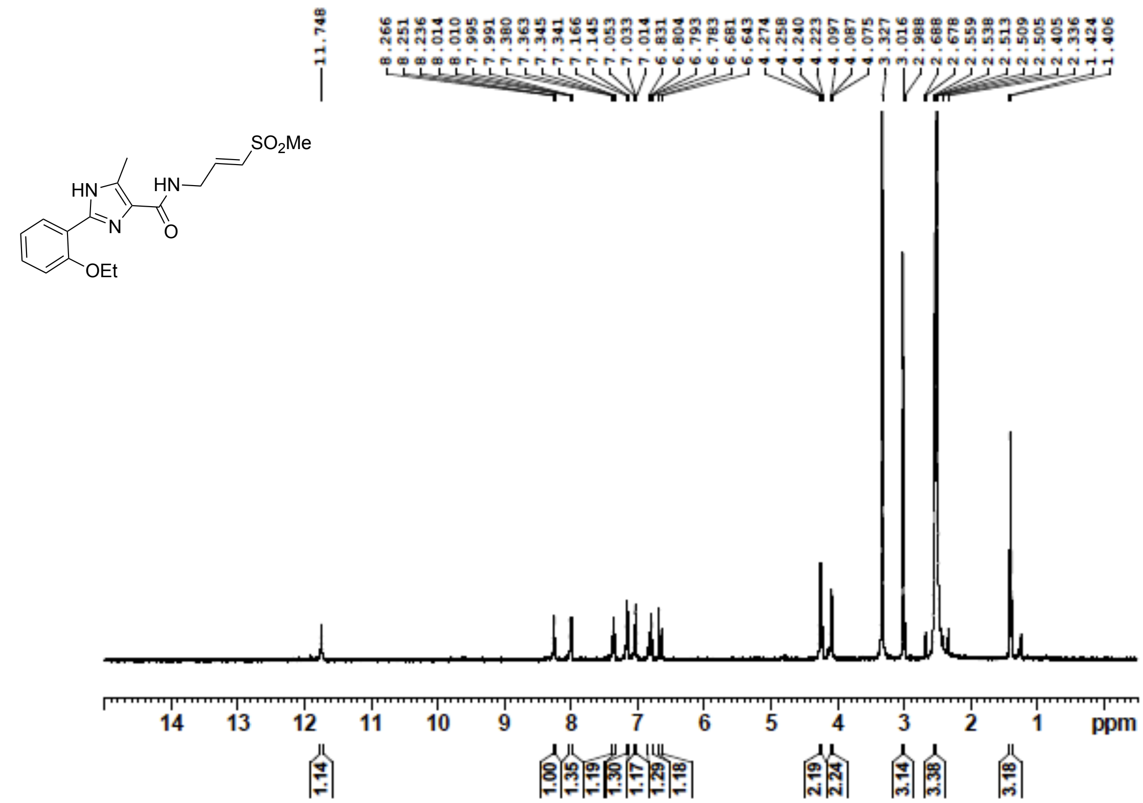

**Figure S103**: $^{13}$C NMR (100 MHz, DMSO-$d_6$) for **22**

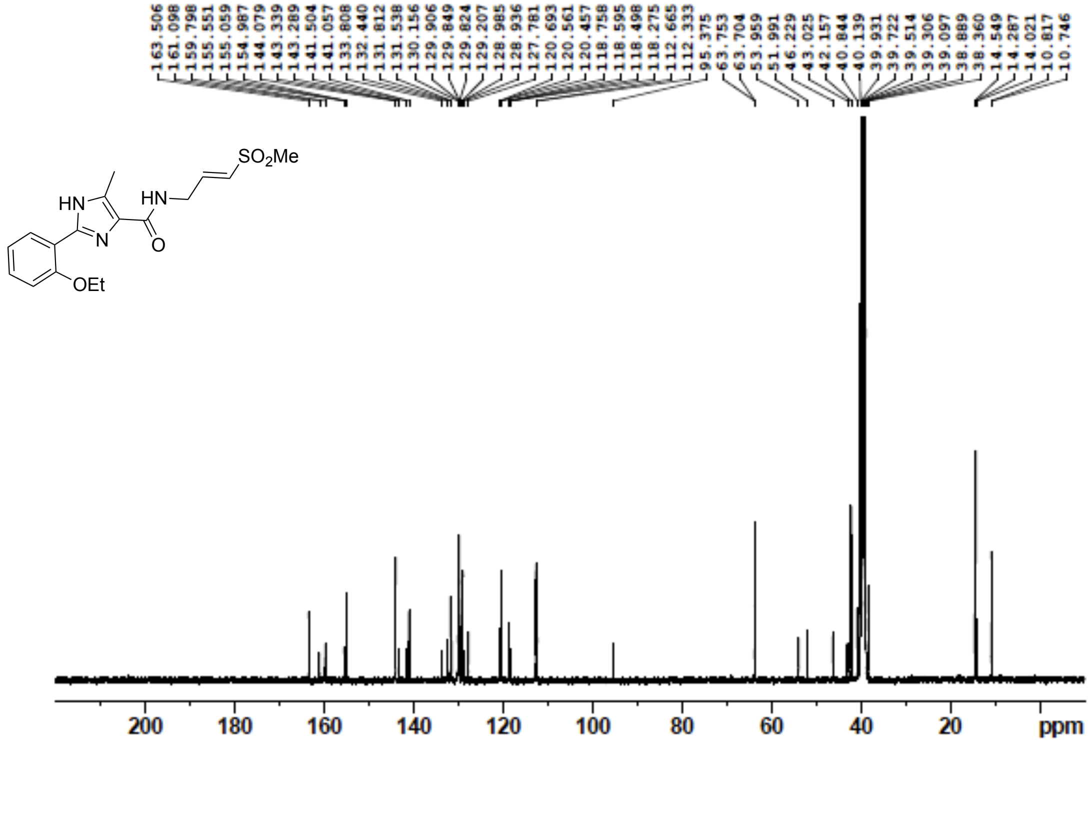

**Figure S104**: $^{1}$H NMR (400 MHz, DMSO-$d_6$) for **23a**

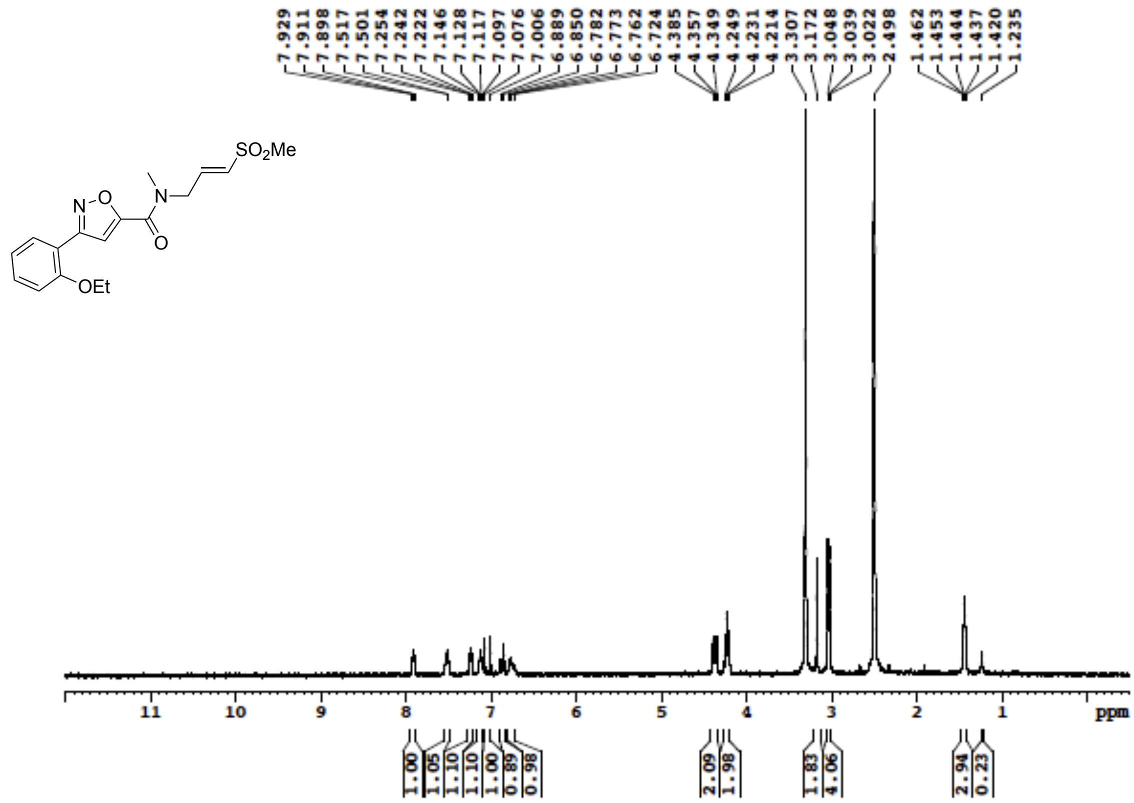

**Figure S105**: $^{13}$C NMR (100 MHz, DMSO-$d_6$) for **23a**

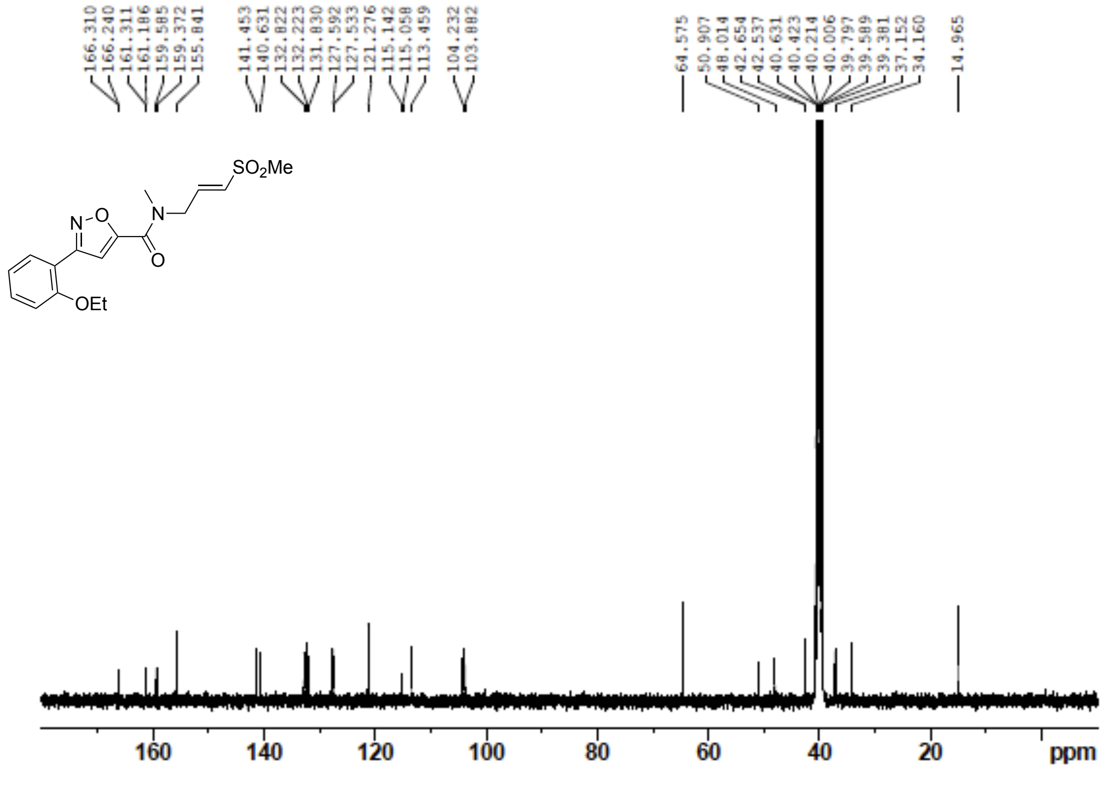

**Figure S106**: $^{1}$H NMR (400 MHz, DMSO-$d_6$) for **23b**

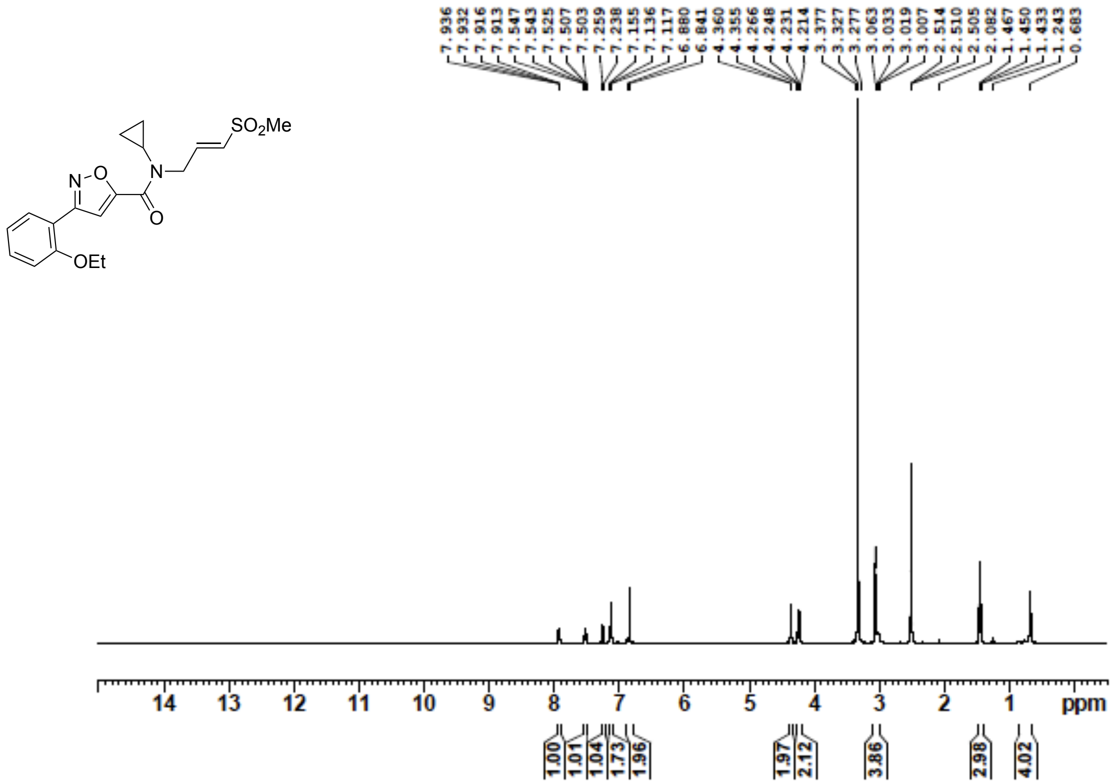


**Figure S107**: $^{13}$C NMR (100 MHz, DMSO-$d_6$) for **23b**

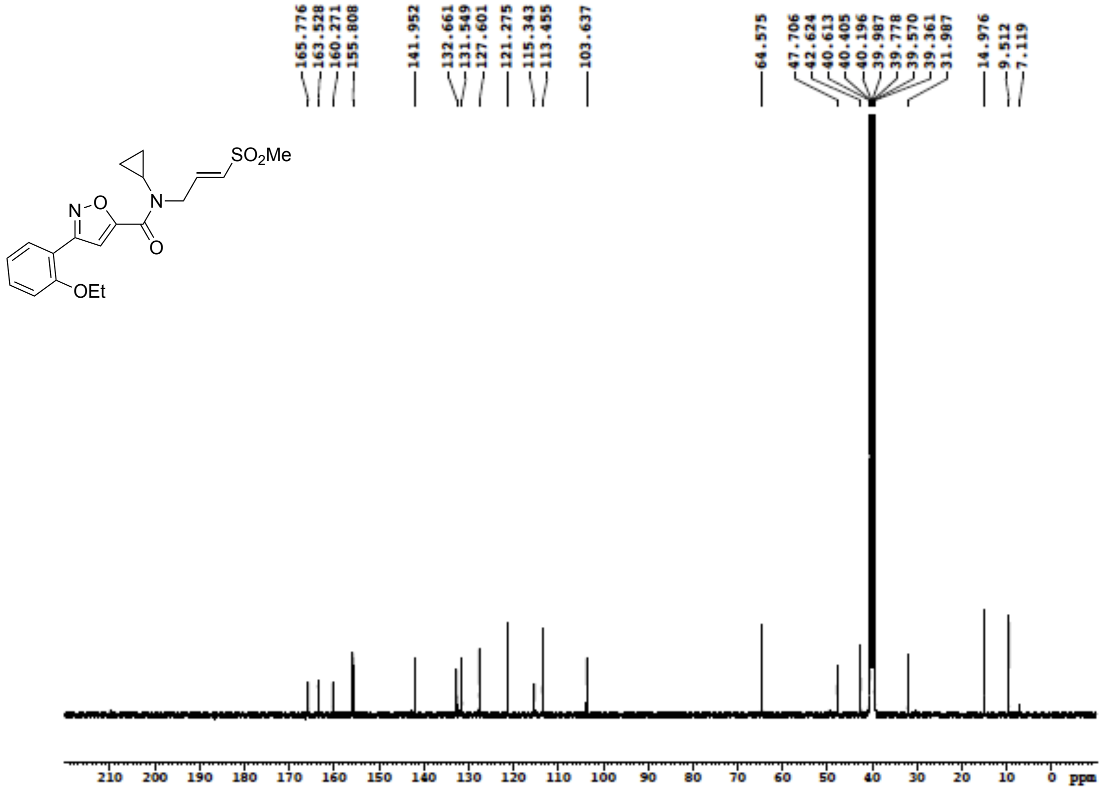

**Figure S108**: $^{1}$H NMR (400 MHz, DMSO-$d_6$) for **23c**

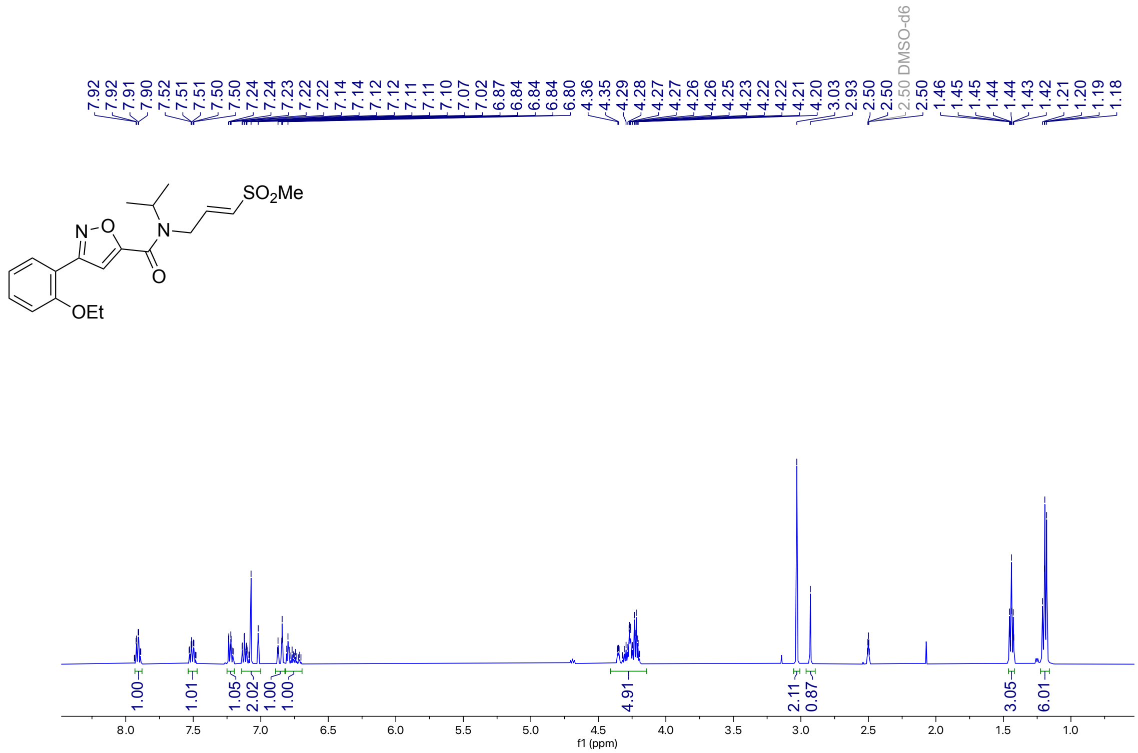

**Figure S109**: $^{13}$C NMR (100 MHz, DMSO-$d_6$) for **23c**

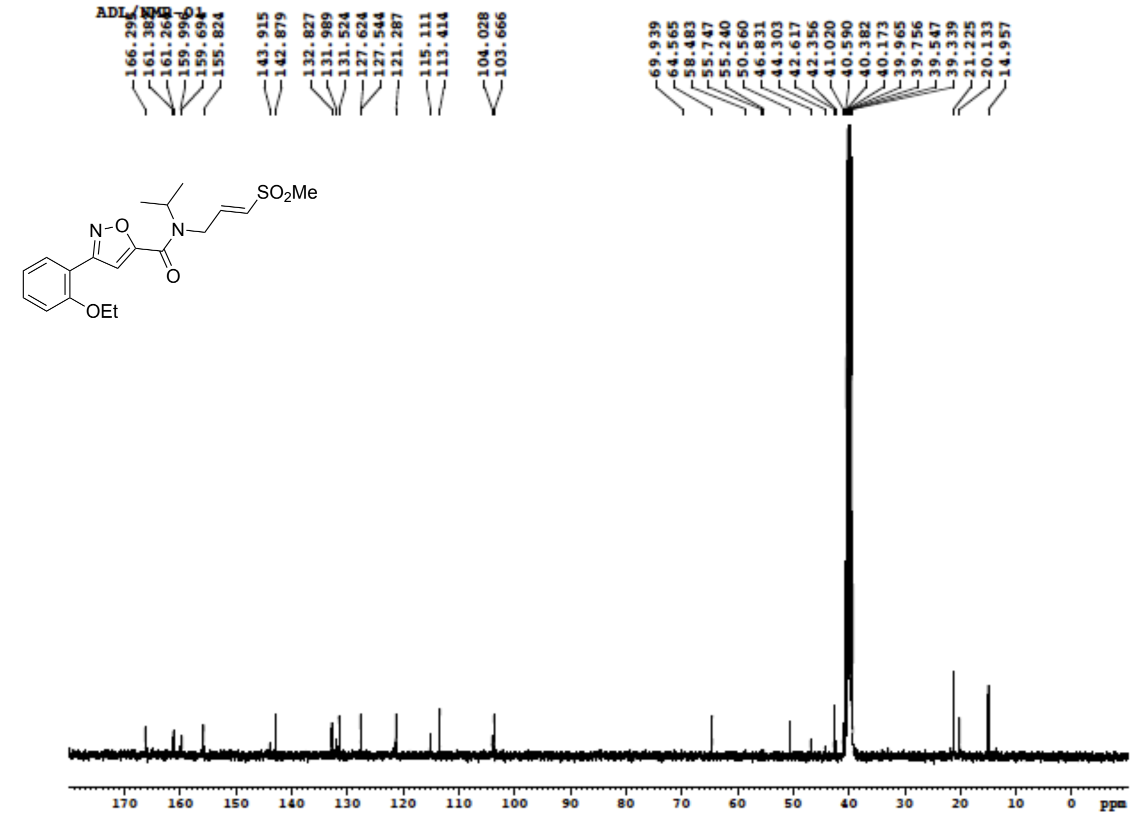

**Figure S110**: $^{1}$H NMR (400 MHz, DMSO-$d_6$) for **23d**

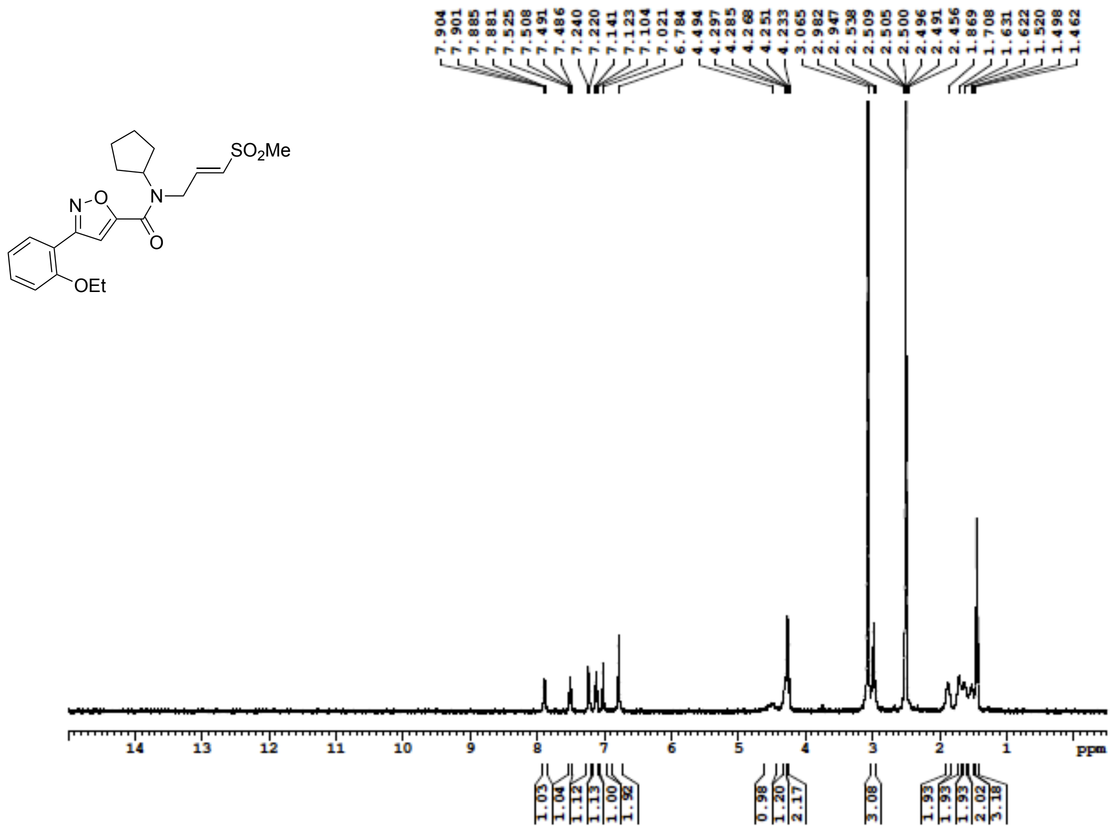

**Figure S111**: $^{13}$C NMR (100 MHz, DMSO-$d_6$) for **23d**

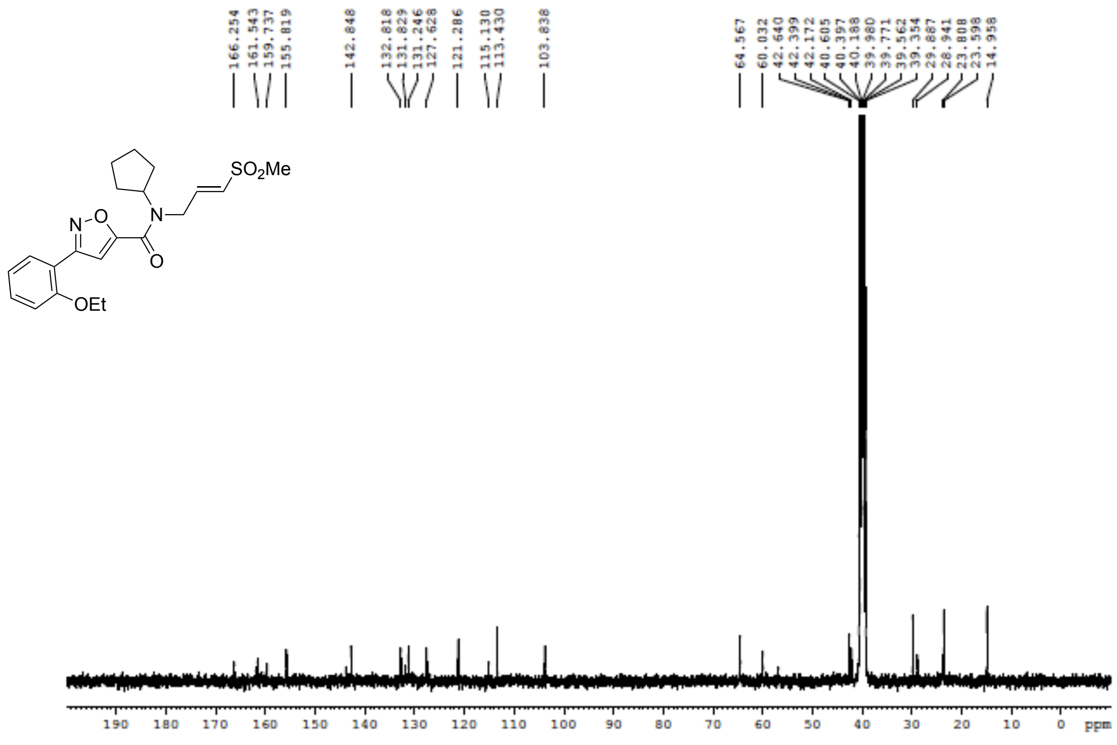

**Figure S112**: $^{1}$H NMR (400 MHz, DMSO-$d_6$) for **23e**

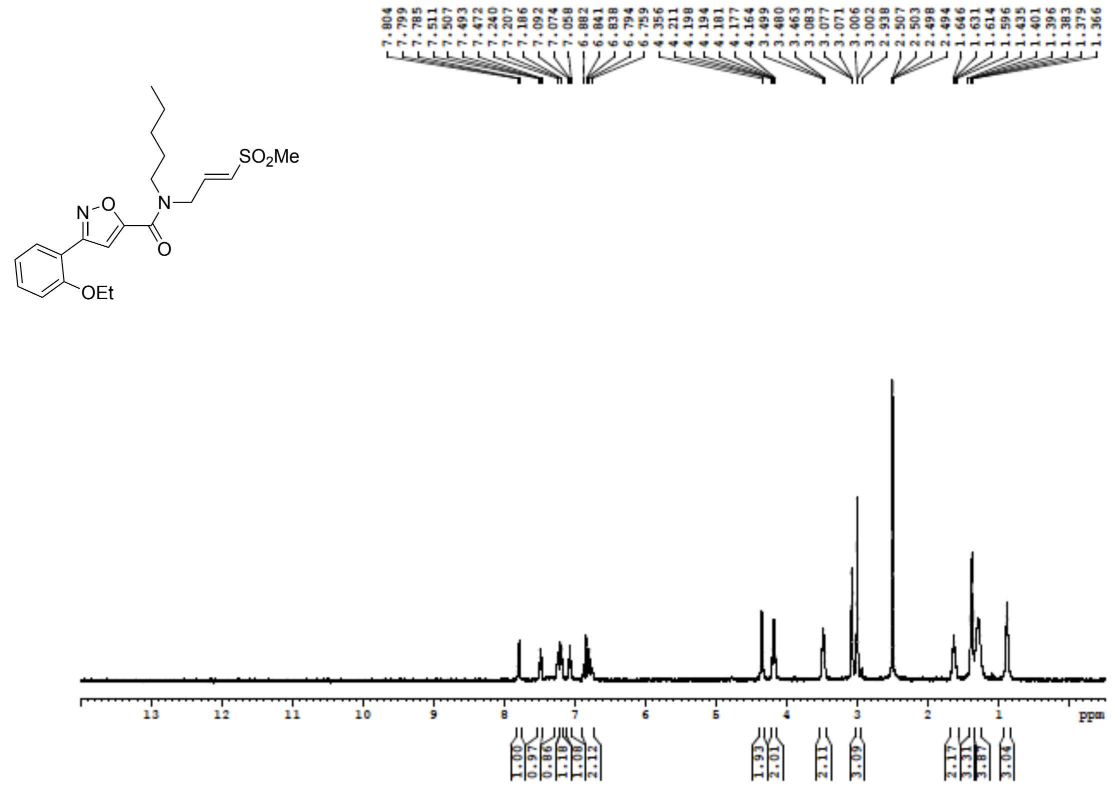

**Figure S113**: $^{13}$C NMR (100 MHz, DMSO-$d_6$) for **23e**

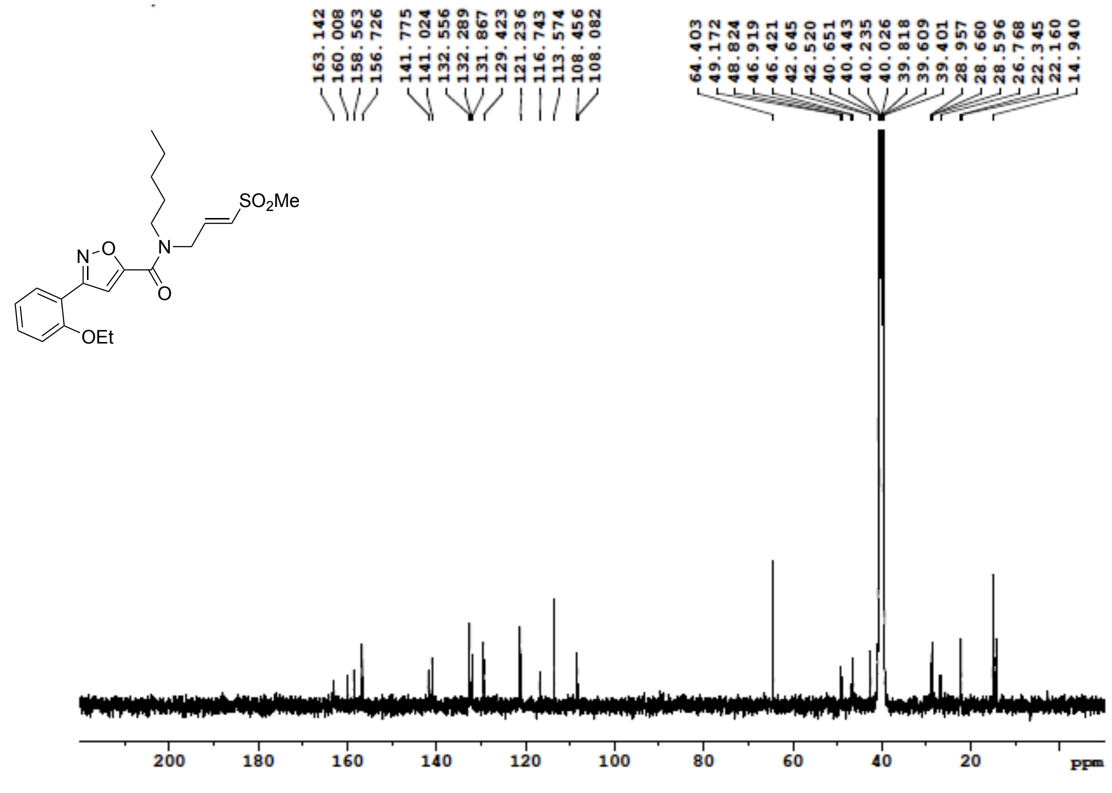

**Figure S114**: $^{1}$H NMR (400 MHz, DMSO-$d_6$) for **23f**

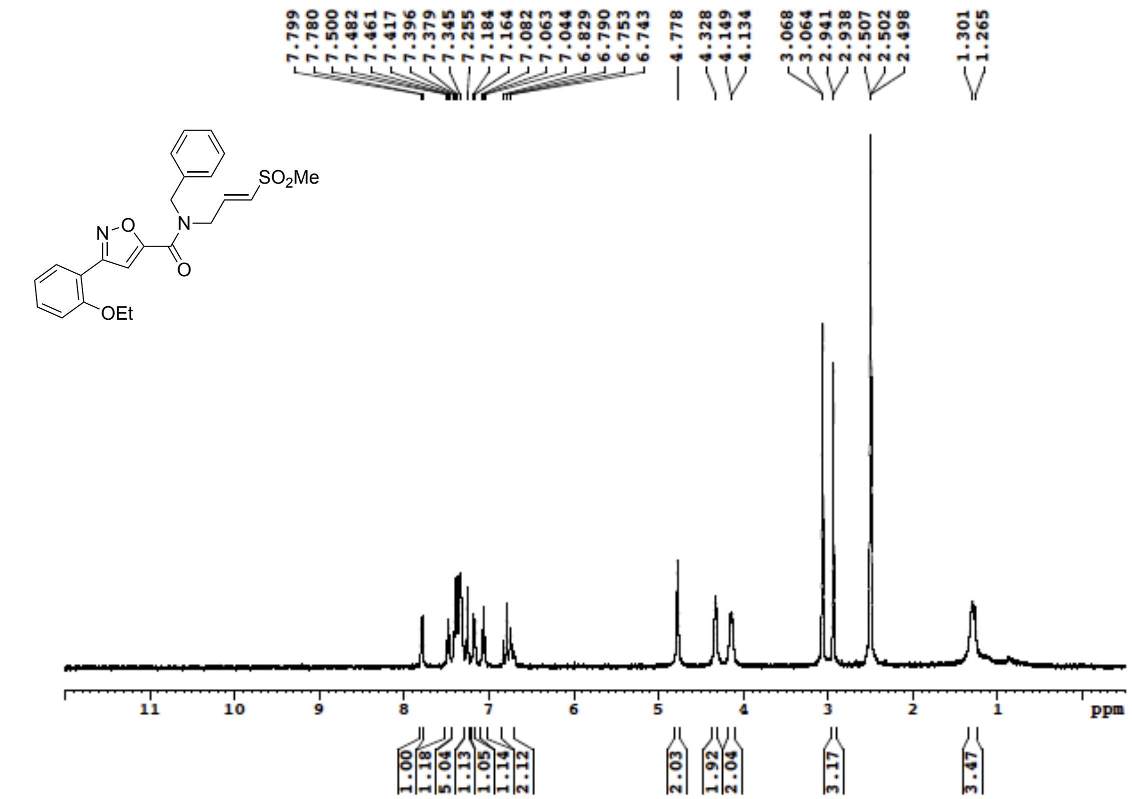


**Figure S115**: $^{13}$C NMR (100 MHz, DMSO-$d_6$) for **23f**

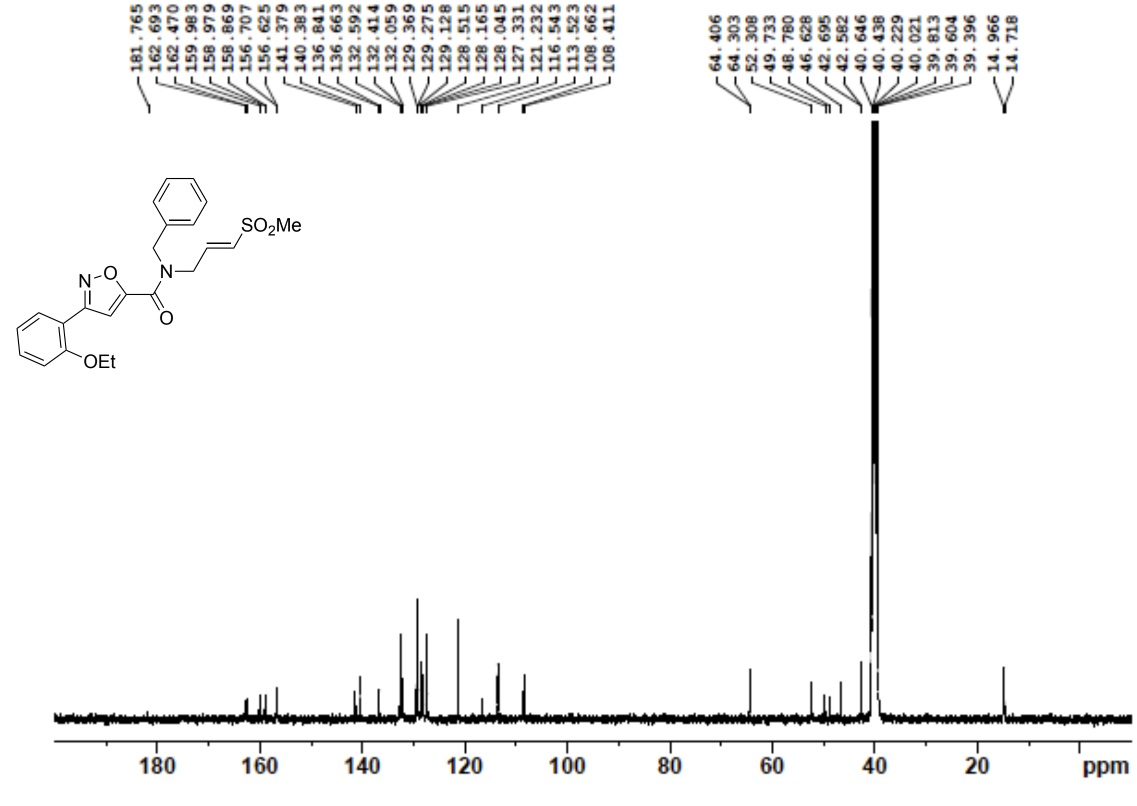

**Figure S116**: $^{1}$H NMR (400 MHz, DMSO-$d_6$) for **24a**

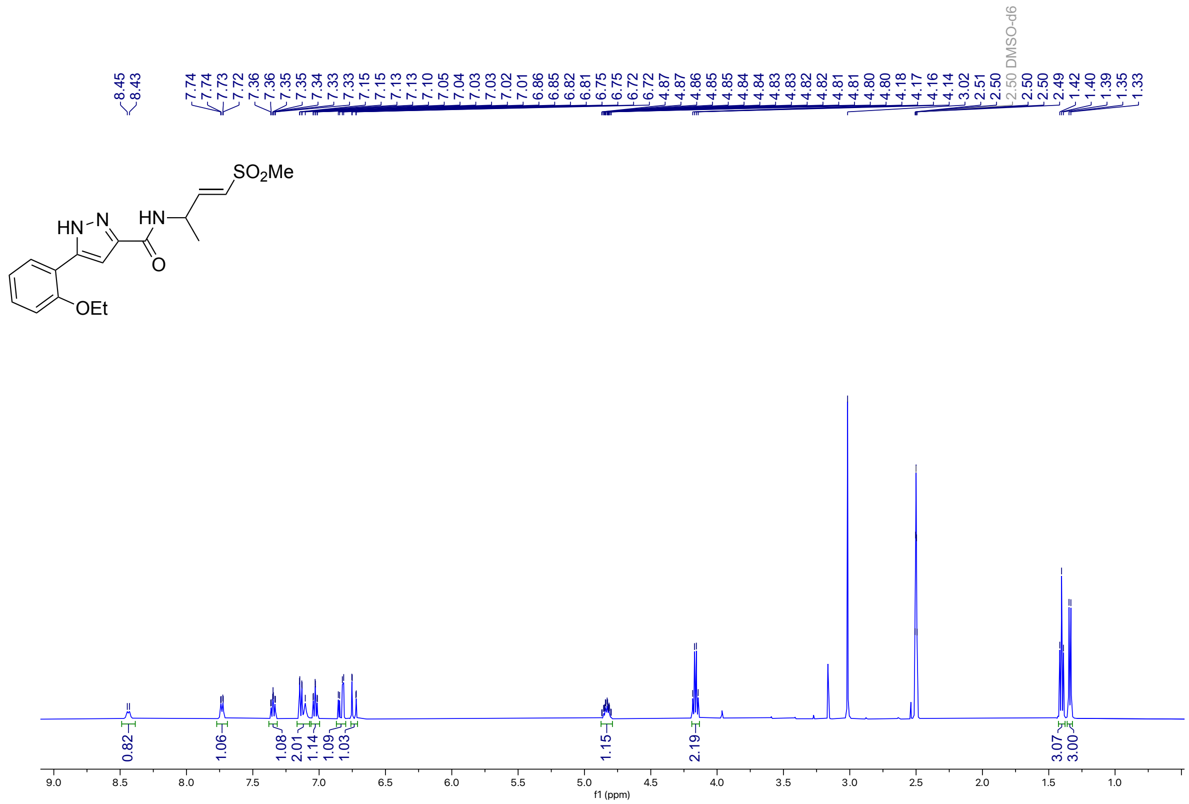

**Figure S117**: $^{13}$C NMR (100 MHz, DMSO-$d_6$) for **24a**

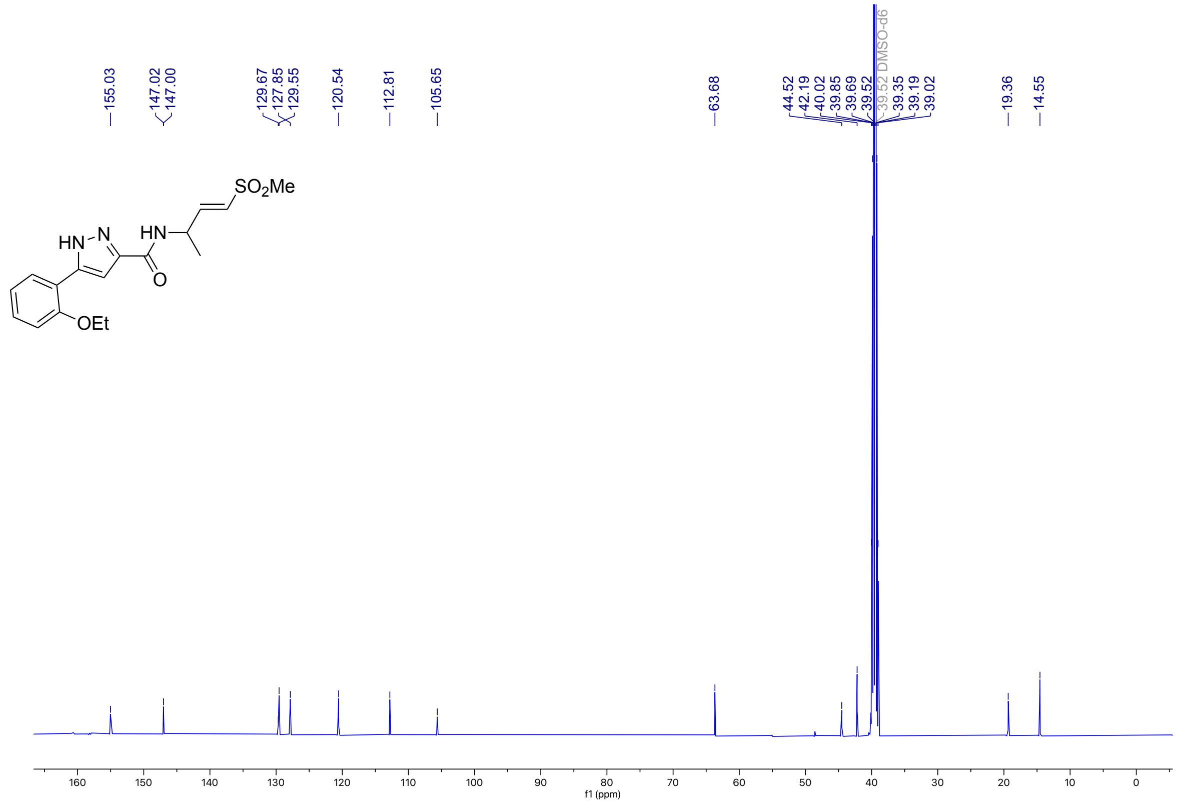

**Figure S118**: $^{1}$H NMR (400 MHz, DMSO-$d_6$) for **24b**

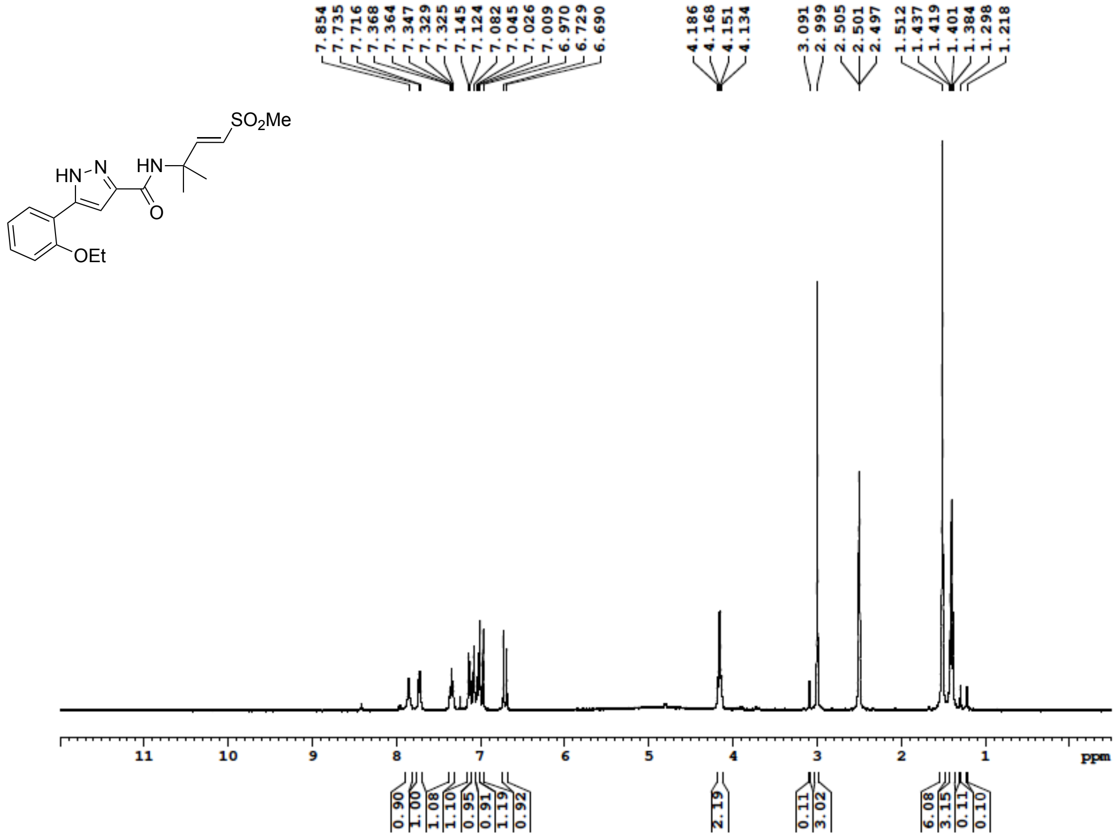

**Figure S119**: $^{13}$C NMR (100 MHz, DMSO-$d_6$) for **24b**

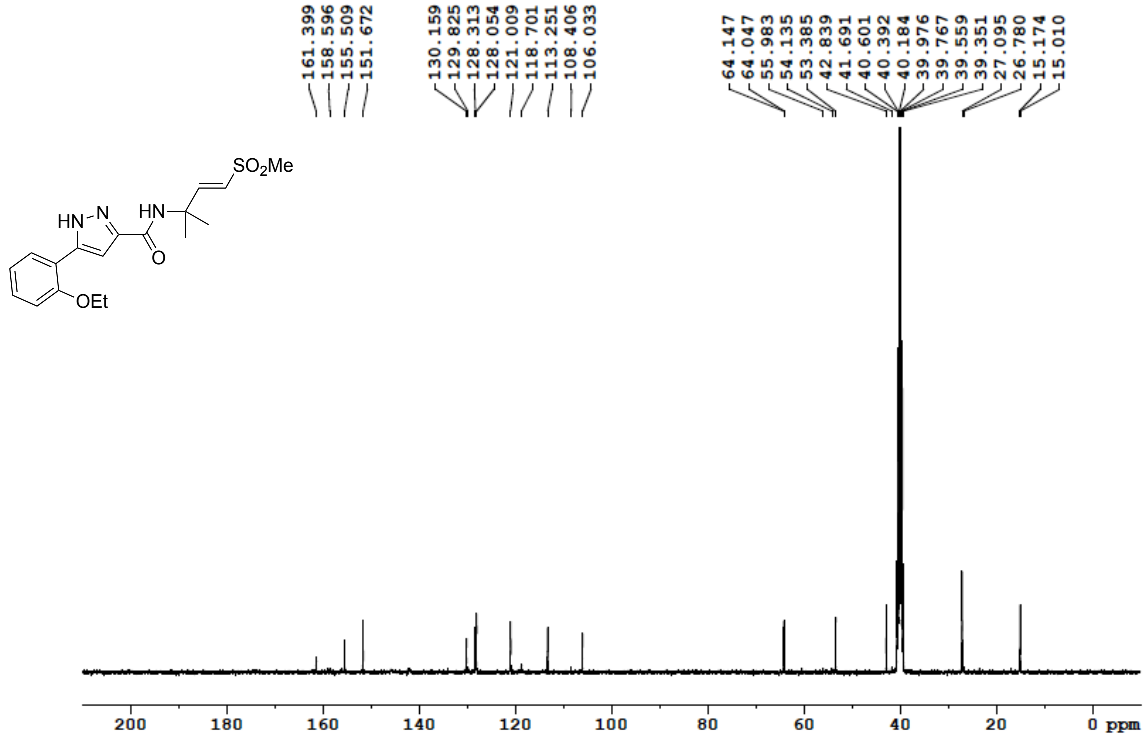

**Figure S120**: $^{1}$H NMR (500 MHz, MeOD) for **24c**

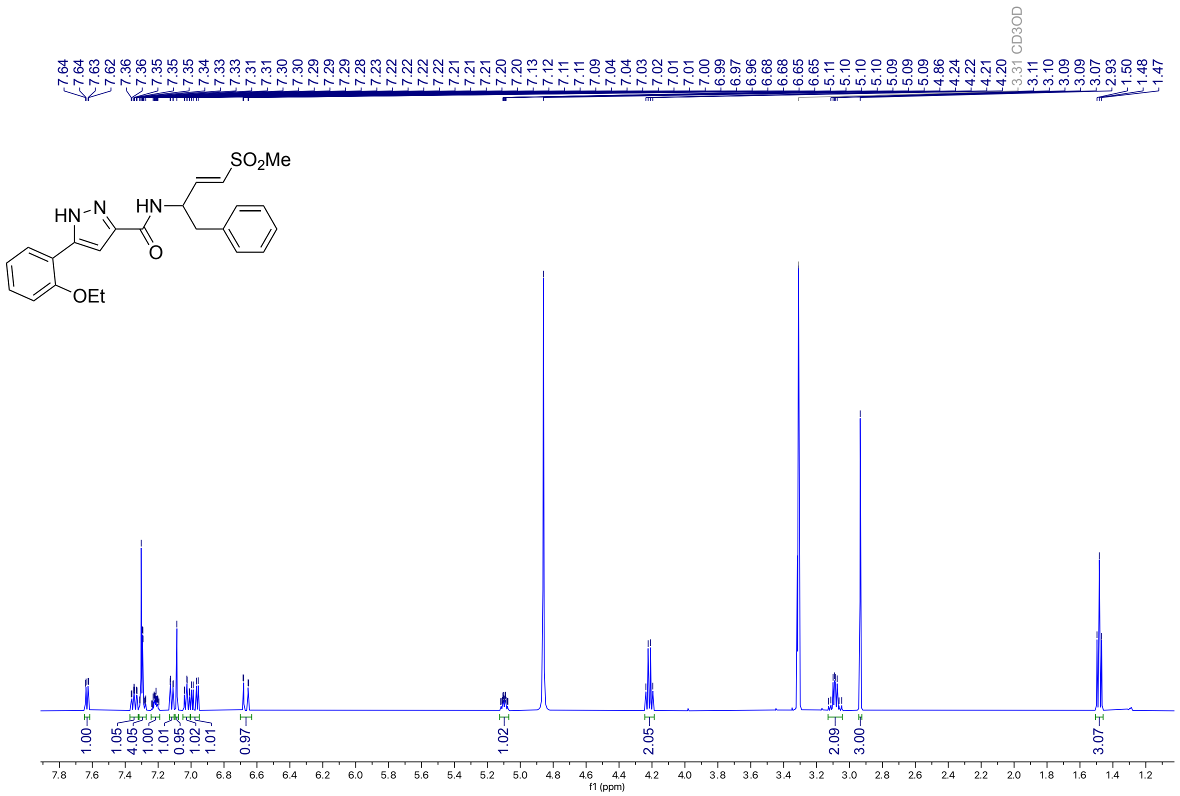


**Figure S121**: $^{13}$C NMR (126 MHz, MeOD) for **24c**

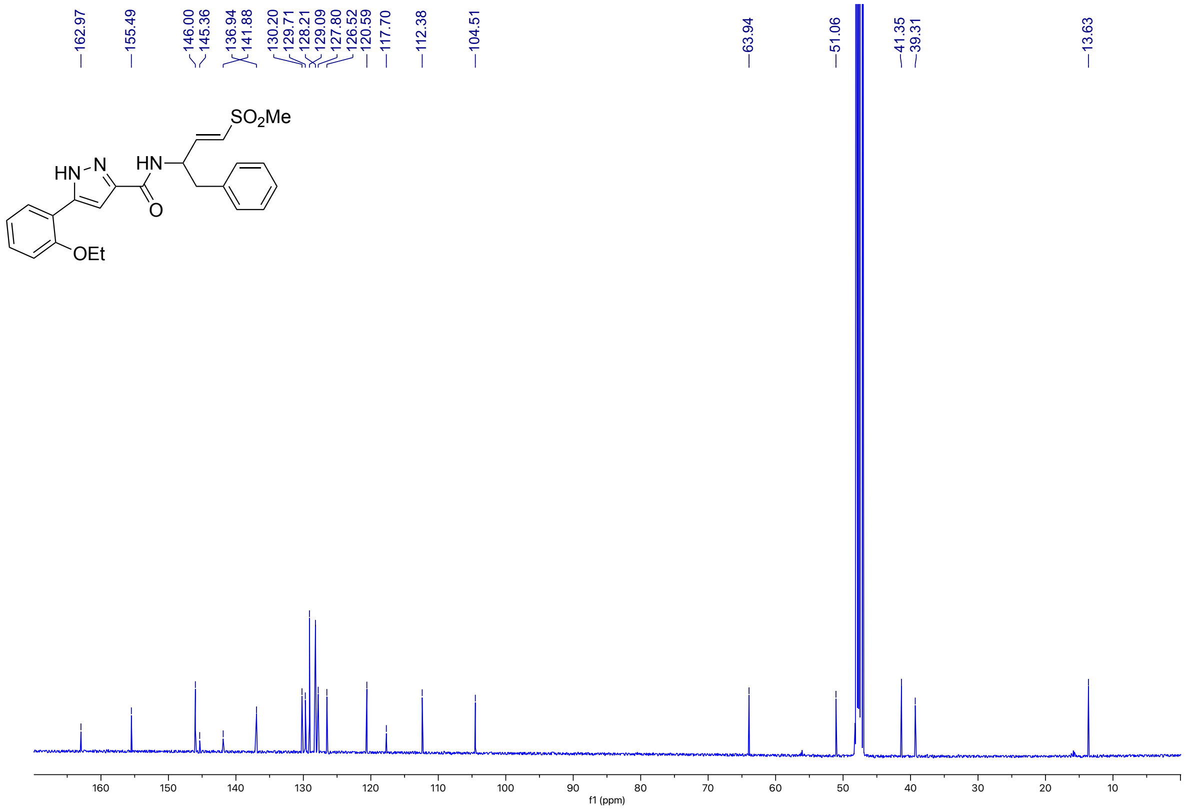

**Figure S122**: $^1$H NMR (400 MHz, DMSO-$d_6$) for **24d**

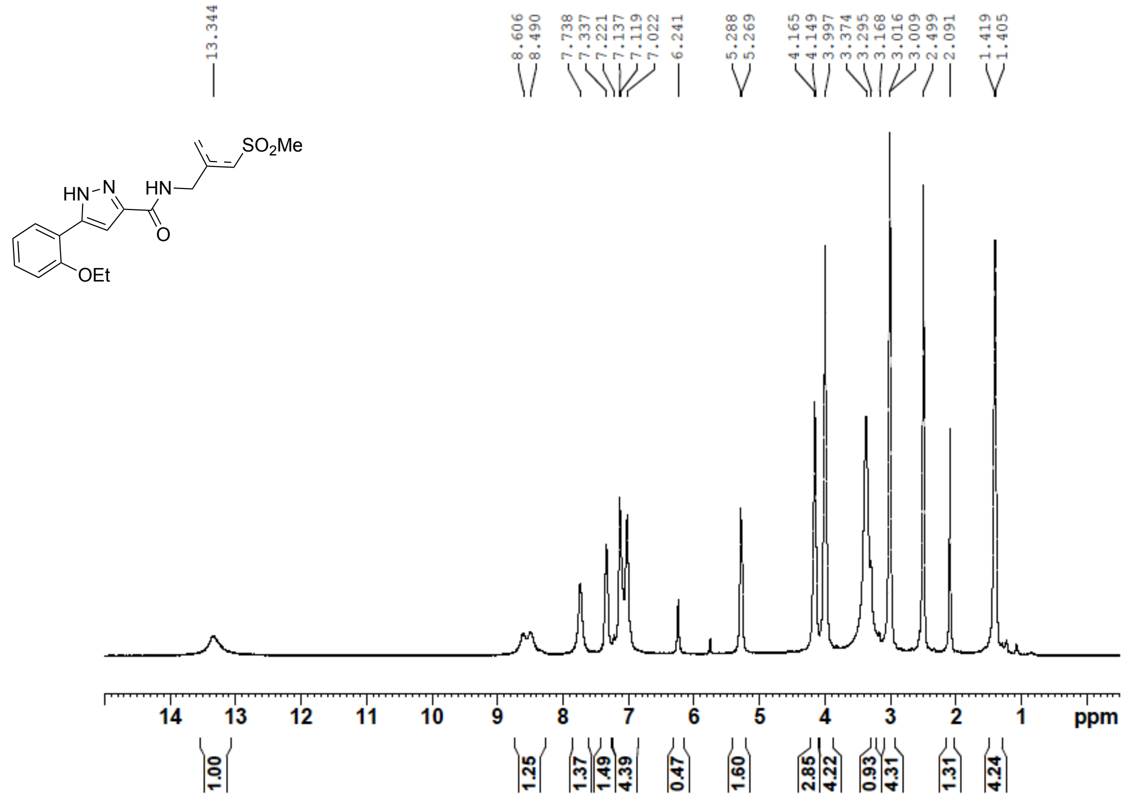

**Figure S123**: $^{13}$C NMR (100 MHz, DMSO-$d_6$) for **24d**

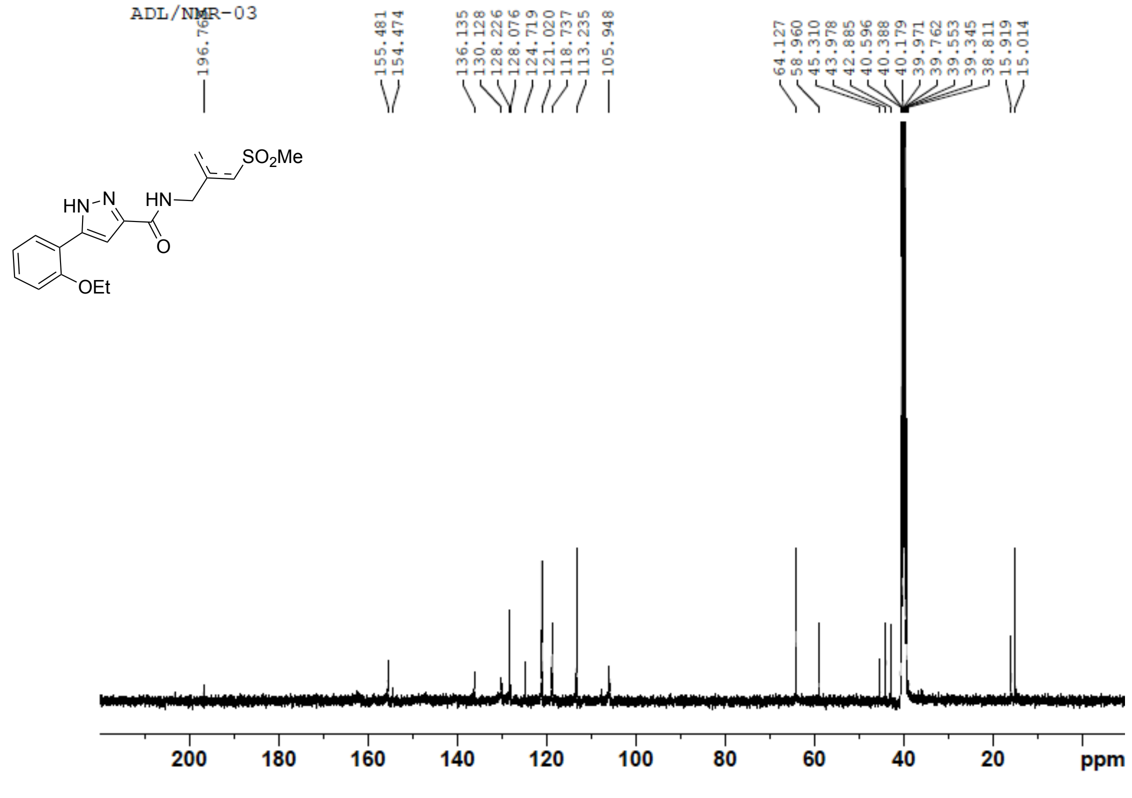

**Figure S124**: $^{1}$H NMR (400 MHz, DMSO-$d_6$) for **24**e

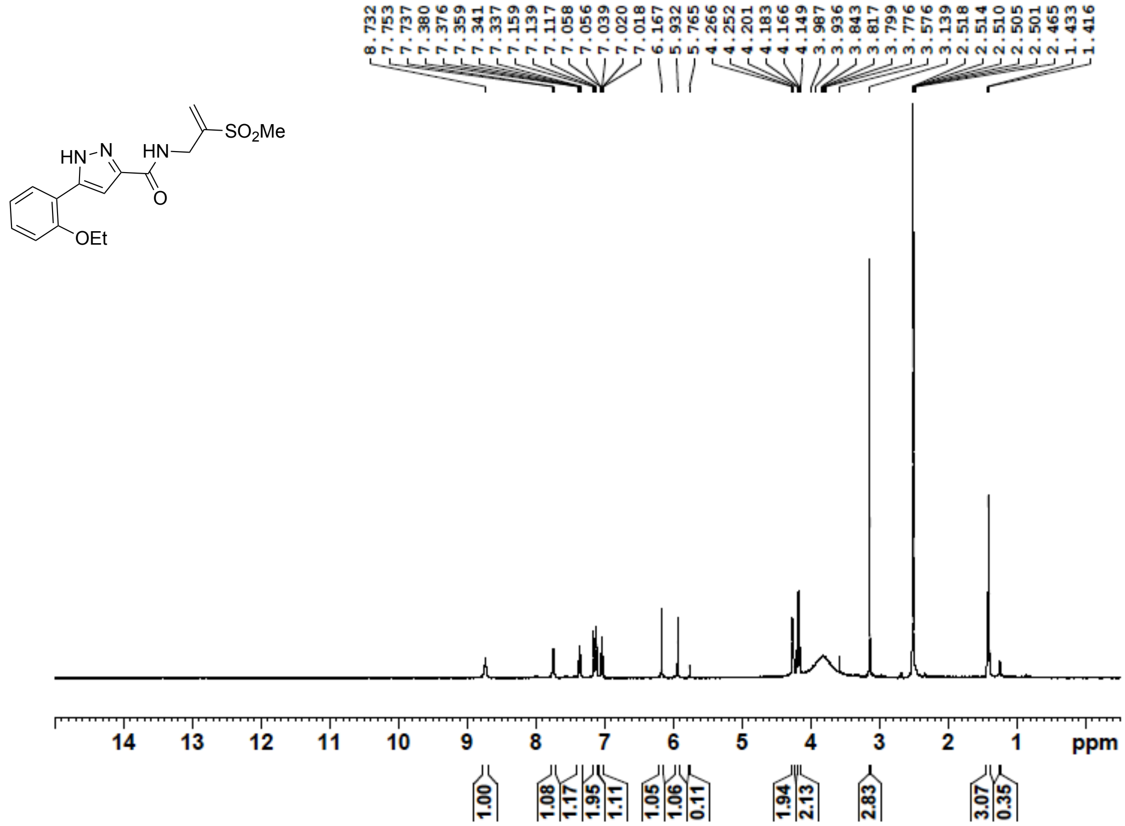


**Figure S125**: $^{13}$C NMR (100 MHz, DMSO-$d_6$) for **24e**

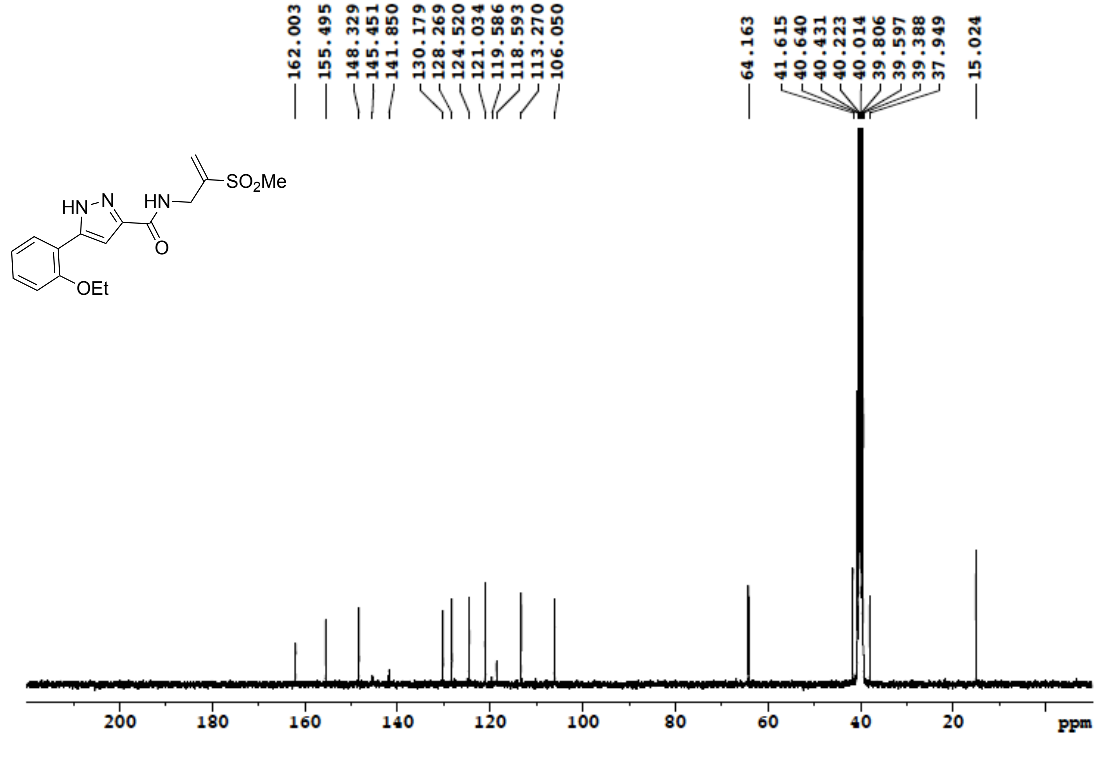

**Figure S126**: $^{1}$H NMR (400 MHz, DMSO-$d_6$) for **24f**

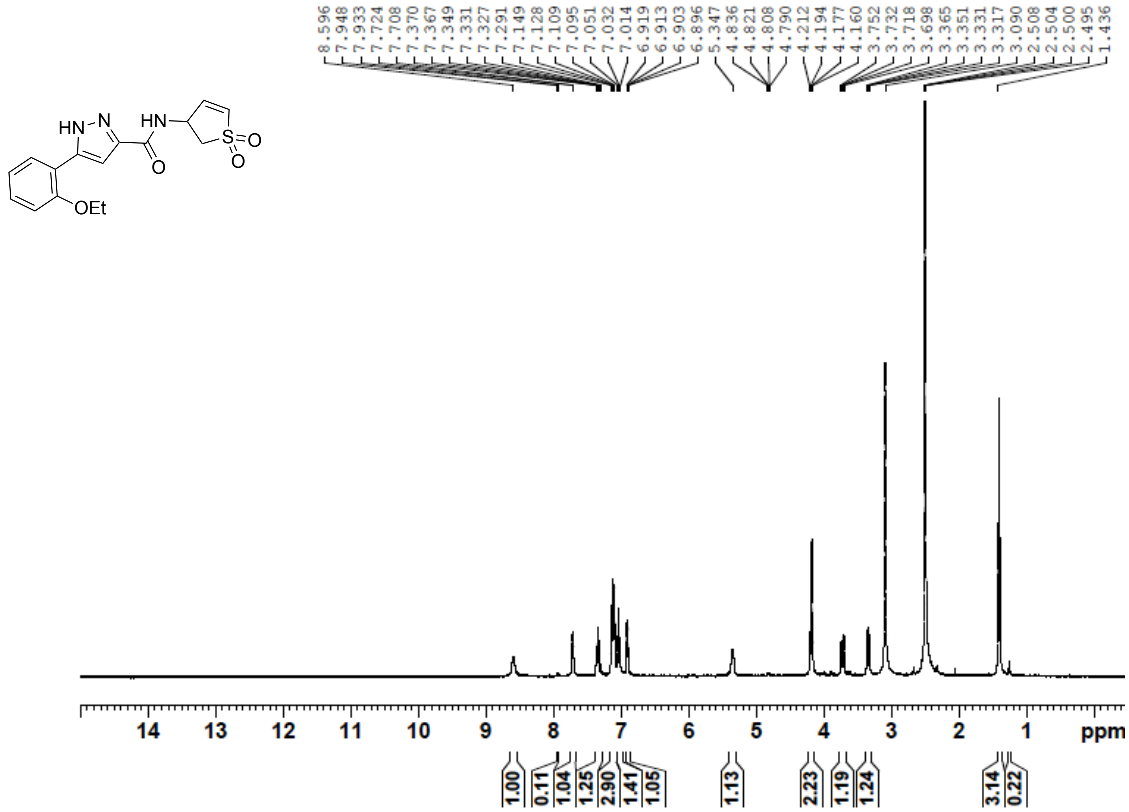

**Figure S127**: $^{13}$C NMR (100 MHz, DMSO-$d_6$) for **24f**

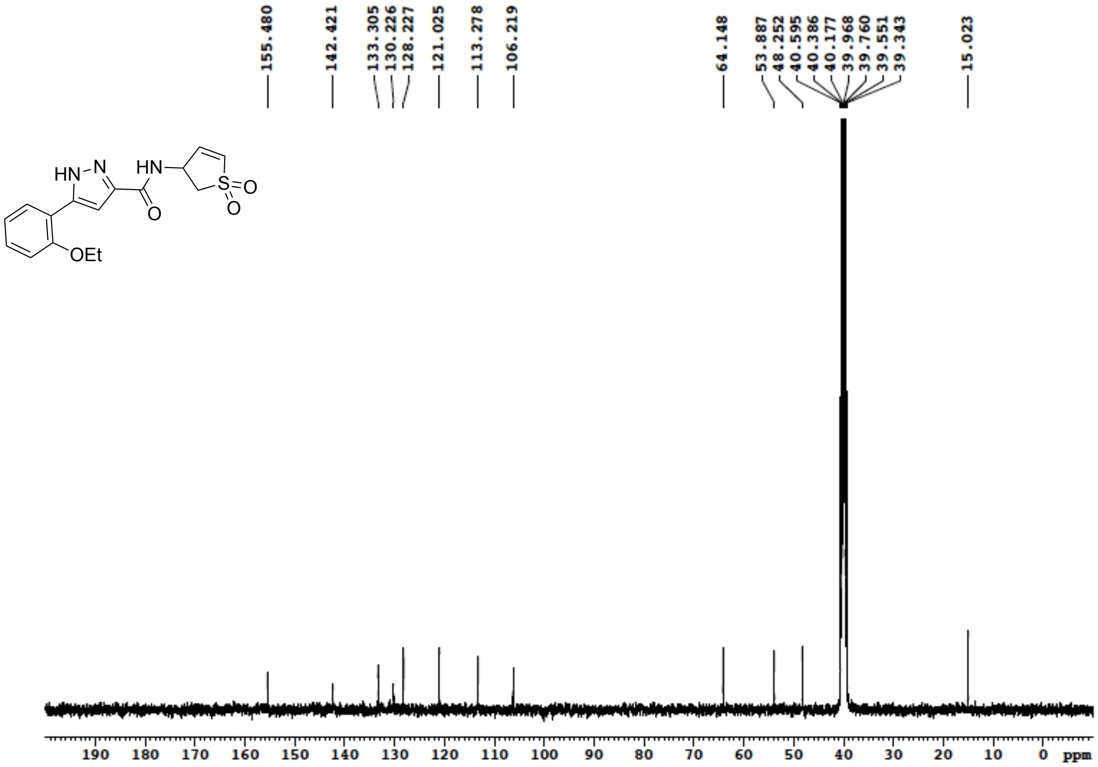

**Figure S128**: $^{1}$H NMR (400 MHz, DMSO-$d_6$) for **25a**

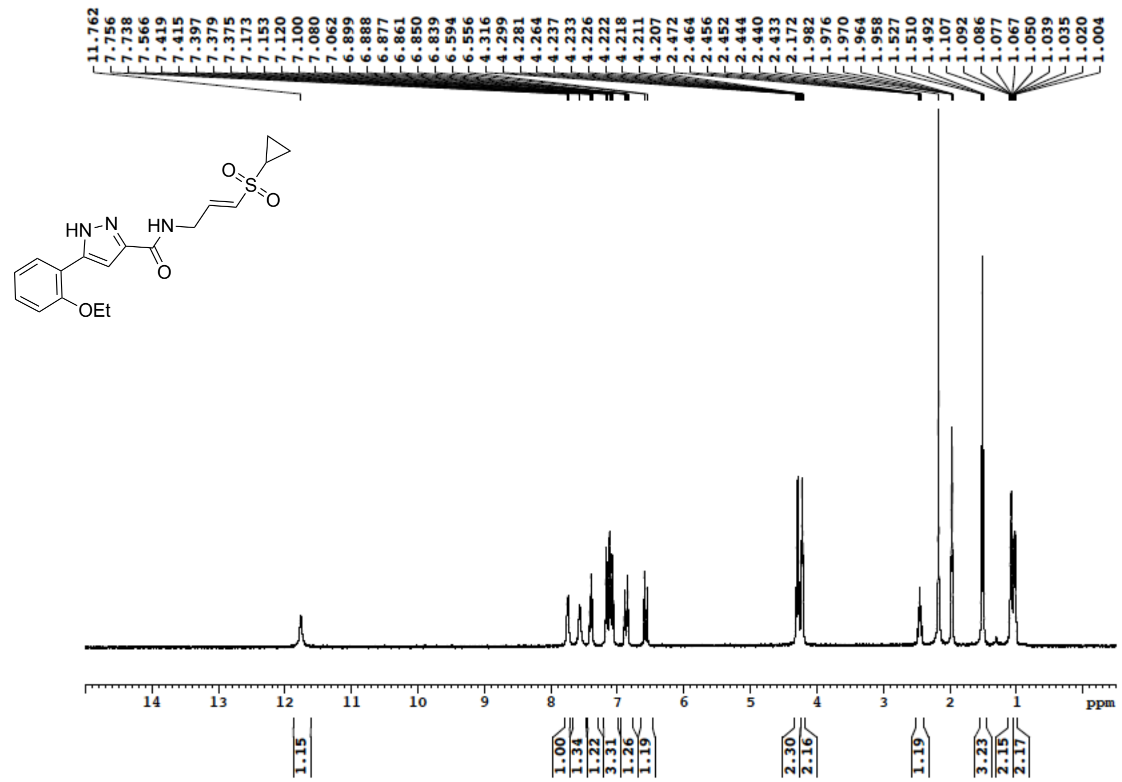


**Figure S129**: $^{13}$C NMR (100 MHz, DMSO-$d_6$) for **25a**

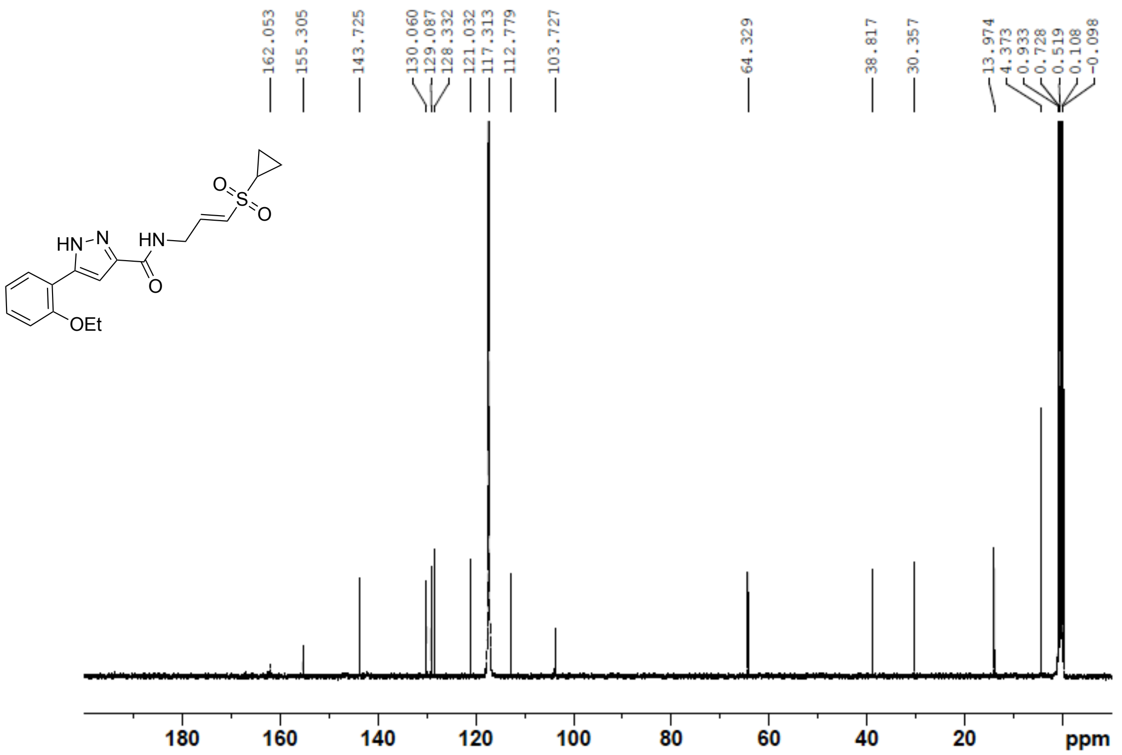

**Figure S130**: $^{1}$H NMR (400 MHz, DMSO-$d_6$) for **25b**

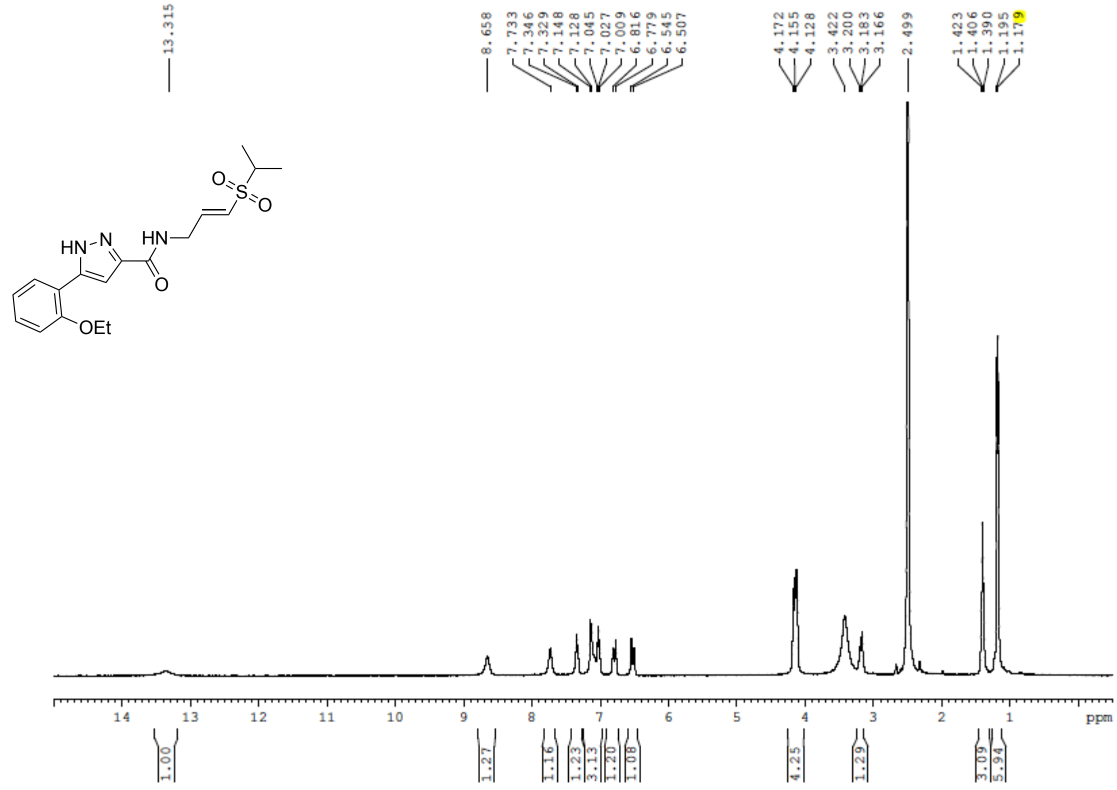

**Figure S131**: $^{13}$C NMR (100 MHz, DMSO-$d_6$) for **25b**

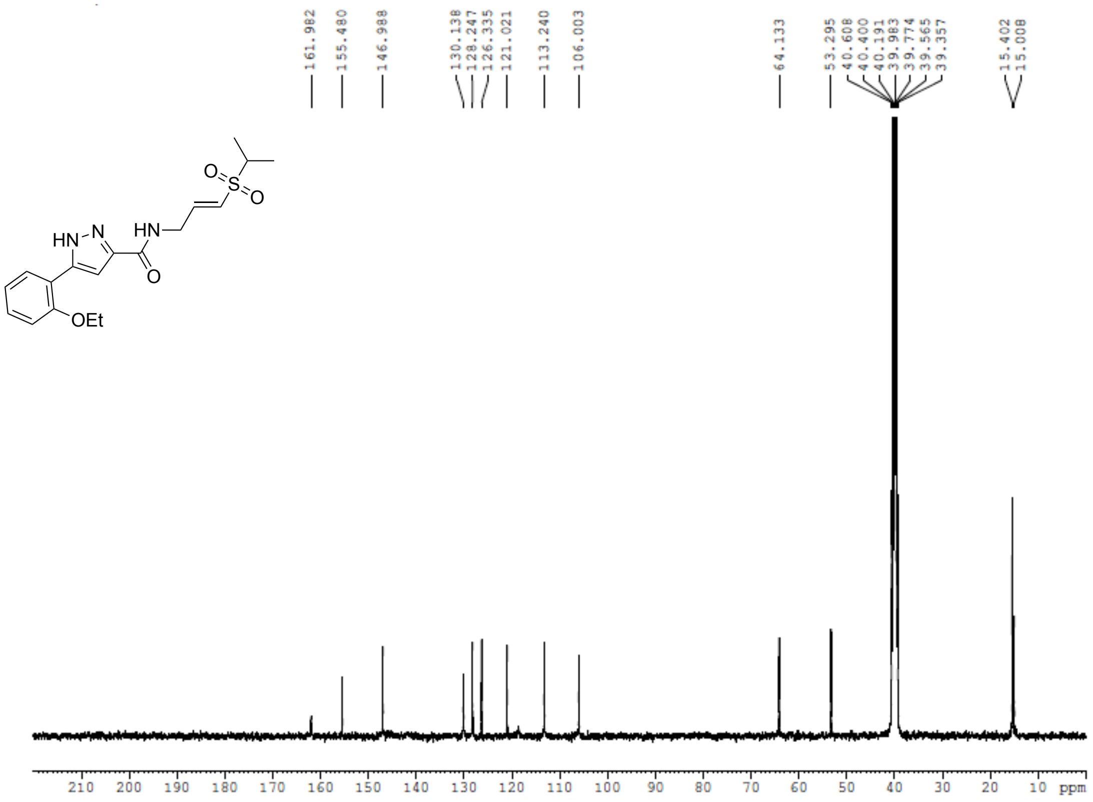

**Figure S132**: $^{1}$H NMR (400 MHz, DMSO-$d_6$) for **25c**

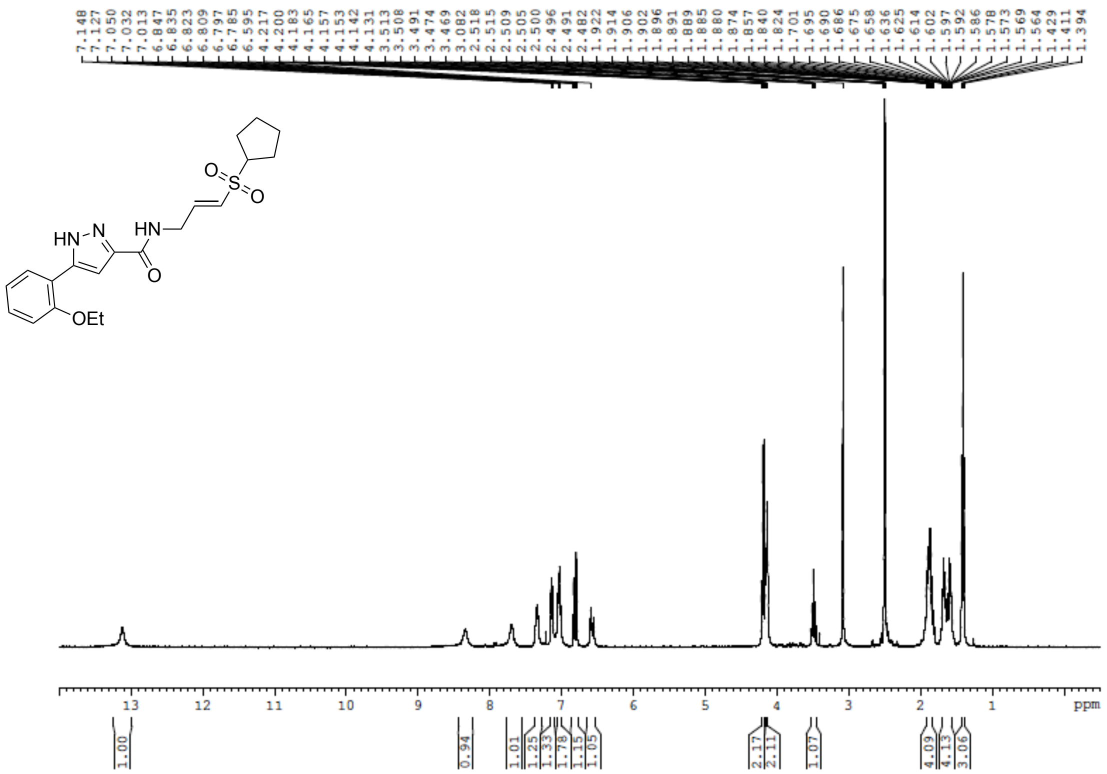

**Figure S133**: $^{13}$C NMR (100 MHz, DMSO-$d_6$) for **25c**

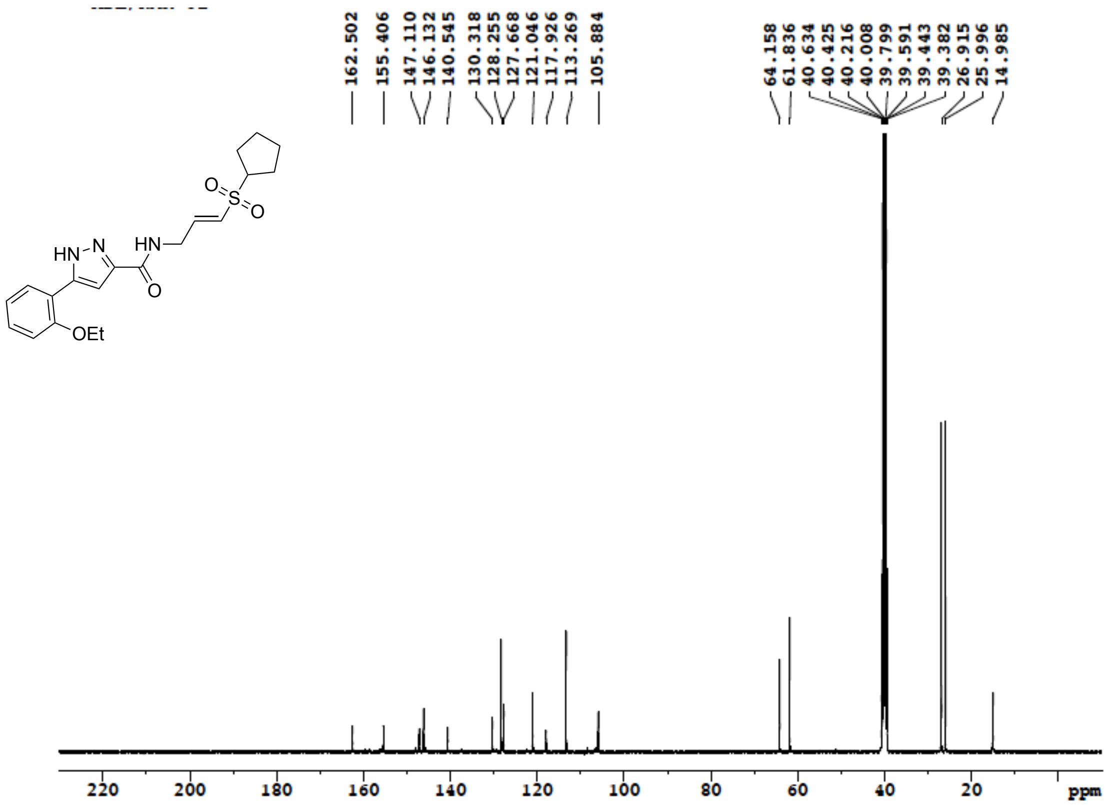

**Figure S134**: [1]H NMR (500 MHz, DMSO-$d_6$) for **25d**

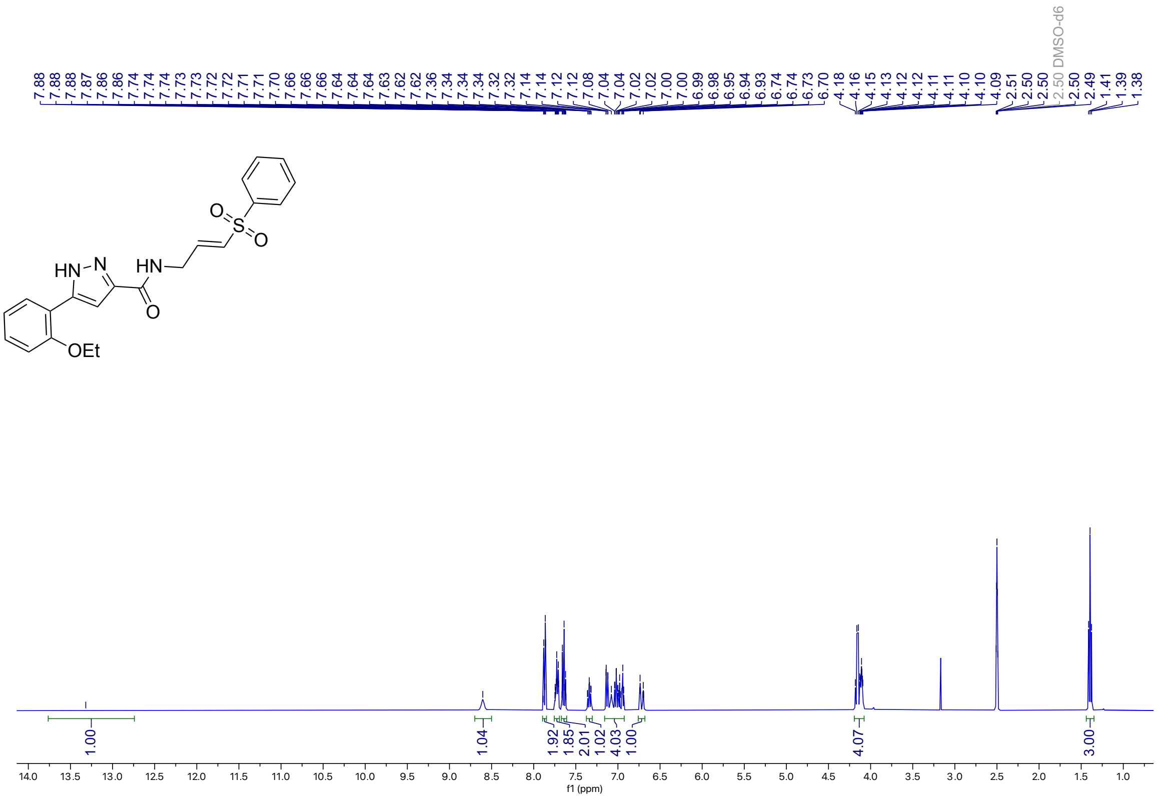

**Figure S135**: [13]C NMR (126 MHz, DMSO-$d_6$) for **25d**

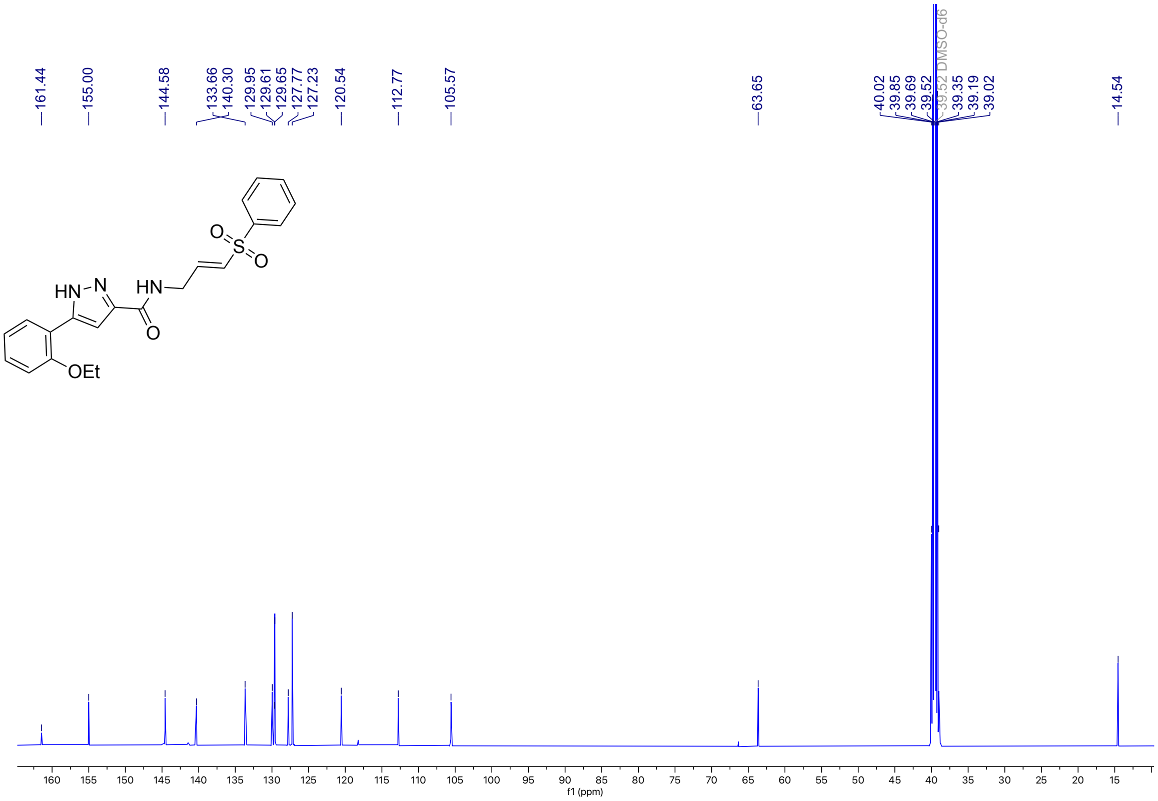

**Figure S136**: [1]H NMR (400 MHz, DMSO-$d_6$) for **25e**

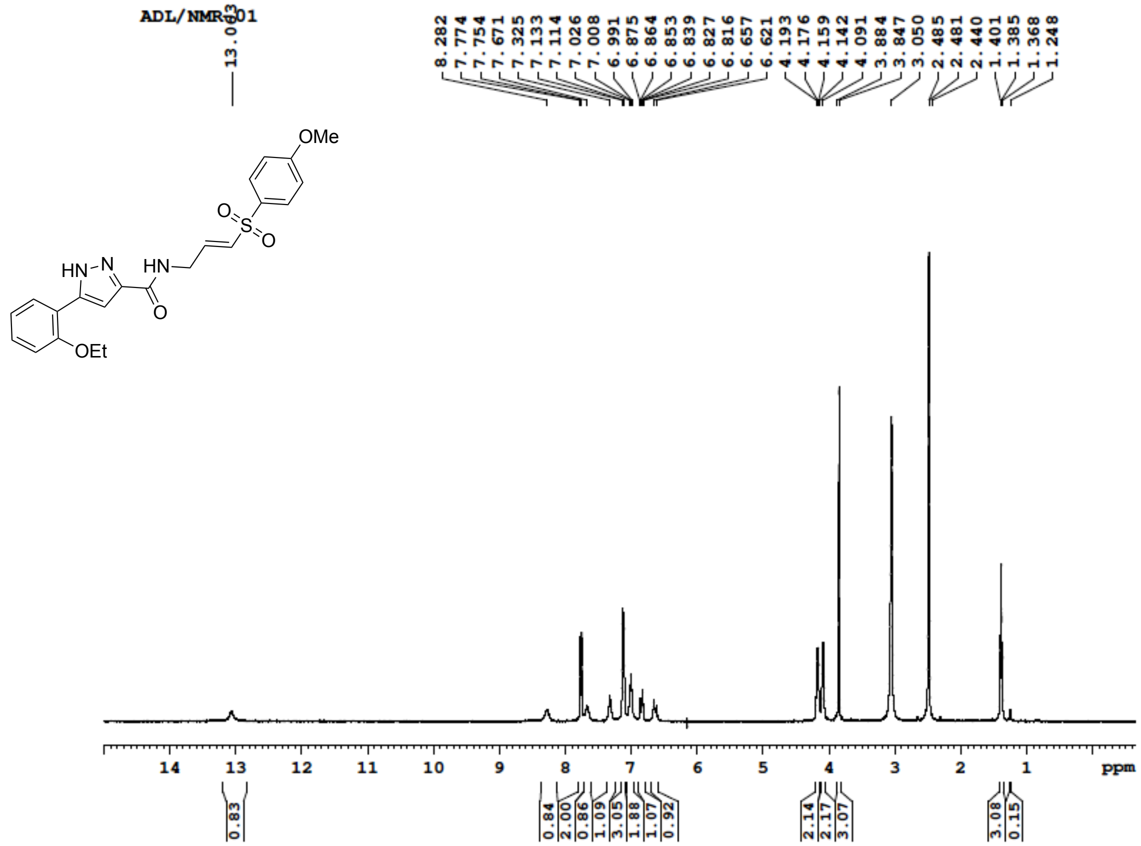


**Figure S137**: [13]C NMR (100 MHz, DMSO-$d_6$) for **25e**

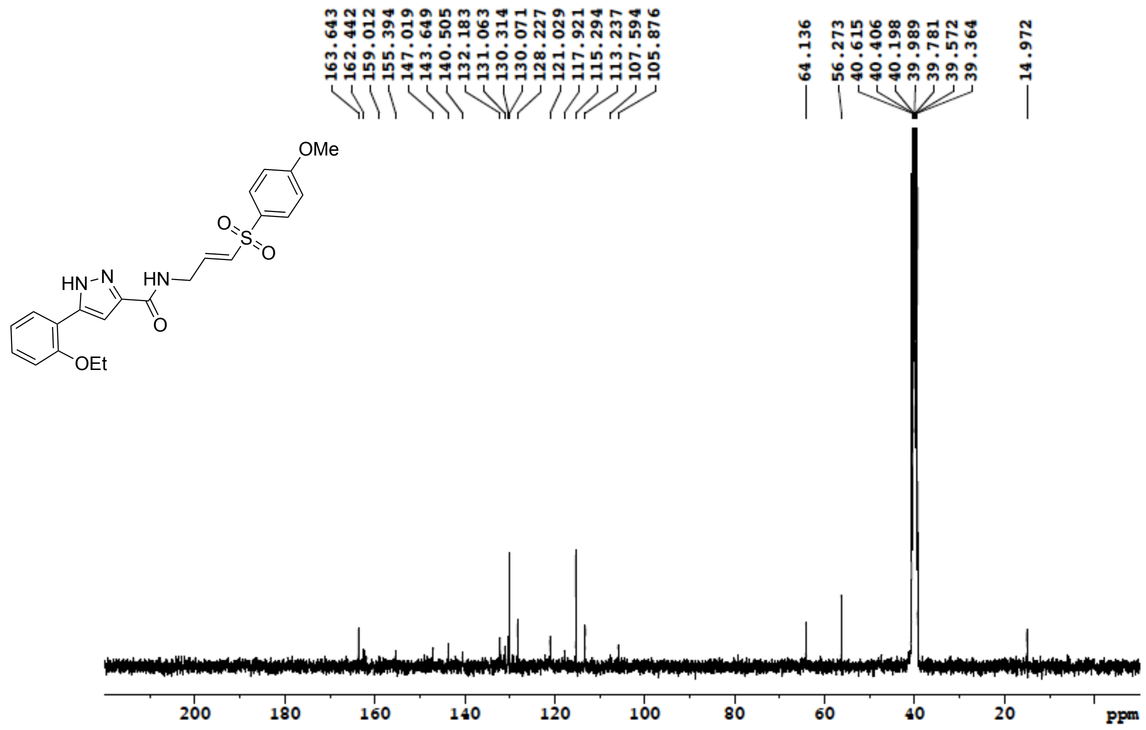

**Figure S138**: $^{1}$H NMR (500 MHz, MeOD) for **25f**

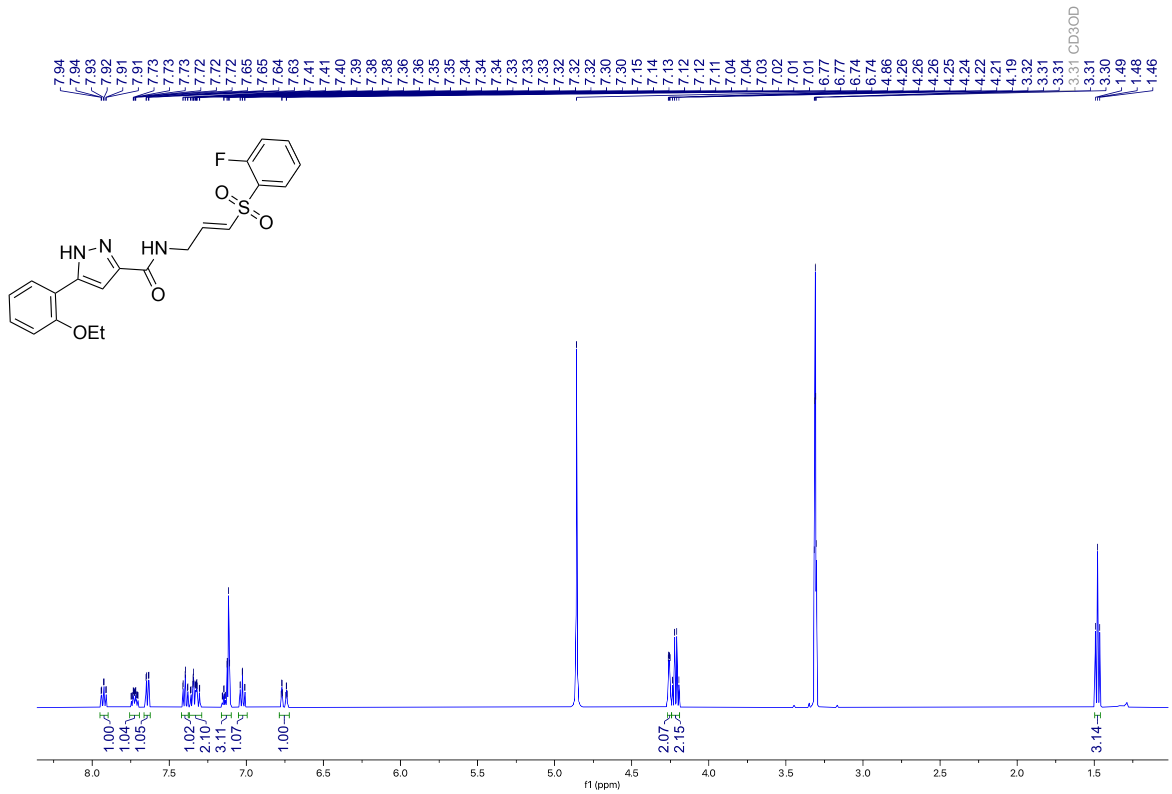

**Figure S139**: $^{13}$C NMR (100 MHz, DMSO-$d_6$) for **25f**

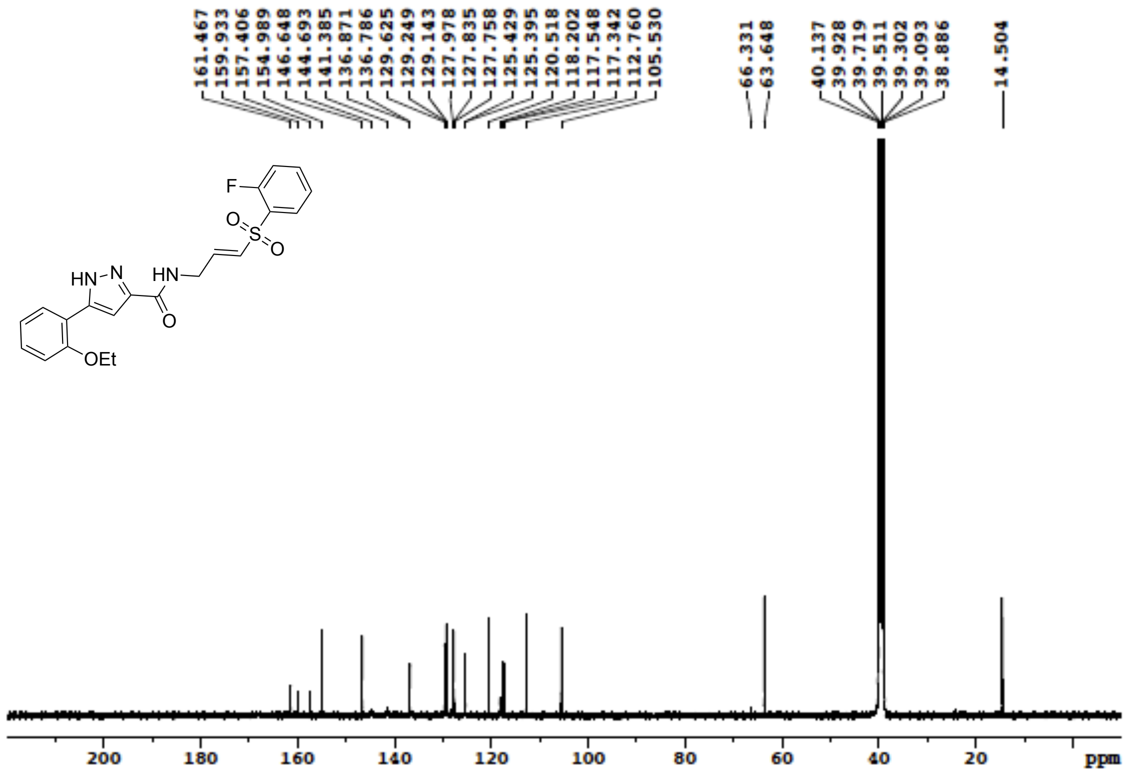

**Figure S140**: $^{1}$H NMR (400 MHz, DMSO-$d_6$) for **25g**

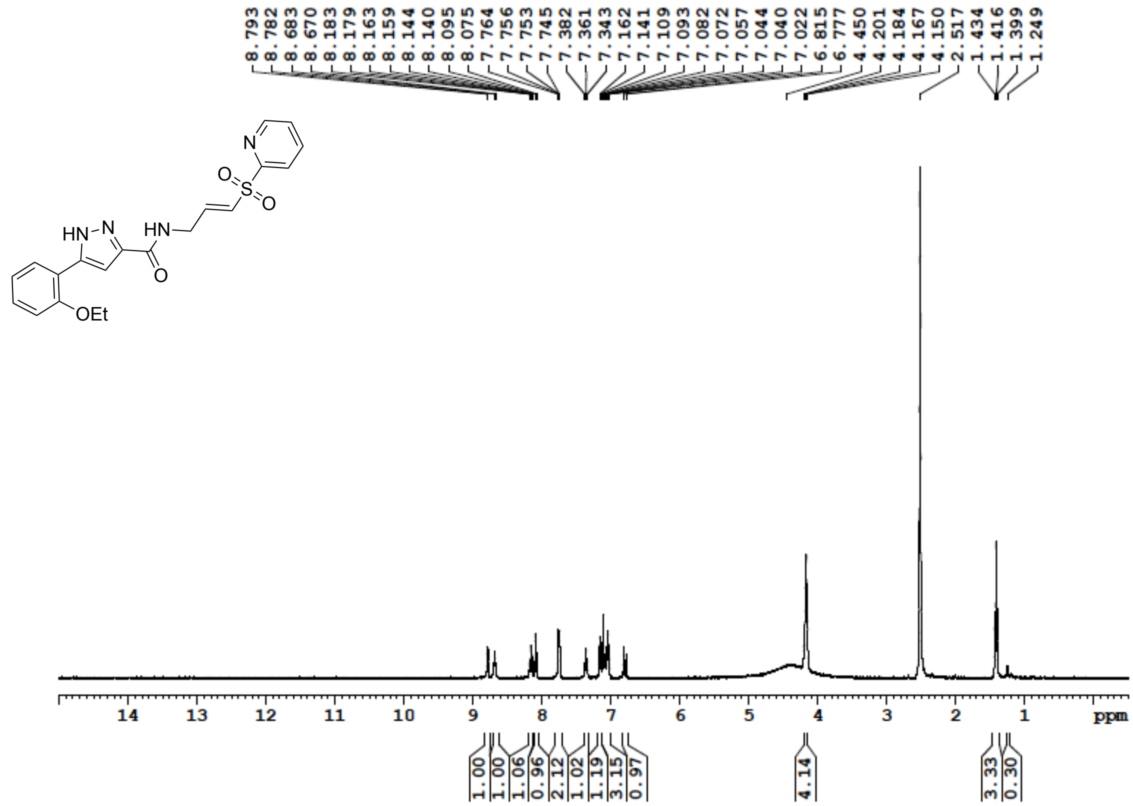


**Figure S141**: $^{13}$C NMR (100 MHz, DMSO-$d_6$) for **25g**

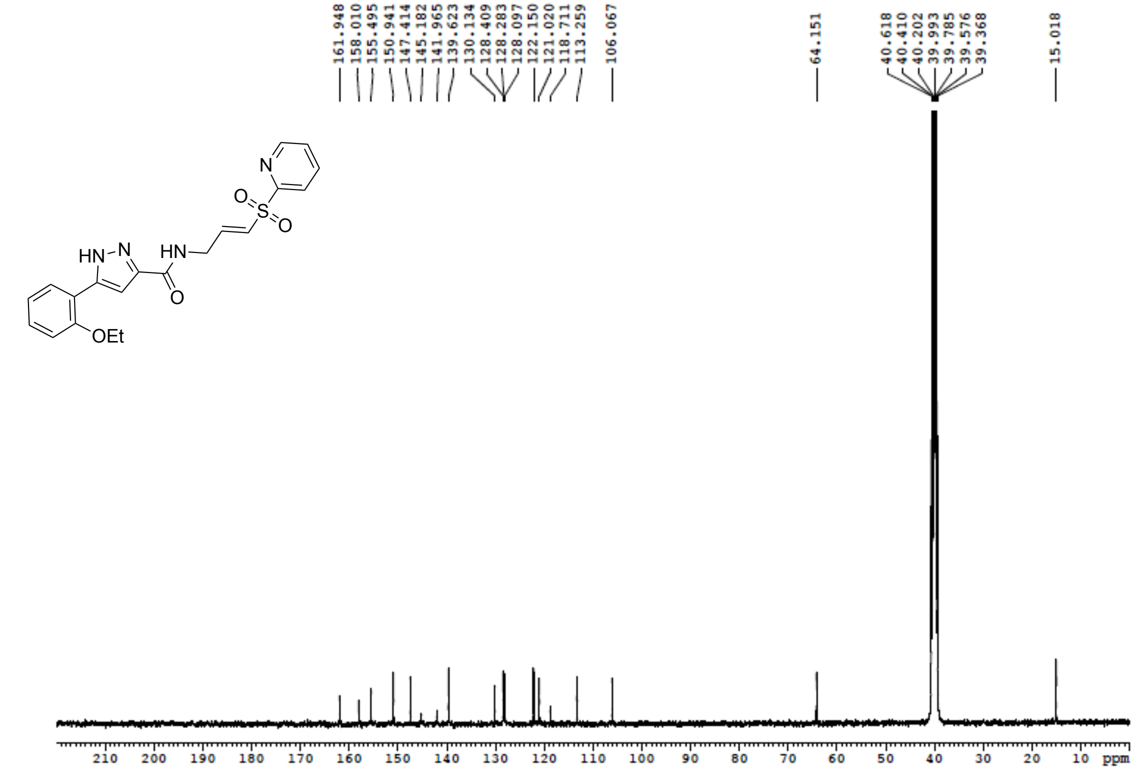

**Figure S142**: $^{1}$H NMR (400 MHz, DMSO-$d_6$) for **26**

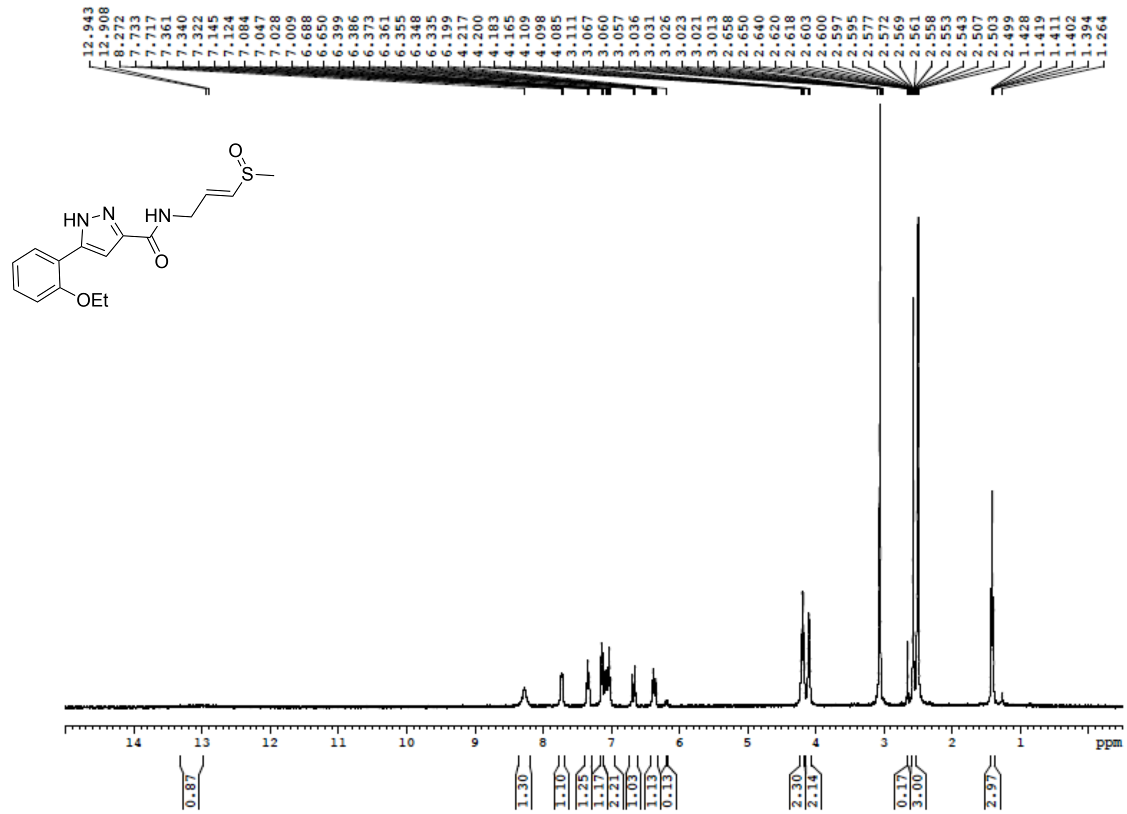

**Figure S143**: $^{13}$C NMR (100 MHz, DMSO-$d_6$) for **26**

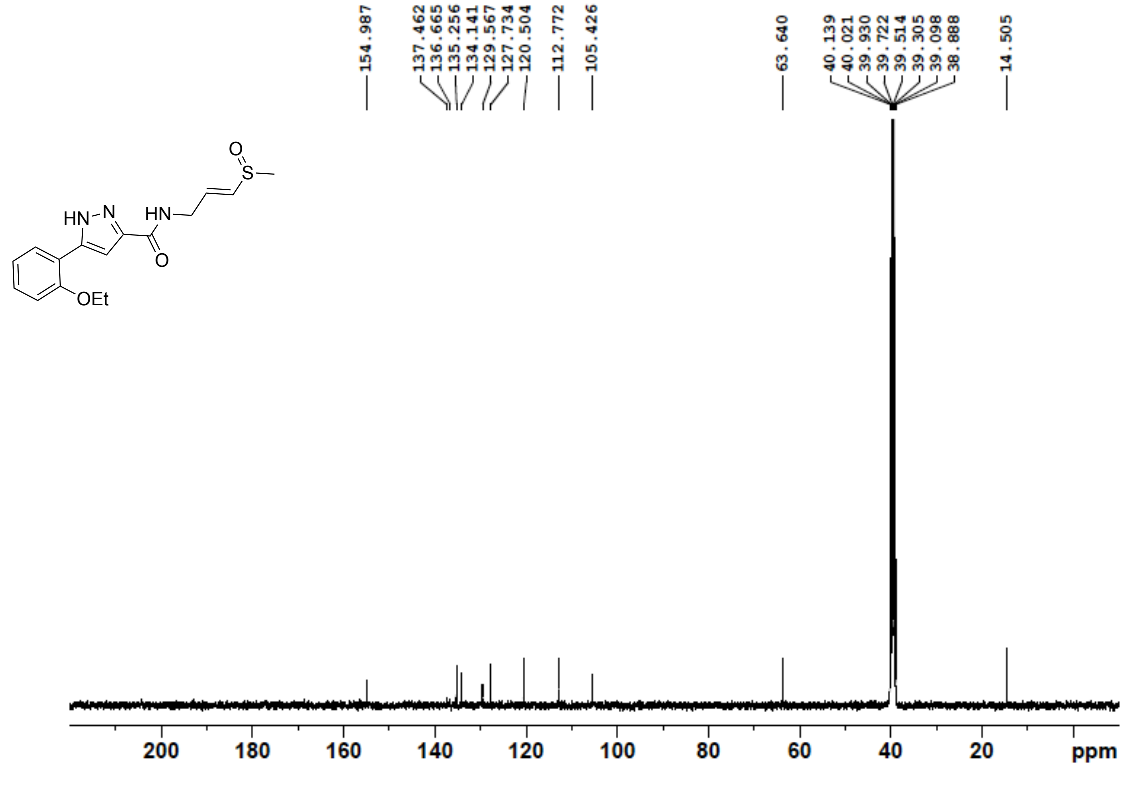

**Figure S144**: $^{1}$H NMR (400 MHz, DMSO-$d_6$) for **27**

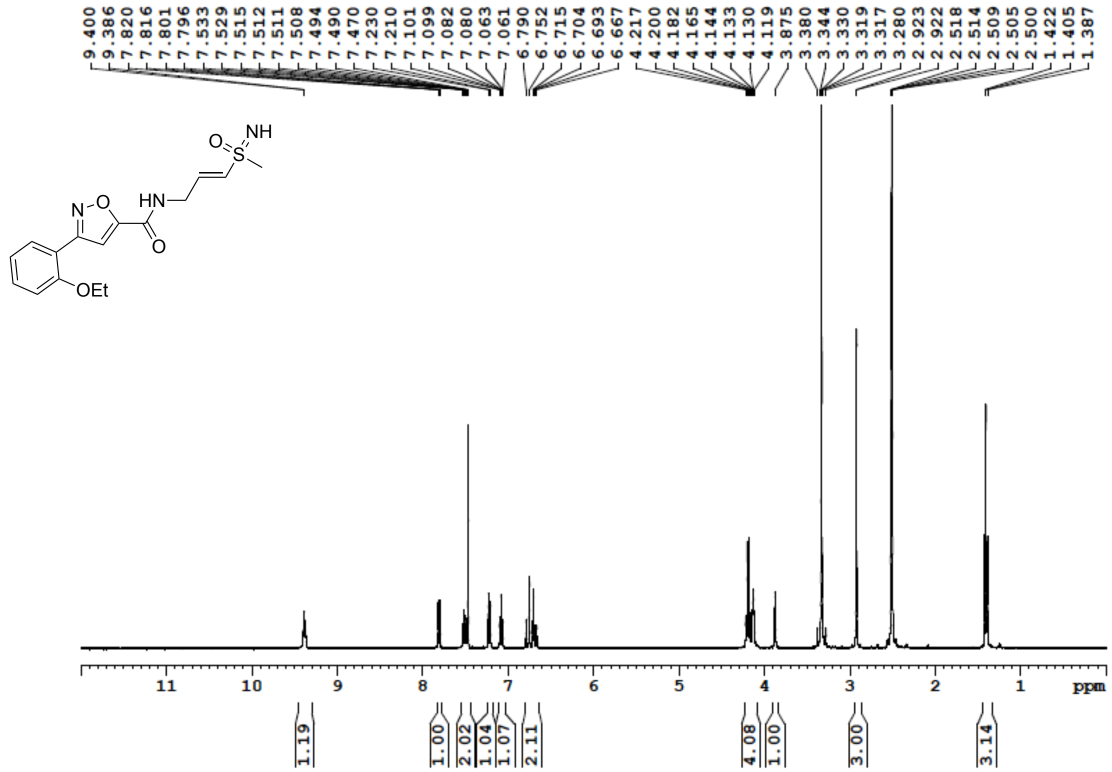

**Figure S145**: $^{13}$C NMR (100 MHz, DMSO-$d_6$) for **27**

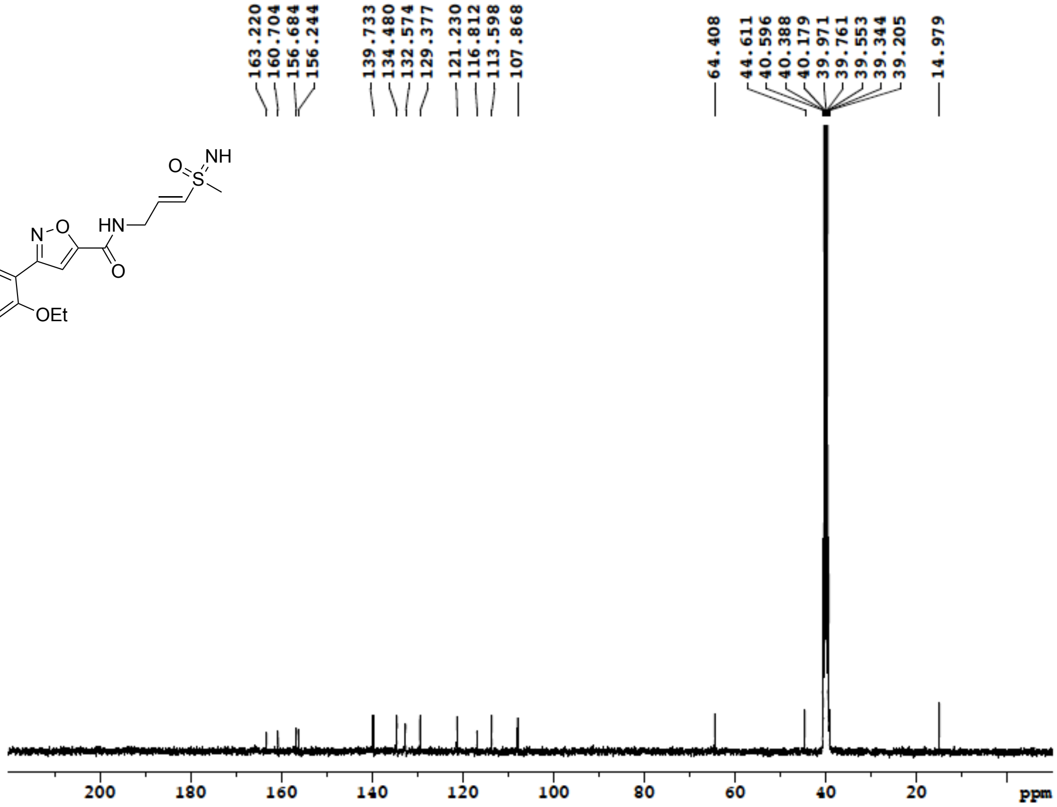

**Figure S146**: $^{1}$H NMR (400 MHz, DMSO-$d_6$) for **28**

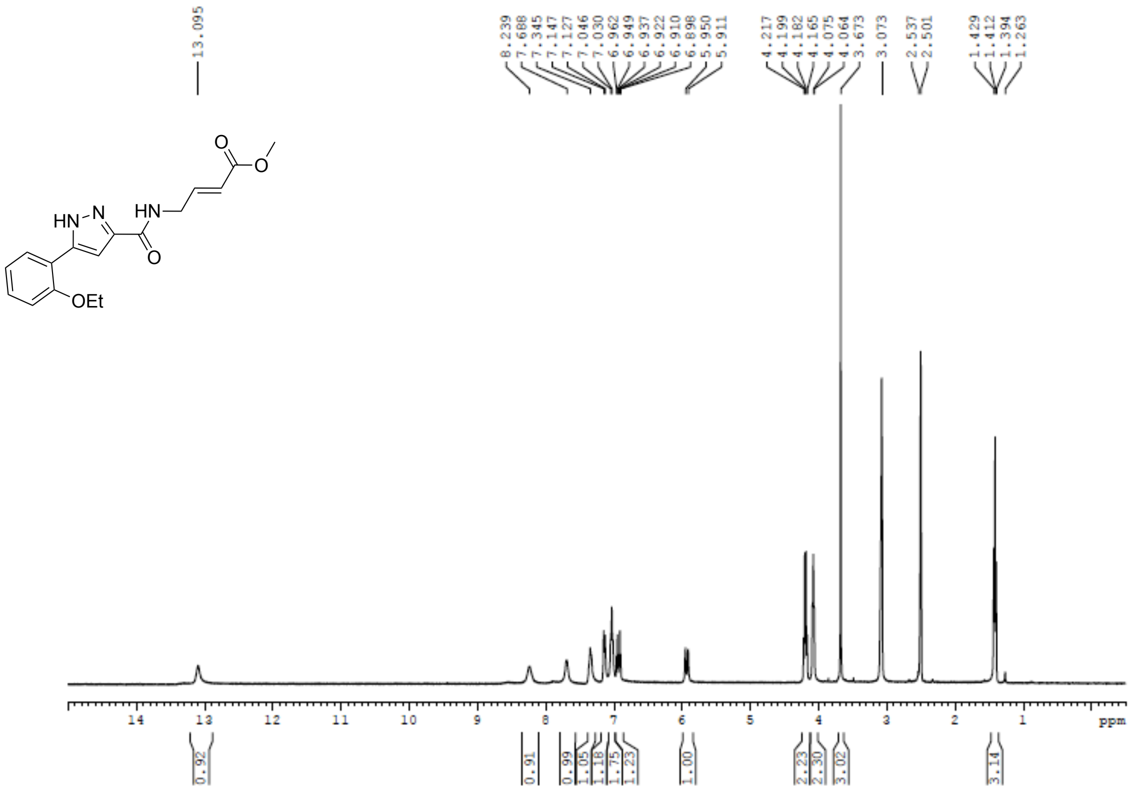


**Figure S147**: $^{13}$C NMR (100 MHz, DMSO-$d_6$) for **28**

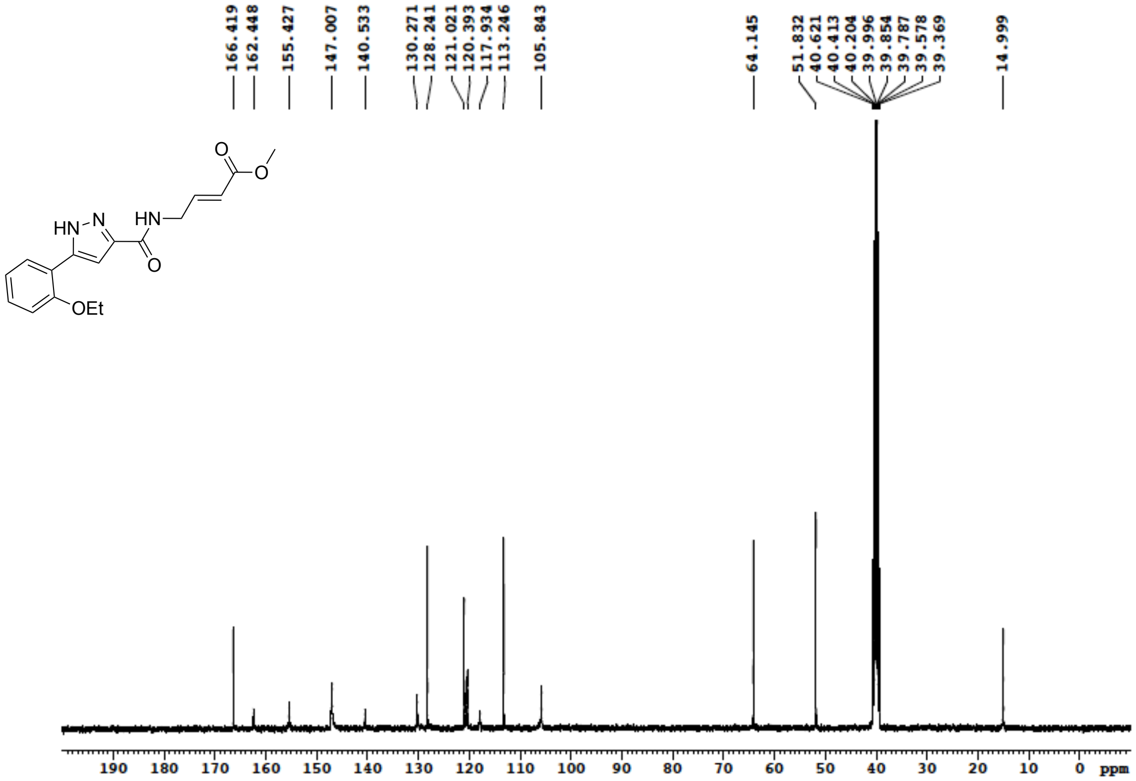

**Figure S148**: $^{1}$H NMR (400 MHz, DMSO-$d_6$) for **29**

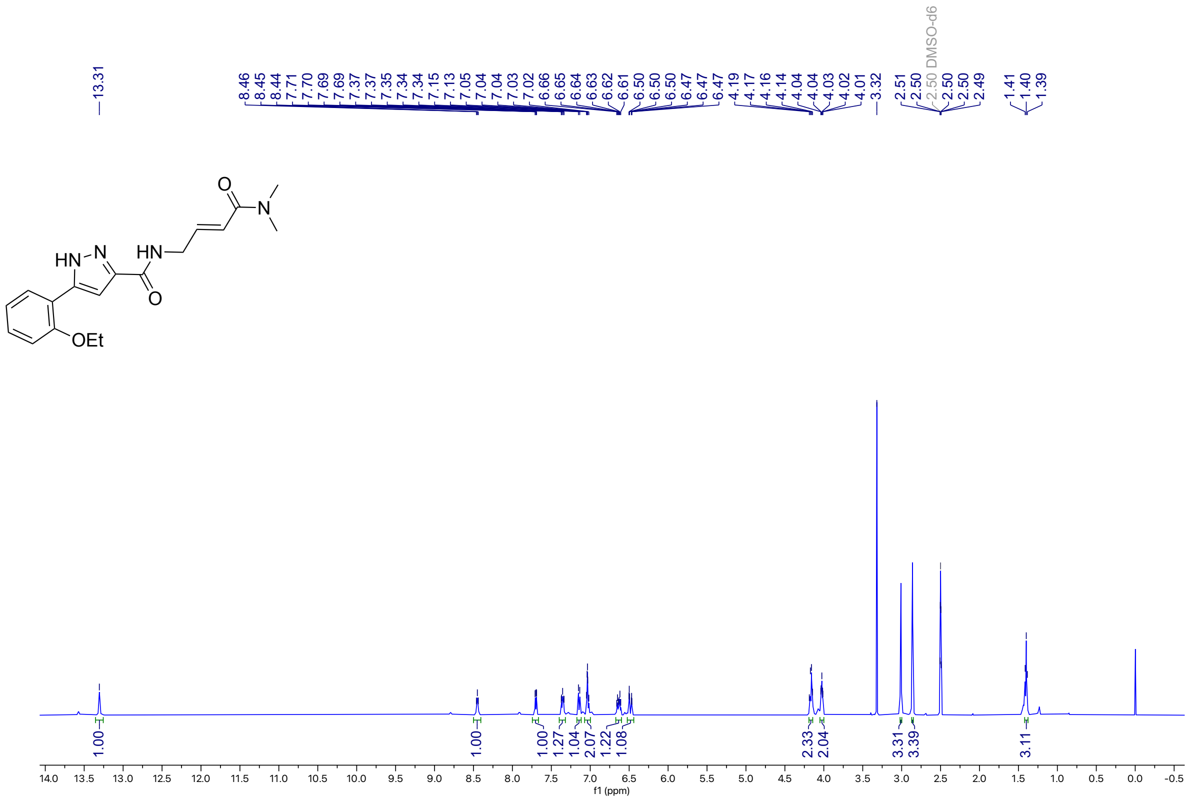

**Figure S149**: $^{13}$C NMR (100 MHz, DMSO-$d_6$) for **29**

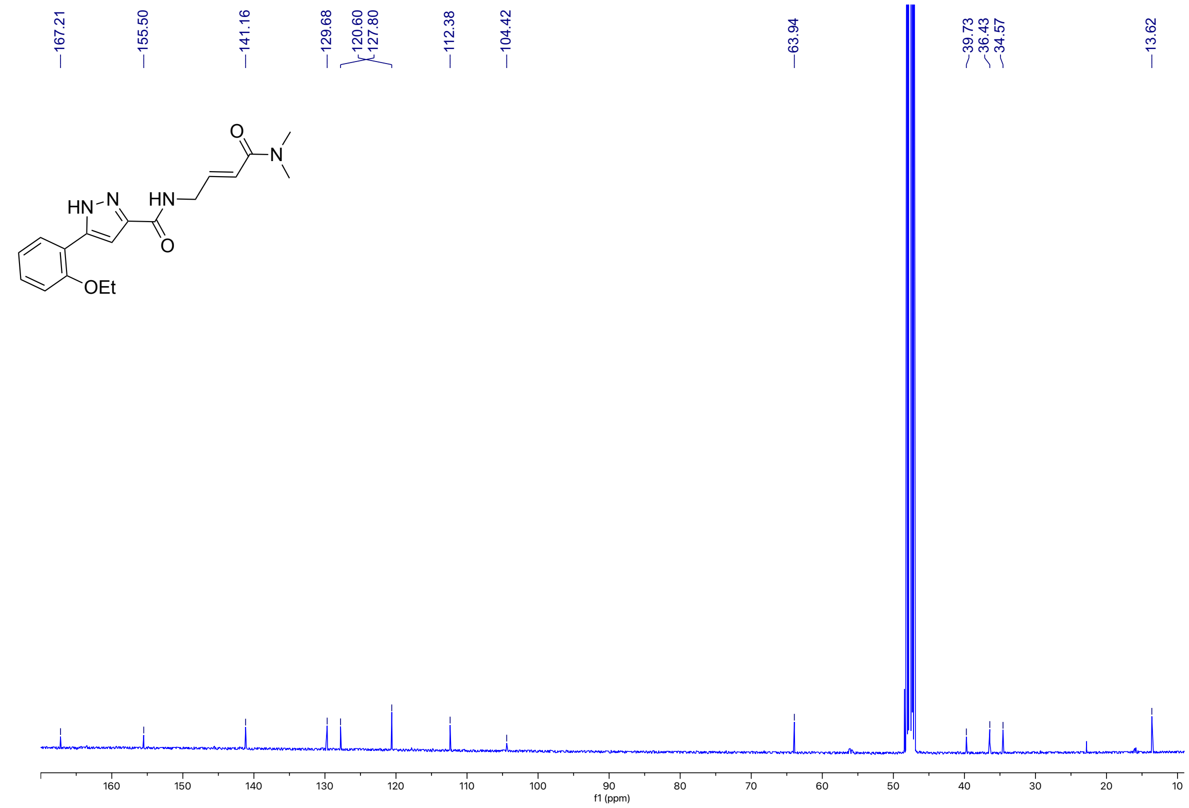

**Figure S150**: $^{1}$H NMR (400 MHz, DMSO-$d_6$) for **30**

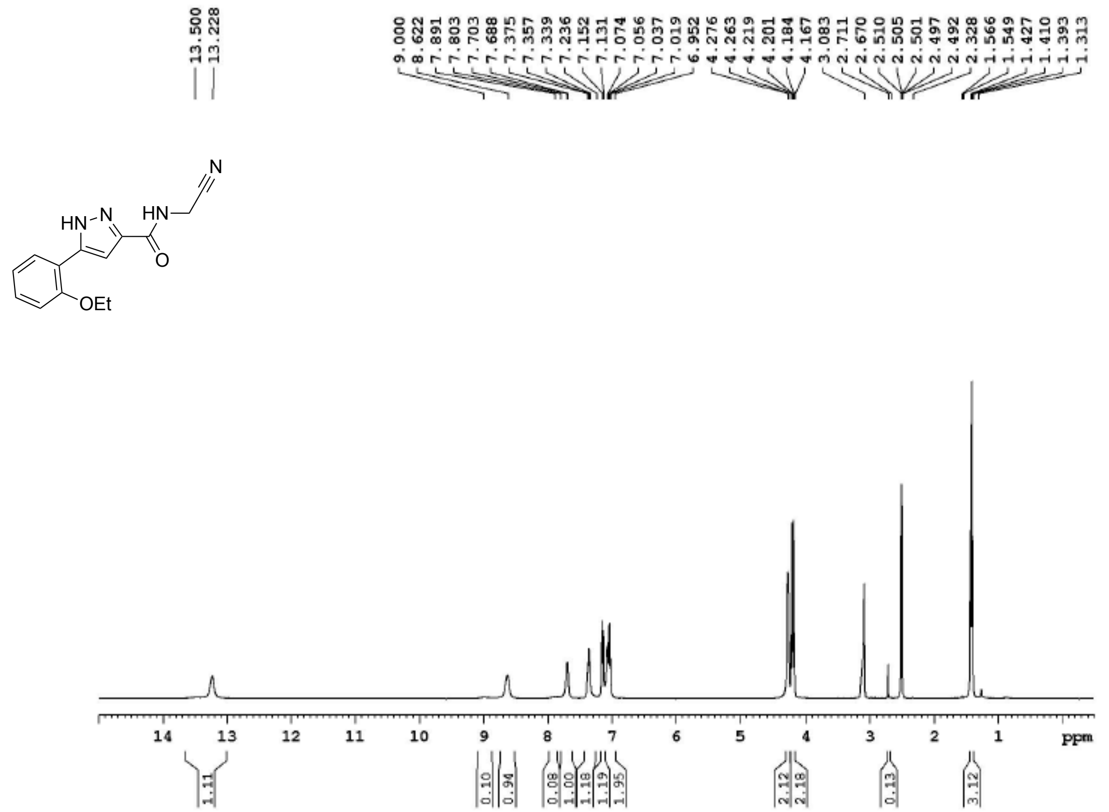


**Figure S151**: $^{13}$C NMR (100 MHz, DMSO-$d_6$) for **30**

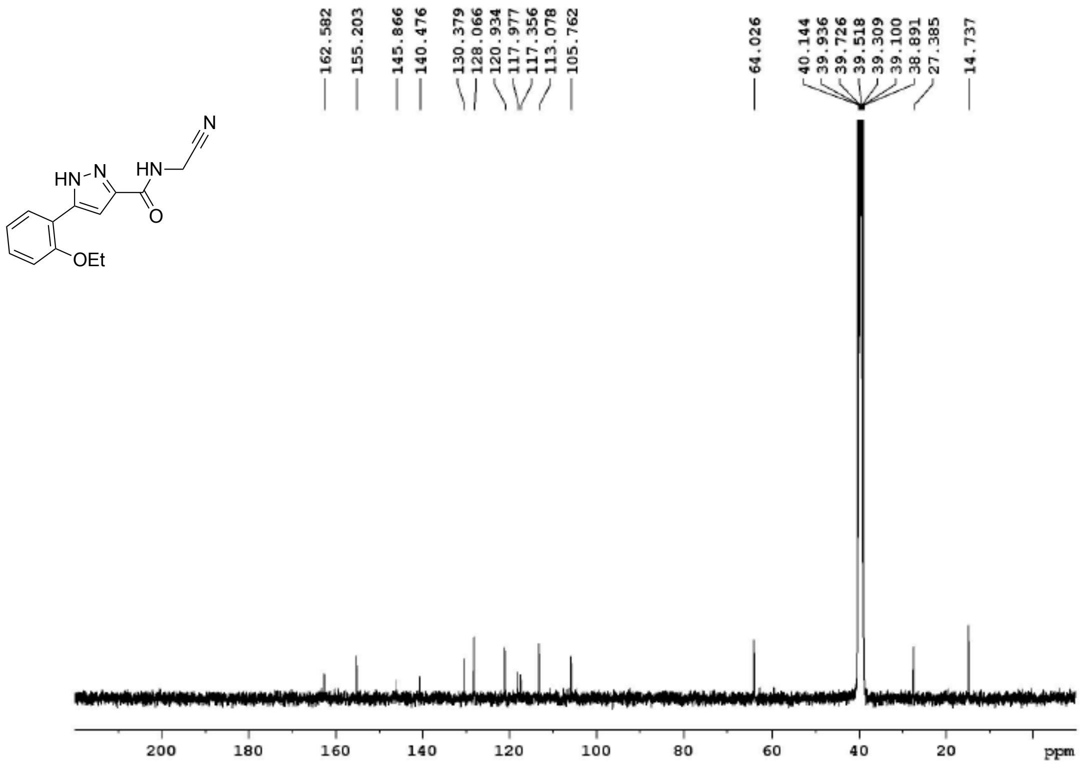

**Figure S152**: $^{1}$H NMR (400 MHz, DMSO-$d_6$) for **31**

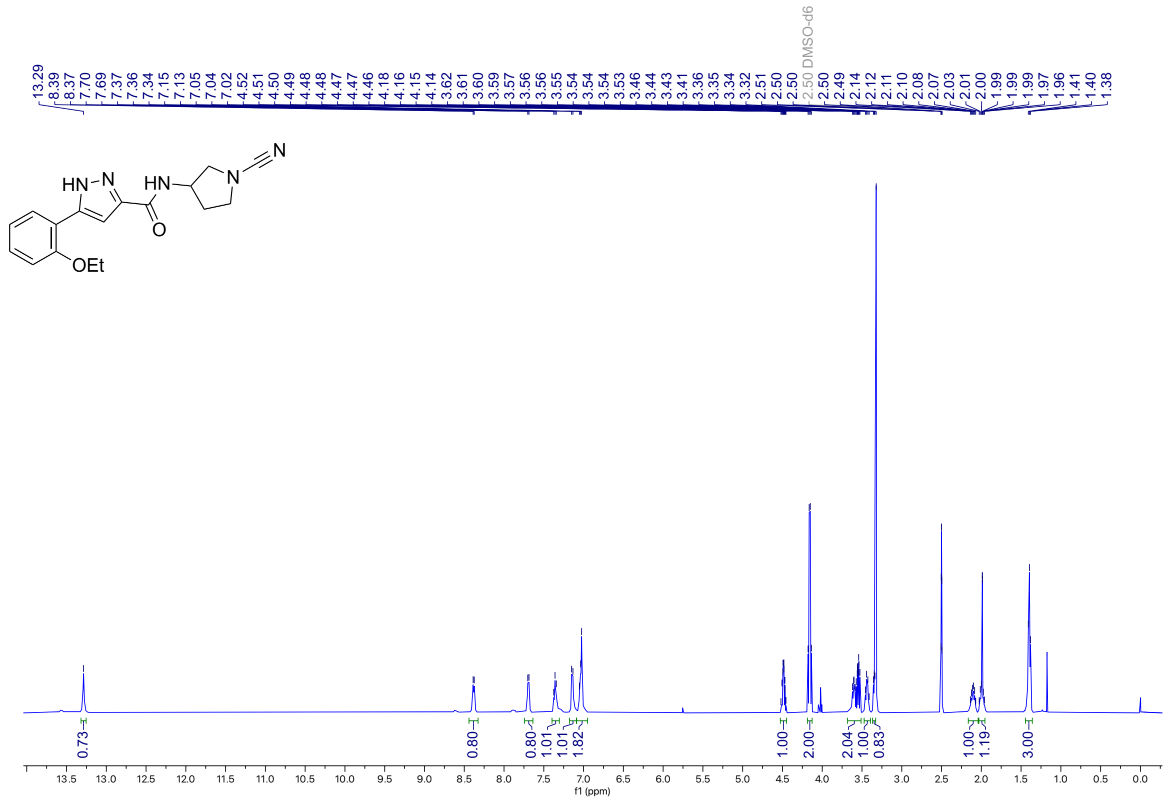

**Figure S153**: $^{13}$C NMR (100 MHz, DMSO-$d_6$) for **31**

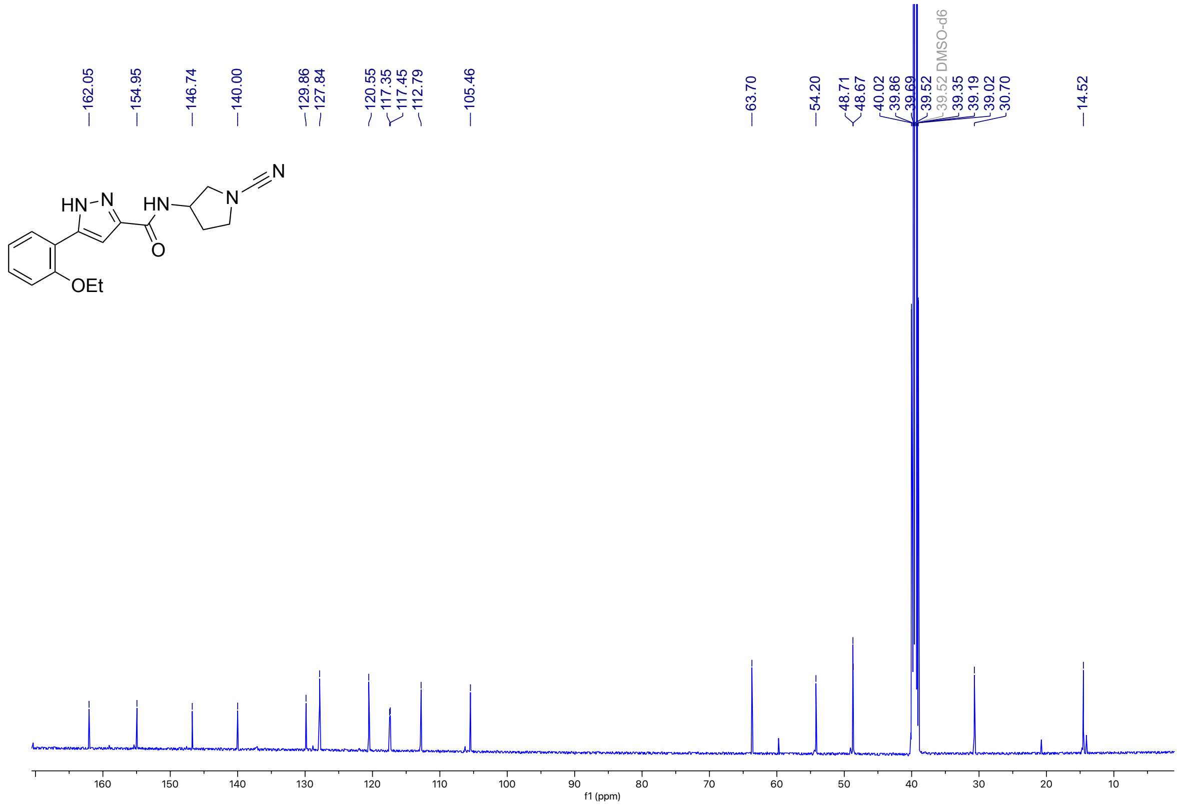

### HPLC analysis of pyrazole analogs

**Figure S154**: HPLC trace of **1a**

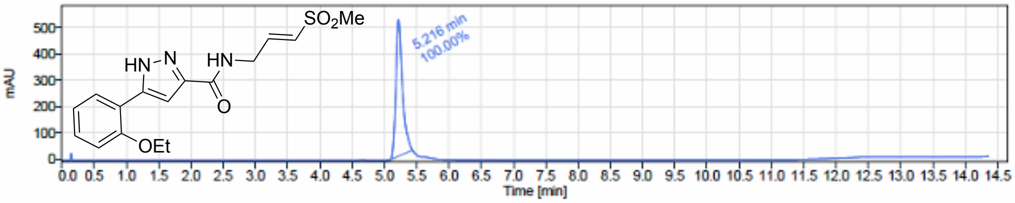


Signal Name DAD1A

| RT (min) | Signal description | m/z | Purity UV(‰) / MS(%) | Area | Area% |
|---|---|---|---|---|---|
| 5.216 | DAD1A,Sig=254.0,5.0 Ref=360.0,80.0 | | | 3652.8 | 100.00 |

**Figure S155**: HPLC trace of **1b**

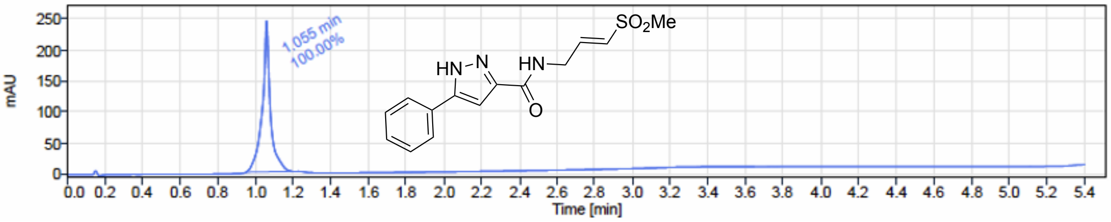


Signal Name DAD1A

| RT (min) | Signal description | m/z | Purity UV(‰) / MS(%) | Area | Area% |
|---|---|---|---|---|---|
| 1.055 | DAD1A,Sig=254.0,5.0 Ref=360.0,80.0 | | | 734.1 | 100.00 |

**Figure S156**: HPLC trace of **1c**

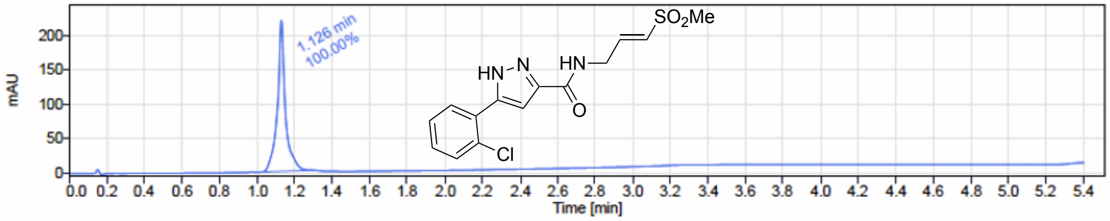


Signal Name DAD1A

| RT (min) | Signal description | m/z | Purity UV(‰) / MS(%) | Area | Area% |
|---|---|---|---|---|---|
| 1.126 | DAD1A,Sig=254.0,5.0 Ref=360.0,80.0 | | | 642.7 | 100.00 |

**Figure S157**: HPLC trace of **1d**

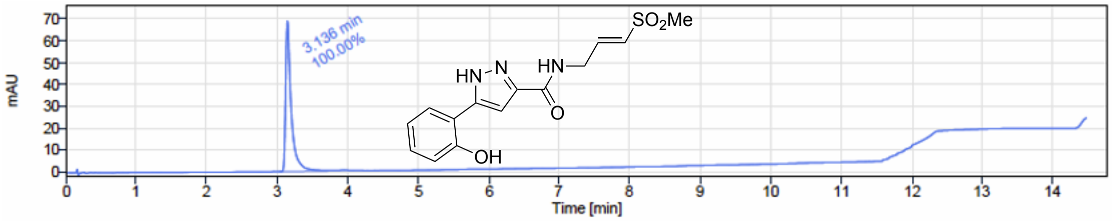


Signal Name DAD1A

| RT (min) | Signal description | m/z | Purity UV(‰) / MS(%) | Area | Area% |
|---|---|---|---|---|---|
| 3.136 | DAD1A,Sig=254.0,5.0 Ref=360.0,80.0 | | | 378.6 | 100.00 |

**Figure S158**: HPLC trace of **1e**

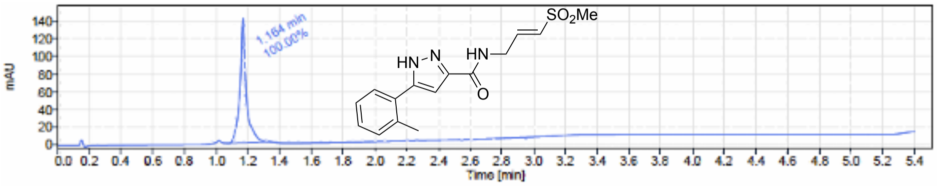


| Signal Name | DAD1A | | | | |
|---|---|---|---|---|---|
| RT (min) | Signal description | m/z | Purity UV(‰) / MS(%) | Area | Area% |
| 1.164 | DAD1A,Sig=254.0,5.0 Ref=360.0,80.0 | | | 395.4 | 100.00 |

**Figure S159**: HPLC trace of **1f**

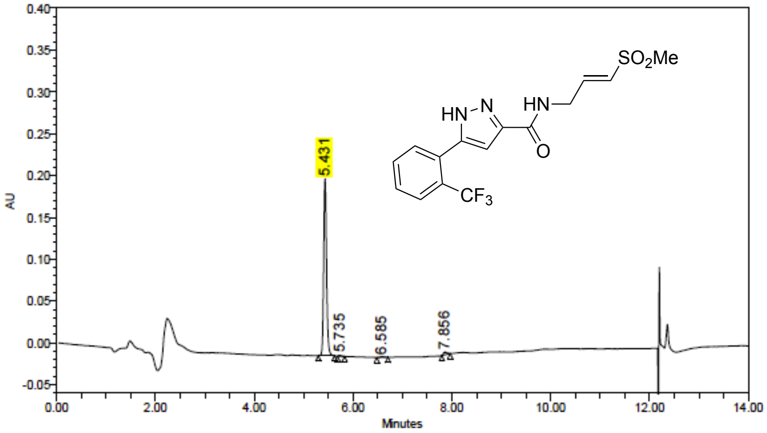


Peak Results

| | Name | RT | Height (µV) | Area (µV*sec) | % Area |
|---|---|---|---|---|---|
| 1 | | 5.431 | 211624 | 855150 | 97.00 |
| 2 | | 5.735 | 1387 | 5316 | 0.60 |
| 3 | | 6.585 | 841 | 5208 | 0.59 |
| 4 | | 7.856 | 3196 | 15943 | 1.81 |

**Figure S160**: HPLC trace of **1g**

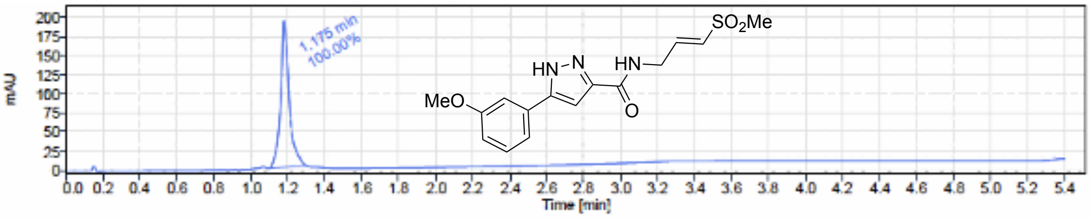


| Signal Name | DAD1A | | | | |
|---|---|---|---|---|---|
| RT (min) | Signal description | m/z | Purity UV(‰) / MS(%) | Area | Area% |
| 1.175 | DAD1A,Sig=254.0,5.0 Ref=360.0,80.0 | | | 614.4 | 100.00 |

**Figure S161**: HPLC trace of **1h**

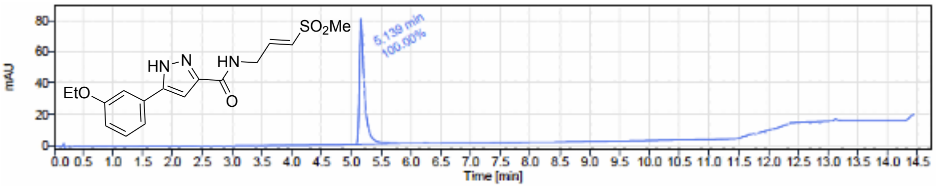


Signal Name DAD1A

| RT (min) | Signal description | m/z | Purity UV(‰) / MS(%) | Area | Area% |
|---|---|---|---|---|---|
| 5.139 | DAD1A,Sig=254.0,5.0 Ref=360.0,80.0 | | | 477.5 | 100.00 |

**Figure S162**: HPLC trace of **1i**

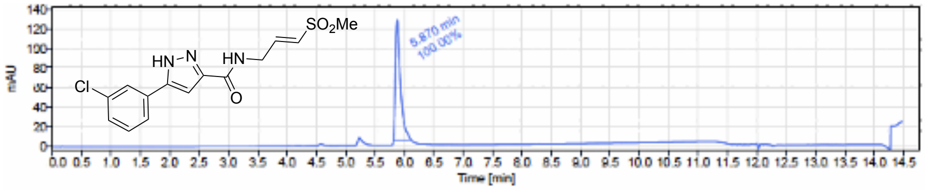


Signal Name DAD1A

| RT (min) | Signal description | m/z | Purity UV(‰) / MS(%) | Area | Area% |
|---|---|---|---|---|---|
| 5.870 | * DAD1A,Sig=254.0,5.0 Ref=360.0,80.0 [b] | | | 731.1 | 100.00 |

**Figure S163**: HPLC trace of **1j**

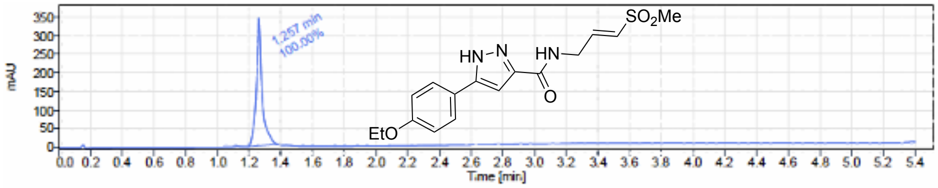


Signal Name DAD1A

| RT (min) | Signal description | m/z | Purity UV(‰) / MS(%) | Area | Area% |
|---|---|---|---|---|---|
| 1.257 | DAD1A,Sig=254.0,5.0 Ref=360.0,80.0 | | | 878.5 | 100.00 |

**Figure S164**: HPLC trace of **1k**

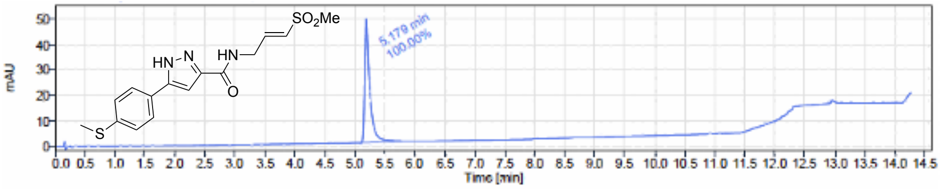


Signal Name DAD1A

| RT (min) | Signal description | m/z | Purity UV(‰) / MS(%) | Area | Area% |
|---|---|---|---|---|---|
| 5.179 | DAD1A,Sig=254.0,5.0 Ref=360.0,80.0 | | | 290.4 | 100.00 |

**Figure S165**: HPLC trace of **1l**

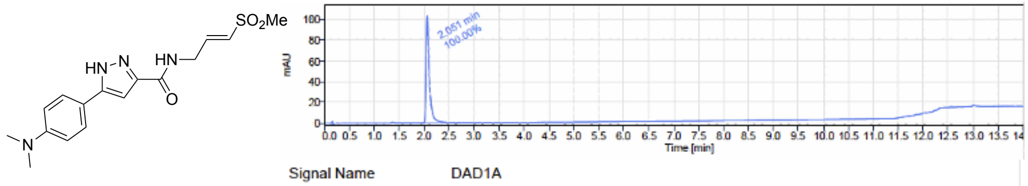


Signal Name DAD1A

| RT (min) | Signal description | m/z | Purity UV(‰) / MS(%) | Area | Are |
|---|---|---|---|---|---|
| 2.051 | DAD1A,Sig=254.0,5.0 Ref=360.0,80.0 | | | 466.7 | 100 |

**Figure S166**: HPLC trace of **1m**

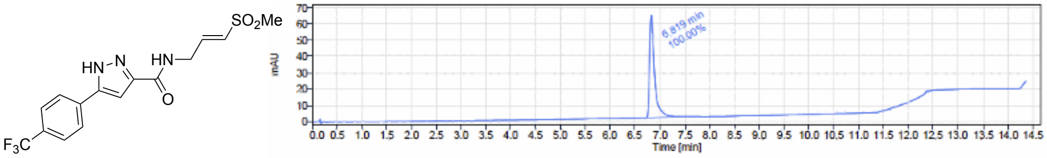


Signal Name DAD1A

| RT (min) | Signal description | m/z | Purity UV(‰) / MS(%) | Area | Area% |
|---|---|---|---|---|---|
| 6.819 | DAD1A,Sig=254.0,5.0 Ref=360.0,80.0 | | | 439.1 | 100.00 |

**Figure S167**: HPLC trace of **1n**

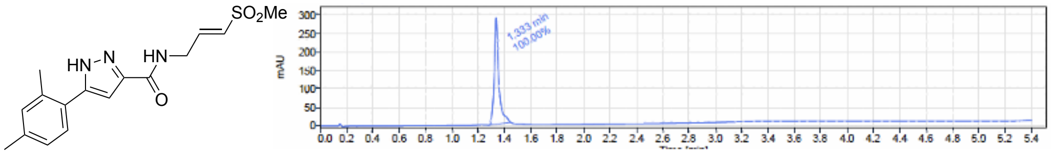


Signal Name DAD1A

| RT (min) | Signal description | m/z | Purity UV(‰) / MS(%) | Area | Area% |
|---|---|---|---|---|---|
| 1.333 | DAD1A,Sig=254.0,5.0 Ref=360.0,80.0 | | | 684.7 | 100.00 |

**Figure S168**: HPLC trace of **1o**

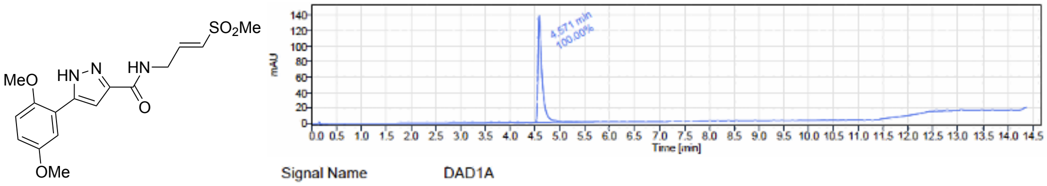


Signal Name DAD1A

| RT (min) | Signal description | m/z | Purity UV(‰) / MS(%) | Area | Area% |
|---|---|---|---|---|---|
| 4.571 | DAD1A,Sig=254.0,5.0 Ref=360.0,80.0 | | | 798.8 | 100.00 |

**Figure S169**: HPLC trace of **1p**

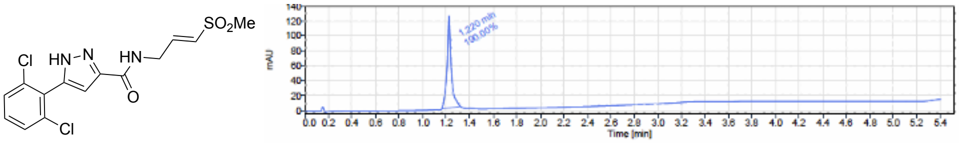


Signal Name DAD1A

| RT (min) | Signal description | m/z | Purity UV(‰) / MS(%) | Area | Area% |
|---|---|---|---|---|---|
| 1.220 | DAD1A,Sig=254.0,5.0 Ref=360.0,80.0 | | | 307.5 | 100.00 |

**Figure S170**: HPLC trace of **1q**

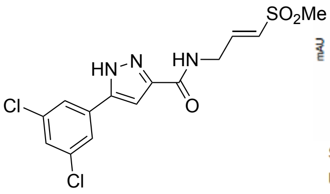


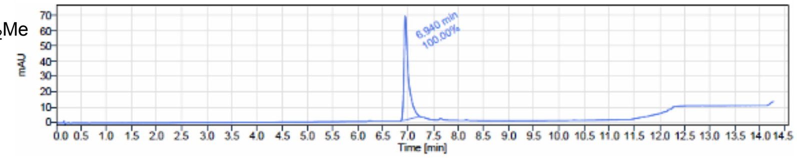


Signal Name DAD1A

| RT (min) | Signal description | m/z | Purity UV(‰) / MS(%) | Area | Area% |
|---|---|---|---|---|---|
| 6.940 | DAD1A,Sig=254.0,5.0 Ref=360.0,80.0 | | | 420.4 | 100.00 |

**Figure S171**: HPLC trace of **2**

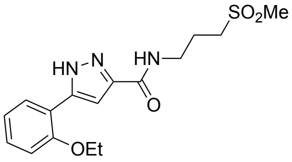


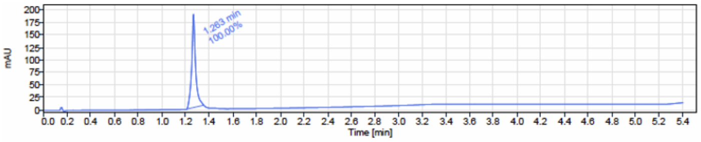


Signal Name DAD1A

| RT (min) | Signal description | m/z | Purity UV(‰) / MS(%) | Area | Area% |
|---|---|---|---|---|---|
| 1.263 | DAD1A,Sig=254.0,5.0 Ref=360.0,80.0 | | | 431.7 | 100.00 |

**Figure S172**: HPLC trace of **3**

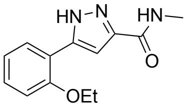


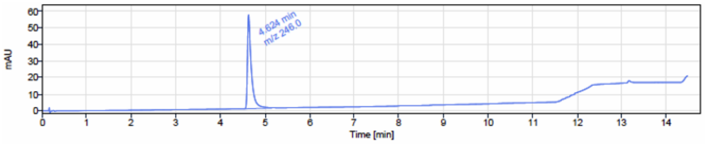


Signal Name DAD1A

| RT (min) | Signal description | m/z | Purity UV(‰) / MS(%) | Area | Area% |
|---|---|---|---|---|---|
| 4.624 | DAD1A,Sig=254.0,5.0 Ref=360.0,80.0 | 246.0 | | 339.8 | 100.00 |

**Figure S173**: HPLC trace of **4a**

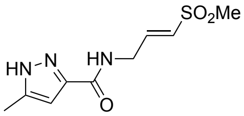


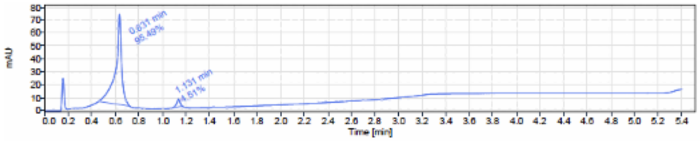


Signal Name DAD1A

| RT (min) | Signal description | m/z | Purity UV(‰) / MS(%) | Area | Area% |
|---|---|---|---|---|---|
| 0.631 | DAD1A,Sig=254.0,5.0 Ref=360.0,80.0 | | | 252.5 | 95.49 |
| 1.131 | DAD1A,Sig=254.0,5.0 Ref=360.0,80.0 | | | 11.9 | 4.51 |

**Figure S174**: HPLC trace of **4b**

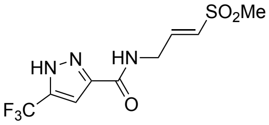


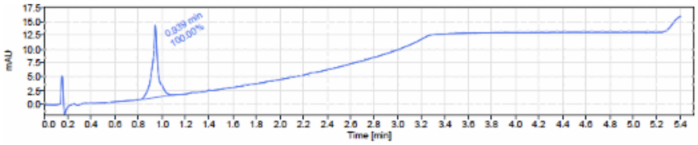


Signal Name DAD1A

| RT (min) | Signal description | m/z | Purity UV(‰) / MS(%) | Area | Area% |
|---|---|---|---|---|---|
| 0.939 | DAD1A,Sig=254.0,5.0 Ref=360.0,80.0 | | | 45.0 | 100.00 |

**Figure S175**: HPLC trace of **4c**

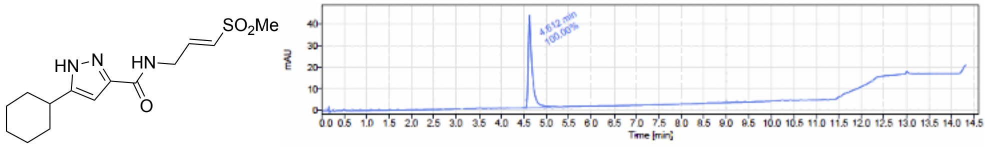


Signal Name DAD1A

| RT (min) | Signal description | m/z | Purity UV(‰) / MS(%) | Area | Area% |
|---|---|---|---|---|---|
| 4.612 | DAD1A,Sig=254.0,5.0 Ref=360.0,80.0 | | | 255.1 | 100.00 |

**Figure S176**: HPLC trace of **4d**

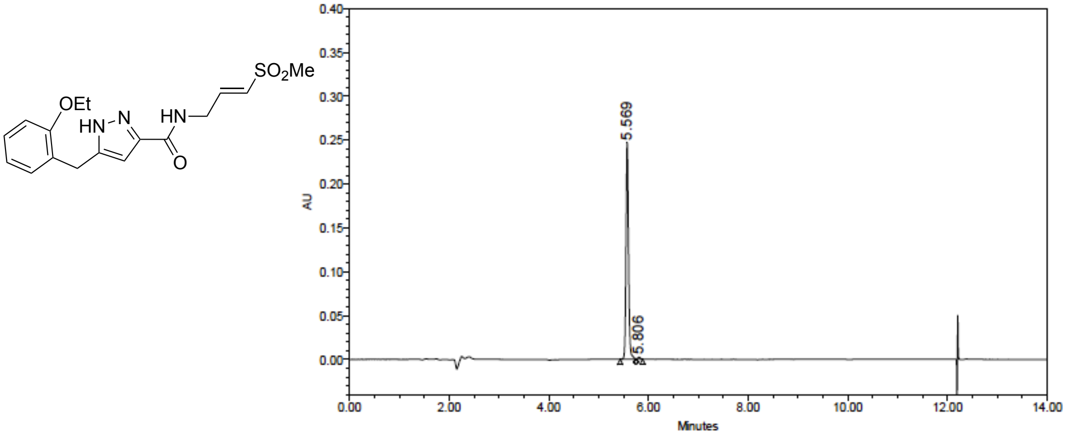


Peak Results

| | Name | RT | Height (μV) | Area (μV*sec) | % Area |
|---|---|---|---|---|---|
| 1 | | 5.569 | 247581 | 863325 | 98.95 |
| 2 | | 5.806 | 2444 | 9195 | 1.05 |

**Figure S177**: HPLC trace of **4e**

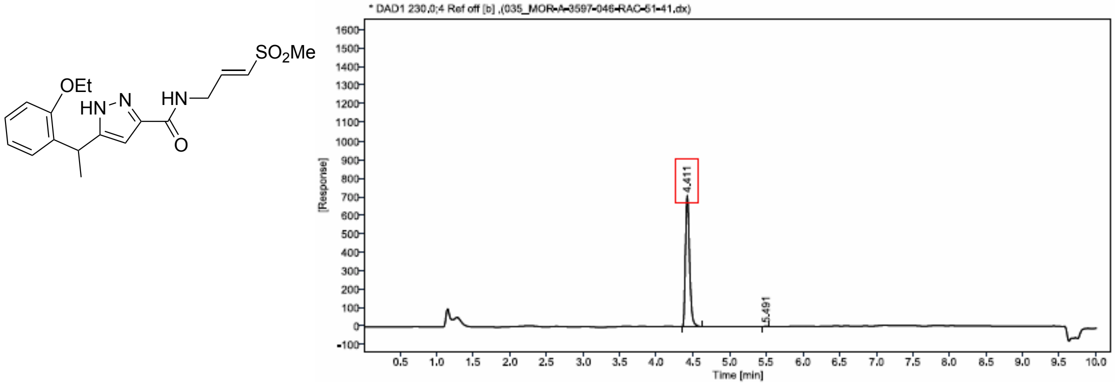


| Sr.No. | RT | Area | Height | % Area |
|---|---|---|---|---|
| 1 | 4,41 | 2712 | 705 | 99,90 |
| 2 | 5,49 | 3 | 1 | 0,10 |

**Figure S178**: HPLC trace of **4f**

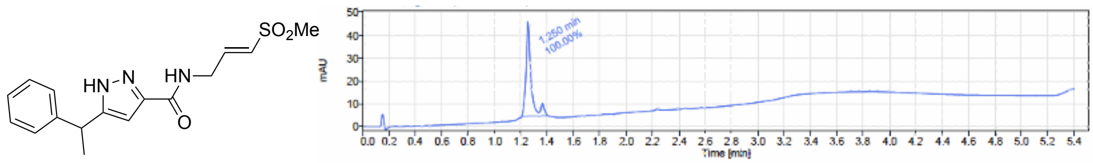


| Signal Name | DAD1A | | | | |
|---|---|---|---|---|---|
| RT (min) | Signal description | m/z | Purity UV(‰) / MS(%) | Area | Area% |
| 1.250 | DAD1A,Sig=254.0,5.0 Ref=360.0,80.0 | | | 112.1 | 100.00 |

**Figure S179**: HPLC trace of **4g**

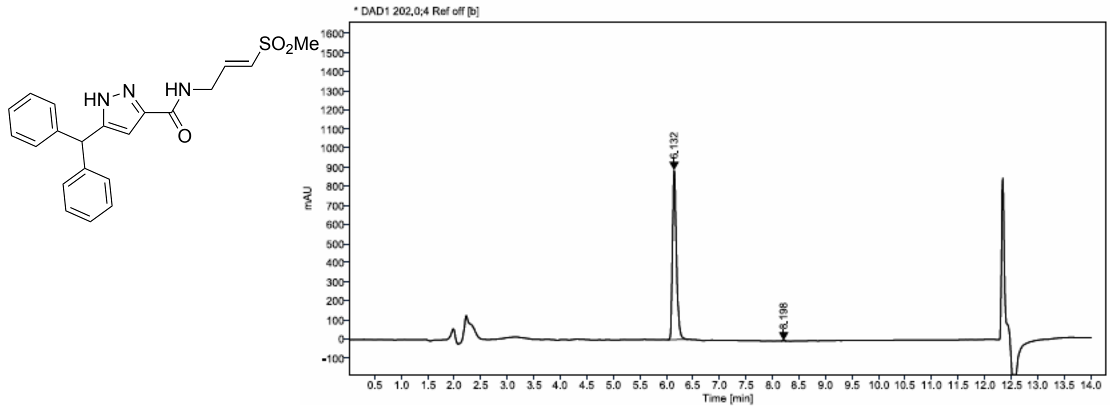


| Sr.No. | RT | Area | Height | % Area |
|---|---|---|---|---|
| 1 | 6.13 | 4622 | 888 | 99.86 |
| 2 | 8.20 | 6 | 2 | 0.14 |

**Figure S180**: HPLC trace of **4h**

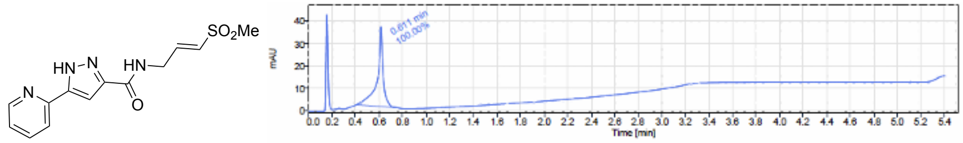


| Signal Name | DAD1A | | | | |
|---|---|---|---|---|---|
| RT (min) | Signal description | m/z | Purity UV(‰) / MS(%) | Area | Area% |
| 0.611 | DAD1A,Sig=254.0,5.0 Ref=360.0,80.0 | | | 125.2 | 100.00 |

**Figure S181**: HPLC trace of **4i**

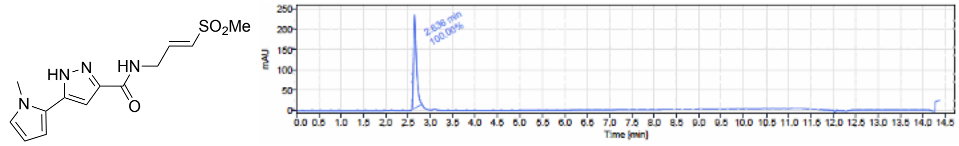


| Signal Name | DAD1A | | | | |
|---|---|---|---|---|---|
| RT (min) | Signal description | m/z | Purity UV(‰) / MS(%) | Area | Area% |
| 2.636 | * DAD1A,Sig=254.0,5.0 Ref=360.0,80.0 [b] | | | 1065.7 | 100.00 |

**Figure S182**: HPLC trace of **5**

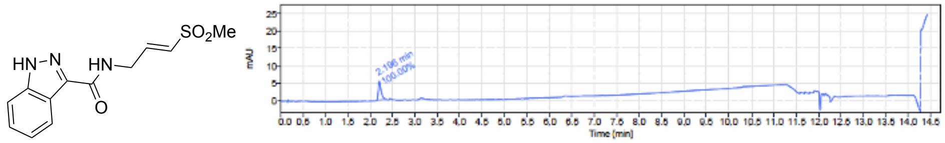


| Signal Name | DAD1A | | | | |
|---|---|---|---|---|---|
| RT (min) | Signal description | m/z | Purity UV(‰) / MS(%) | Area | Area% |
| 2.196 | * DAD1A,Sig=254.0,5.0 Ref=360.0,80.0 [b] | | | 23.3 | 100.00 |

**Figure S183**: HPLC trace of **6**

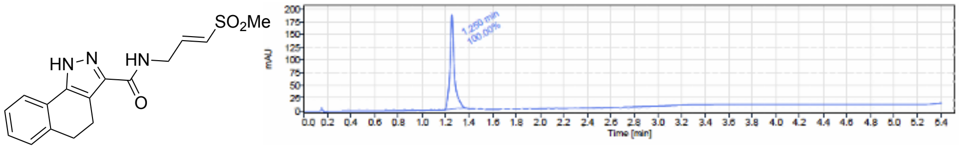


| Signal Name | DAD1A | | | | |
|---|---|---|---|---|---|
| RT (min) | Signal description | m/z | Purity UV(‰) / MS(%) | Area | Area% |
| 1.250 | DAD1A,Sig=254.0,5.0 Ref=360.0,80.0 | | | 474.0 | 100.00 |

**Figure S184**: HPLC trace of **7a**

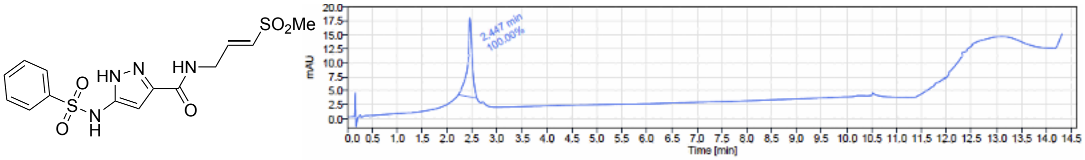


| Signal Name | DAD1A | | | | |
|---|---|---|---|---|---|
| RT (min) | Signal description | m/z | Purity UV(‰) / MS(%) | Area | Area% |
| 2.447 | DAD1A,Sig=254.0,5.0 Ref=360.0,80.0 | | | 94.0 | 100.00 |

**Figure S185**: HPLC trace of **7b**

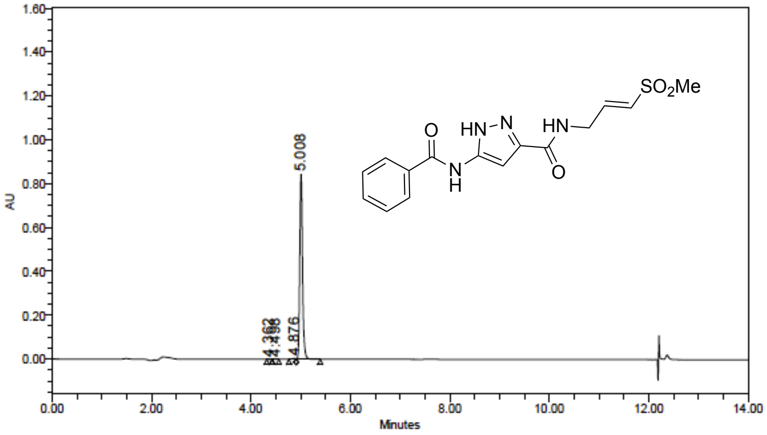


Peak Results

| | Name | RT | Height (µV) | Area (µV*sec) | % Area |
|---|---|---|---|---|---|
| 1 | | 4.362 | 787 | 2309 | 0.07 |
| 2 | | 4.498 | 732 | 2633 | 0.08 |
| 3 | | 4.876 | 4643 | 21707 | 0.66 |
| 4 | | 5.008 | 843836 | 3277199 | 99.19 |

**Figure S186**: HPLC trace of **7c**

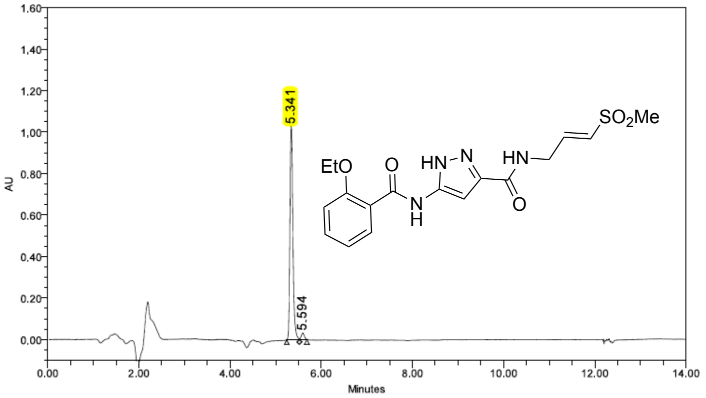

Peak Results

| | Name | RT | Height (µV) | Area (µV*sec) | % Area |
|---|---|---|---|---|---|
| 1 | | 5,341 | 1028710 | 4618285 | 97,00 |
| 2 | | 5,594 | 30827 | 142877 | 3,00 |

**Figure S187**: HPLC trace of **7d**

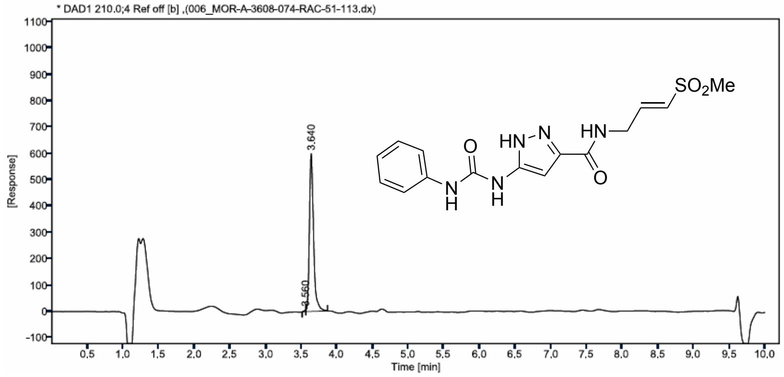

| Sr.No. | RT | Area | Height | % Area |
|---|---|---|---|---|
| 1 | 3,56 | 9 | 7 | 0,36 |
| 2 | 3,64 | 2435 | 598 | 99,64 |

**Figure S188**: HPLC trace of **7e**

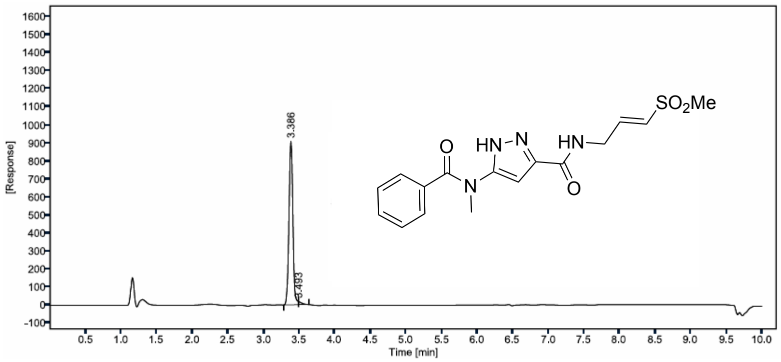

| Sr.No. | RT | Area | Height | % Area |
|---|---|---|---|---|
| 1 | 3,39 | 3486 | 906 | 98,47 |
| 2 | 3,49 | 54 | 18 | 1,53 |

**Figure S189**: HPLC trace of **7f**

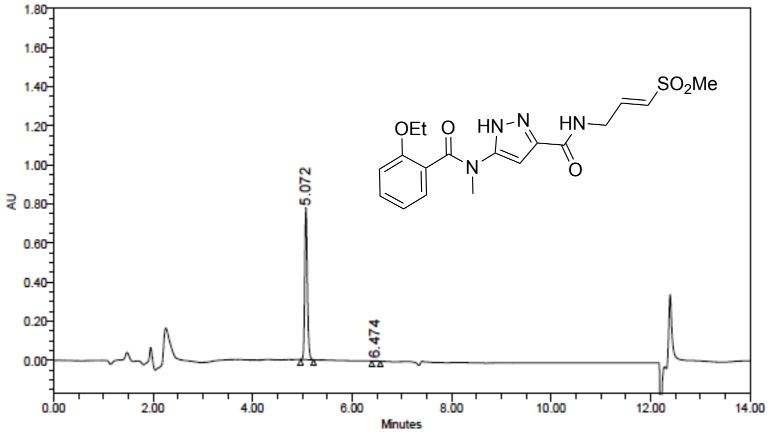


**Peak Results**

| | Name | RT | Height (μV) | Area (μV*sec) | % Area |
|---|---|---|---|---|---|
| 1 | | 5.072 | 775596 | 2834909 | 99.63 |
| 2 | | 6.474 | 2035 | 10557 | 0.37 |

**Figure S190**: HPLC trace of **8a**

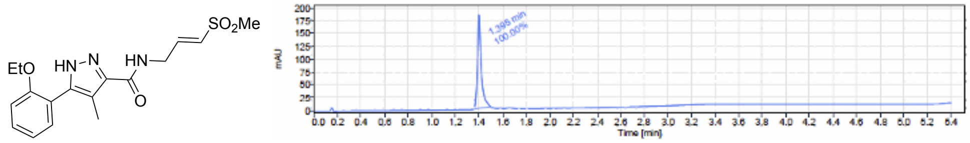


Signal Name DAD1A

| RT (min) | Signal description | m/z | Purity UV(‰) / MS(%) | Area | Area% |
|---|---|---|---|---|---|
| 1.395 | DAD1A,Sig=254.0,5.0 Ref=360.0,80.0 | | | 408.6 | 100.00 |

**Figure S191**: HPLC trace of **8b**

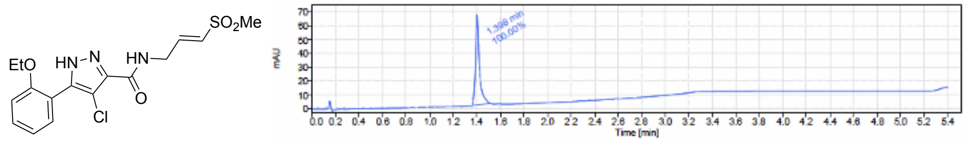


Signal Name DAD1A

| RT (min) | Signal description | m/z | Purity UV(‰) / MS(%) | Area | Area% |
|---|---|---|---|---|---|
| 1.398 | DAD1A,Sig=254.0,5.0 Ref=360.0,80.0 | | | 150.6 | 100.00 |

**Figure S192**: HPLC trace of **8c**

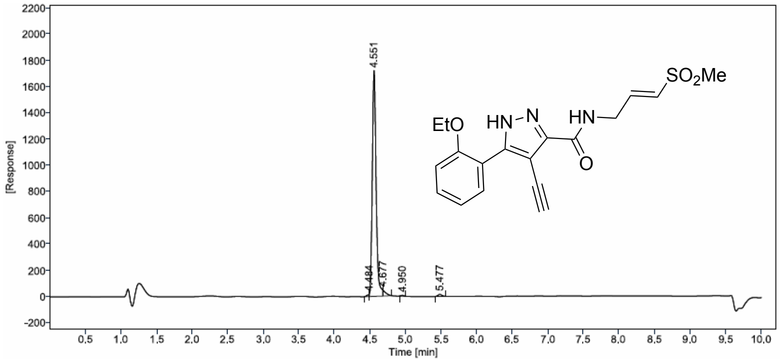


| Sr.No. | RT | Area | Height | % Area |
|---|---|---|---|---|
| 1 | 4.48 | 32 | 16 | 0.48 |
| 2 | 4,55 | 6471 | 1716 | 96,48 |
| 3 | 4.68 | 128 | 45 | 1.91 |
| 4 | 4.95 | 18 | 6 | 0.26 |
| 5 | 5,48 | 58 | 16 | 0,87 |

**Figure S193**: HPLC trace of **8d**

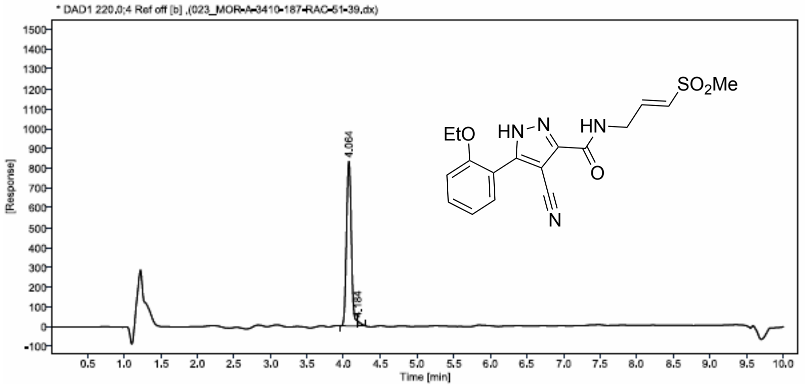


| Sr.No. | RT | Area | Height | % Area |
|---|---|---|---|---|
| 1 | 4,06 | 3792 | 831 | 98,33 |
| 2 | 4,18 | 64 | 24 | 1,67 |

**Figure S194**: HPLC trace of **24a**

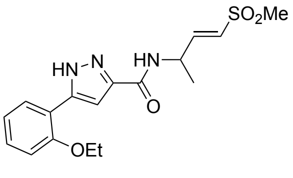


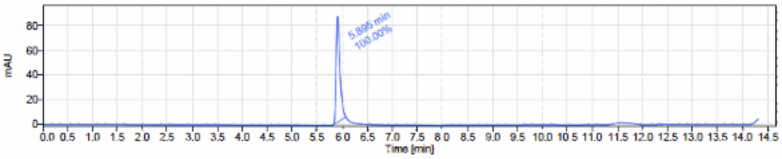


Signal Name DAD1A

| RT (min) | Signal description | m/z | Purity UV(‰) / MS(%) | Area | Area% |
|---|---|---|---|---|---|
| 5.895 | * DAD1A,Sig=254.0,5.0 Ref=360.0,80.0 [b] | | | 454.4 | 100.00 |

**Figure S195**: HPLC trace of **24b**

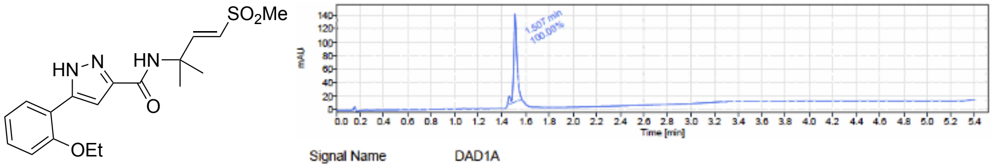


| Signal Name | DAD1A | | | | |
|---|---|---|---|---|---|
| RT (min) | Signal description | m/z | Purity UV(‰) / MS(%) | Area | Area% |
| 1.507 | DAD1A,Sig=254.0,5.0 Ref=360.0,80.0 | | | 258.8 | 100.00 |

**Figure S196**: HPLC trace of **24c**

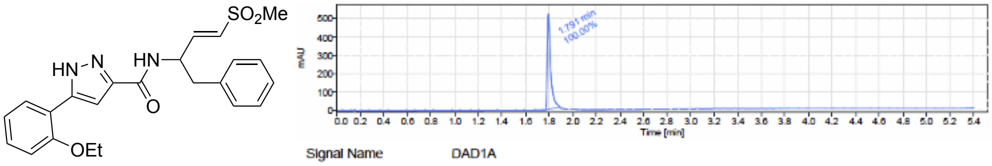


| Signal Name | DAD1A | | | | |
|---|---|---|---|---|---|
| RT (min) | Signal description | m/z | Purity UV(‰) / MS(%) | Area | Area% |
| 1.791 | DAD1A,Sig=254.0,5.0 Ref=360.0,80.0 | | | 1014.0 | 100.00 |

**Figure S197**: HPLC trace of **24d**

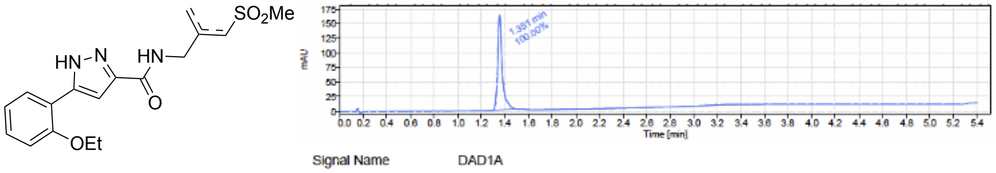


| Signal Name | DAD1A | | | | |
|---|---|---|---|---|---|
| RT (min) | Signal description | m/z | Purity UV(‰) / MS(%) | Area | Area% |
| 1.351 | DAD1A,Sig=254.0,5.0 Ref=360.0,80.0 | | | 432.6 | 100.00 |

**Figure S198**: HPLC trace of **24e**

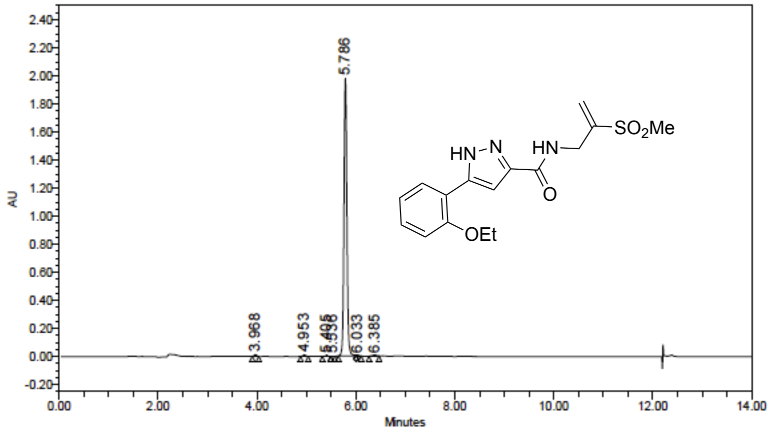


Peak Results

| | Name | RT | Height (µV) | Area (µV*sec) | % Area |
|---|---|---|---|---|---|
| 1 | | 3.968 | 12629 | 43316 | 0.52 |
| 2 | | 4.953 | 11227 | 40908 | 0.49 |
| 3 | | 5.405 | 5007 | 18787 | 0.23 |
| 4 | | 5.536 | 719 | 1895 | 0.02 |
| 5 | | 5.786 | 1979110 | 8113963 | 97.85 |
| 6 | | 6.033 | 7091 | 27342 | 0.33 |
| 7 | | 6.385 | 7974 | 46424 | 0.56 |

**Figure S199**: HPLC trace of **24f**

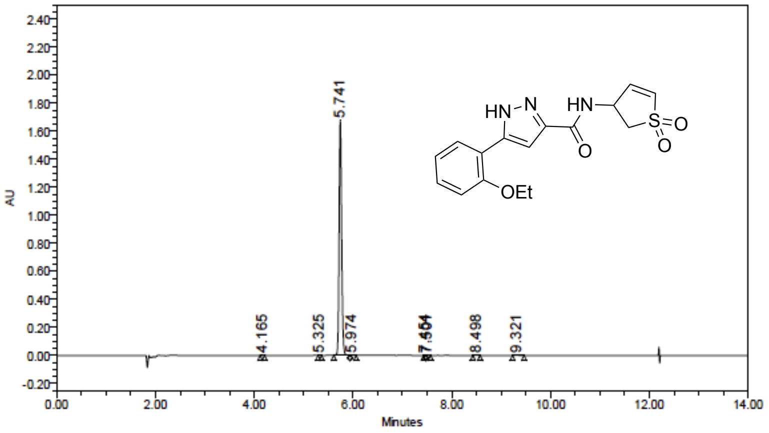

Peak Results

| | Name | RT | Height (µV) | Area (µV*sec) | % Area |
|---|---|---|---|---|---|
| 1 | | 4.165 | 1627 | 4408 | 0.07 |
| 2 | | 5.325 | 605 | 1471 | 0.02 |
| 3 | | 5.741 | 1678516 | 5999346 | 99.74 |
| 4 | | 5.974 | 1001 | 3939 | 0.07 |
| 5 | | 7.454 | 200 | 416 | 0.01 |
| 6 | | 7.501 | 292 | 858 | 0.01 |
| 7 | | 8.498 | 602 | 2221 | 0.04 |
| 8 | | 9.321 | 418 | 2103 | 0.03 |

**Figure S200**: HPLC trace of **25a**

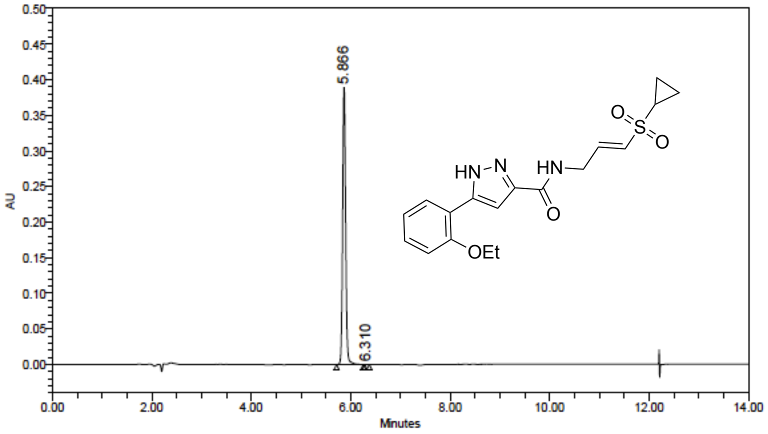

Peak Results

| | Name | RT | Height (µV) | Area (µV*sec) | % Area |
|---|---|---|---|---|---|
| 1 | | 5.866 | 389753 | 1626082 | 99.99 |
| 2 | | 6.310 | 67 | 231 | 0.01 |

**Figure S201**: HPLC trace of **25b**

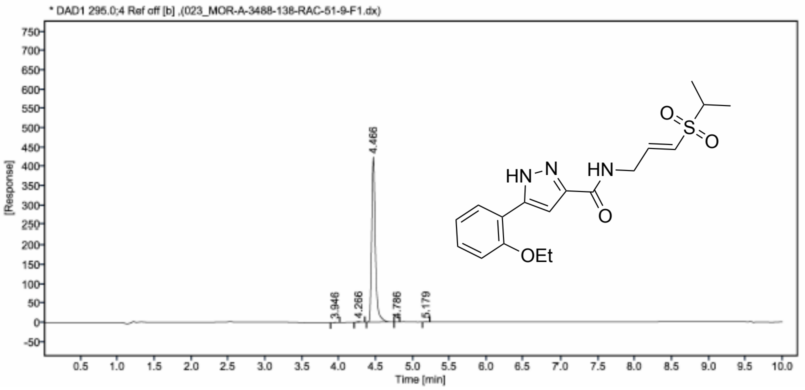


| Sr.No. | RT | Area | Height | % Area |
|---|---|---|---|---|
| 1 | 3.95 | 2 | 1 | 0.16 |
| 2 | 4.27 | 6 | 2 | 0.39 |
| 3 | 4.47 | 1486 | 424 | 99.25 |
| 4 | 4.79 | 1 | 0 | 0.07 |
| 5 | 5.18 | 2 | 1 | 0.12 |

**Figure S202**: HPLC trace of **25c**

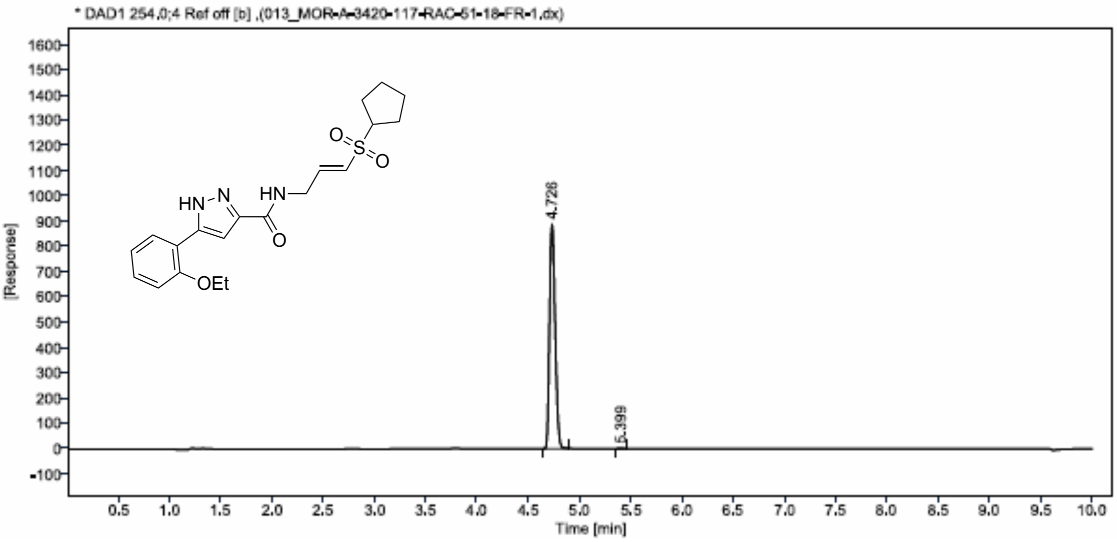


| Sr.No. | RT | Area | Height | % Area |
|---|---|---|---|---|
| 1 | 4,73 | 3383 | 889 | 99,81 |
| 2 | 5,40 | 7 | 2 | 0,19 |

**Figure S203**: HPLC trace of **25d**

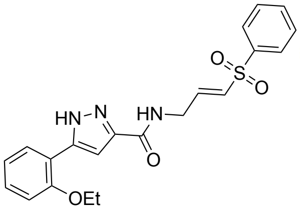


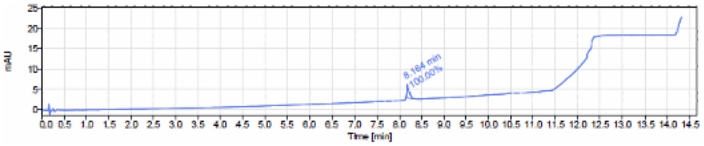


Signal Name DAD1A

| RT (min) | Signal description | m/z | Purity UV(%) / MS(%) | Area | Area% |
|---|---|---|---|---|---|
| 8.164 | DAD1A,Sig=254.0,5.0 Ref=360.0,80.0 | | | 11.4 | 100.00 |

**Figure S204**: HPLC trace of **25e**

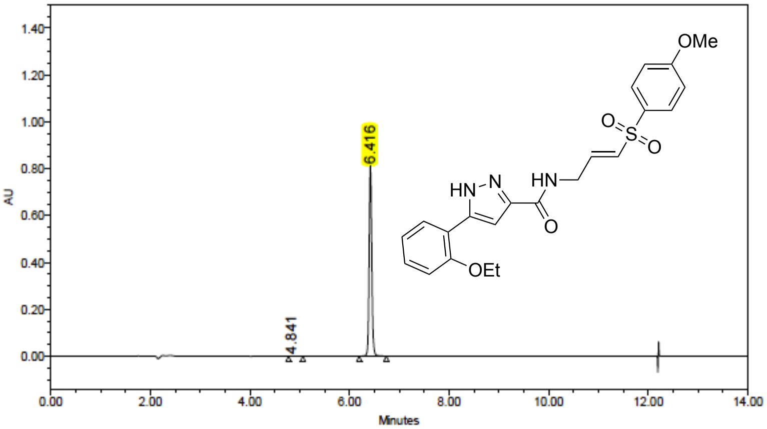


Peak Results

| | Name | RT | Height (µV) | Area (µV*sec) | % Area |
|---|---|---|---|---|---|
| 1 | | 4.841 | 107 | 925 | 0.03 |
| 2 | | 6.416 | 810051 | 3004935 | 99.97 |

**Figure S205**: HPLC trace of **25f**

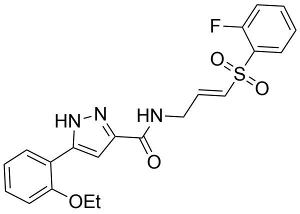


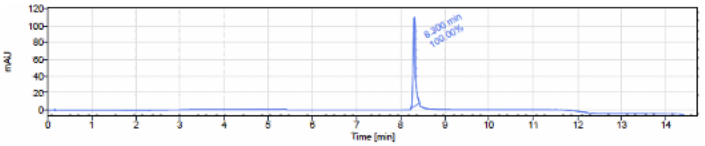


Signal Name DAD1A

| RT (min) | Signal description | m/z | Purity UV(%) / MS(%) | Area | Area% |
|---|---|---|---|---|---|
| 8.300 | * DAD1A,Sig=254.0,5.0 Ref=360.0,80.0 [b] | | | 399.6 | 100.00 |

**Figure S206**: HPLC trace of **25g**

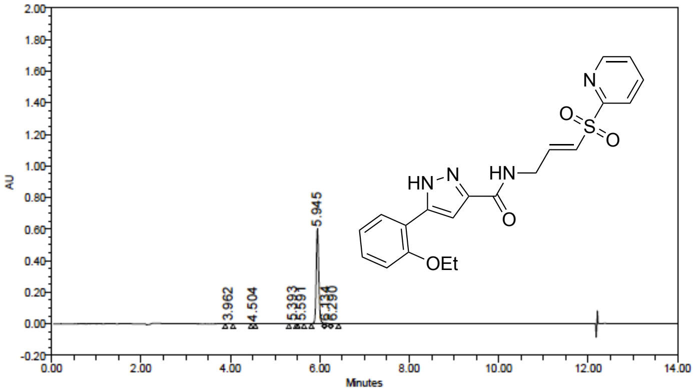


Peak Results

| | Name | RT | Height (µV) | Area (µV*sec) | % Area |
|---|---|---|---|---|---|
| 1 | | 3.962 | 2116 | 7405 | 0.31 |
| 2 | | 4.504 | 386 | 992 | 0.04 |
| 3 | | 5.393 | 813 | 2776 | 0.12 |
| 4 | | 5.591 | 693 | 2328 | 0.10 |
| 5 | | 5.945 | 599754 | 2359373 | 98.53 |
| 6 | | 6.134 | 3328 | 16267 | 0.68 |
| 7 | | 6.290 | 1024 | 5466 | 0.23 |

**Figure S207**: Chiral SFC trace of (*R*)-**24a**

| Sample | Sample Well | Inj. Vol.(ul) |
|---|---|---|
| MOR-A-3543-191-RAC-51-99-FR-01 | 2,5,A | 12 |

| File Name | Instrument Method | Inj. Time |
|---|---|---|
| MOR-A-3543-191-RAC-51-99-FR-01_1.tta | 04_SFC_05_100 | 11-03-2024 17:40:13 |

| Flow(ml/min) | Co-Solvent% Values | Back Pressure | Col. Temp. (C) | Report Date |
|---|---|---|---|---|
| 4.00 | 5.0 | 100 Bar | 40 | 11-03-2024 |
| 4.00 | 50.0 | | | |
| 4.00 | 50.0 | | | |

| Column | Co-Solvent |
|---|---|
| CHIRALPAK IH (250*4.6mm, 5um) | 0.1%NH3 IN IPA:ACN (50:50) |

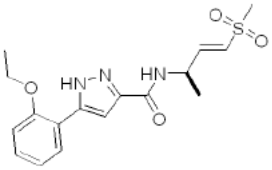


Molecular Weight: 363.43

Single Absorbance (250nm) Plot

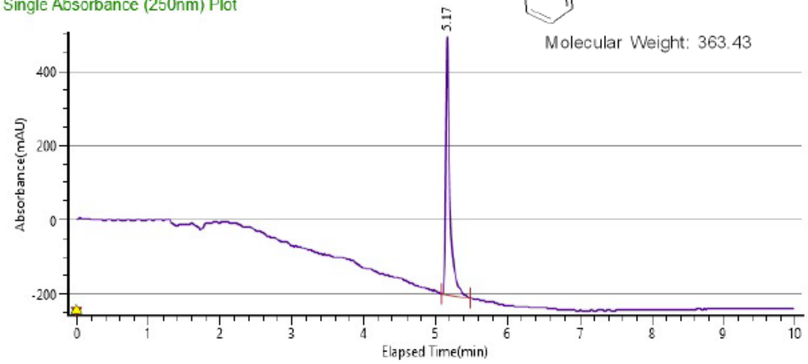


**Peak Info[(W2998)SingleAbsorbance250]**

| Peak # | Ret. Time | Area | Height | Area % |
|---|---|---|---|---|
| 1 | 5.17 min | 2738.3965 | 693.9703 | 100.0000 |

**Figure S208**: Chiral SFC trace of (*S*)-**24a**

| Sample | Sample Well | Inj. Vol.(ul) |
|---|---|---|
| MOR-A-3543-192-RAC-51-99-FR-02 | 2,6,A | 12 |

| File Name | Instrument Method | Inj. Time |
|---|---|---|
| MOR-A-3543-192-RAC-51-99-FR-02_1.tta | 04_SFC_05_100 | 11-03-2024 17:53:23 |

| Flow(ml/min) | Co-Solvent% Values |
|---|---|
| 4.00 | 5.0 |
| 4.00 | 50.0 |
| 4.00 | 50.0 |

| Back Pressure | Col. Temp. (C) | Report Date |
|---|---|---|
| 99 Bar | 40 | 11-03-2024 |

| Column | Co-Solvent |
|---|---|
| CHIRALPAK IH (250*4.6mm, 5um) | 0.1%NH3 IN IPA:ACN (50:50) |

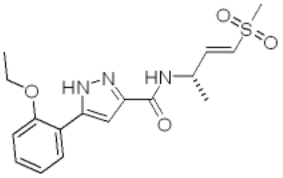

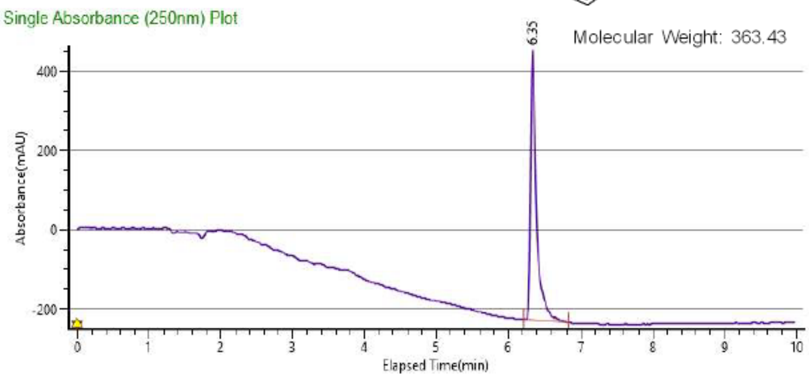


**Peak Info[(W2998)SingleAbsorbance250]**

| Peak # | Ret. Time | Area | Height | Area % |
|---|---|---|---|---|
| 1 | 6.35 min | 3880.3869 | 680.3875 | 100.0000 |
